# Higher-order epistasis creates idiosyncrasy, confounding predictions in protein evolution

**DOI:** 10.1101/2022.09.07.505194

**Authors:** Karol Buda, Charlotte M. Miton, Nobuhiko Tokuriki

## Abstract

Epistasis shapes evolutionary outcomes during protein adaptation. In particular, when the effects of single mutations or mutational interactions are idiosyncratic, that is, unique to a genetic background, the predictability of protein evolution becomes greatly impaired. Here, we unveil a quantitative picture of the prevalence and role of idiosyncrasy in protein evolution by analysing 45 protein fitness landscapes, generated from seven enzymes. We found that mutational effects and epistasis are highly idiosyncratic across the landscapes. Idiosyncrasy obscured functional predictions of mutated proteins when using limited mutational data, and often continued to impair prediction upon incorporation of epistatic information. We show that idiosyncrasy stems from higher-order epistasis, and highlight examples where it permits, or restricts, evolutionary accessibility of certain genotypes. Our work suggests that idiosyncrasy deeply confounds predictions in protein evolution necessitating its incorporation into predictive models and in-depth exploration of its underlying molecular mechanisms.

## Introduction

The evolution of new protein functions often requires the gradual accumulation of adaptive mutations. Accordingly, the fixation of multiple mutations along an evolutionary trajectory may cause a rewiring of intramolecular amino acid interaction networks that results in non-additive mutational effects, or epistasis^1–3^. Epistasis causes a deviation from smooth, additive, and predictable mutational behaviours that – when rendered on a fitness landscape – creates a rugged mountain range with jagged peaks and valleys^4–6^. As evolution proceeds *via* the stepwise accumulation of single mutations, a rugged landscape topography dictates the ‘opening’ or ‘closing’ of accessible mutational paths across the landscape towards a given fitness peak^7,8^. Thus, understanding, let alone predicting, protein evolution requires a robust description of epistasis and its ability to distort the sequence-function relationship^4^.

One approach to survey intramolecular epistasis in proteins relies on the functional characterisation of genotypes encompassing all possible combinations of a mutational subset that collectively alter protein function^9–11^. Such investigations have demonstrated that there are a limited number of accessible mutational paths between the starting and endpoint genotypes, highlighting the evolutionary constraints exerted by epistasis^11–17^. Recently, novel statistical approaches have been developed to obtain a more quantitative picture of epistasis, including higher-order epistasis (the non-additive effects of three or more mutations), embedded in these landscapes^12–19^. These statistical analyses typically provide a ‘global’ view of epistasis, where the averages of mutational and epistatic effects are captured across the combinatorial space^12,15,17^. Such global approaches have demonstrated that the overall evolutionary changes within a combinatorial fitness landscape can be statistically recapitulated using the average effects of all individual mutations, supplemented only by the epistasis between pairwise and triplet mutations^15,17^.

The global view of fitness landscapes can, however, have substantial shortcomings in scenarios where the effects of single mutations and epistasis are highly variable across different genotypes. For instance, if the effect of a mutation is idiosyncratic – *i*.*e*., showing diverse functional contributions depending on the background genotype – its function in each genotype will significantly deviate from its global effect across the entire landscape^20,21^. Since the majority of available mutational datasets characterise mutational effects in only a few backgrounds (generally single and double mutants in the protein’s wild-type (WT) background) most predictions made in protein evolutionary studies and engineering campaigns have the propensity to be heavily skewed by idiosyncrasy. Indeed, if the accessibility of a genotype is opened up or closed off by a strong, highly idiosyncratic, mutational effect, which is only present in a single (or very few) genotype(s) in the landscape, this idiosyncrasy will simultaneously dictate the evolutionary path followed by the protein whilst being lost in the global view, as it cannot be attributed to any epistasis captured by global effects. Hence, ubiquitous idiosyncrasy has the potential to confound our predictions and our understanding of evolutionary outcomes. However, we have little knowledge regarding the prevalence and roles of idiosyncrasy in protein fitness landscapes. If prevalent, what causes idiosyncrasy? To what extent does idiosyncrasy impact predictions in protein evolution?

In this study, we address these questions by systematically characterising 45 functionally annotated, combinatorially complete (for a subset of mutations), mutational landscapes. We explored the levels of idiosyncrasy across protein fitness landscapes and explore its impact on the accessibility of adaptive trajectories (Fig. 1). We first inferred idiosyncrasy from the extent of heterogeneity in single mutational effects and epistasis in these datasets. We then quantified the levels of WT idiosyncrasy by comparing the magnitude of every single mutation and epistatic contribution in the WT genotype to their average across the landscape. Armed with a global and local statistical model, we unveiled the impact of idiosyncrasy on our ability to predict evolutionary-intermediates and endpoints across various fitness landscapes. Finally, we examined how idiosyncrasy affects the accessibility of a given evolutionary trajectory by permitting or restricting certain paths, and how these patterns translate to the molecular level.

**Fig. 1.**
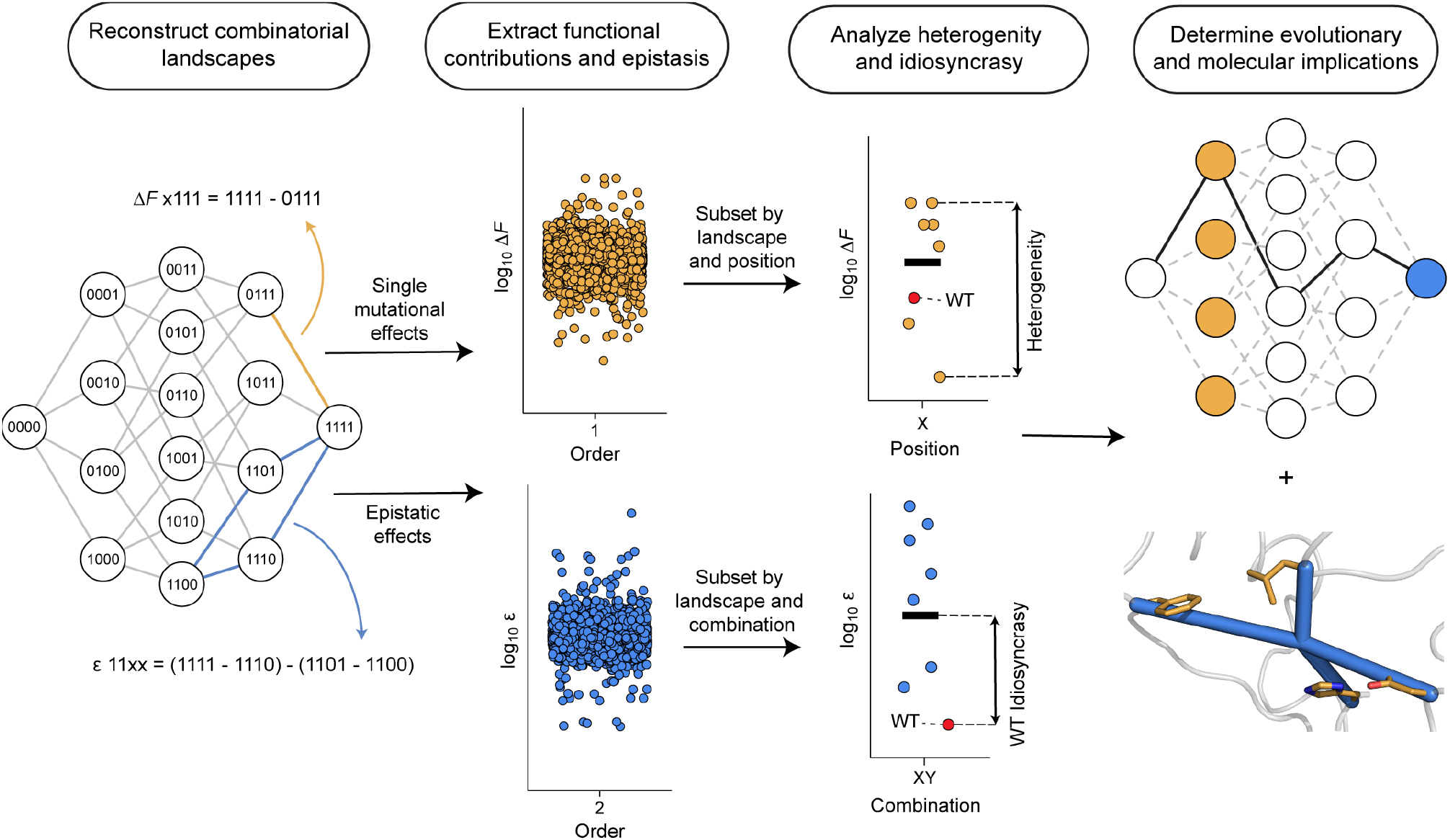
Analytical pipeline for characterising idiosyncrasy in protein combinatorial landscapes. From left to right, the data from empirical studies were processed to reconstruct combinatorial landscapes; in genotypes, 0 and 1 represent ancestral (WT) and derived (mutated) positions, respectively. We extracted the functional effect of mutations (Δ*F*) for each position (< 2 mutations), and epistasis (ε) for each combination (≥ 2 mutations). Heterogeneity and WT idiosyncrasy for single mutational- and epistatic-effects were captured using a statistical metric of spread and WT deviation from the mean (black bars), respectively. The impact of idiosyncrasy on evolutionary trajectories was analyzed and translated to the structural and molecular levels, *e*.*g*., single mutational (yellow) and epistatic (blue) data can be visualized on combinatorial landscapes and a protein structure to shed light on their evolutionary relevance.

## Results

### Statistical characterisation of 45 combinatorial landscapes

We collected several experimental studies that characterised the changes in protein function for a set of combinatorial mutations along an adaptive trajectory. The analyzed fitness landscapes – herein referred to as combinational landscapes – were restricted to datasets probing single mutations per position. These studies were filtered to ensure that the landscapes explored four or more positions (n ≥ 4) and functionally characterised all possible combinations of these mutations (2^n^ variants). Of those studies, we only retained those reporting continuous data. Using these cut-offs, we obtained a working set of ten studies exploring seven different enzymes (Table 1). For some studies, the mutations were accumulated through directed evolution or protein engineering of a novel function (phosphotriesterase, PTE; β-lactamase, OXA-48; nitroreductase, NfsA)^13,22–24^. For the remaining studies, the positions were identified from naturally occurring evolutionary trajectories, either through a retrospectively identified path using ancestral sequence reconstruction (methyl-parathion hydrolase, MPH)^14,25^, the presence of clinically relevant mutations (dihydrofolate reductase, DHFR and β-lactamase, TEM-1)^11,26–28^, or in the case of alkaline phosphatase (AP), by using previously characterised active site residue mutations^29^. The final dataset consisted of 56 unique mutations, of which 54 are located within the protein open reading frame, and two are located in a promoter region (in DHFR and TEM-1)^11,23^. These were analyzed in 45 separate combinatorial landscapes with a total of 1,504 genotype-phenotype data points, some of which explored a set of positions for the same enzyme using a different substrate, inhibitor, or metal cofactor (Table 1 and Supplementary File 1).

**Table 1.** Combinatorial landscapes analyzed in this study.

| Enzyme | No. of mutations | Conditions | No. of total measurements | Measured trait | Reference |
| --- | --- | --- | --- | --- | --- |
| OXA-48<sup>a</sup> | 4, 6, 6 | 2, 2, 2 | 288 | $IC_{50}$ | 22 |
| TEM-1<sup>b</sup> | 5, 4 | 1, 15 | 272 | MIC, Growth rate | 11,28 |
| AP | 5 | 1 | 32 | $k_{cat}/K_M$ | 29 |
| NfsA<sup>c</sup> | 7, 7 | 1, 1 | 256 | $EC_{50}$ | 24 |
| DHFR<sup>d</sup> | 4, 6, 5, 5 | 5, 1, 2, 2 | 272 | $IC_{50}$, $IC_{75}$, $k_{cat}/K_M$ and $K_i$ | 23,26,27 |
| MPH<sup>e</sup> | 5 | 8 | 256 | Lysate activity | 25 |
| PTE<sup>f</sup> | 6 | 2 | 128 | Lysate activity | 13 |

<sup>a</sup> Three mutationally unique trajectories, where each was probed using two inhibitors
<sup>b</sup> Ref. <sup>28</sup> explored 15 inhibitors for the same set of four mutations
<sup>c</sup> Two separate mutational trajectories for the same protein and substrate
<sup>d</sup> Ref. <sup>26</sup> explored four mutations using five different substrates; ref. <sup>27</sup> explored both $k_{cat}/K_M$ and $K_i$ for two mutational trajectories
<sup>e</sup> Ref. <sup>25</sup> explored the same five mutations under eight different metal environments
<sup>f</sup> Six mutations were explored using two substrates, one in ref. <sup>13</sup> and one outlined in this study (see **Methods**)

Next, the mutational data were processed to allow for a streamlined analysis using our computational pipeline (see Code Availability). Trajectories that explored different mutational combinations for the same enzyme were treated as separate combinatorial landscapes, as were the combinatorial landscapes characterising the function of the same subset of mutants across different substrates, ligands, or metals. Due to the variety of measured functions, ranging from direct physicochemical properties of the enzymes to indirect effects on the cellular phenotype, all enzyme functions were normalized relative to their WT background, providing us with the fold-change in enzyme function (*F*), which was then log_10_ transformed (see Methods). Not accounting for the WT genotypes where *F* = 0, we obtained 1,459 *F* values for further analysis (Supplementary Data 1).

### Idiosyncrasy in single mutational effects

To paint a comprehensive picture of the prevalence of epistasis in the selected proteins, we first extracted the functional effect of every single mutation at a given position across all available genetic backgrounds. The data were processed to provide the change in function (Δ*F*), or single mutational effect, of a given mutation across every genotype (see Methods). In a combinatorially complete fitness landscape of *n* mutations, a particular mutation occurs in 2^n-1^ distinct genetic backgrounds, hence, each combinatorial landscape contains *n* × 2^n-1^ Δ*F* values. Using this approach, we collected the 4,064 Δ*F* (Fig. 2a) and faceted them by the 214 unique mutation-substrate/inhibitor/metal pairs, simply referred to as ‘positions’, *e*.*g*., the effect of a mutation at the same amino acid position for two different substrates is treated as a different ‘position’ (Supplementary Data 2). We chose to use a significance threshold of 1.5-fold for all analyses; this was the median error rate (calculated as 2 × standard deviation, or 2*σ*) for all replicate measurements available in our data. Using this threshold, we found that the signs of the Δ*F* values were 21% negative, 36% neutral, and 43% positive across all genotypes. This constitutes a relatively even split for the sign of the single mutational effects, despite a slight bias toward a positive effect, consistent with the fact that most mutations were originally identified due to their beneficial effect on function (Fig. 2a).

**Fig. 2.**
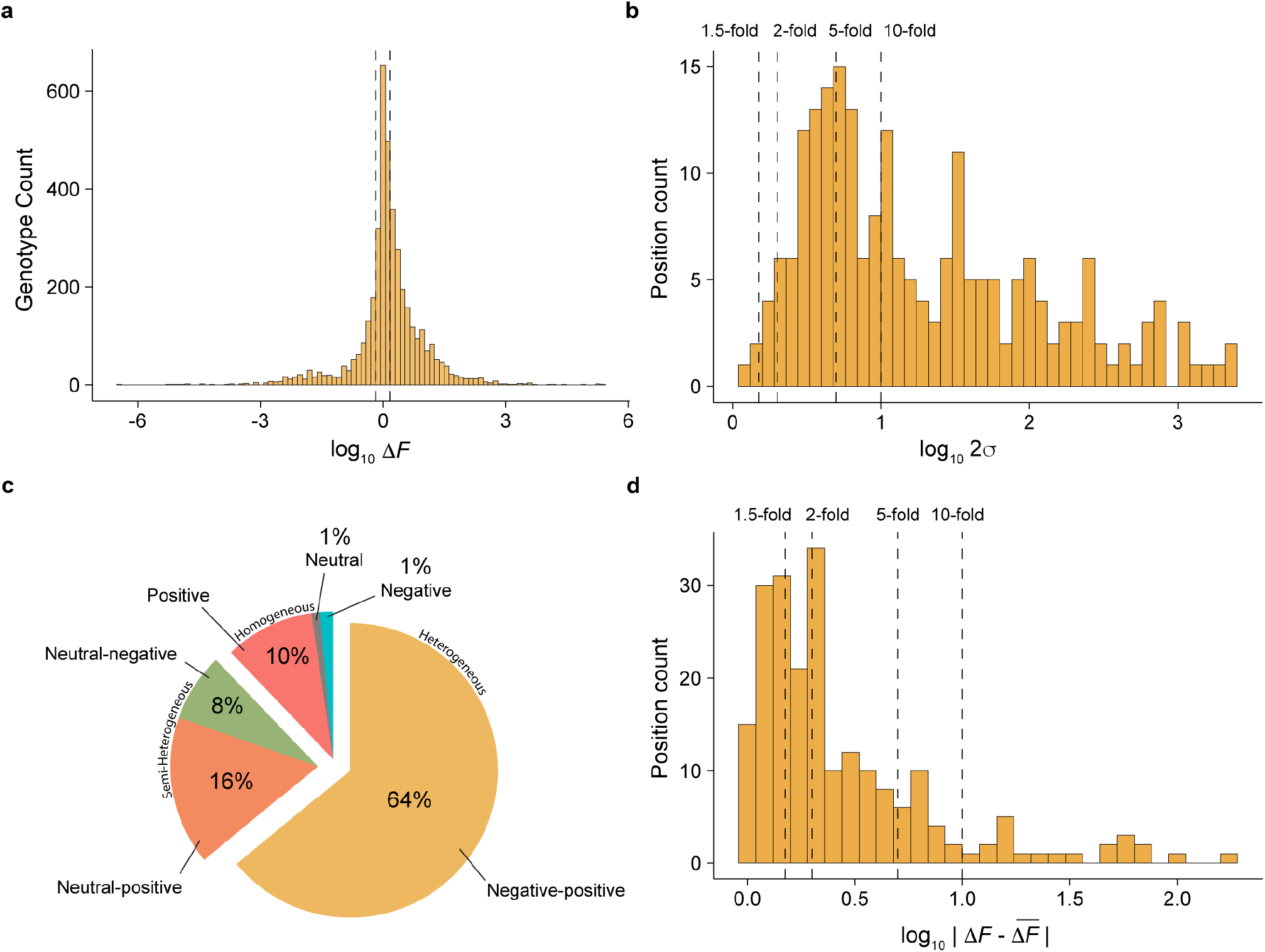
Descriptive statistics of single mutational effects. **a**, Distribution of single mutational effects (log_10_ Δ*F*) at every position across all genotypes for every combinatorial landscape. Dashed lines represent the 1.5-fold significance threshold that distinguishes negative, neutral, and positive effects, respectively. **b**, Distribution of the heterogeneity of the single mutational effects (log_10_ 2σ for the distribution of Δ*F*s) at each of the 214 positions. Dashed lines represent 1.5-, 2-, 5-, and 10-fold significance thresholds, respectively. **c**, Categorisation of the sign-changing behavior in Δ*F* distribution for each of the 214 mutations. **d**, Distribution of the WT idiosyncrasy quantified by the deviation of each of the 214 Δ*F*_wt_ from 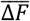 at each position. Dashed lines representing 1.5-, 2-, 5-, and 10-fold significance thresholds, respectively.

Next, we characterised the variability in single mutational effects for a given position, herein referred to as heterogeneity, by plotting and analysing the spread in the distribution of Δ*F* at each position (Supplementary File 2). The heterogeneity in Δ*F* across different genetic backgrounds demonstrates the presence of extensive idiosyncrasy; accordingly, the degree of Δ*F* scatter for a position across various genotypes reflects the existence of epistasis of pairwise, and likely higher-order, interactions. To capture this spread, we computed 2*σ* for the Δ*F* at each position, which should encompass ∼95% of the heterogeneity and inform on the spread around the mean single mutational effect for a given position. We deemed positions to be significantly heterogeneous when their 2*σ* > 1.5-fold. We found that heterogeneity prevails in our dataset: 99% (211/214) of positions exhibit significant heterogeneity (Fig. 2b and Extended Data Table 2). Many positions exhibit much higher heterogeneity than the threshold: 70% (150/214) show a 2*σ* >5-fold, and 53% (114/214) show a 2*σ* >10-fold (Fig. 2b and Extended Data Table 2). We also characterised the extent of sign heterogeneity, *i*.*e*., whether the sign of the single mutational effect varies between positive, negative, and/or neutral in different genotypic backgrounds. Only 12% of the positions showed sign homogeneity (retaining the same sign in all genotypes) and the remaining positions showed at least moderate sign heterogeneity, with 8% of positions contributing either a neutral or negative effect and 16% of positions displaying either a neutral or positive effect. Interestingly, 64% of positions showed full sign heterogeneity, showing background-dependent variance between positive and negative effects (Fig. 2c and Extended Data Table 2). These observations suggest that single mutational effects in the combinatorial landscapes are highly idiosyncratic – likely a result of pervasive epistasis.

Highly heterogeneous Δ*F* also suggest that the effect of a mutation can be idiosyncratic in a particular background compared to its global effect in the combinatorial landscape. We computed the difference in magnitude and sign of Δ*F* for a given mutation in the WT background (Δ*F*_wt_) *versus* the mean 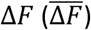 of that mutation across all representative genotypes to determine the levels of WT idiosyncrasy (Fig. 2d). For 68% of positions, Δ*F*_wt_ deviates from 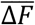 by >1.5-fold (Fig. 2d and Extended Data Table 3), while the sign of Δ*F*_*wt*_ remains similar to that 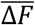 (only 3% of positions show a significant difference in sign). The idiosyncrasy in single mutational effects does not only apply to Δ*F*_wt_, but also to the Δ*F* in other genotypes – 59% (2396/4064) of single mutants across every genotypic background also deviated >1.5-fold from their 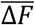 (Extended Data Table 4). This suggests that the Δ*F*_wt_, or Δ*F* of any other genotypic background, is inaccurately captured by the global effect of that mutation within the combinatorial landscape. In other words, the magnitude of a previously observed single mutational effect is likely to vary in new genotypic backgrounds due to idiosyncrasy.

### Idiosyncrasy in pairwise- and higher-order epistasis

The extensive idiosyncrasy in Δ*F* essentially stems from epistasis, therefore, we expanded our survey of idiosyncrasy in single mutational effects to interactions between mutations. Epistasis (ε), at any order, can be mathematically defined as the additional functional contribution originating from a new mutational interaction relative to all constituent lower-order interactions and single mutational effects. For example, our calculation of ε at the 3^rd^ order (and higher orders) accounts for ε observed in the 2^nd^ order, in contrast to the traditional calculation of the difference between observed function and predicted function using only the sum of the single mutational effects. Thus, we used the following equation to calculate 2^nd^ order epistasis between mutations:

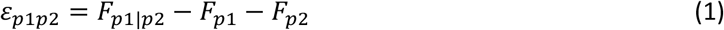

Where p1 and p2 represent the single mutations at hypothetical positions ‘1’ and ‘2’, respectively, p1|p2 represents the combination of mutations p1 and p2, ε is the epistasis coefficient, and *F* is the normalized function. Higher-order epistasis was calculated using a similar approach as in equation 1 (see Methods). The extracted ε coefficients for all combinations across all landscapes were probed using the same metrics applied to the Δ*F*, to capture the extent of idiosyncrasy in epistasis.

Compared to the 214 surveyed positions, we extracted 419 pairwise-, 438 triple-, and 267 quadruple-combinations. Since a few landscapes only covered 4 mutations, 22/267 quadruple combinations could not be examined as they only occurred in one genetic background, leaving 245 quadruple combinations viable for analysis. Ultimately, we collected 8968 values for ε encompassing the 2^nd^ to the 4^th^ order (Supplementary Data 3). Surprisingly, we found that nearly all (99% (413/419)) pairwise interactions are significantly heterogeneous (Fig. 3a). This proportion decreased at less stringent thresholds, nonetheless, even with a 10-fold significance cut-off, 41% of pairwise interactions remained significantly heterogeneous (Extended Data Table 2). Interestingly, this trend remains consistent at higher orders of epistasis: 96% (420/438) of triple interactions, and 98% (240/245) of the quadruple interactions exhibit significant heterogeneity (Fig. 3a). The impact of high heterogeneity in epistasis was also seen in their sign variability – for pairwise combinations, 71% (295/419) show full sign heterogeneity (Fig. 3b). Only a handful of them (5%) exhibited sign homogeneity, and the remaining combinations were either neutral-negative (9%) or neutral-positive (15%). For higher-order combinations, sign heterogeneity remains high: 54% for 3^rd^ order, and 52% for 4^th^ order combinations (Fig. 3b and Extended Data Table 2).

**Fig. 3.**
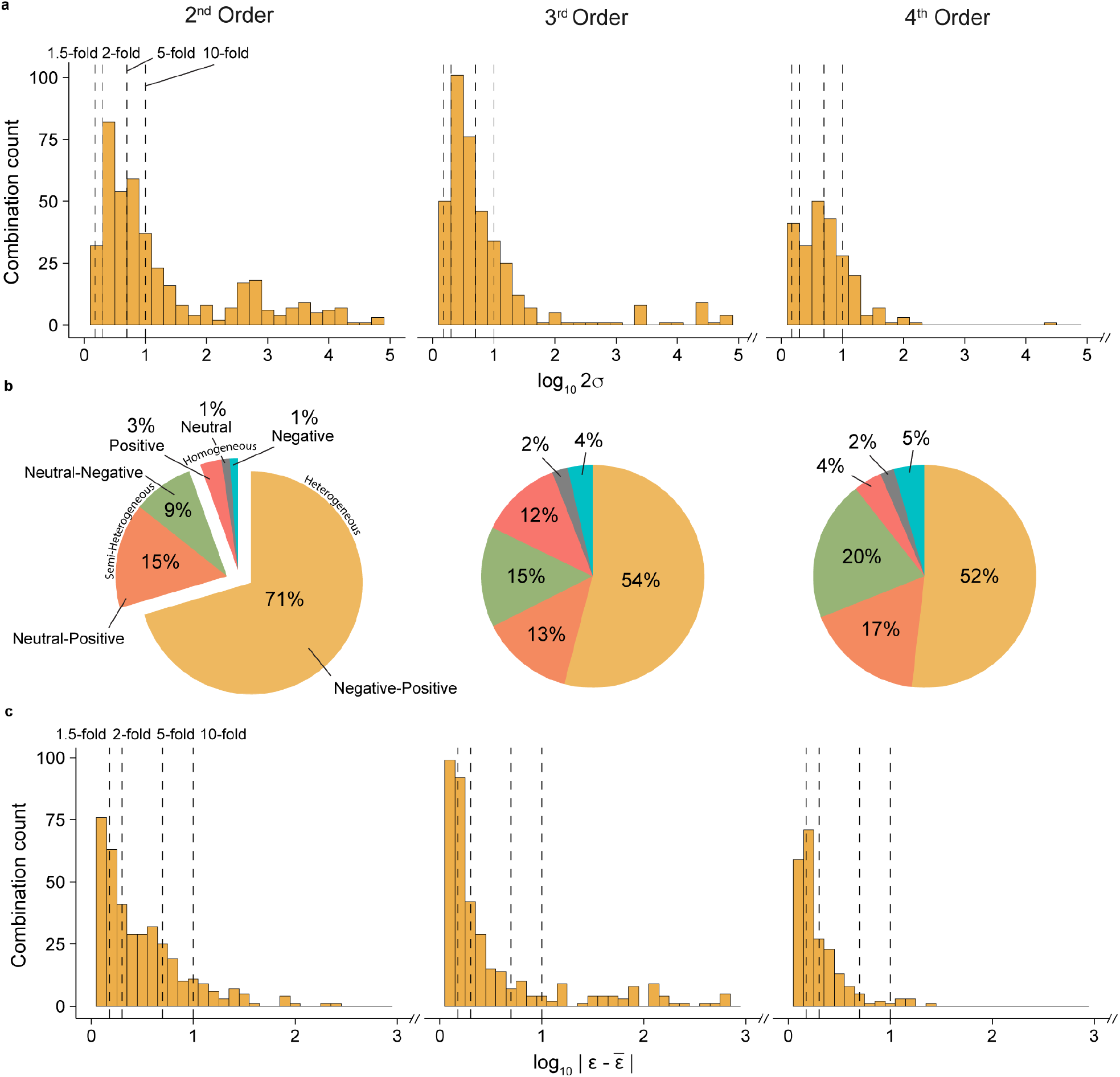
Descriptive statistics of epistasis. **a**, Histogram of log_10_ 2σ of ε at each order for every mutational combination, with annotated dashed lines representing 1.5-fold, 2-fold, 5-fold, and 10-fold significance thresholds, respectively. Data omitted due to x-axis scaling can be found in **Extended Data Fig. 1a. b**, Categorisation of the ε at each order for all mutational combinations by sign. **c**, The distribution of the WT idiosyncrasy in epistasis quantified *via* the absolute difference of ε_*wt*_ from 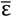 at each combination. Dashed lines represent 1.5-, 2-, 5-, and 10-fold significance thresholds, respectively. Data omitted due to x-axis scaling can be found in **Extended Data Fig. 1b**.

Like single mutational effects, the WT background epistasis (ε_wt_) showed a high deviation from the mean 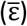 for each combination. More than half (64% (270/419)) of pairwise, (56% (244/438)) of triplet, and (62% (153/245)) of quadruplet ε_wt_ deviated from 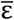 by more than 1.5-fold (Fig. 3c). As with the Δ*F*, this high deviation was also representative of other non-WT backgrounds (Extended Data Table 4). Interestingly, more combinations showed significant sign discrepancy in ε_wt_ *versus* 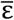 than those for Δ*F*. Although still in the minority, 13% (53/419) of pairwise-, 5% (34/438) of triplet-, and 8% (12/245) of quadruplet-interactions had a significantly different sign effect in the ε_wt_ than the 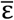. These findings suggest that epistasis itself remains idiosyncratic, even at increasing orders. The persistence of idiosyncrasy at the 3^rd^ and 4^th^ orders of epistasis suggests that epistasis at the 5^th^ (and likely higher) order(s) continues to impose a genotypic context dependence even on triplet and quadruplet mutational interactions.

### Idiosyncrasy confounds predictability

Given the strong prevalence of idiosyncrasy in single mutational effects and epistasis, we asked to what extent does idiosyncrasy impair the predictability of genotypic functions in the landscape? We computed functional predictions using two different models and examined the discrepancies between them. The first model collected the Δ*F*_wt_ and ε_wt_ for every position and combination and computed the function at every genotype using the additive assumption of that genotype’s constituent Δ*F* and ε values (Fig. 4a and Methods). We refer to this as the ‘WT-background model’, because the model relies on limited data from the WT background, namely Δ*F*_wt_ and ε_wt_, as opposed to any other, or all, Δ*F* and ε across the landscape (Fig. 4b). This model is inspired by the local view of epistasis^16,17^, which stems from the stepwise process employed in protein engineering and evolutionary studies. We compared the WT-background model to a global model that used 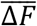 and 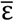 by averaging Δ*F* and ε values across the landscape this model aims to minimize the idiosyncratic noise of the system. Our global model of choice was a linear regression with interaction terms, simply referred to as the global model, which is conventionally employed in combinatorial landscape analyses (Fig. 4a)^11,13,14,16,17,25^. This comparison aims at exposing how a deviation between mutational- and epistatic-effects in WT background *versus* their corresponding mean effects can confound predictions of evolutionary outcomes.

**Fig. 4.**
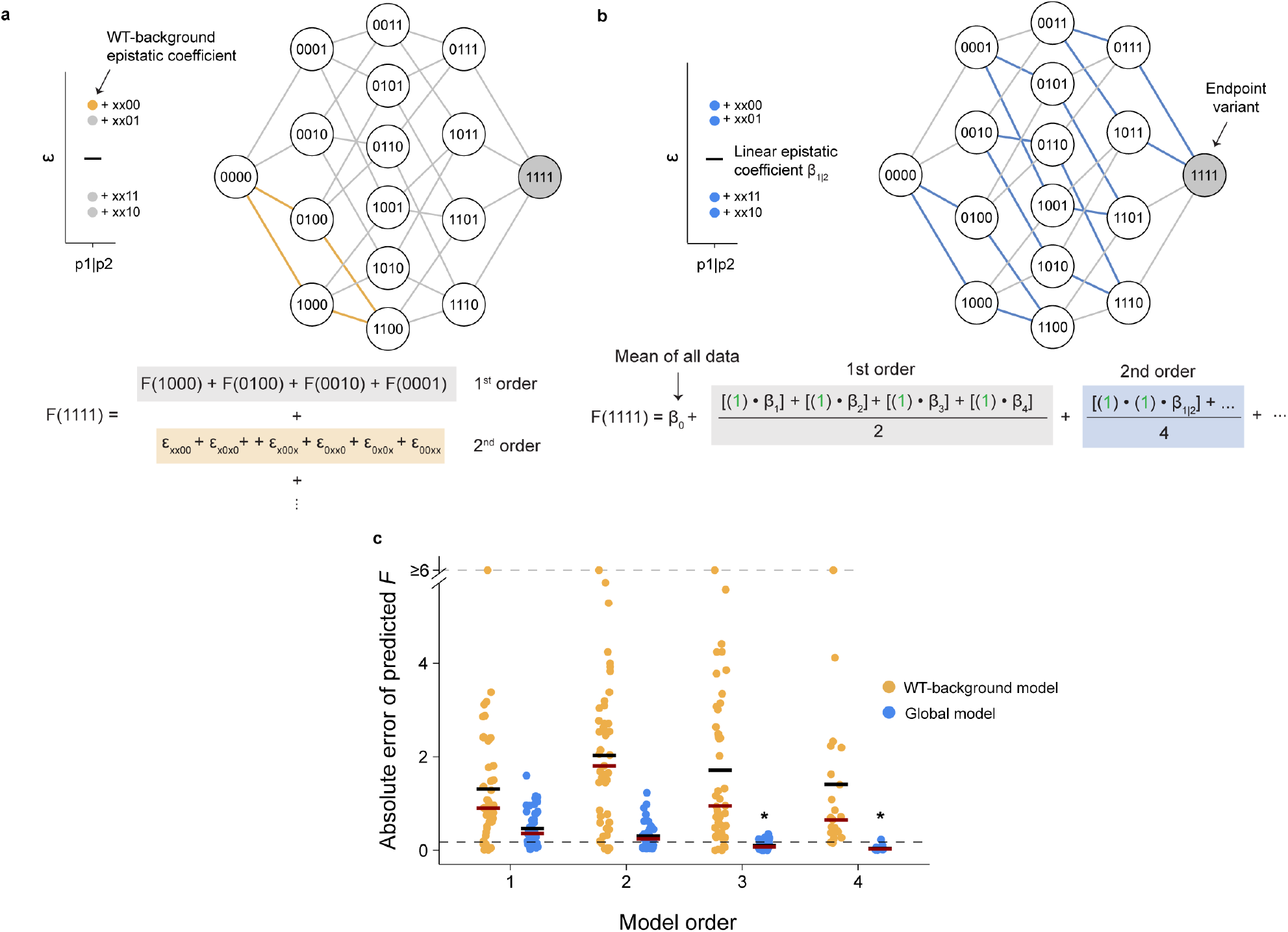
Differences in WT-background and global models. **a-b**, Predicting the function of an endpoint variant in a hypothetical landscape using the WT-background (**a**) and global (**b**) model. **a**, The WT-background only uses epistasis in the WT background to extract the epistatic coefficient (ε_xx00_) and uses it to predict the function (equation below and **Methods**). **b**, The global model computes epistatic coefficients for an interaction, here for positions 1 and 2 (β_1|2_), from the mean epistasis between mutations across all genotypes that contain this interaction (equation below and **Methods**). **c**, Absolute error of the predicted function (*F*) at orders 1, 2 and 3 (n=45), and at order 4 (n=23). The absolute error means and medians from the global (blue) and WT-background models (yellow) are shown as black and red bars, respectively. The 1.5-fold significance threshold is depicted as a dashed line, with an asterisk (*) marking the model and order with a significant mean (one-sided t-test; p = 9.67 × 10^−7^ and 1.83 × 10^−12^ for global model order 3 and 4, respectively).

We computed the predicted functions, using both the WT-background and global models, with gradually increasing orders up to the 4^th^ order (Fig. 4a and 4b). To assess the quality of the functional prediction performed by each model, we measured the absolute error (AE) of the predicted *versus* observed function for the “endpoint”, or most derived, genotype in each landscape (Fig. 4c and Methods).

Given the discrepancy in the amount of data accessed by the models, as expected, the mean AE of the global model outperformed the WT-background model at the lower orders. In the 1^st^ order, 13% (6/45) and 29% (13/45) of variants were successfully predicted by the WT-background and global model, respectively. The global model’s prediction accuracy increased with each incorporated order. Furthermore, the incorporation of 3^rd^ order information was largely sufficient for predicting the function of most endpoint variants below the significance threshold (p < 0.05; Fig. 4b and Extended Data Table 5). This suggests that average effects, encompassing relatively minimal epistatic information, can recapitulate the topology of the combinatorial landscapes. By contrast, the WT-background model showed high AEs with a marginal improvement in median AE, but not mean AE, even upon introducing 4^th^ order epistatic information (Fig. 4b and Extended Data Table 5). This poor performance of the WT-background model suggests that the Δ*F*_wt_ and ε_wt_ values were not sufficient predictors of the function of genotypes with multiple mutations, highlighting how idiosyncrasy confounds functional predictions in protein fitness landscapes.

### Idiosyncrasy disrupts the interpretability of adaptive trajectories

Next, we performed several analyses to assess the extent to which idiosyncrasy impairs the predictability of adaptive trajectories. To this end, we focused on 11 adaptive landscapes, *i*.*e*., landscapes where the substrate or ligand is assumed to be the primary selection pressure that led to the accumulation of the probed mutations. For each landscape, we retained a single most accessible path, in which the most functionally advantageous mutation is fixed at each step. We then computed a predicted function for each genotype along the most accessible path using ascending orders of the WT-background model (Fig. 5a and Extended Data Table 6). However, much as with the endpoint prediction, we again saw that for most genotypes the WT-background model failed to predict function accurately at all orders. Interestingly, some highly mutated genotypes along the trajectory were, in fact, better predicted by lower-order information. This indicates that when the WT-background model incorporates extensive idiosyncrasy at a given order, its predictive power is impaired relative to a model with less mutational information (Fig. 5a and Extended Data Table 6).

**Fig. 5.**
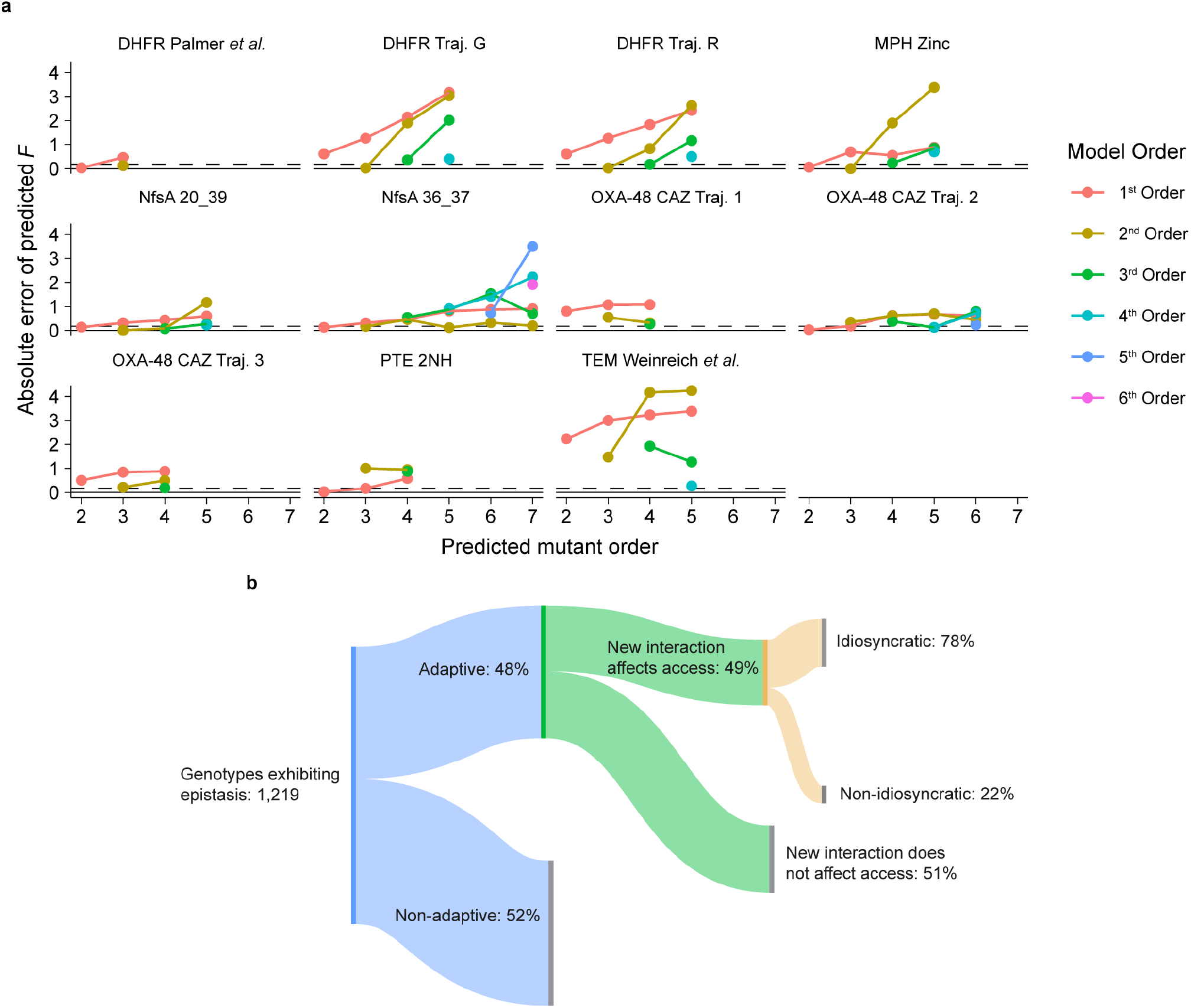
Impact of idiosyncratic epistasis on prediction and genotype accessibility. **a**, Absolute error of the functional prediction for each genotype along the most accessible path in 11 adaptive landscapes. Colors indicate the order at which the WT-background model was used. **b**, Sankey plot of epistasis and its link to genotype accessibility in adaptive landscapes. Epistasis encompasses all ε values of newly introduced interactions between ≥ 2 mutations that show > 1.5-fold magnitude in effect. Genotypes with new significant epistasis were filtered by their occurrence in the 11 adaptive, or 34 non-adaptive landscapes, then by modulation of accessibility by new interactions, and finally by idiosyncrasy.

Next, we sought to gain a statistical picture elucidating how idiosyncrasy can open or close certain evolutionary trajectories in the adaptive landscapes. First, we defined the genotype *n*+1 as accessible if its function increases, or does not decrease by more than 1.5-fold, compared to the ancestral *n* genotype. In other words, we enabled nearly neutral steps to occur by considering that a 1.5-fold decrease in function is insignificant. Then, we examined how newly introduced interactions (*e*.*g*., 4^th^ order epistasis for a quadruple mutant, but not 3^rd^ order or lower) affect the accessibility of a genotype, and marked the genotype as ‘affected by epistasis’ if removing the new interaction (ε_new_ = 0) meant that the genotype’s accessibility was affected. Furthermore, a genotype was considered ‘affected by idiosyncratic epistasis’ if its accessibility was affected upon converting the effect of the new epistatic interaction to the average epistasis of that interaction 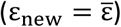 across the landscape.

By looking at all genotypes across the adaptive landscapes, we found that new epistatic interactions affected the accessibility of 49% (288/583) of genotypes. Furthermore, 78% (211/288) of these genotypes showed different accessibility specifically due to idiosyncratic epistasis (Fig. 5b). Of course, many of these genotypes are unlikely to be accessed at all, as early mutations with greater functional contributions will be selected first, consequently affecting the chance of visiting many intermediate genotypes. Thus, we further refined our accessibility metrics by focusing on the genotypes in the most accessible path for each adaptive landscape. Of these most accessible genotypes, the accessibility of 26% (11/42) was significantly affected by new interactions, and 73% (8/11) of those genotypes were idiosyncratic. Hence, although in the most accessible trajectories new epistatic interactions are generally less likely to affect the accessibility of a genotype, when they do impact the evolutionary path, they are usually idiosyncratic.

Through this analysis, we identified two trajectories exhibiting particularly interesting evolutionary dynamics caused by idiosyncrasy, and decided to probe their genetic and molecular underpinnings. The first trajectory was the most accessible path along the PTE evolution toward the arylesterase activity^13,30–32^. Along this trajectory, the highest functional peak is reached at genotype D233E-H254R-L271F-F306I with ∼1,700-fold improvement in function relative to WT (Fig. 6a). The accessibility of this genotype is permitted by a strong quadruple epistatic interaction between these positions, resulting in a 7.4-fold increase in function relative to the contribution from all single mutational, pairwise- and triple-interaction effects (Fig. 6b). Without this 4^th^ order epistasis, this genotype would become inaccessible, as it would lead to a ∼4.5-fold decrease in function relative to D233E-H254R-F306I. Interestingly, this genotype would also be inaccessible if the effect of the quadruple interaction was represented by its 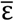 (Fig. 6b). In fact, 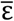 captures the incompatibility of the quadruple epistasis with L272M and I313F, illustrating the idiosyncratic nature of this quadruple interaction within the PTE landscape. In other words, this idiosyncratic quadruple interaction provides an evolutionary solution that permits access to the functional peak, but can quickly become disrupted by negative higher-order epistasis from other mutations. Idiosyncrasy can also, on the other hand, be restrictive. In the MPH landscape, five adaptive mutations were found to confer methyl-parathion hydrolase activity to MPH^14,25^. Yet, a pair of individually beneficial mutations exhibit negative epistasis when combined: the L72R-F273L interaction restricts the transition from the single (F273L) to double mutant (L72R-F273L) due to a ∼1.7-fold decrease in function (Fig. 6c). However, the mean pairwise epistasis between L72R and F273L is significantly positive (+4.0-fold), suggesting that the L72R-F273L pair becomes beneficial later in the trajectory. Indeed, this sign change in ε results from the early fixation of Δ193S, which greatly stabilizes the L72R-F273L interaction, and is further reinforced by positive higher-order interactions with H258L and I271T (Fig. 6d). Thus, higher-order epistasis appears to be the key mechanism underlying the compensation of this initially idiosyncratic antagonism, later fostering the pair’s synergy. These two examples demonstrate how idiosyncrasy is a by-product of higher-order interactions – whether synergistic or antagonistic – and how it can create localized permissive or restrictive dynamics in combinatorial landscapes that are concealed by the reductionism of the global view.

**Fig. 6.**
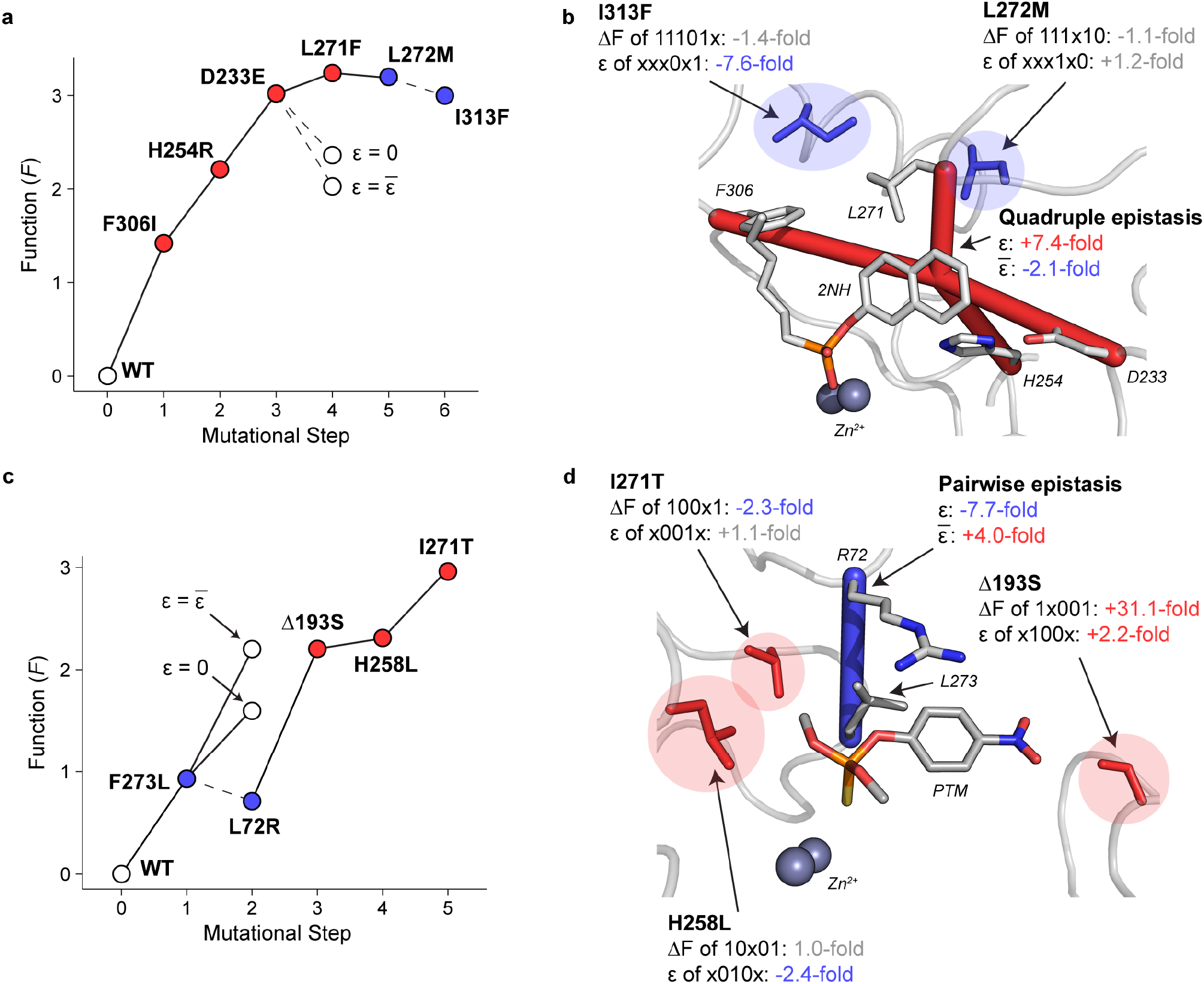
Structural representation of idiosyncrasy and its effect on evolutionary trajectory accessibility. **a**,**c**, Function of genotypes along **a**, the PTE trajectory toward arylester (2NH) hydrolysis, and **c**, the MPH trajectory toward parathion-methyl (PTM) hydrolysis. Key steps with no epistasis (*ε* = 0) or with mean epistasis 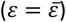 are highlighted along each trajectory for comparison. **b**, Structure of WT PTE (PDB: 4PCP) overlaid with an evolved PTE variant containing a 2-naphthyl hexanoate (2NH) transition state analogue (PDB: 43ET). **d**, Structure of a derived MPH variant (PDB: 1P9E) with molecular docking of parathion-methyl (PTM)^*14*^. Red bars represent positive epistasis, while blue bars indicate negative epistasis. Idiosyncratic (*ε*) and mean epistasis (*ε*) are displayed for each interaction. Blue and red residues represent destabilizing and stabilizing mutations with respect to idiosyncratic epistasis, respectively. The blue and red color coding is consistent across all panels.

Lastly, we explored whether the high levels of idiosyncrasy could be explained by other features in our adaptive trajectories. For example, it has been debated whether the functional contribution of a mutation is affected by the functional magnitude of the genotype in which it occurs^33^. Indeed, several adaptive trajectories in our study exhibited diminishing returns topology, where the functional contribution of mutations decreased as they were introduced into fitter, and more derived, backgrounds (Extended Data Fig. 2a). This trend has often been attributed to negative epistasis arising between mutations that originally appeared beneficial, a phenomenon called diminishing return epistasis^18,20,21,34^. However, upon probing ε in the adaptive landscapes, we found no trend in ε based on the functional level of intermediates, suggesting no evidence of the dominance of diminishing return epistasis (Extended Data Fig. 2b). As such, the patterns of diminishing returns observed in the adaptive landscapes are likely the result of other mechanisms, such as beneficial mutational exhaustion, whereby the most beneficial adaptive mutations are fixed in early rounds of evolution, quickly depleting the pool of beneficial mutations as evolution progresses.

We also explored idiosyncrasy from the context of protein structures. Since intramolecular protein epistasis is attributed to the interaction between mutations, we analyzed the spatial proximity between the epistatic positions as a function of the magnitude of epistasis between said positions. However, we observed no significant correlation between the distance connecting the a-carbons of two residues and the absolute magnitude of their pairwise ε (Extended Data Fig. 3). Thus, other factors have to be considered when characterising the relationship between spatial proximity of mutations and epistasis, for instance, non-specific epistatic effects^35^, side chain interactions, or perhaps complex mutational interactions including conformational dynamics or ligand and substrate relays^1^.

## Discussion

In this study, we demonstrated high levels of idiosyncrasy in both single mutational effects and epistasis across 45 combinatorial landscapes representing seven enzymes. The success in predicting the functional effects within combinatorial landscapes by the global model, using epistatic coefficients as low as the 3^rd^ order (Fig. 4c), may suggest that fitness landscapes (up to seven mutational combinations) can be explained by relatively simple interaction terms^16,17^. In this study, however, we find that the global view is not always useful to deconvolute specific evolutionary trajectories, because each mutational path can contain idiosyncrasy that permits or restricts the accessibility of certain genotypes. The idiosyncrasy is inherently a by-product of higher-order epistasis; thus, higher-order epistasis should not just be perceived as an additional epistatic contribution stemming from three or more characterised mutations, but more importantly, as the source of evolutionary idiosyncrasy. Indeed, we observed high idiosyncrasy even in the 3^rd^ and 4^th^ orders of epistasis suggesting that epistasis at the 5^th^ – and higher – order(s) is creating strong idiosyncrasy which, in turn, leads to a strong deviation between the epistasis arising in a particular genotype and its global (or average) effect (Fig. 3c and Extended Data Figs. 3-4). Thus, as a protein gradually accumulates mutations, the single mutational effects and epistasis are expected to be steadily idiosyncratic, and therefore predicting their evolutionary behaviours at any given genotype is challenging.

The underlying biophysical basis of idiosyncrasy has yet to be uncovered. We suspect that the observed epistatic interactions are partitioned into subnetworks within the grand interaction network of the protein, including the protein’s ligands, cofactors, and substrates^1,36–38^. If the modularity of protein epistatic networks is indeed a key factor in dictating idiosyncrasy, and therefore a major component of understanding higher-order epistasis, it is essential to combine epistatic analysis with techniques probing the molecular and structural bases of these networks^39^, *e*.*g*., using tools such as molecular dynamics (MD)^38,40^ and nucleic magnetic resonance (NMR)^41^, particularly in the context of interactions through other molecules^1^. By deepening our understanding of these parameters, we may be able to account for idiosyncrasy *a priori* and smoothen the apparent noise in our protein fitness landscapes.

Our observations also provide important implications for the engineering of new proteins. Recent advances in high-throughput mutational characterisation approaches, such as deep mutational scanning (DMS), provide access to a large mutational dataset around the WT background but are generally limited to single and double mutants^42,43^. However, engineered targets generally stray further from their genotype, and, as we have shown, local information is insufficient for the accurate prediction of evolutionary intermediates and endpoints (Fig. 4c and 5a). In some instances, this caveat has been overcome using *in silico* techniques employing co-evolutionary based models such as direct coupling analysis^44^, or natural language processing^45^, which demonstrate great success in predicting sequences with enhanced functions. Albeit, such techniques are currently unsatisfactory when it comes to predictions of evolutionary paths towards non-native ligands and substrates. We acknowledge that there are more sophisticated machine-learning (ML) based models that may provide greater success in functional prediction than the ones used in this study^46,47^. However, explicit incorporation of higher-order epistasis into these models has shown impaired prediction, possibly due to the incorporation of ‘noise’ stemming from idiosyncratic effects^46^. Nevertheless, our results advocate attempts to incorporate idiosyncrasy into ML models, or at least warrant an acknowledgement of the ability to which idiosyncrasy can distort predictions. Likewise, the prevalence and impact of idiosyncrasy shown in our study reaffirm the importance of broader fitness landscape exploration, where ML can also be employed^47^. We hope that in the future it may be feasible to use the predictive power of ML models and broader characterisation of fitness landscapes, combined with other biophysical and biochemical inputs such as DMS, structural prediction tools, and biophysical analyses, to gain a deeper understanding of idiosyncrasy and higher-order epistasis, greatly elevating the level of prediction for protein engineering efforts.

## Methods

### PTE Combinatorial landscape

During the directed evolution of PTE for higher arylesterase activity, we previously identified a cluster of six function-altering mutations^30–32^. Here, we construct one of the combinatorial landscapes analysed in this study. We explored these six positions on the genetic background of WT PTE (64 variants), and tested all of their combinations for activity against 2-naphthyl hexanoate (2NH), as described previously^13^. Briefly, the 64 variants were constructed by site-directed mutagenesis and subcloned into a pET-27-STREP vector^13^. The variants were then transformed into *E. coli* BL21(DE3) carrying the pGro7 plasmid (Takara, Shiga, Japan) for GroEL/ES chaperones co-expression^48^. Variants were individually inoculated in 96-deep well plates containing lysogeny broth (LB) media, 100 μg/mL ampicillin, and 34 μg/mL chloramphenicol, then grown overnight at 30°C. Overnight cultures were transferred to a new deep well plate containing LB, supplemented with 100 μg/mL ampicillin, 34 μg/mL chloramphenicol, 200 μM ZnCl_2_, and 0.2% (w/v) arabinose for chaperone co-expression, then induced with 1 mM IPTG. Pellets were lysed with lysis buffer (50 mM Tris-HCl buffer, 100 mM NaCl, pH 7.5, 0.1% (w/v) Triton-X100, 200 μM ZnCl_2_, 100 μg/mL lysozyme and 1 μL benzonase (25 U/μL) per 100 mL of lysis buffer). Lysates were incubated with 200 μM 2NH + 1 mM Fast Red and hydrolysis was monitored at 500 nm.

### Data processing

Data from various studies were compiled and standardized to a spreadsheet format described previously^13^. The values of reported functions were divided by the WT-background function, or, in the presence of replicates, by the mean of the WT-background functions, then log_10_ transformed. However, we raised 10 to the power of all the TEM-1 growth rate values from Mira *et al*. before using them in our standard pipeline, as the growth rates from this study are assumed to be additive and not multiplicative. All processed files are provided (Supplementary File 1). We also chose not to apply the non-linear power transform^19,49^, which we address in the supplementary material (Supplementary File 3).

### Δ*F* and ε calculation

Genotypes were represented by a string of amino acids that underwent mutation, then encoded using ‘0’ for ancestral states, and ‘1’ for derived states at the given amino acid positions. The single mutational effect (Δ*F*) of a mutation at position *i* was calculated for each genetic background by computing the difference between *F*_i=0_ and *F*_i=1_. The mutational transition for the Δ*F* is denoted with ‘x’ which represents a transition from ‘0’ to ‘1’, *e*.*g*., the Δ*F*_x001_ represents Δ*F* of mutating position 1 in the position 4 mutant background and is equal to F_1001_ - F_0001_. Epistasis (ε) between two positions *i* and *j* were calculated for each genetic background by computing the difference between the Δ*F*_i=x, j=0_ and Δ*F*_i=x, j=1_. For example, the ε_xx00_ represents the additional functional contribution of the combination of the first and second position in the WT background and is equal to Δ*F*_x100_ – Δ*F*_x000_. This is equivalent to Δ*F*_1×00_ – Δ*F*_0×00_. As with pairwise interactions, higher-order ε was calculated by taking the difference between ε of the previous order, *e*.*g*., ε_xxx0_ = ε_xx10_ – ε_xx00_.

### WT-background model

The function of a genotype was predicted using the WT-background model as a sum of all mutational (Δ*F*) and epistatic (ε) effects in the WT-background:

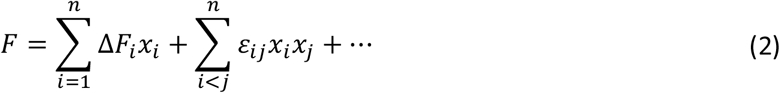

Where *i* (and *j*) represents the index of residue position, x_i_ is either ‘0’ or ‘1’ depending on the mutational state of the residue in the given genotype, and *F* is the function. This is analogous to the biochemical view of epistasis from Poelwijk *et al*.^16^, however, our model uses *F* values that do not represent ΔG of the protein or enzymatic reaction.

### Global model

The linear regression model, or global model, employed for epistatic analysis was the same as those used previously^13,14,25^. Briefly, mutations at *n* residue positions were annotated with variables ‘-1’ for the ancestral state or ‘1’ for the derived state. These were used as x variables in the linear model, *F* is the log_10_ transformed and WT-normalized function of the variant. The linear model was constructed such that:

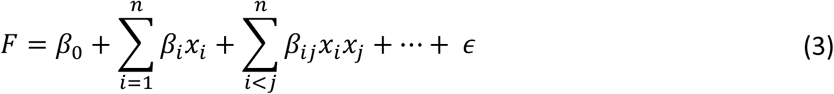

Where *i* (and *j*) represents the index of residue position, x_i_ is either ‘-1’ or ‘1’ depending on the mutational state of the residue in the given genotype, β are the linear coefficients, and *ε* is the error^16^.

### Absolute Error (AE) calculation for the endpoint variant

For each of the 45 combinatorial landscapes, the endpoint was selected as the genotype furthest mutationally removed from the WT-background. AE of the prediction from each order of the WT background and linear model was calculated using the following formula:

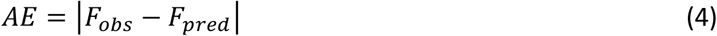

Where *F*_*obs*_ is the log_10_ WT-normalized function of the final variant in the landscape, and *F*_*pred*_ is the log_10_ WT-normalized predicted function of the final variant. The mean and median of the AEs in Fig. 4c represents the arithmetic mean and median of the AEs for each model and order pairing.

## Code and Data Availability

Data for all combinatorial landscapes are provided in log_10_-transformed WT-normalized format (Supplementary Data 1). Processed data for functional contributions (Supplementary Data 2) and epistasis (Supplementary Data 3) are also available. Scripts for individual combinatorial landscape analysis and the global statistical analysis are publicly available via GitHub at https://github.com/karolbuda/epistasis_analysis_v2. The scripts for plotting epistasis on PDB structures were adapted from Miton *et al*.^13^. The scripts utilize the R language (https://www.R-project.org/), along with R packages: tidyverse (https://CRAN.R-project.org/package=tidyverse), igraph (https://CRAN.R-project.org/package=igraph), gtools (https://CRAN.R-project.org/package=gtools), e1071 (https://CRAN.R-project.org/package=e1071), svglite (https://CRAN.R-project.org/package=svglite), ggrepel (https://CRAN.R-project.org/package=ggrepel), ggpubr (https://CRAN.R-project.org/package=ggpubr), and knitr (https://yihui.org/knitr).

## Extended Data

**Extended Data Fig 1.**
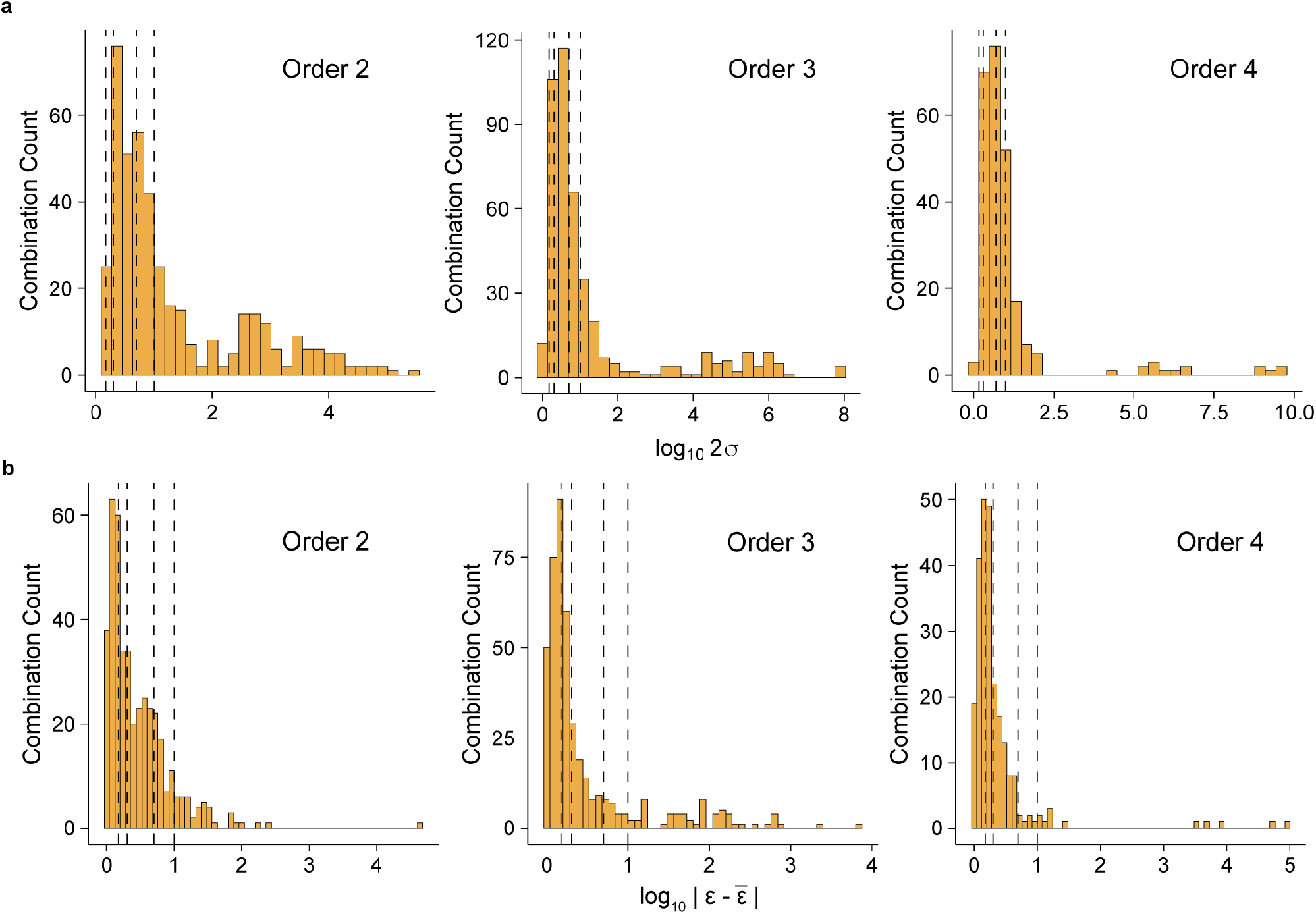
Expanded histograms from Figs. 3a and 3c. **a**, Distribution of log10 2σ of ε at each order for all mutational combinations, with annotated dashed lines representing 1.5-fold, 2-fold, 5-fold, and 10-fold significance thresholds, respectively. **b**, Distribution of the WT-background idiosyncratic epistasis quantified via the absolute difference of *ε* from 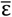 at each combination. Dashed lines represent 1.5-, 2-, 5-, and 10-fold significance thresholds, respectively.

**Extended Data Fig 2.**
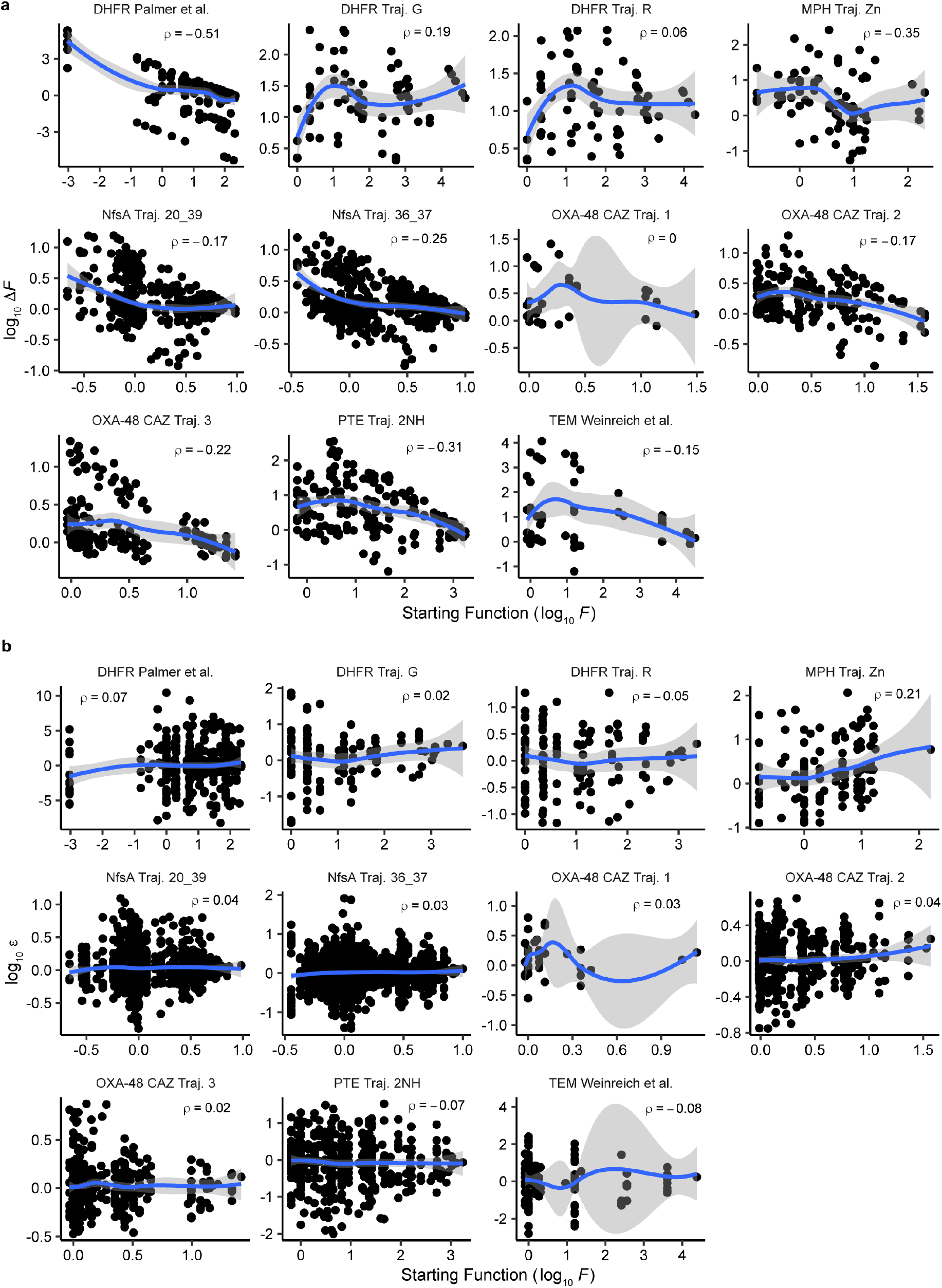
Trends in mutational and epistatic effects with respect to a genotype’s starting function. **a**, Relationship between stating function and mutational functional contribution (Δ*F*) for 11 adaptive landscapes. **b**, Relationship between stating function and epistasis (*ε*) for 11 adaptive landscapes. Lines represent local regressions using the LOESS method. Rho (*ρ*) represents the spearman correlation coefficient for each landscape.

**Extended Data Fig 3.**
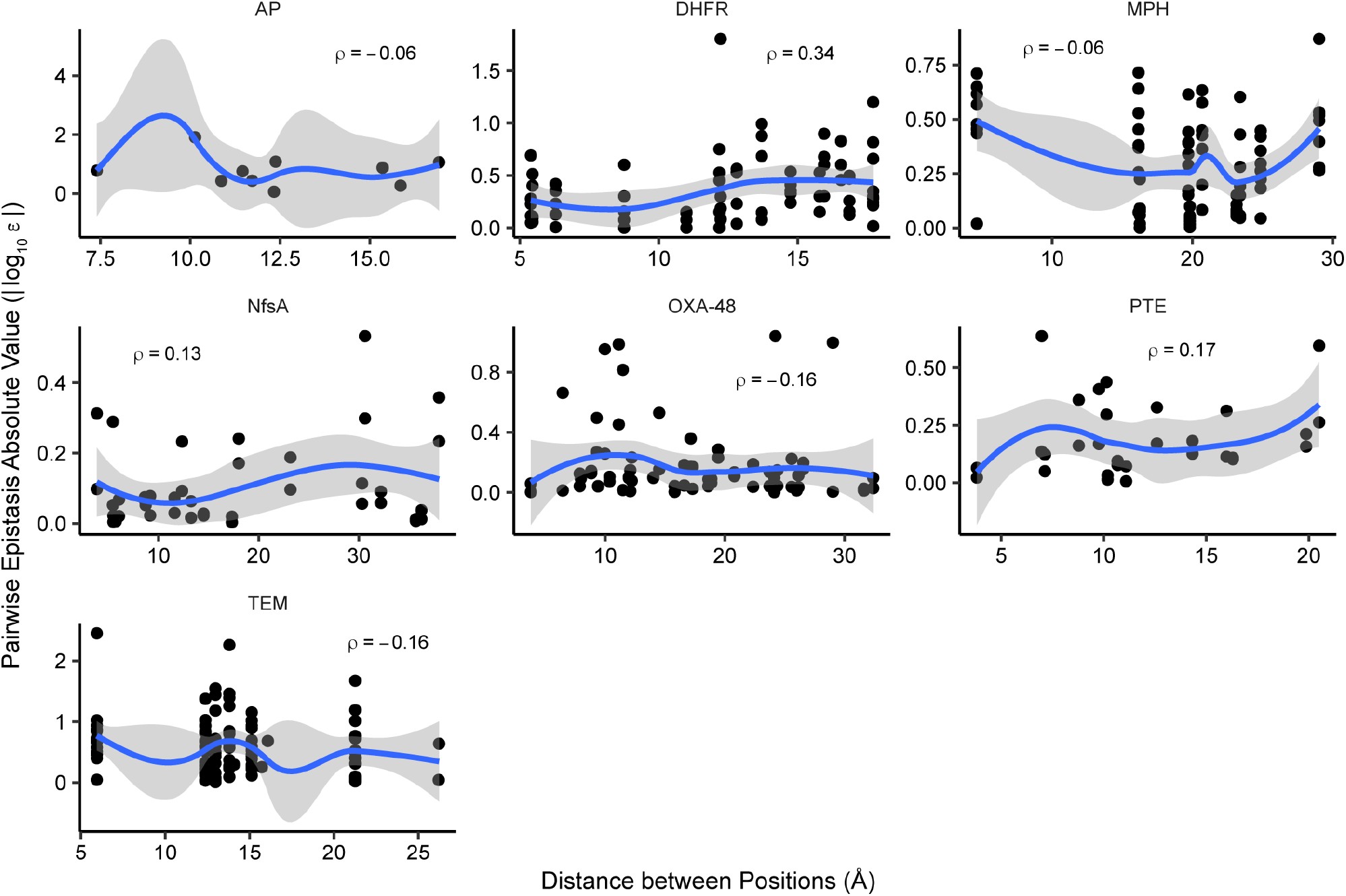
Trends in pairwise epistasis with respect to alpha carbon distance between the two positions. Only epistatic values from the 11 adaptive landscapes were used. Structures were obtained from PDB IDs (3TG0, 3CXK, 6A2K, 5HIF, 7NMP, 3HBR, 4PCP, 1XPB) for AP, *E*.*coli* DHFR, *P. falciparum* DHFR, MPH, NfsA, OXA-48, PTE, and TEM respectively. Lines represent local regressions using the LOESS method. Rho (*ρ*) represent the spearman correlation coefficient for each landscape.

**Extended Data Table 1.** Sign variation of the positions’ (1^st^ order) and combinations’ (2^nd^, 3^rd^, and 4^th^ orders) Δ*F’*s and ε.

| Order | Homogeneous |  |  | Semi-Homogeneous |  | Heterogeneous |
| --- | --- | --- | --- | --- | --- | --- |
|  | Negative | Neutral | Positive | Negative-Neutral | Neutral-Positive | Negative-Positive |
| 1 | 1.4% (3/214) | 0.9% (2/214) | 9.8% (21/214) | 7.5% (16/214) | 16.4% (35/214) | 64.0% (137/214) |
| 2 | 1.2% (5/419) | 1.2% (5/419) | 3.1% (13/419) | 8.8% (37/419) | 15.3% (64/419) | 70.4% (295/419) |
| 3 | 3.7% (16/438) | 2.3% (10/438) | 11.9% (52/438) | 14.6% (64/438) | 13.5% (59/438) | 54.1% (237/438) |
| 4 | 4.5% (11/245) | 2.0% (5/245) | 4.1% (10/245) | 20.4% (50/245) | 17.1% (42/245) | 51.8% (127/245) |

**Extended Data Table 2.** Percent of positions (1^st^ order) and combinations (2^nd^, 3^rd^, and 4^th^ order) that show a 2σ for Δ*F* or ε above the listed thresholds.

| Order | Threshold |  |  |  |
| --- | --- | --- | --- | --- |
|  | 1.5-fold | 2-fold | 5-fold | 10-fold |
| 1 | 98.6% (211/214) | 95.8% (205/214) | 70.1% (150/214) | 53.3% (114/214) |
| 2 | 98.6% (413/419) | 92.4% (387/419) | 60.1% (252/419) | 40.6% (170/419) |
| 3 | 95.9% (420/438) | 85.4% (374/438) | 45.4% (199/438) | 29.7% (130/438) |
| 4 | 98.0% (240/245*) | 83.3% (204/245*) | 49.8% (122/245*) | 24.9% (61/245*) |

\* 4<sup>th</sup> order considers 245 out of the total 267 combinations as some landscapes have a single 4<sup>th</sup> order data point and a 2σ cannot be computed

**Extended Data Table 3.**
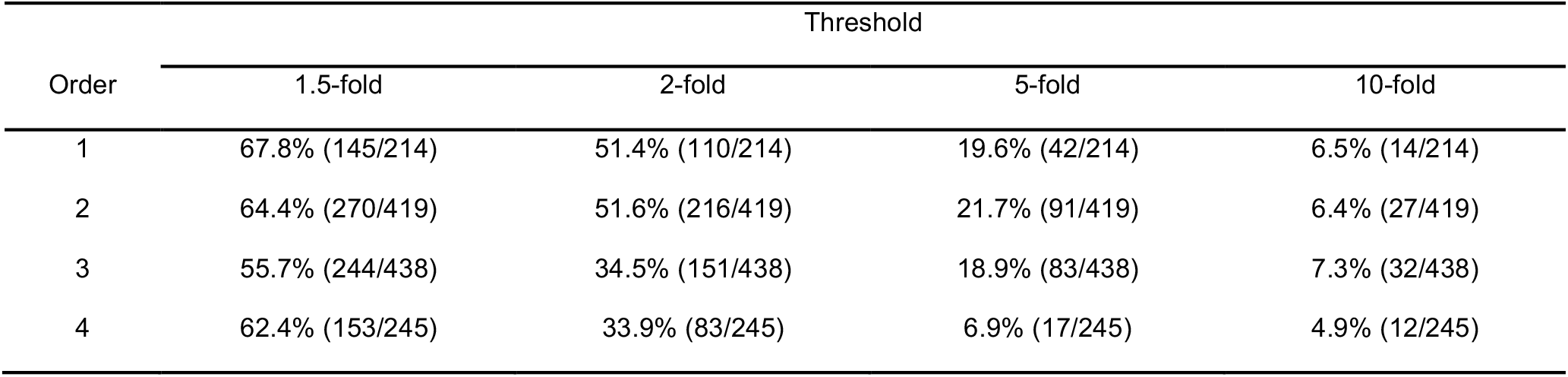
Percent of positions and combinations that show a WT Δ*F* or WT ε to mean Δ*F* or mean ε deviation above the listed thresholds.

| Order | Threshold |  |  |  |
| --- | --- | --- | --- | --- |
|  | 1.5-fold | 2-fold | 5-fold | 10-fold |
| 1 | 67.8% (145/214) | 51.4% (110/214) | 19.6% (42/214) | 6.5% (14/214) |
| 2 | 64.4% (270/419) | 51.6% (216/419) | 21.7% (91/419) | 6.4% (27/419) |
| 3 | 55.7% (244/438) | 34.5% (151/438) | 18.9% (83/438) | 7.3% (32/438) |
| 4 | 62.4% (153/245) | 33.9% (83/245) | 6.9% (17/245) | 4.9% (12/245) |

**Extended Data Table 4.** Percent of genotypes that show a Δ*F* or ε to mean Δ*F* or mean ε deviation above the listed thresholds.

| Order | Threshold |  |  |  |
| --- | --- | --- | --- | --- |
|  | 1.5-fold | 2-fold | 5-fold | 10-fold |
| 1 | 59.0% (2396/4064) | 40.5% (1646/4064) | 14.8% (601/4064) | 8.1% (331/4064) |
| 2 | 52.8% (2466/4672) | 34.9% (1630/4672) | 13.4% (624/4672) | 7.7% (358/4672) |
| 3 | 49.3% (1467/2976) | 30.7% (913/2976) | 11.6% (345/2976) | 7.6% (227/2976) |
| 4 | 60.0% (627/1120) | 34.6% (387/1120) | 9.0% (101/1120) | 5.0% (56/1120) |

**Extended Data Table 5.**
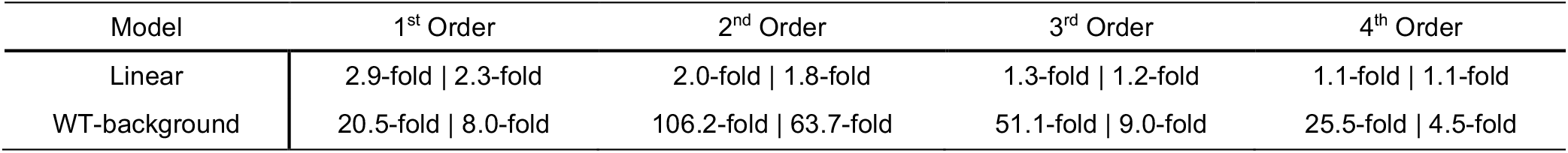
Mean and median absolute error of the models’ prediction for the derived variant at each model order.

**Extended Data Table 6.** WT-background model absolute error in predicted log-transformed function (log_10_ *F*) of each variant in the most accessible path by model order.

| Predicted Variant Mutations | Trajectory | Order |  |  |  |  |  |
| --- | --- | --- | --- | --- | --- | --- | --- |
|  |  | 1<sup>st</sup> | 2<sup>nd</sup> | 3<sup>rd</sup> | 4<sup>th</sup> | 5<sup>th</sup> | 6<sup>th</sup> |
| 2 | DHFR IC<sub>75</sub> | **<u>0.03</u>** | - | - | - | - | - |
| 3 |  | 0.46 | **<u>0.12</u>** | - | - | - | - |
| 2 | DHFR G-Traj. | **0.61** | - | - | - | - | - |
| 3 |  | 1.26 | **<u>0.02</u>** | - | - | - | - |
| 4 |  | 2.14 | 1.90 | **0.36** | - | - | - |
| 5 |  | 3.17 | 3.04 | 2.02 | **0.40** | - | - |
| 2 | DHFR R-Traj | **0.61** | - | - | - | - | - |
| 3 |  | 1.26 | **<u>0.02</u>** | - | - | - | - |
| 4 |  | 1.84 | 0.83 | **<u>0.17</u>** | - | - | - |
| 5 |  | 2.43 | 2.63 | 1.17 | **0.50** | - | - |
| 2 | MPH Zn PTM | **<u>0.06</u>** | - | - | - | - | - |
| 3 |  | 0.69 | **<u>0.00</u>** | - | - | - | - |
| 4 |  | 0.55 | 1.90 | **0.23** | - | - | - |
| 5 |  | 0.88 | 3.38 | 0.85 | **0.70** | - | - |
| 2 | NfsA 20_39 | **<u>0.14</u>** | - | - | - | - | - |
| 3 |  | 0.32 | **<u>0.01</u>** | - | - | - | - |
| 4 |  | 0.44 | **<u>0.09</u>** | **<u>0.07</u>** | - | - | - |
| 5 |  | 0.60 | 1.17 | 0.27 | **0.23** | - | - |
| 2 | NfsA 36_37 | **<u>0.13</u>** | - | - | - | - | - |
| 3 |  | 0.32 | **<u>0.17</u>** | - | - | - | - |
| 4 |  | 0.47 | **0.46** | 0.54 | - | - | - |
| 5 |  | 0.81 | **<u>0.12</u>** | 0.88 | 0.91 | - | - |
| 6 |  | 0.87 | **0.33** | 1.53 | 1.42 | 0.74 | - |
| 7 |  | 0.91 | **0.2** | 0.71 | 2.23 | 3.50 | 1.91 |
| 2 | OXA-48 CAZ Traj. 1 | **0.80** | - | - | - | - | - |
| 3 |  | 1.07 | **0.55** | - | - | - | - |
| 4 |  | 1.08 | 0.32 | **0.27** | - | - | - |
| 2 | OXA-48 CAZ Traj. 2 | **<u>0.03</u>** | - | - | - | - | - |
| 3 |  | **0.18** | 0.34 | - | - | - | - |
| 4 |  | 0.63 | 0.62 | **0.38** | - | - | - |
| 5 |  | 0.68 | 0.70 | **<u>0.13</u>** | **<u>0.13</u>** | - | - |
| 6 |  | 0.61 | 0.45 | 0.80 | 0.72 | **0.24** | - |
| 2 | OXA-48 CAZ Traj. 3 | **0.50** | - | - | - | - | - |
| 3 |  | 0.84 | **0.21** | - | - | - | - |
| 4 |  | 0.88 | 0.50 | **0.19** | - | - | - |
| 2 | PTE 2NH | **<u>0.03</u>** | - | - | - | - | - |
| 3 |  | **<u>0.17</u>** | 1.00 | - | - | - | - |
| 4 |  | **0.58** | 0.95 | 0.88 | - | - | - |
| 2 | TEM MIC | **2.23** | - | - | - | - | - |
| 3 |  | 3.00 | **1.47** | - | - | - | - |
| 4 |  | 3.23 | 4.16 | **1.93** | - | - | - |
| 5 |  | 3.38 | 4.24 | 1.27 | **0.27** | - | - |

## Acknowledgements

We thank D. Anderson and members of the Tokuriki lab for valuable comments. This work was supported by the Natural Sciences and Engineering Research Council of Canada (NSERC) Discovery Grant (RGPIN 2017-04909) and Human Frontier Science Program (HFSP) Research Grant (RGP0054/2020).

## Author Contributions

K.B. and N.T. conceived of and designed the study. K.B. and N.T. performed all statistical analyses. C.M.M. performed activity assays for the PTE combinatorial landscape. All authors contributed to writing the paper.

