## Supplementary File 1 for "Higher-order epistasis creates idiosyncrasy, confounding predictions in protein evolution"

Representations of 45 fitness landscapes used in the study. Landscapes depict fold-change in activity from their WT background. The Mira *et al.* TEM landscapes depict $10^{\mu1-\mu2}$, where μ1 is the genotype's growth rate and μ2 is the WT growth rate.

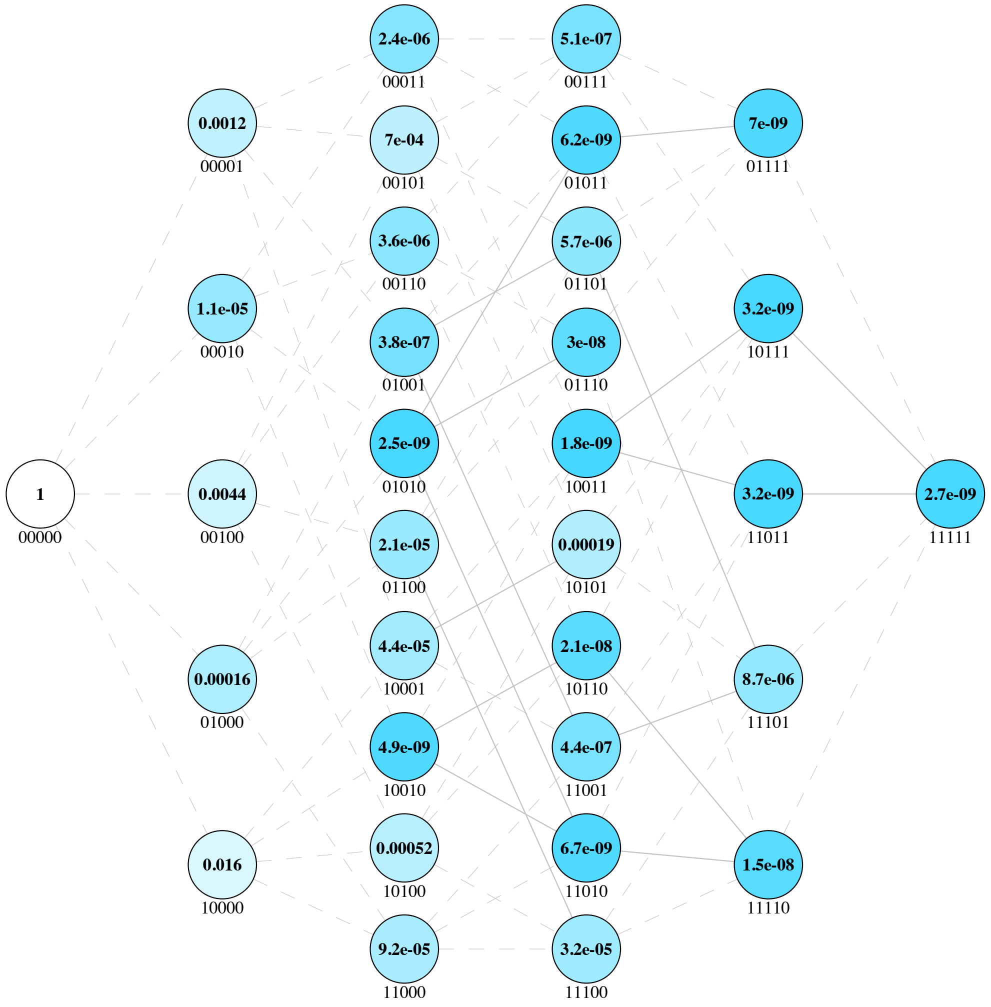


**Landscape 1 | Fitness Landscape for Alkaline Phosphatase (AP) from Sunden *et al.***

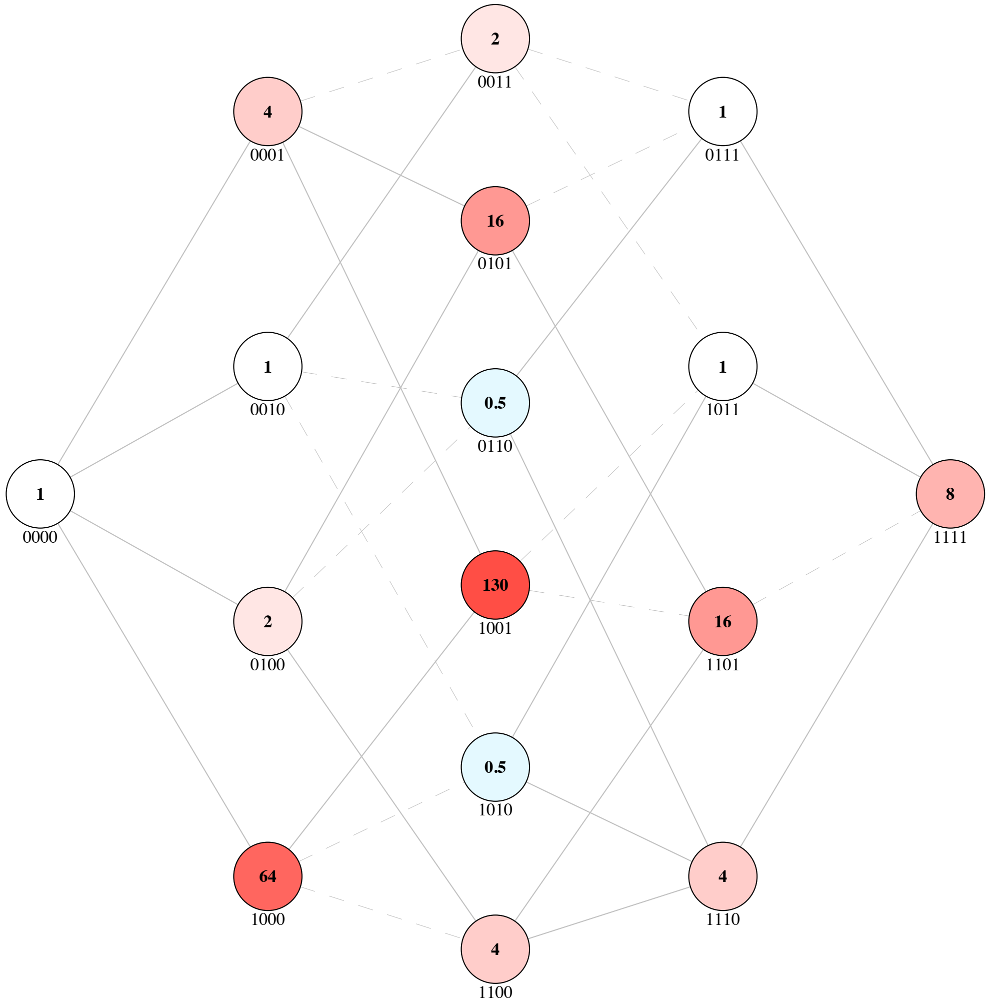


**Landscape 2 | Fitness Landscape of Dihydrofolate Reductase (DHFR) for inhibitor c57 from Lozovsky *et al.***

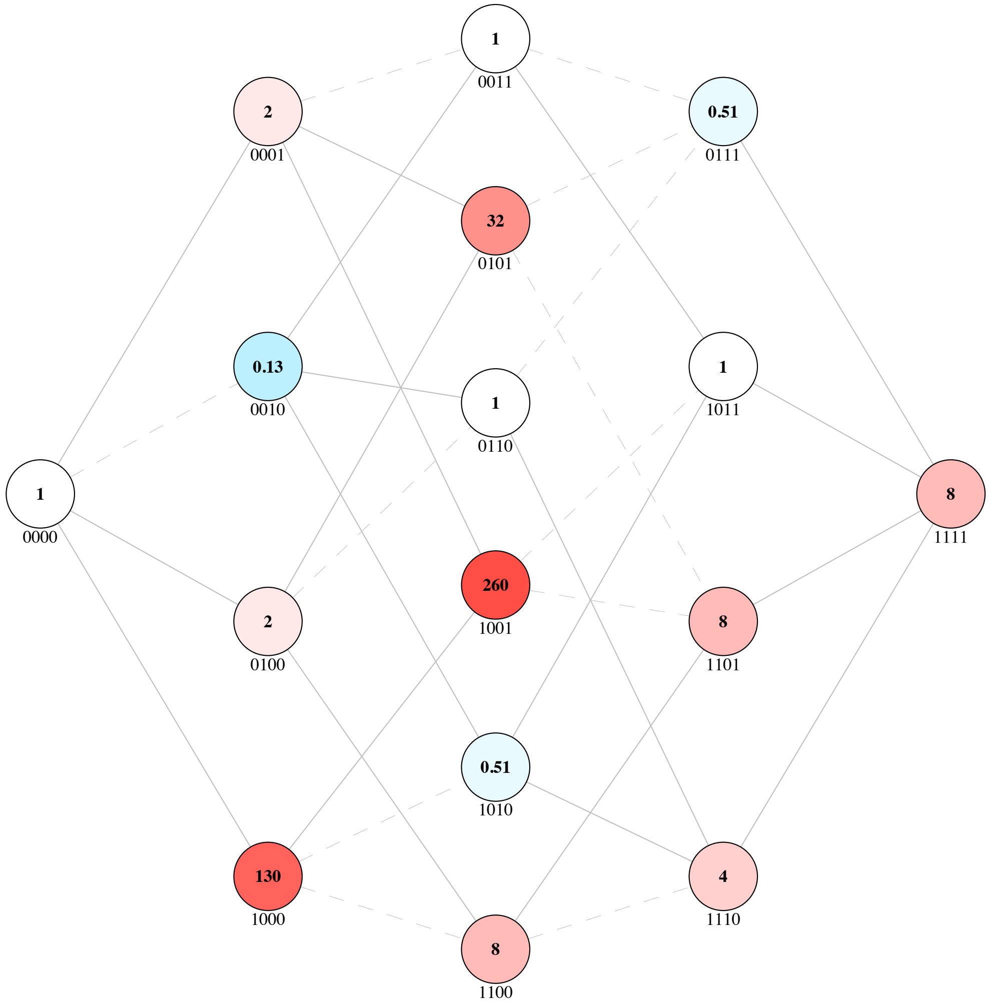


**Landscape 3 | Fitness Landscape of Dihydrofolate Reductase (DHFR) for inhibitor c58 from Lozovsky *et al.***

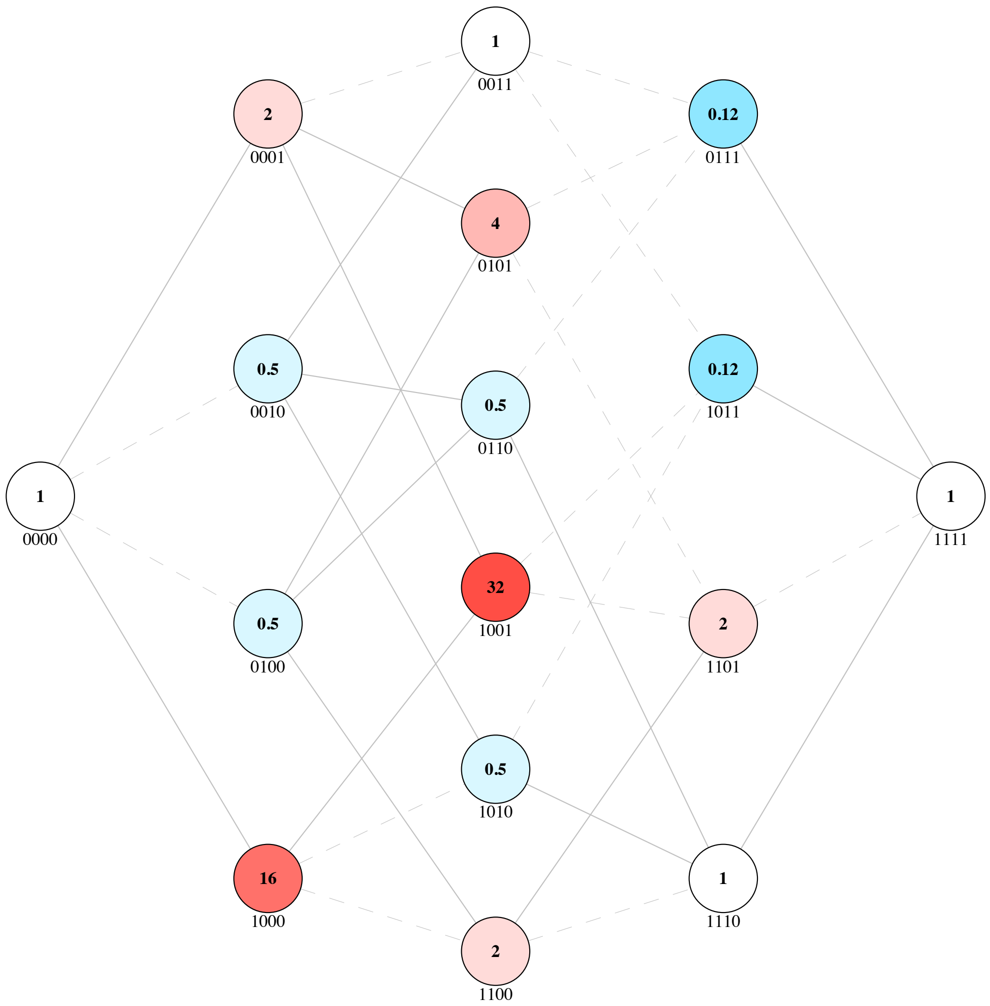


**Landscape 4 | Fitness Landscape of Dihydrofolate Reductase (DHFR) for inhibitor c59 from Lozovsky *et al.***

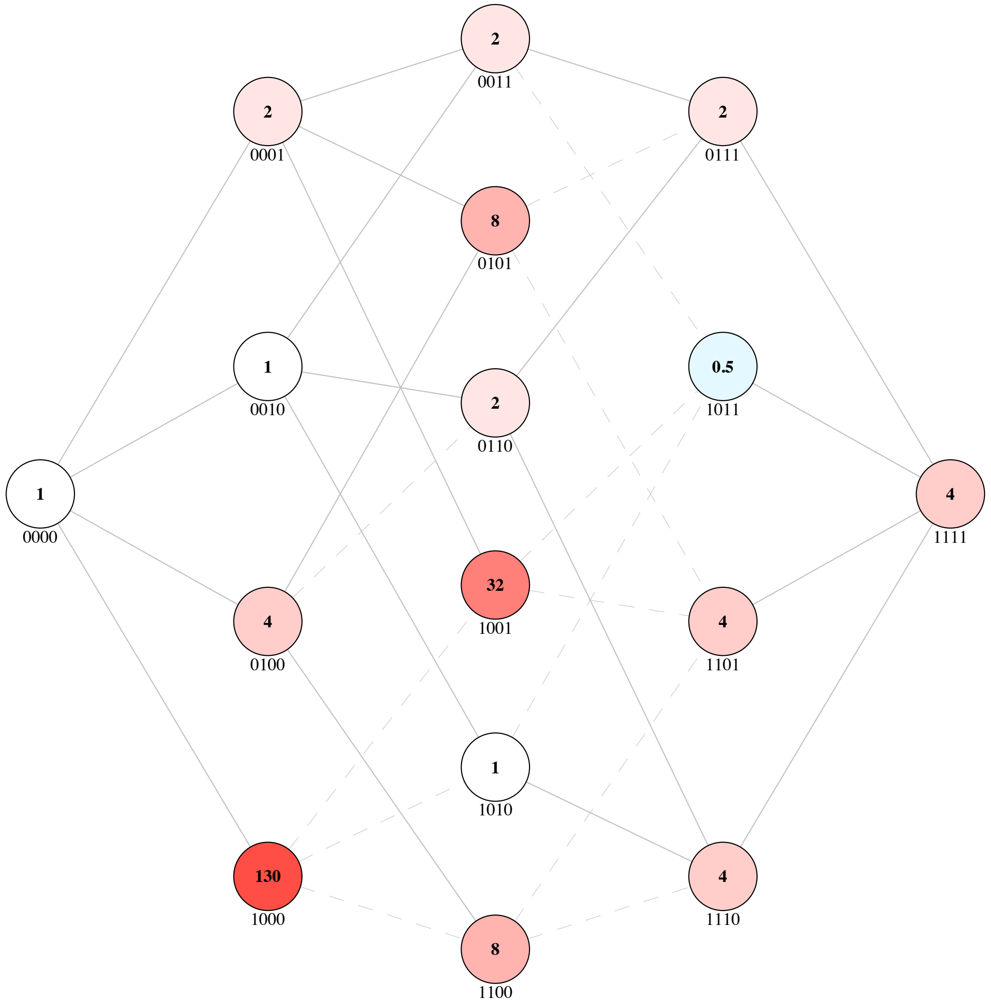


**Landscape 5 | Fitness Landscape of Dihydrofolate Reductase (DHFR) for inhibitor c60 from Lozovsky *et al.***

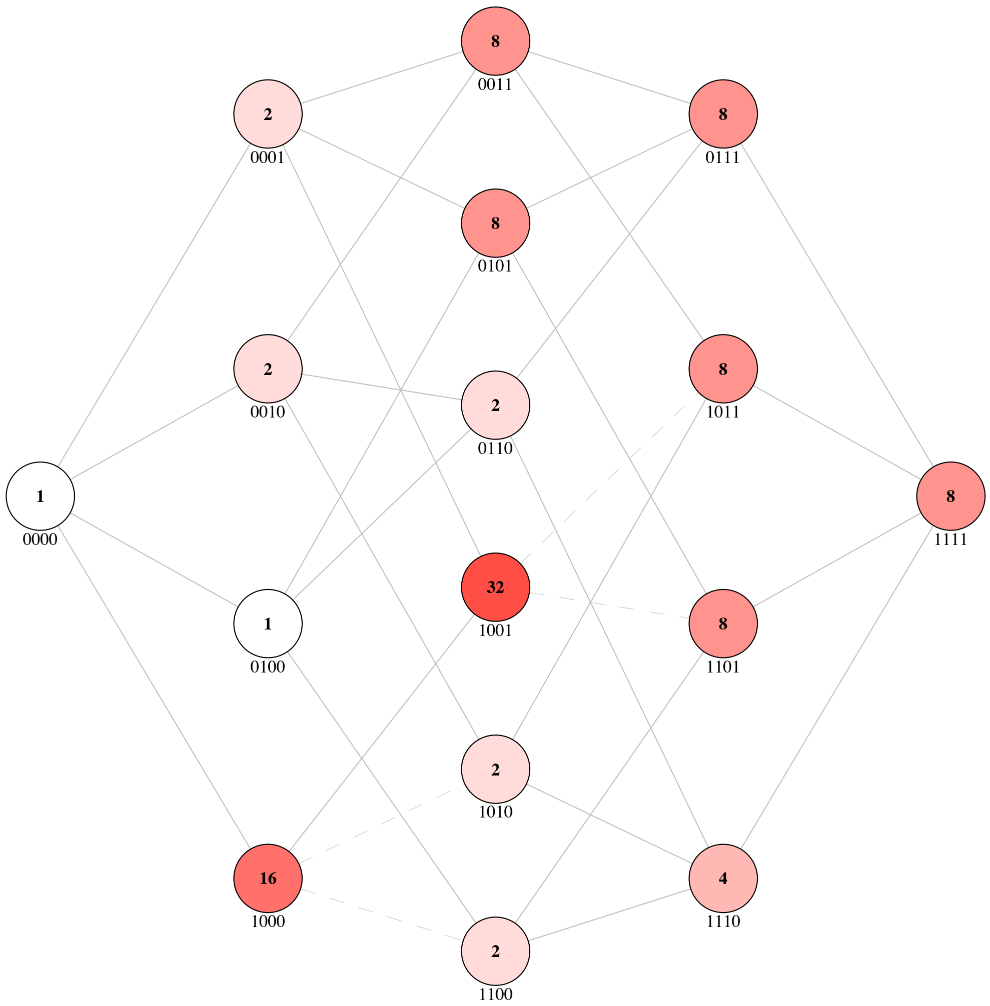


**Landscape 6 | Fitness Landscape of Dihydrofolate Reductase (DHFR) for inhibitor c61 from**

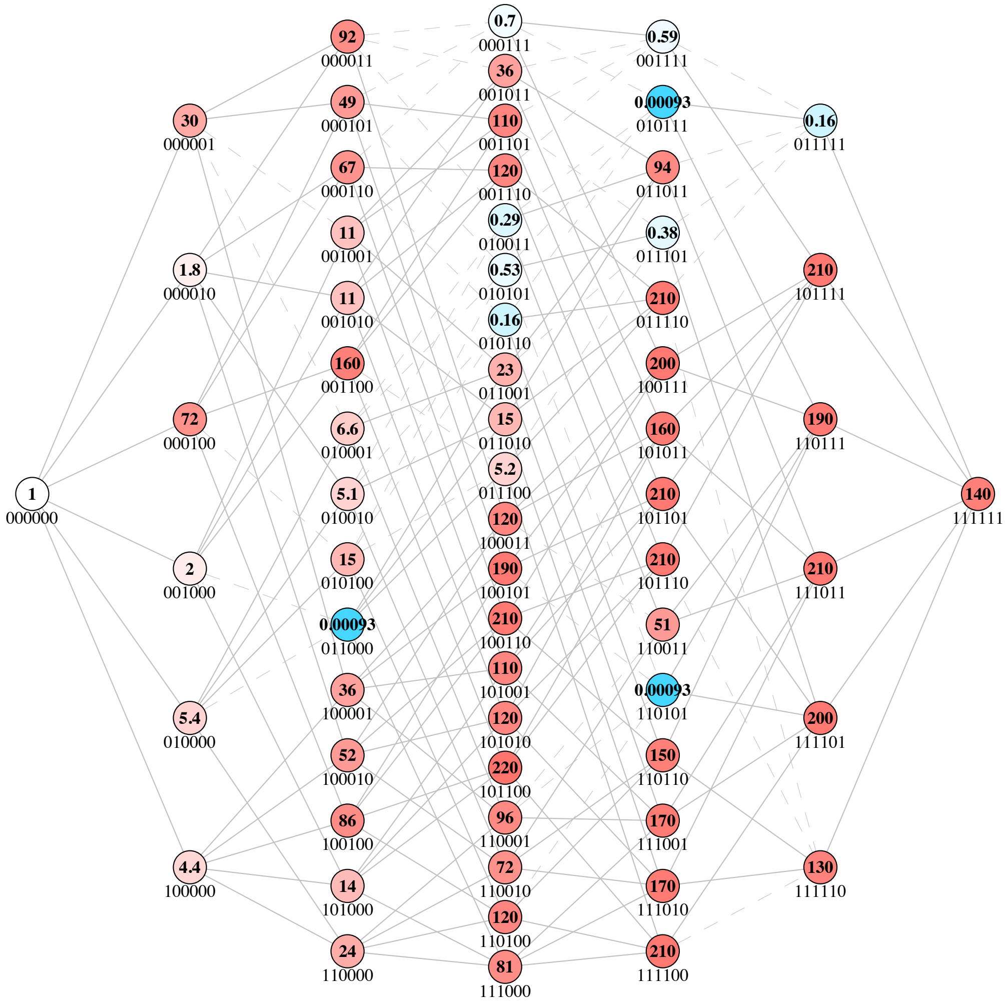


**Landscape 7 | Fitness Landscape of Dihydrofolate Reductase (DHFR) from Palmer *et al.***

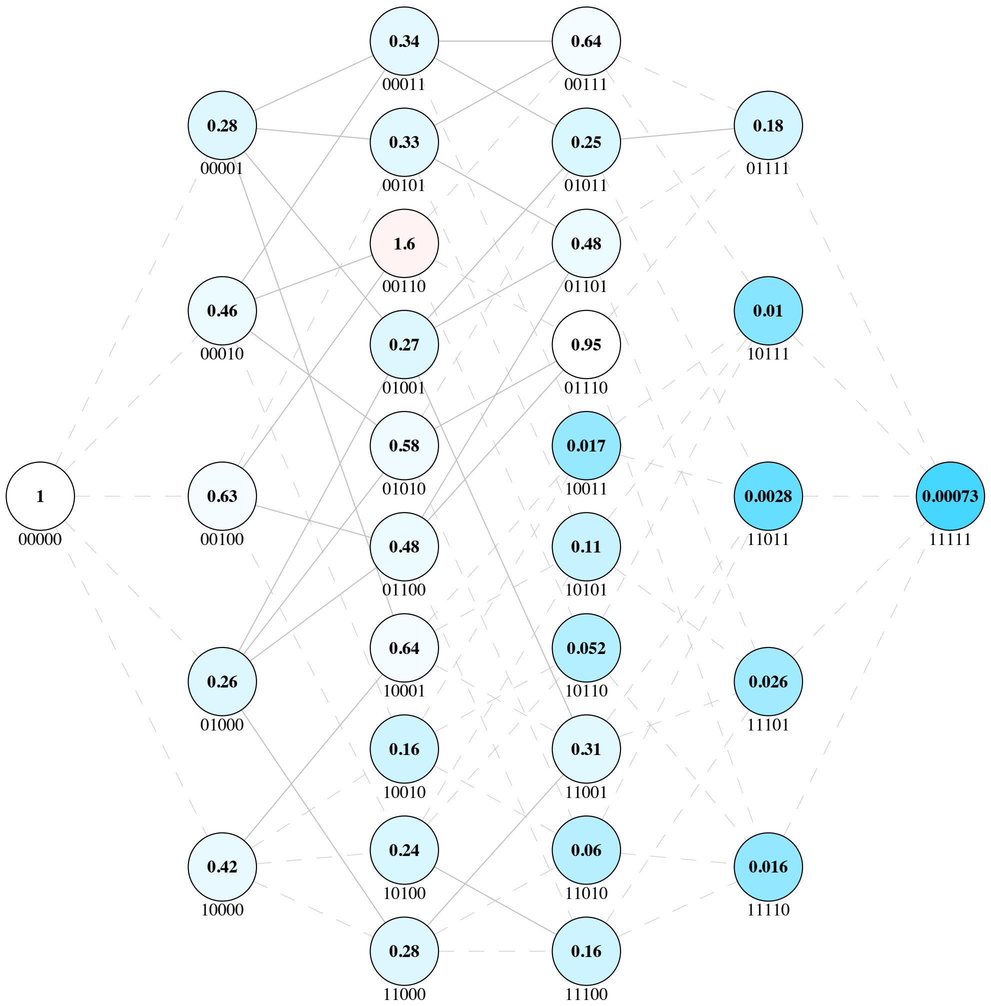


**Landscape 8 | Fitness Landscape of Dihydrofolate Reductase (DHFR) of $k_{cat}$ in the glycine trajectory from Tamer *et al.***

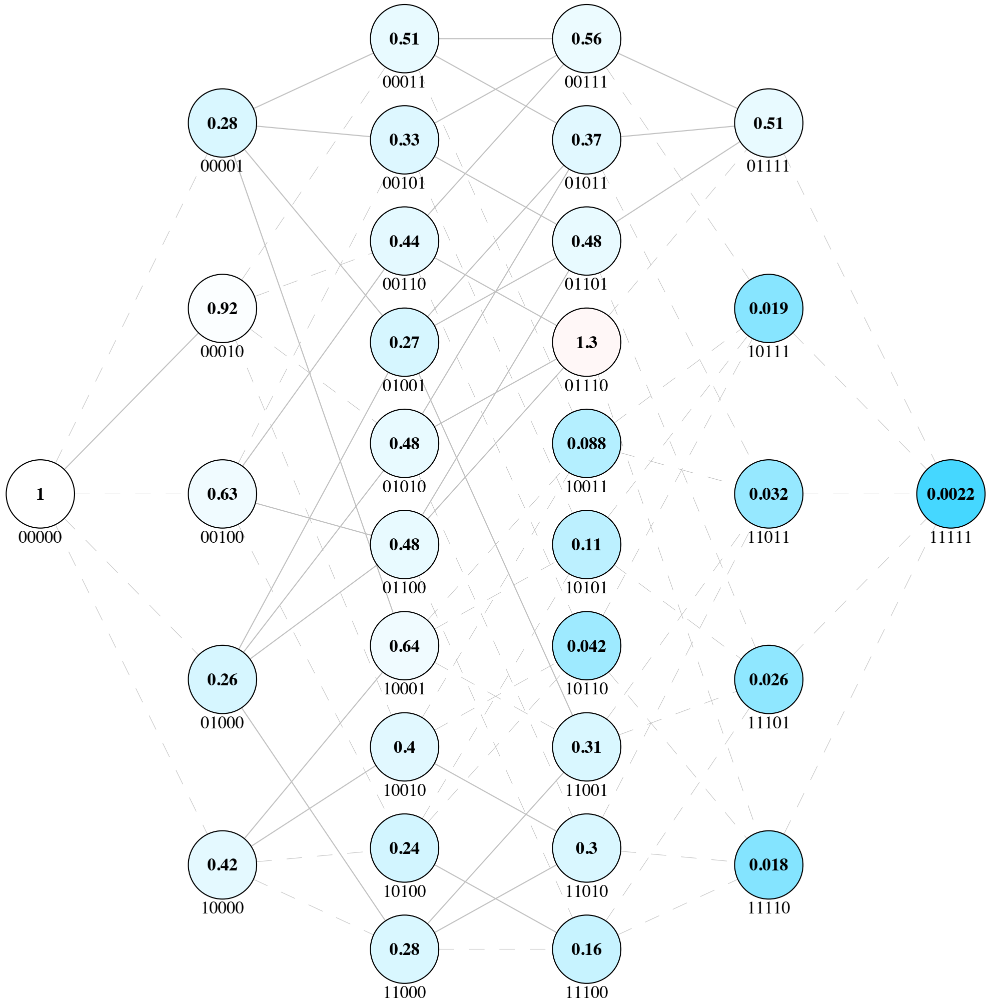


**Landscape 9 | Fitness Landscape of Dihydrofolate Reductase (DHFR) of $k_{cat}$ in the arginine trajectory from Tamer *et al.***

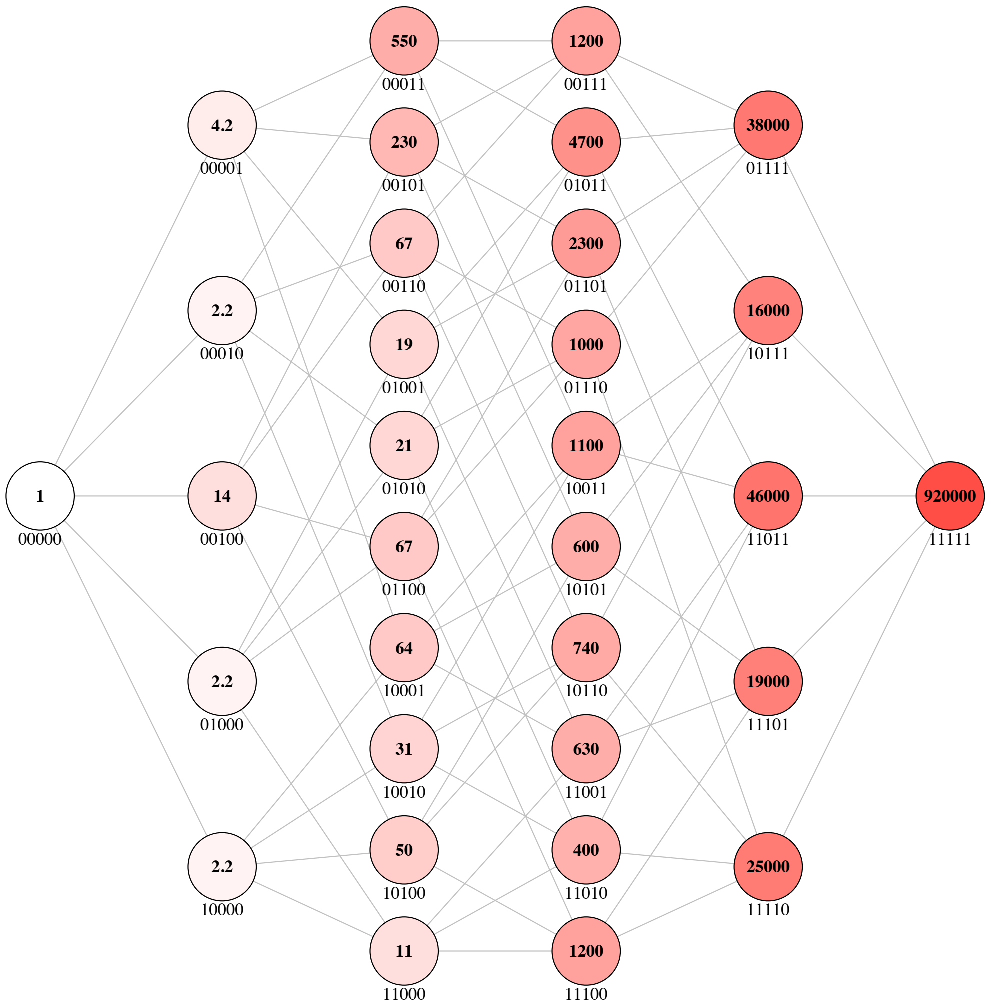


**Landscape 10 | Fitness Landscape of Dihydrofolate Reductase (DHFR) of $K_I$ in the glycine trajectory from Tamer *et al.***

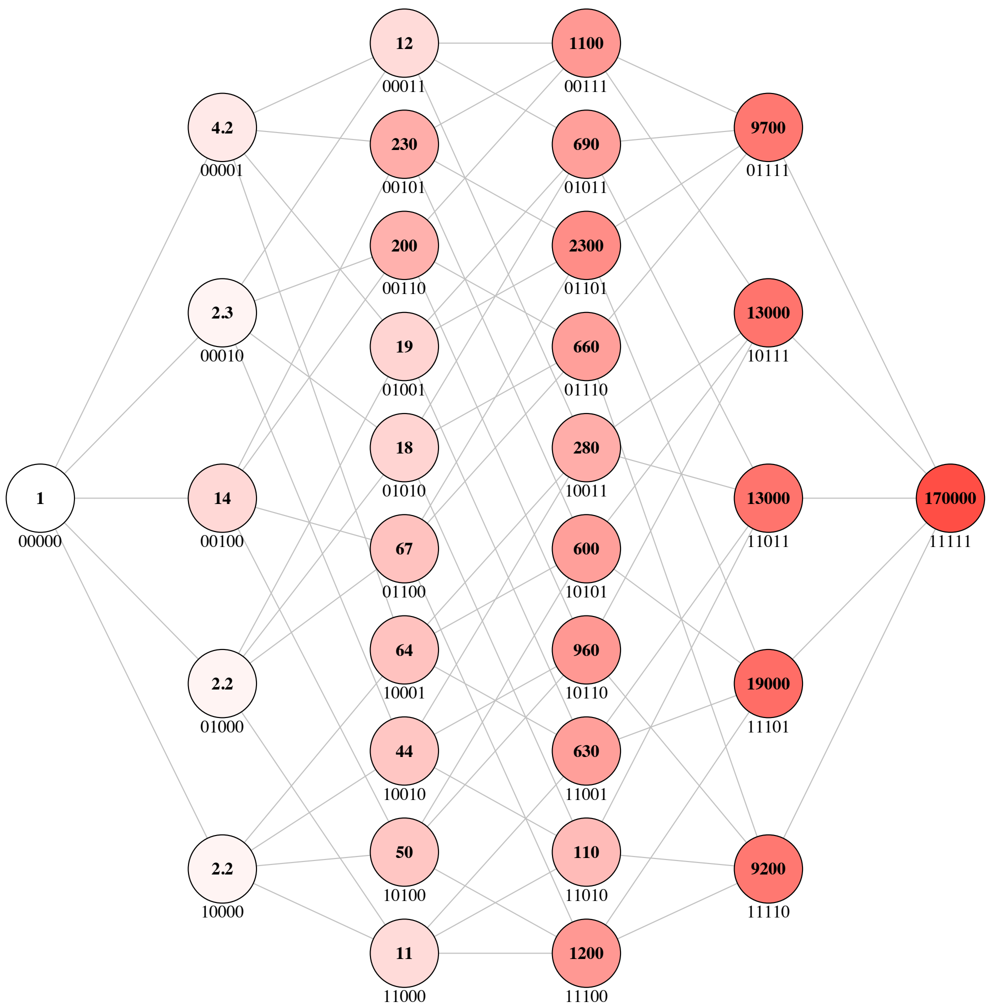


**Landscape 11 | Fitness Landscape of Dihydrofolate Reductase (DHFR) of $K_I$ in the arginine trajectory from Tamer *et al.***

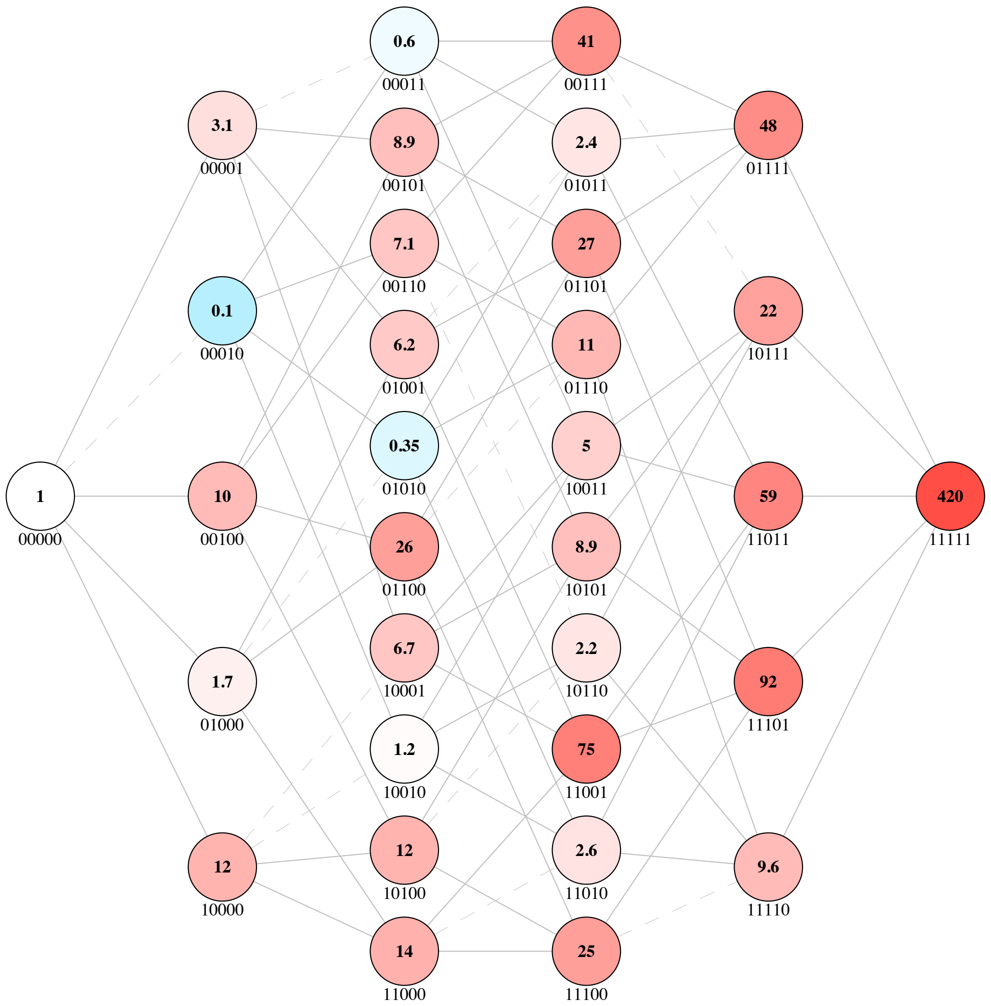


**Landscape 12 | Fitness Landscape of Methyl Parathion Hydrolase (MPH) for methyl parathion in calcium metal conditions from Anderson *et al.***

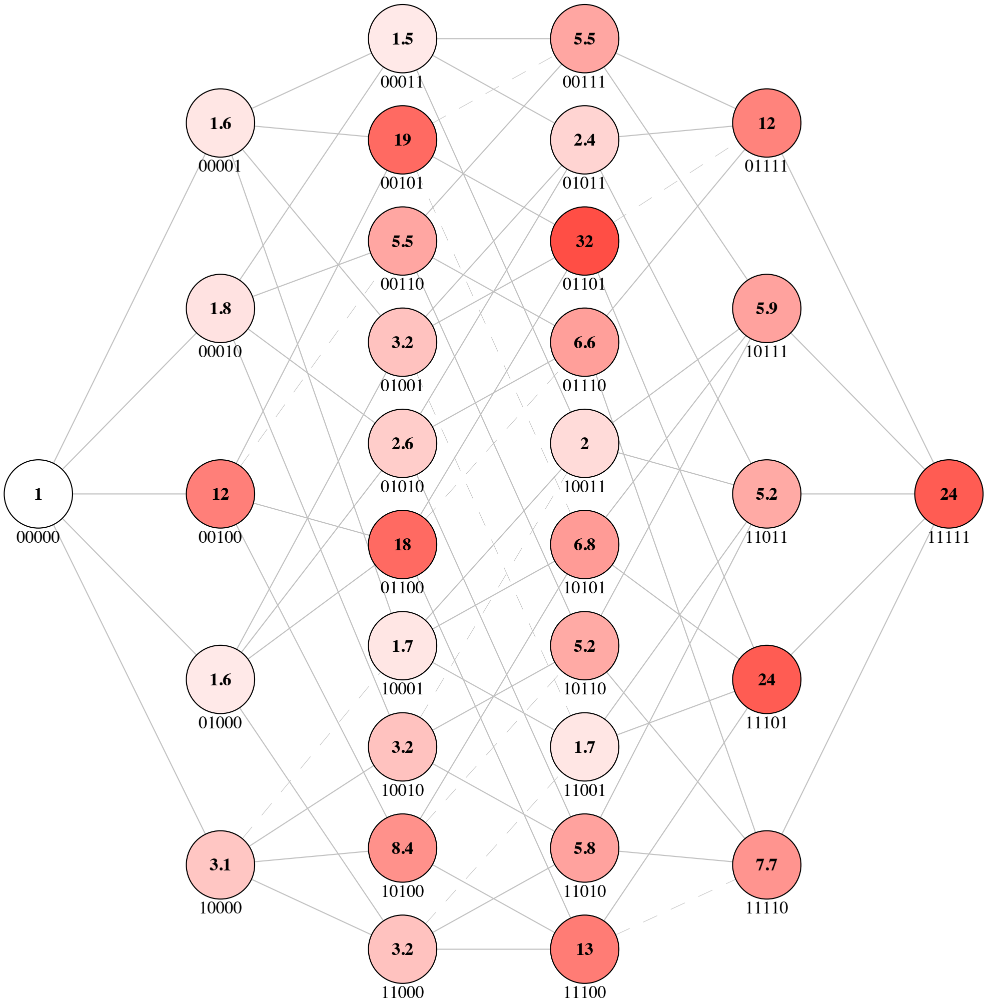


**Landscape 13 | Fitness Landscape of Methyl Parathion Hydrolase (MPH) for methyl parathion in cadmium metal conditions from Anderson *et al.***

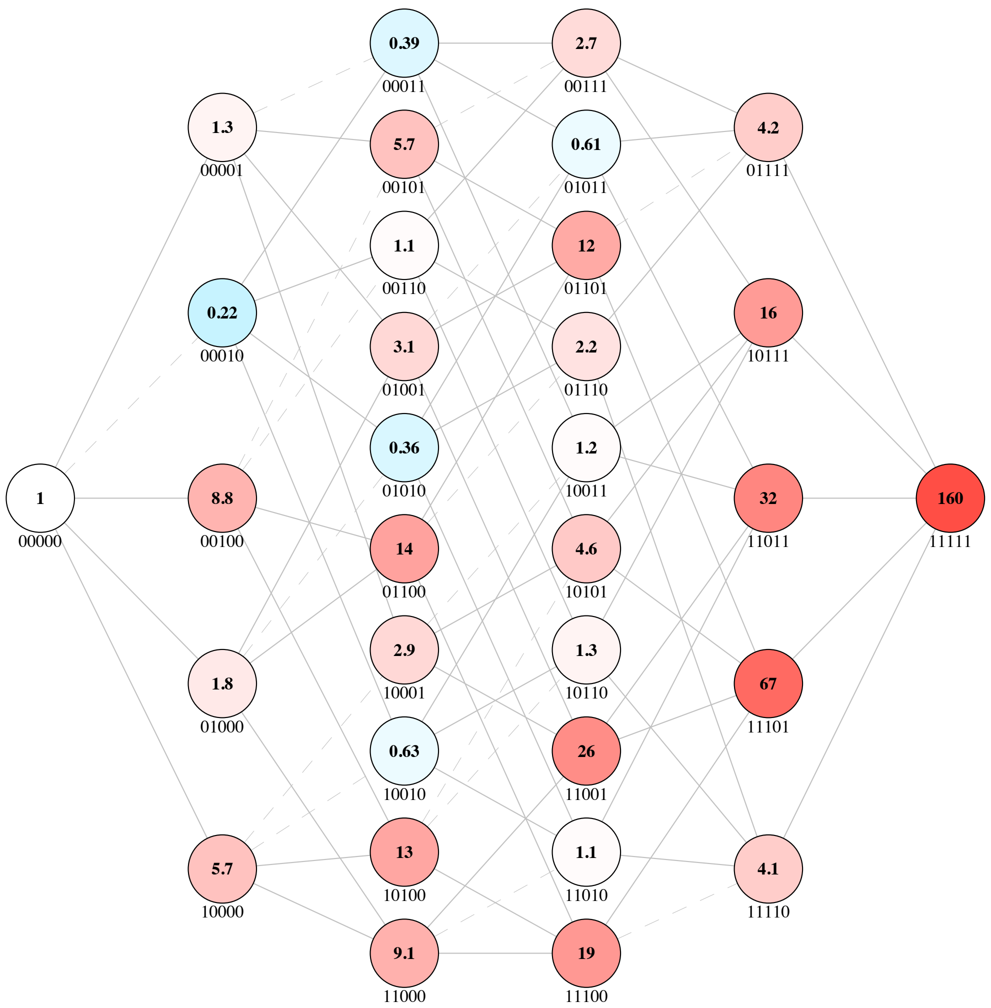


**Landscape 14 | Fitness Landscape of Methyl Parathion Hydrolase (MPH) for methyl parathion in cobalt metal conditions from Anderson *et al.***

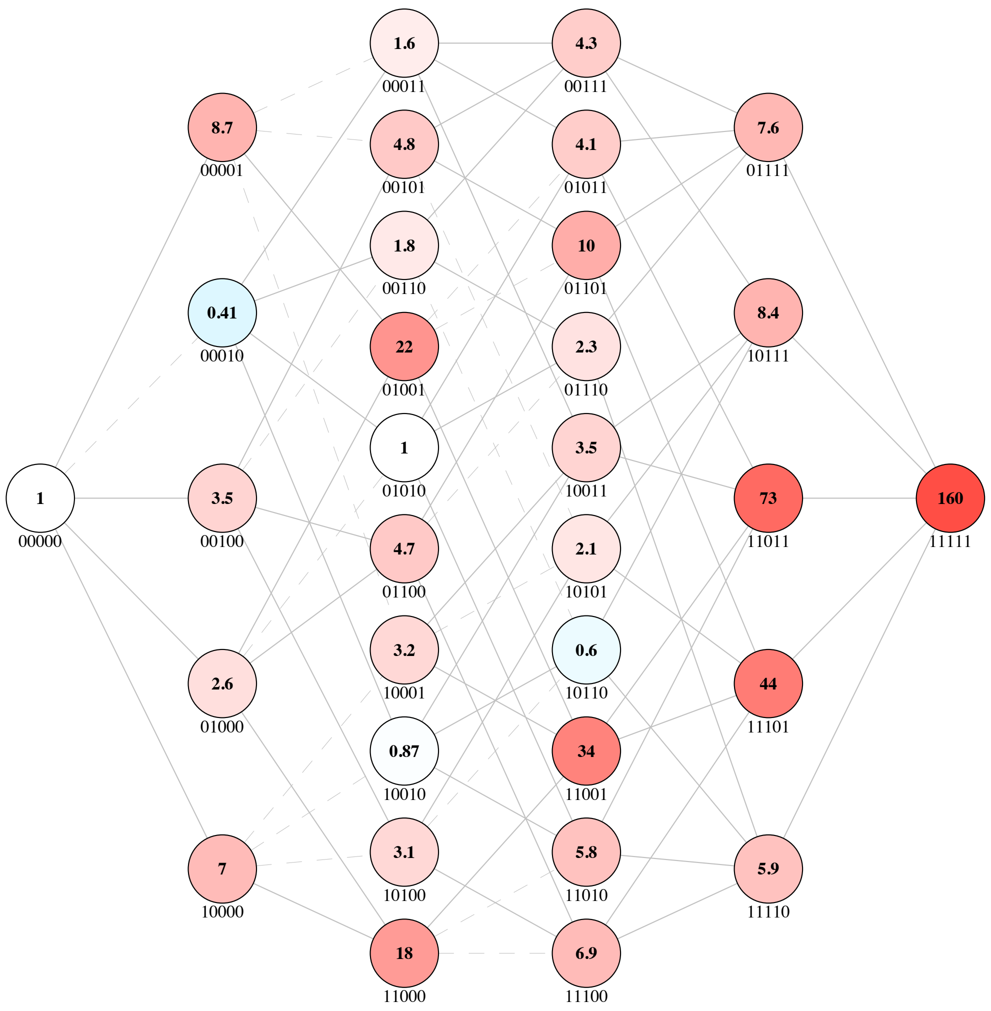


**Landscape 15 | Fitness Landscape of Methyl Parathion Hydrolase (MPH) for methyl parathion in copper metal conditions from Anderson *et al.***

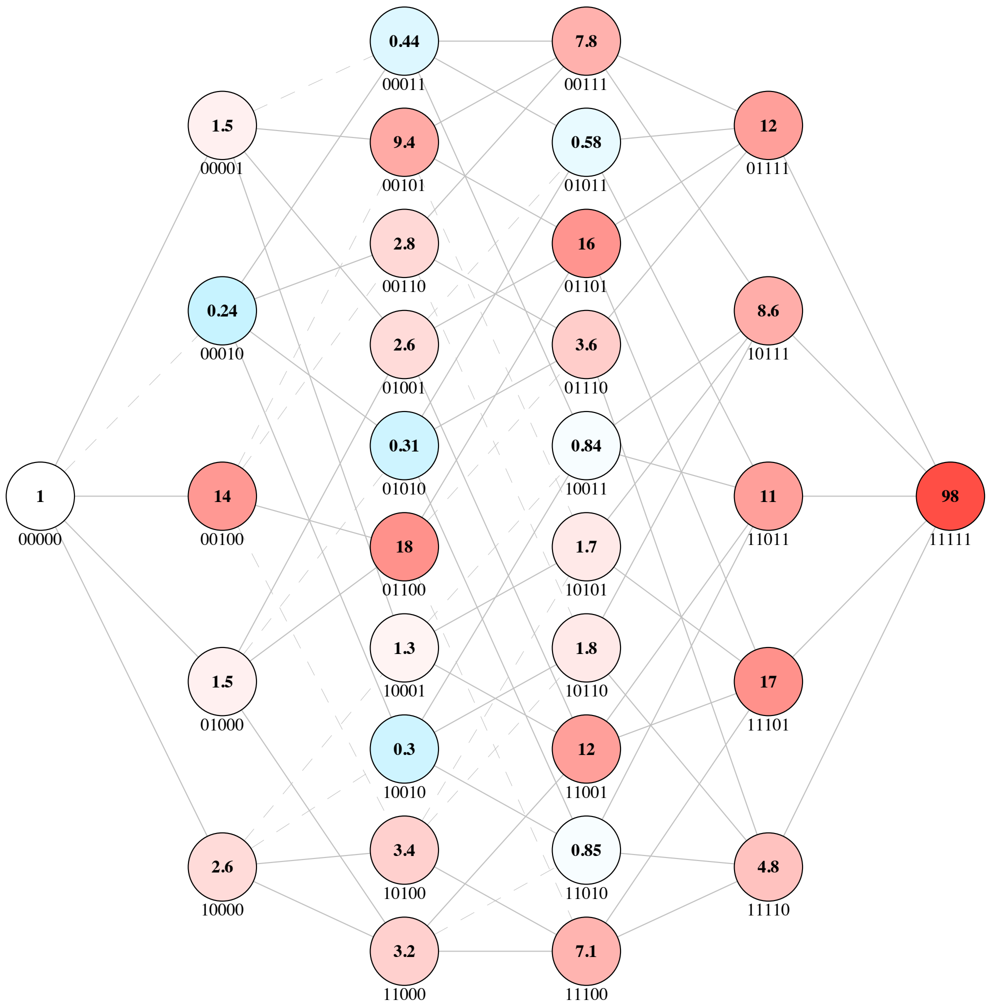


**Landscape 16 | Fitness Landscape of Methyl Parathion Hydrolase (MPH) for methyl parathion in magnesium metal conditions from Anderson *et al.***

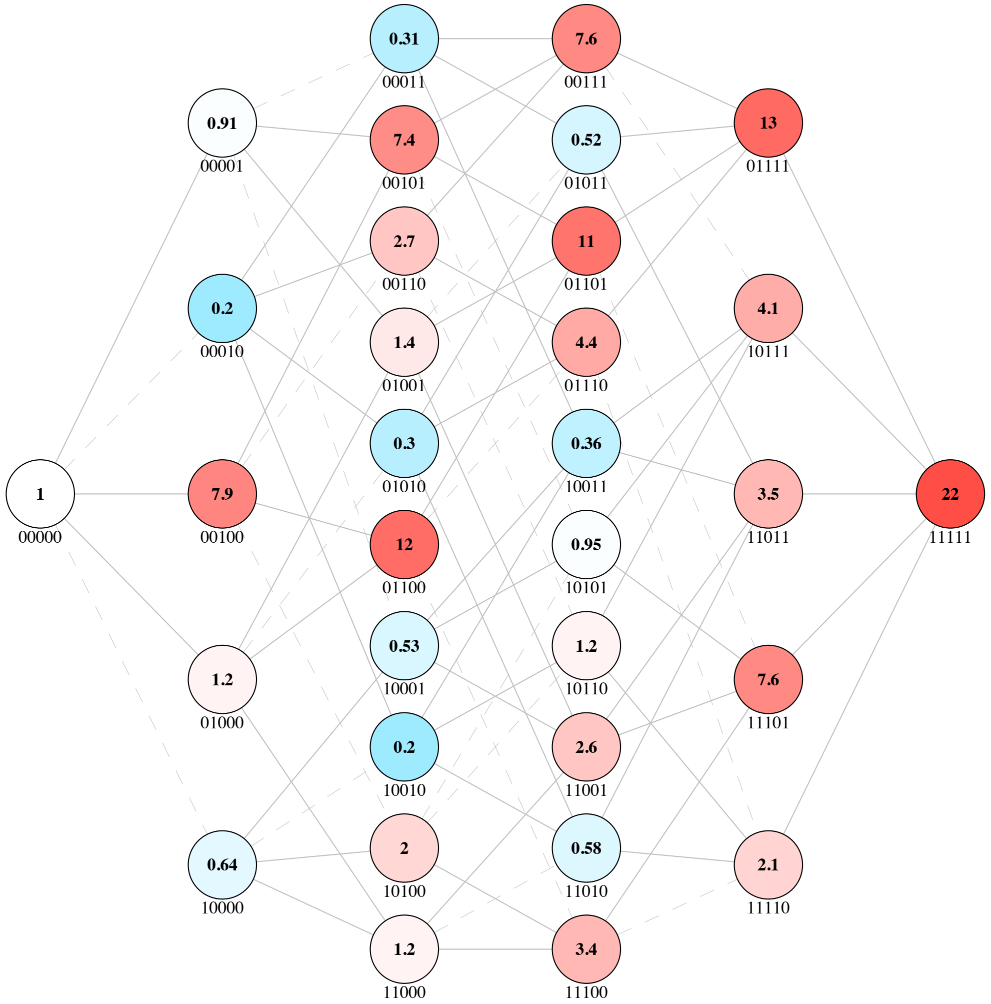


**Landscape 17 | Fitness Landscape of Methyl Parathion Hydrolase (MPH) for methyl parathion in manganese metal conditions from Anderson *et al.***

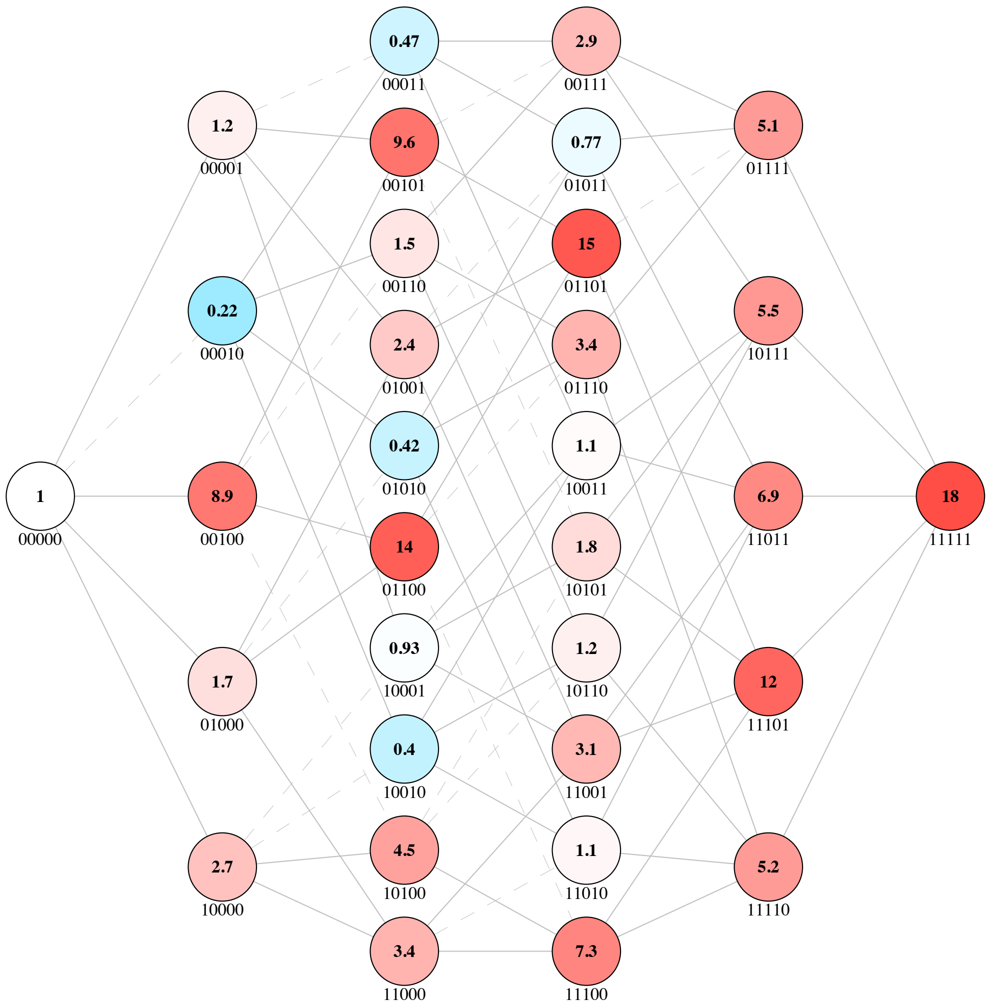


**Landscape 18 | Fitness Landscape of Methyl Parathion Hydrolase (MPH) for methyl parathion in nickel metal conditions from Anderson *et al.***

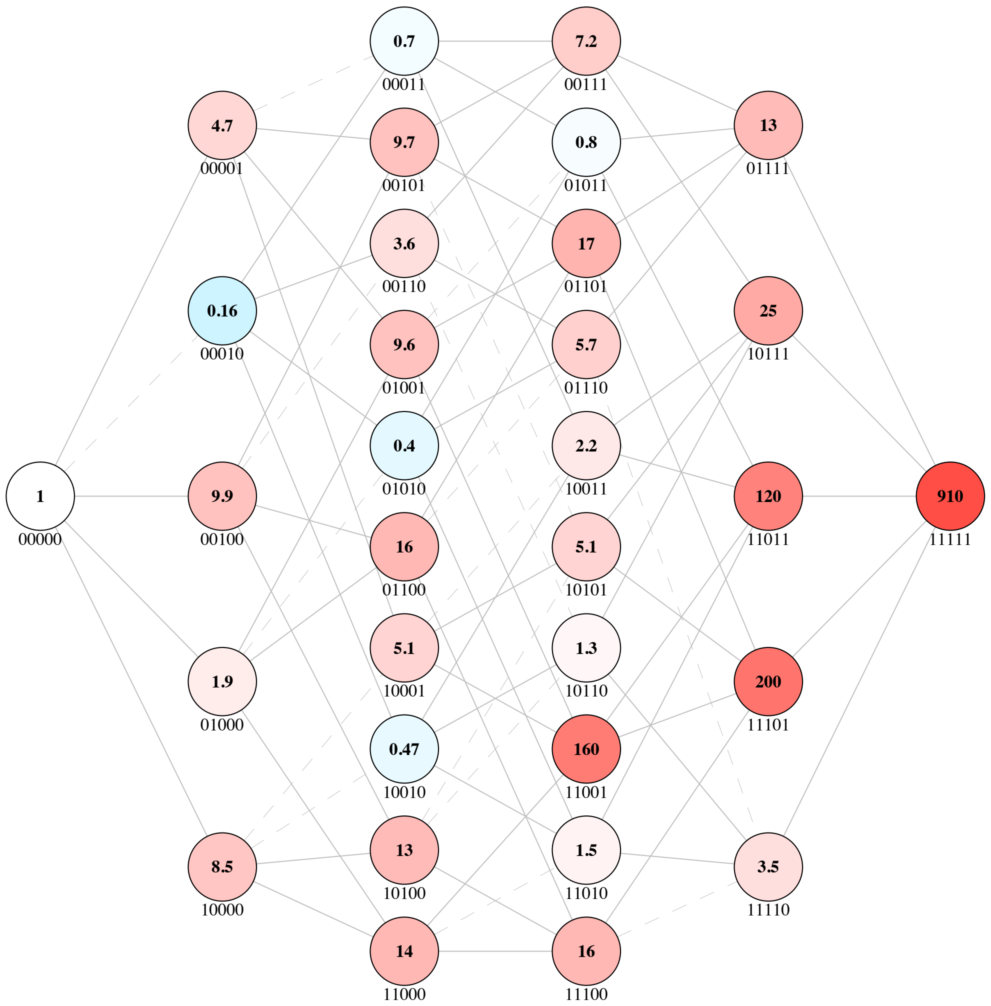


**Landscape 19 | Fitness Landscape of Methyl Parathion Hydrolase (MPH) for methyl parathion in zinc metal conditions from Anderson *et al.***

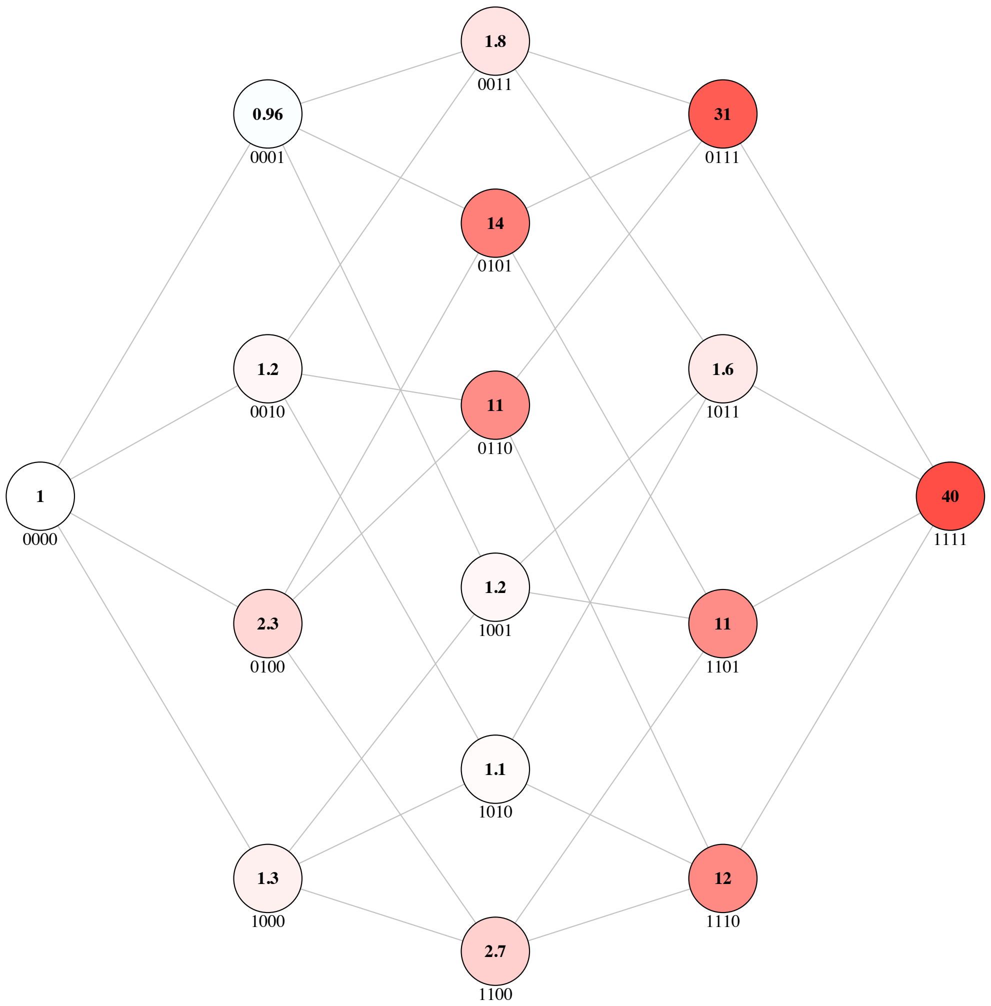


**Landscape 20 | Fitness Landscape of beta lactamase OXA-48 for ceftazidime hydrolysis in trajectory 1 from Fröhlich *et al.***

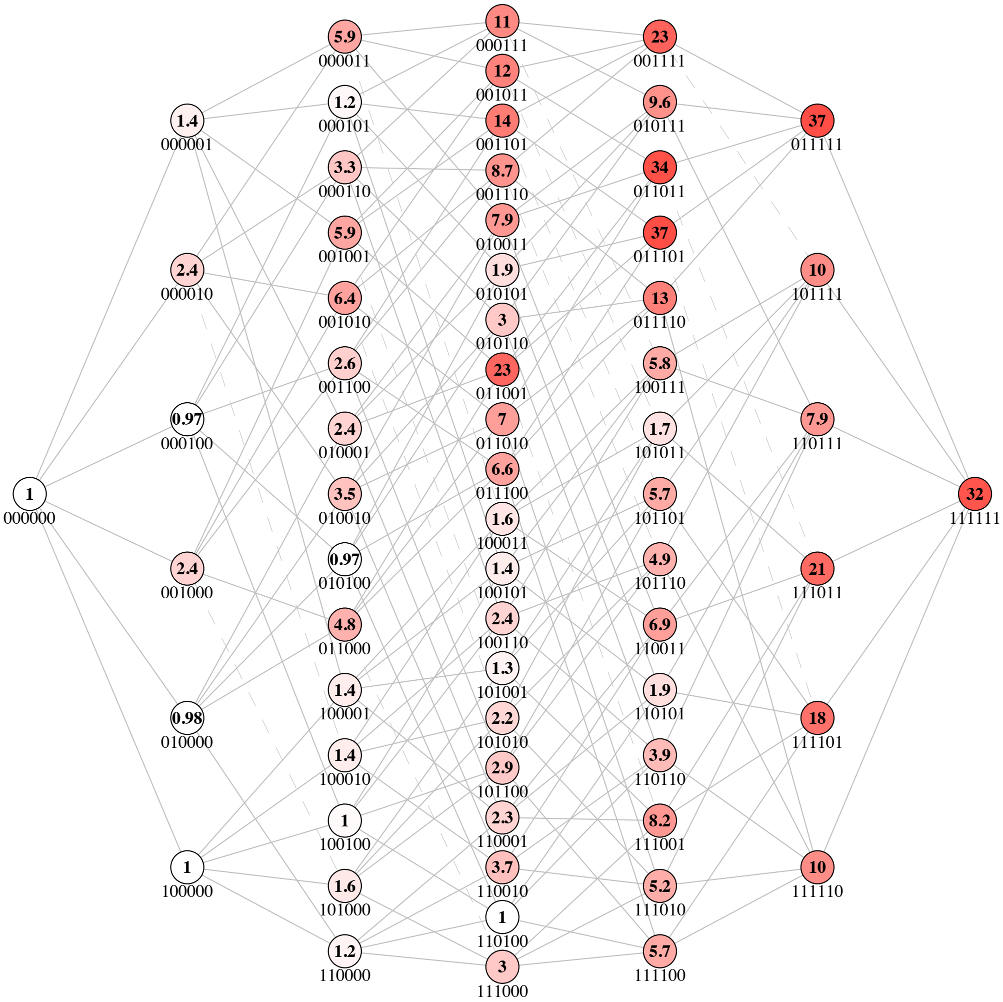


**Landscape 21** | **Fitness Landscape of beta lactamase OXA-48 for ceftazidime hydrolysis in trajectory 2 from Fröhlich *et al.***

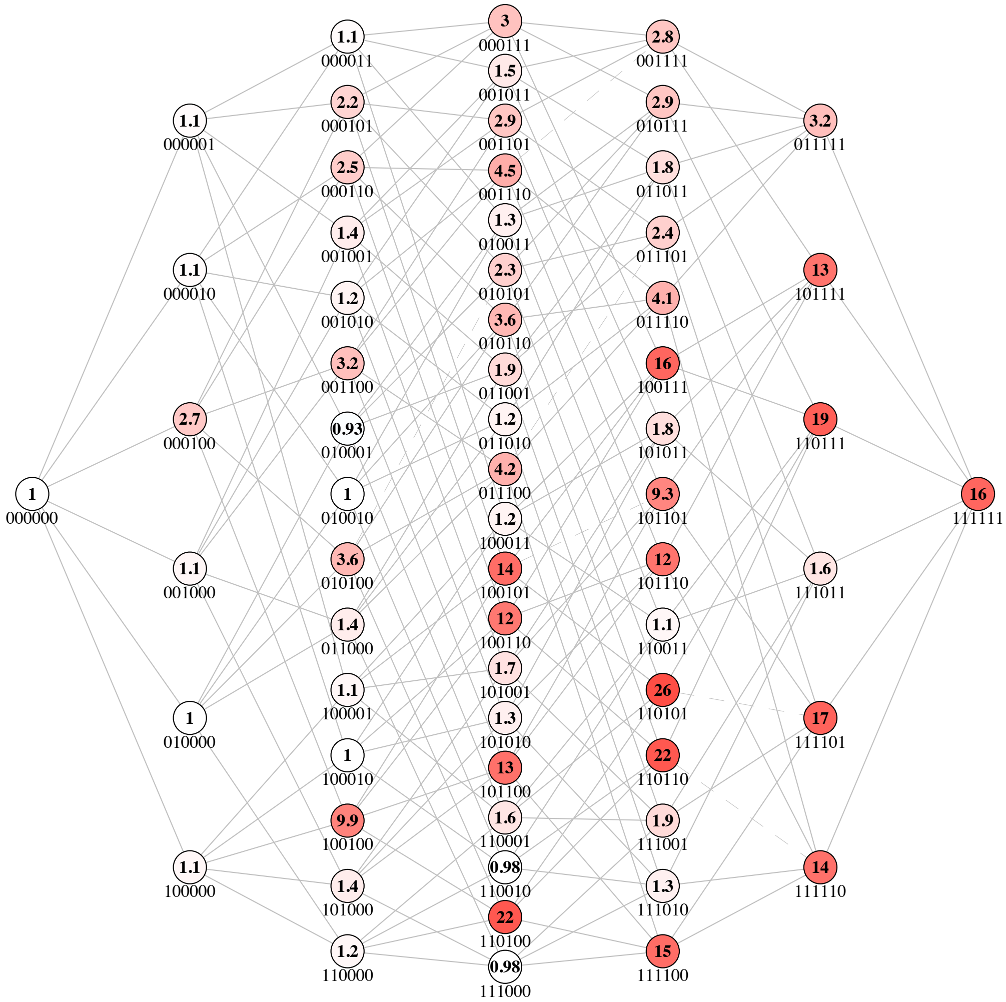


**Landscape 22 | Fitness Landscape of beta lactamase OXA-48 for ceftazidime hydrolysis in trajectory 3 from Fröhlich *et al.***

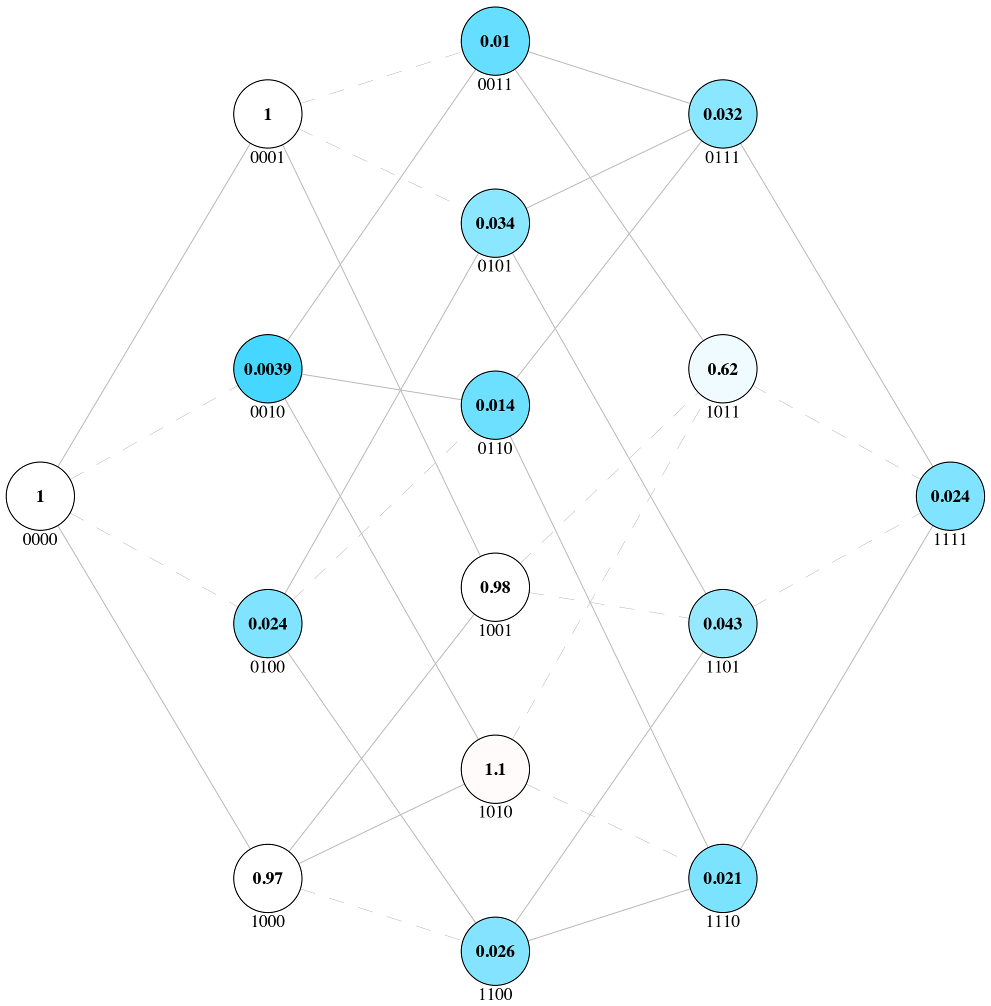


**Landscape 23** | **Fitness Landscape of beta lactamase OXA-48 for piperacillin hydrolysis in trajectory 1 from Fröhlich *et al.***

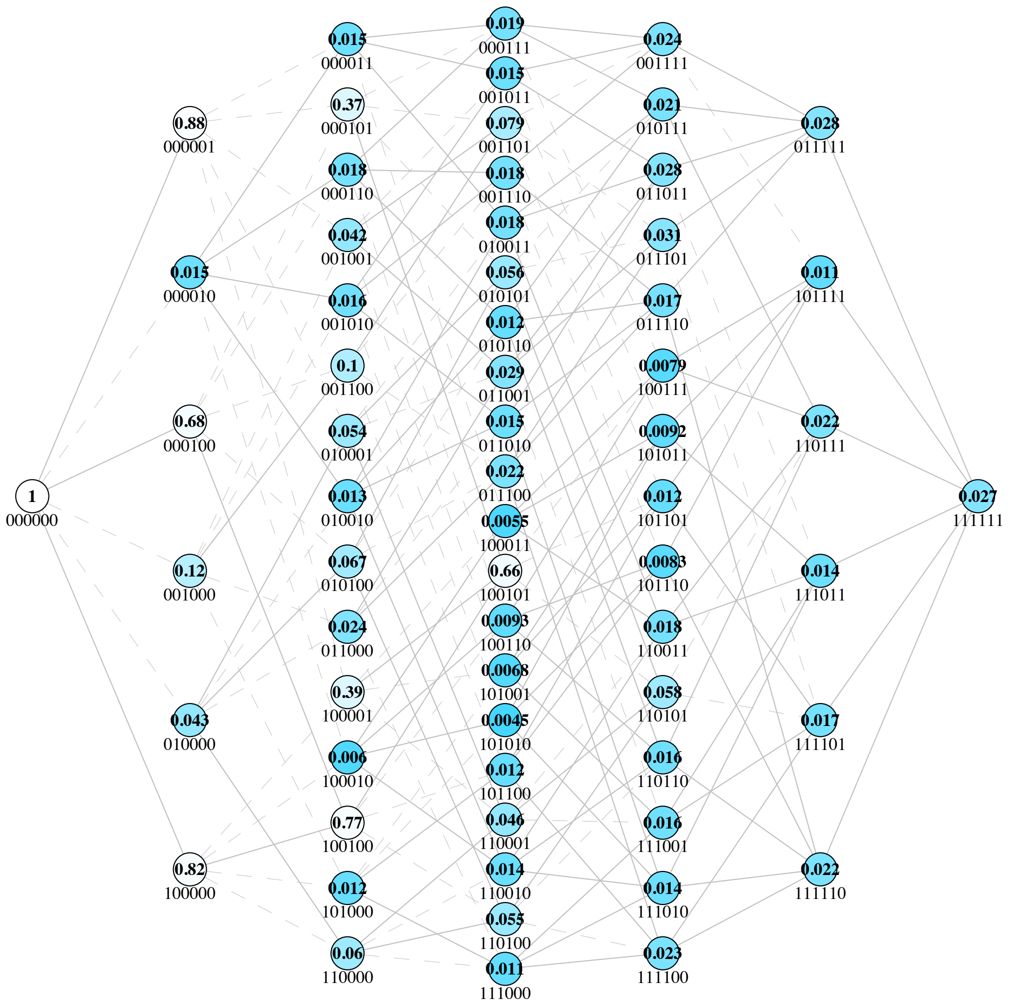


**Landscape 24 | Fitness Landscape of beta lactamase OXA-48 for piperacillin hydrolysis in trajectory 2 from Fröhlich *et al.***

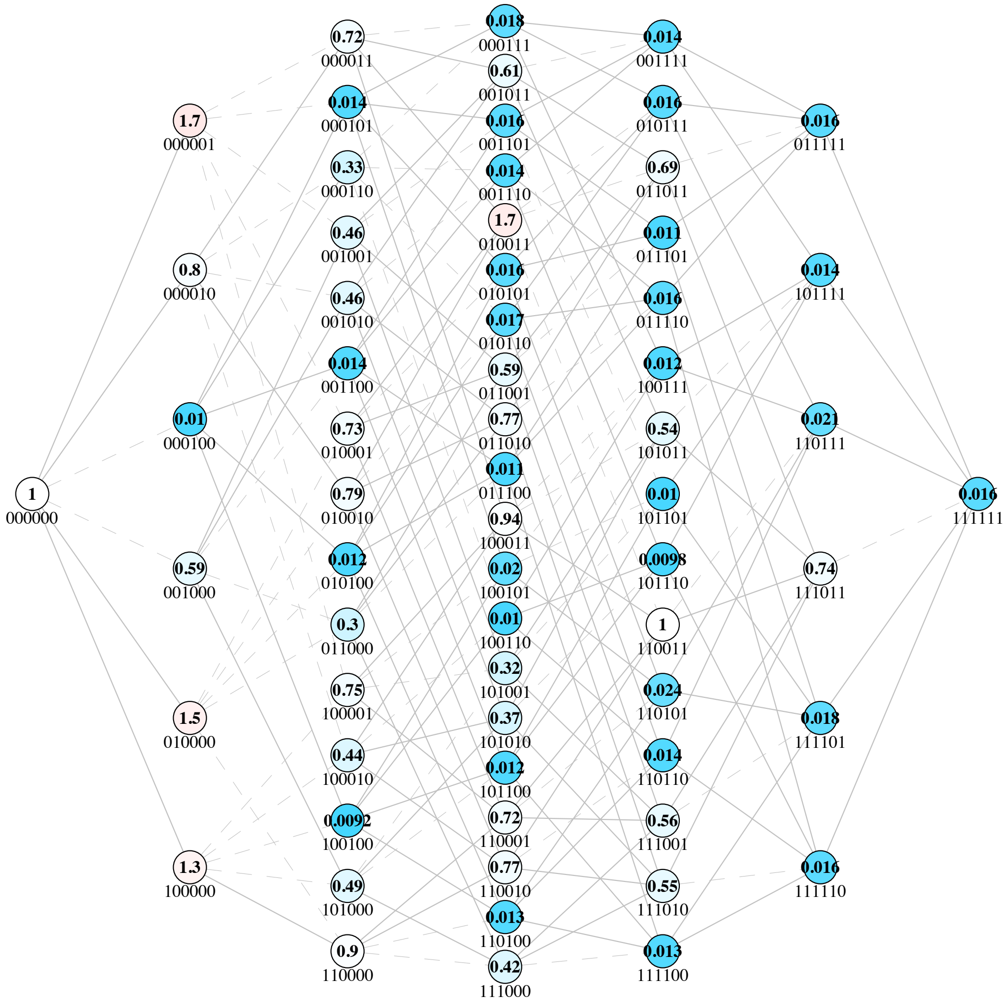


**Landscape 25 | Fitness Landscape of beta lactamase OXA-48 for piperacillin hydrolysis in trajectory 3 from Fröhlich *et al.***

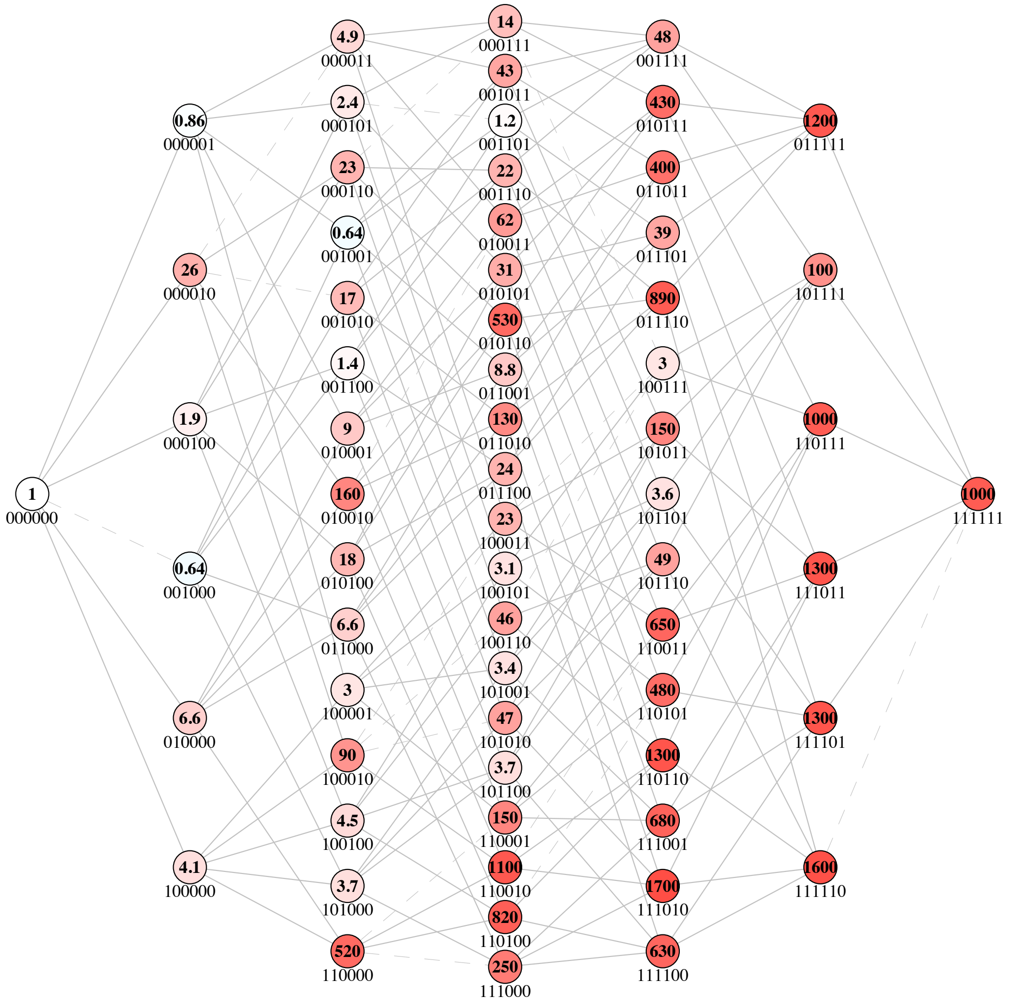


**Landscape 26 | Fitness Landscape of phosphotriesterase (PTE) for 2-napthyl hexanoate (2NH) hydrolysis from this publication**

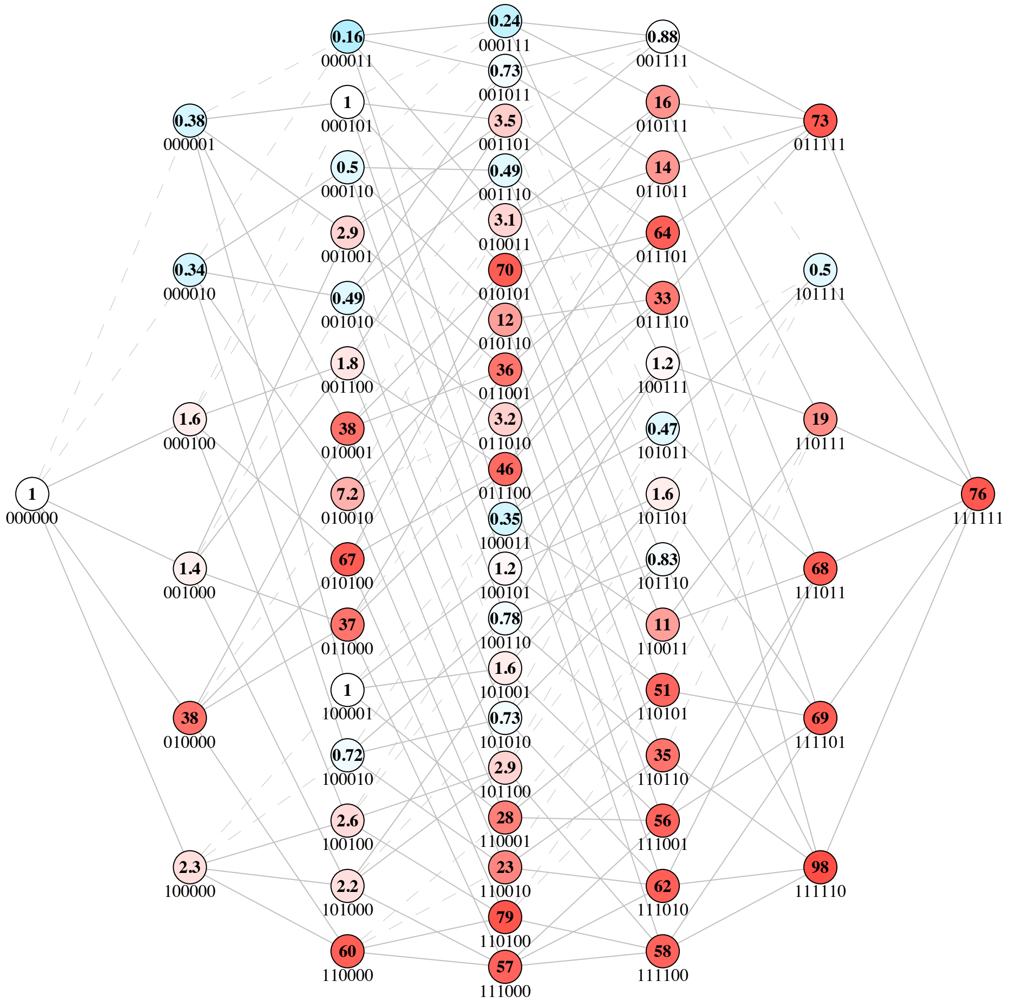


**Landscape 27** | **Fitness Landscape of phosphotriesterase (PTE) for butyrate hydrolysis from Miton *et al.***

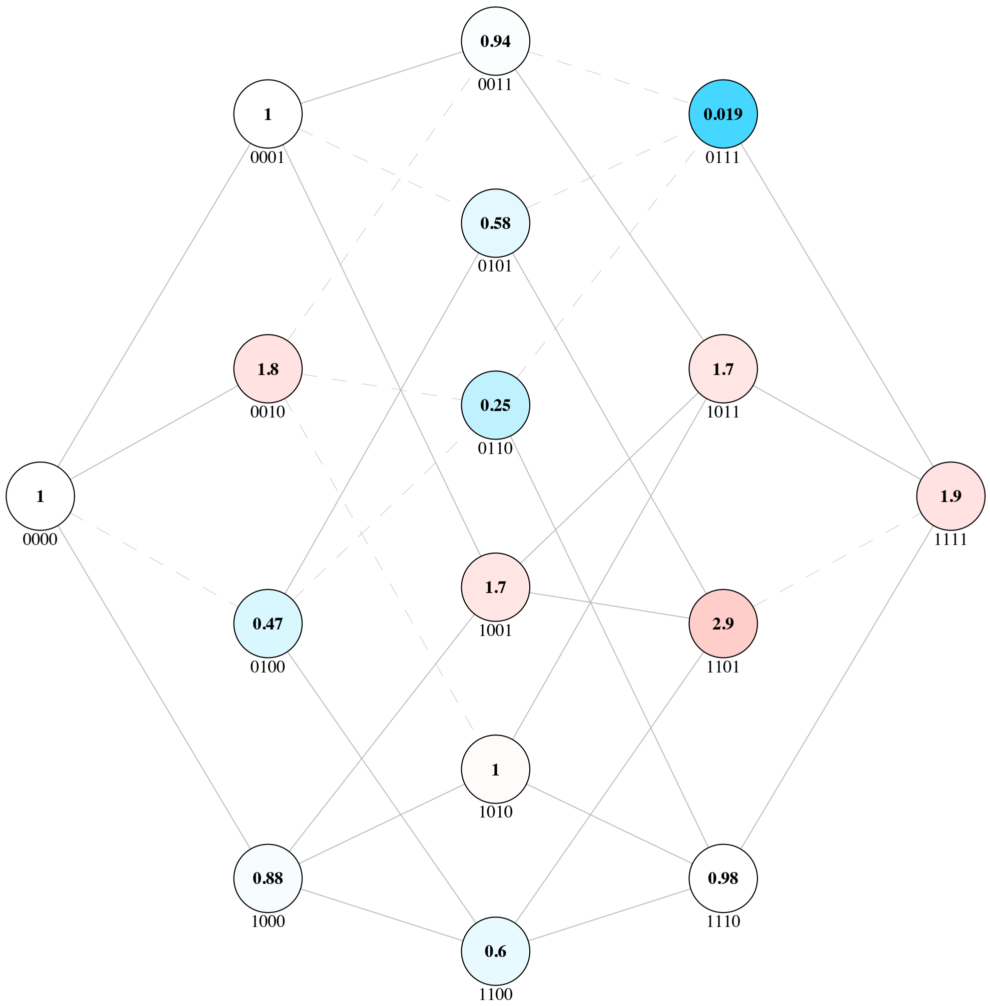


**Landscape 28 | Fitness Landscape of beta lactamase TEM in AM antibiotic from Mira *et al.***

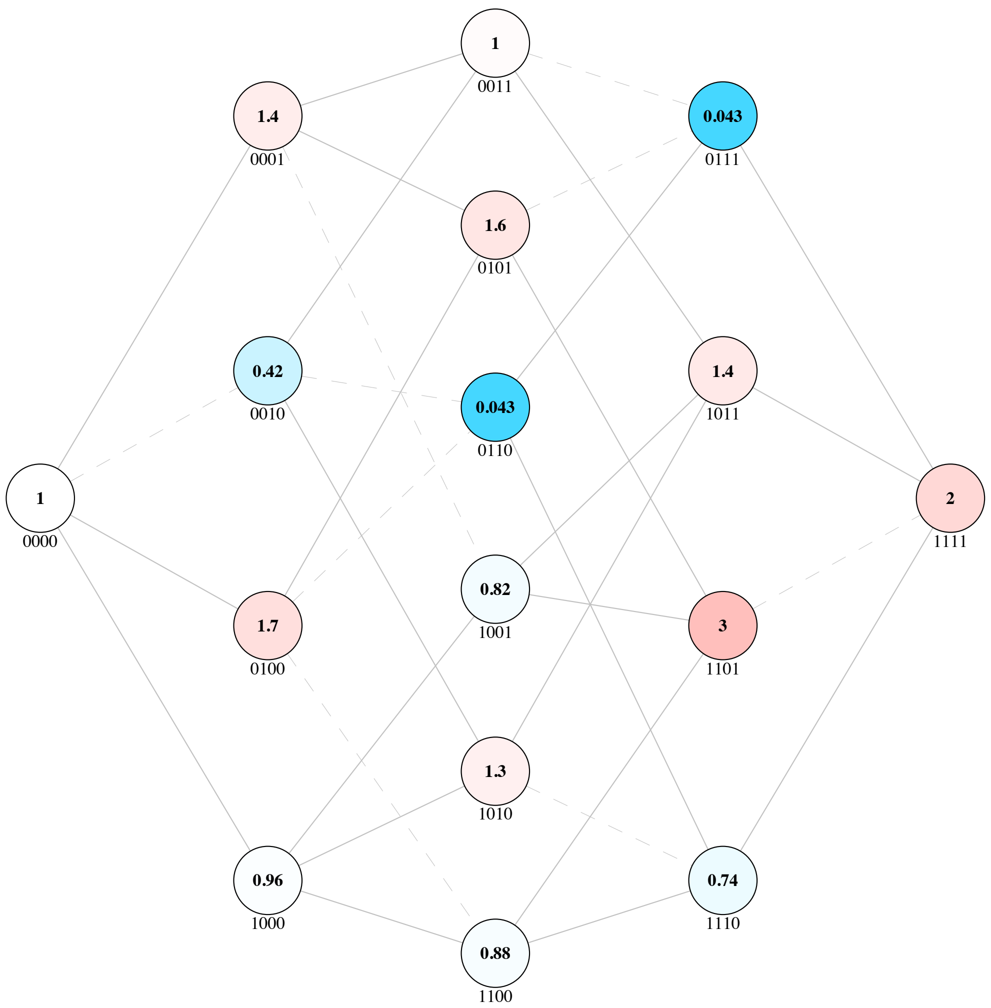


**Landscape 29 | Fitness Landscape of beta lactamase TEM in AMC antibiotic from Mira *et al.***

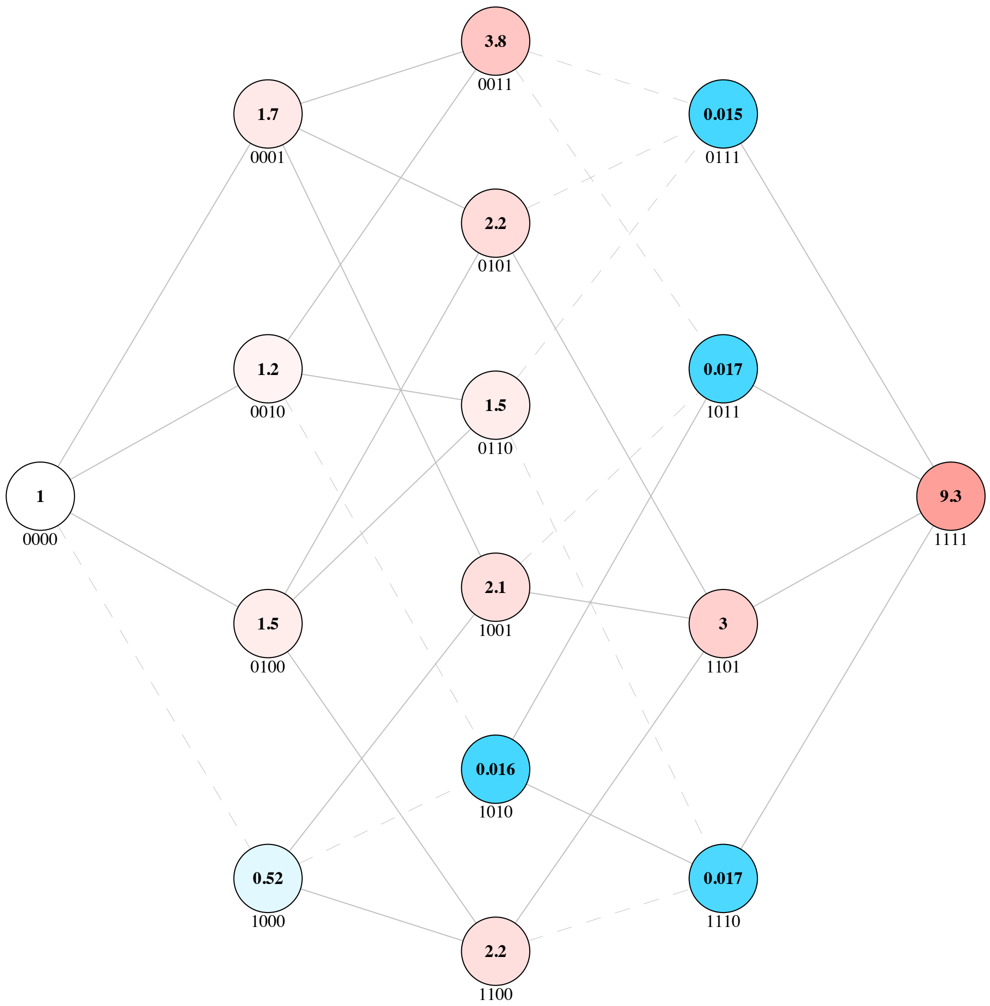


**Landscape 30 | Fitness Landscape of beta lactamase TEM in AMP antibiotic from Mira *et al.***

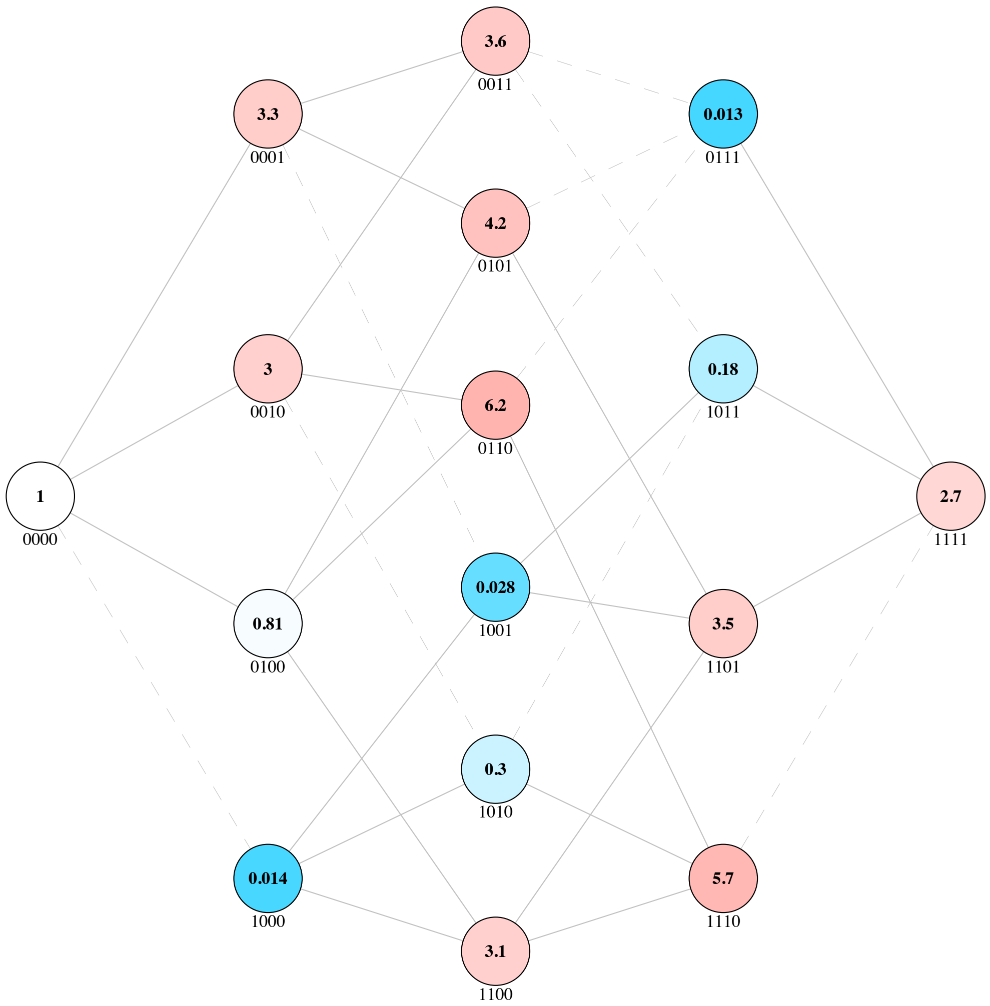


**Landscape 31 | Fitness Landscape of beta lactamase TEM in CAZ antibiotic from Mira *et al.***

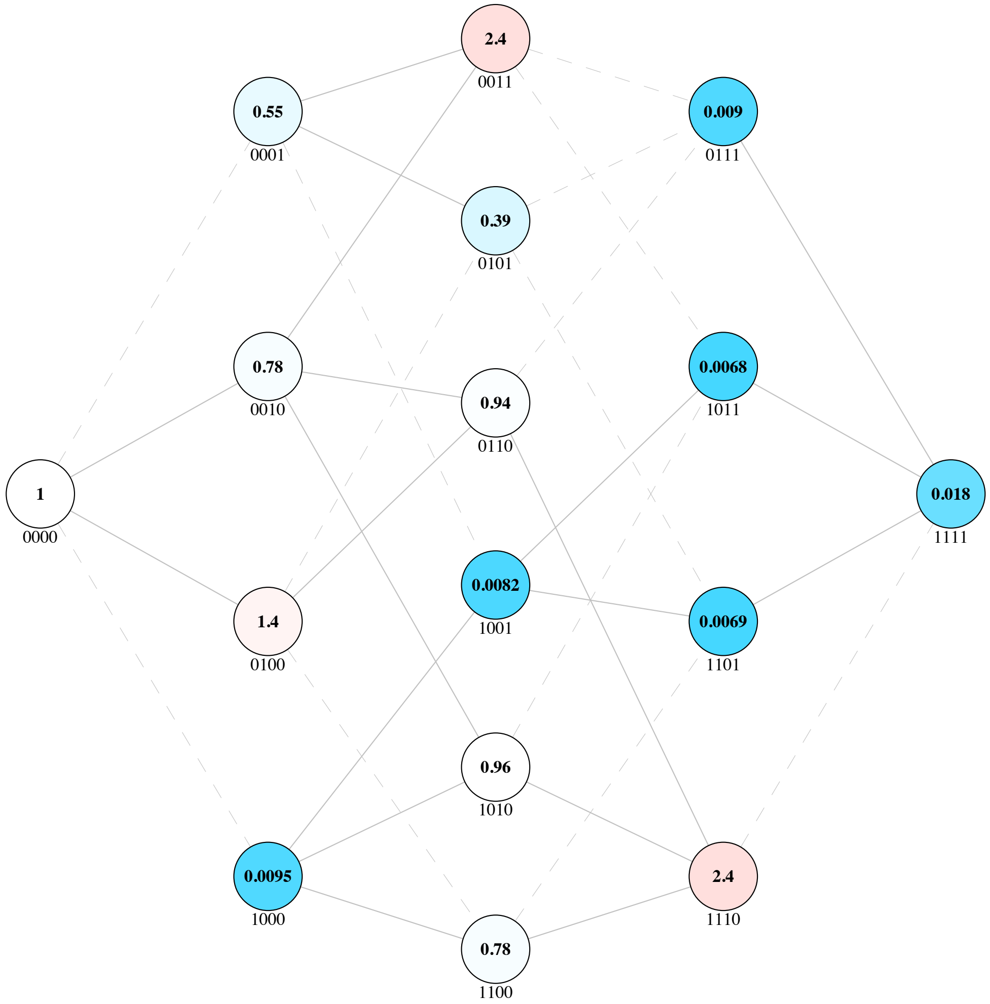


**Landscape 32 | Fitness Landscape of beta lactamase TEM in CEC antibiotic from Mira *et al.***

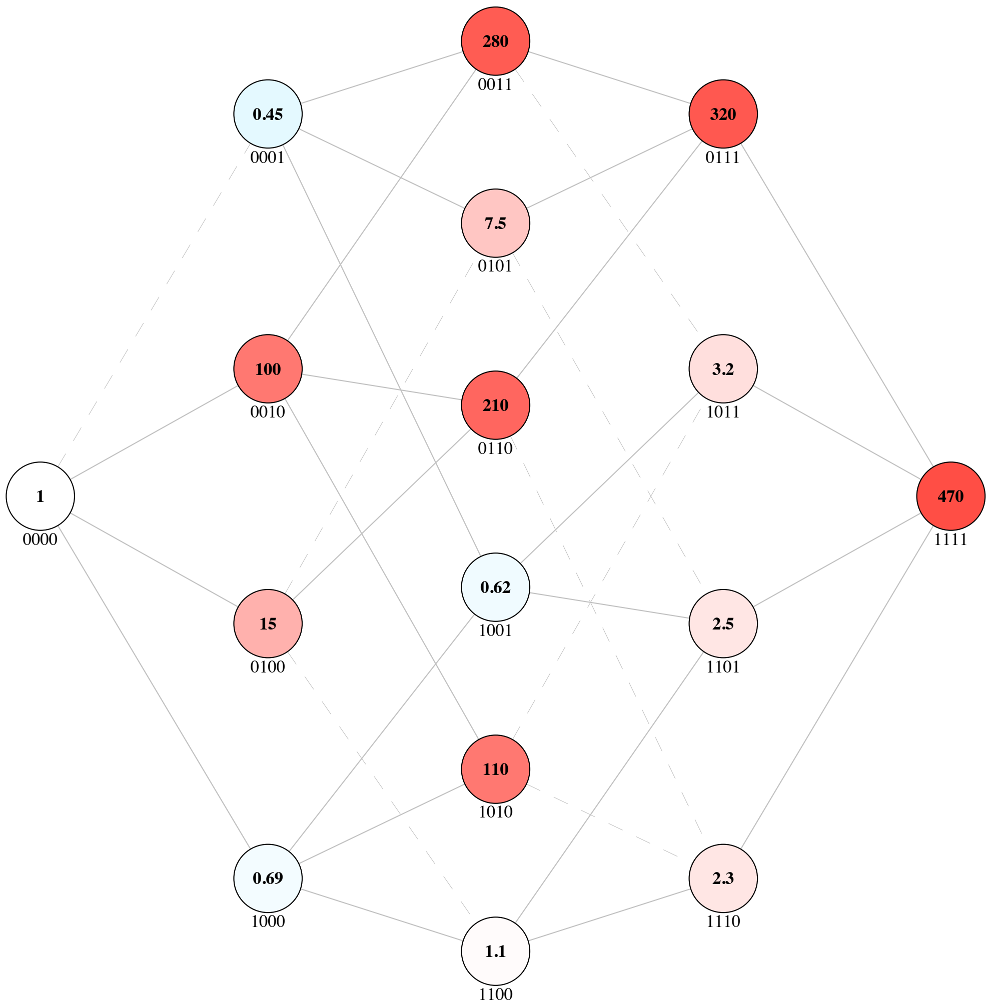


**Landscape 33 | Fitness Landscape of beta lactamase TEM in CPD antibiotic from Mira *et al.***

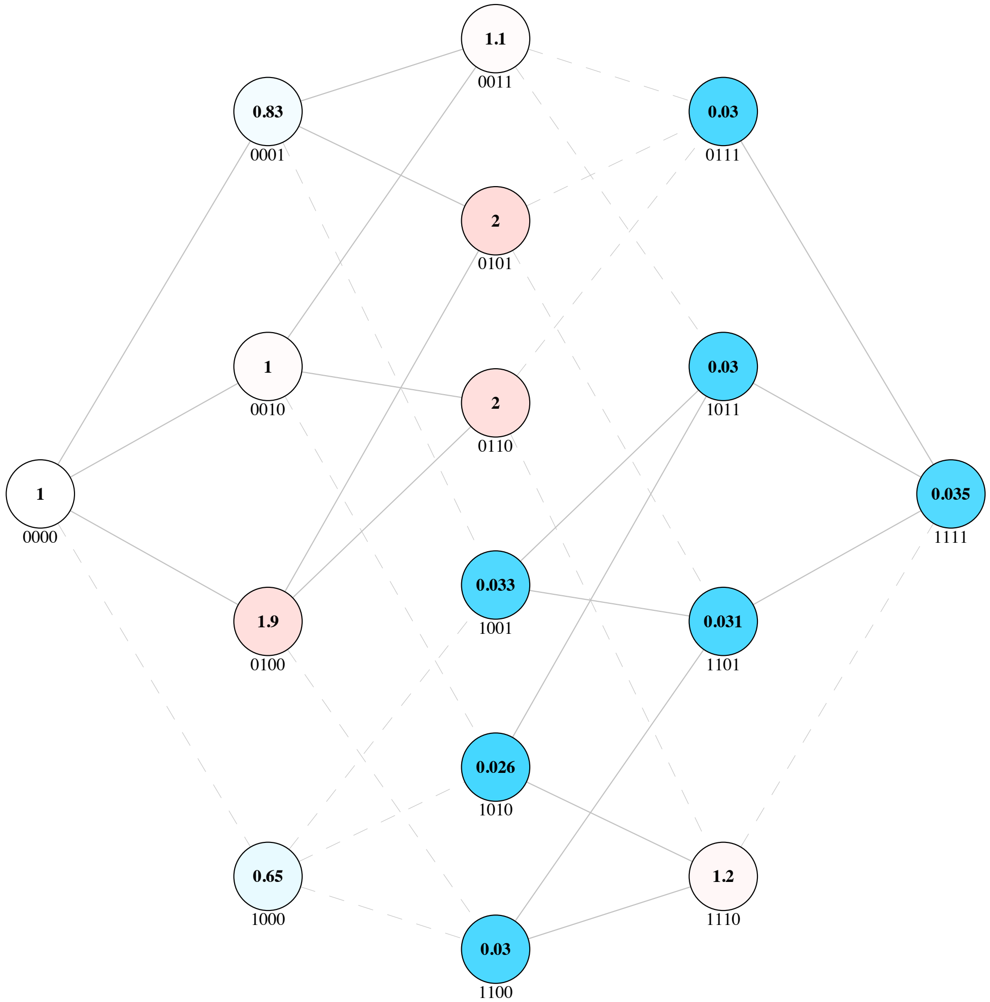


**Landscape 34 | Fitness Landscape of beta lactamase TEM in CPR antibiotic from Mira *et al.***

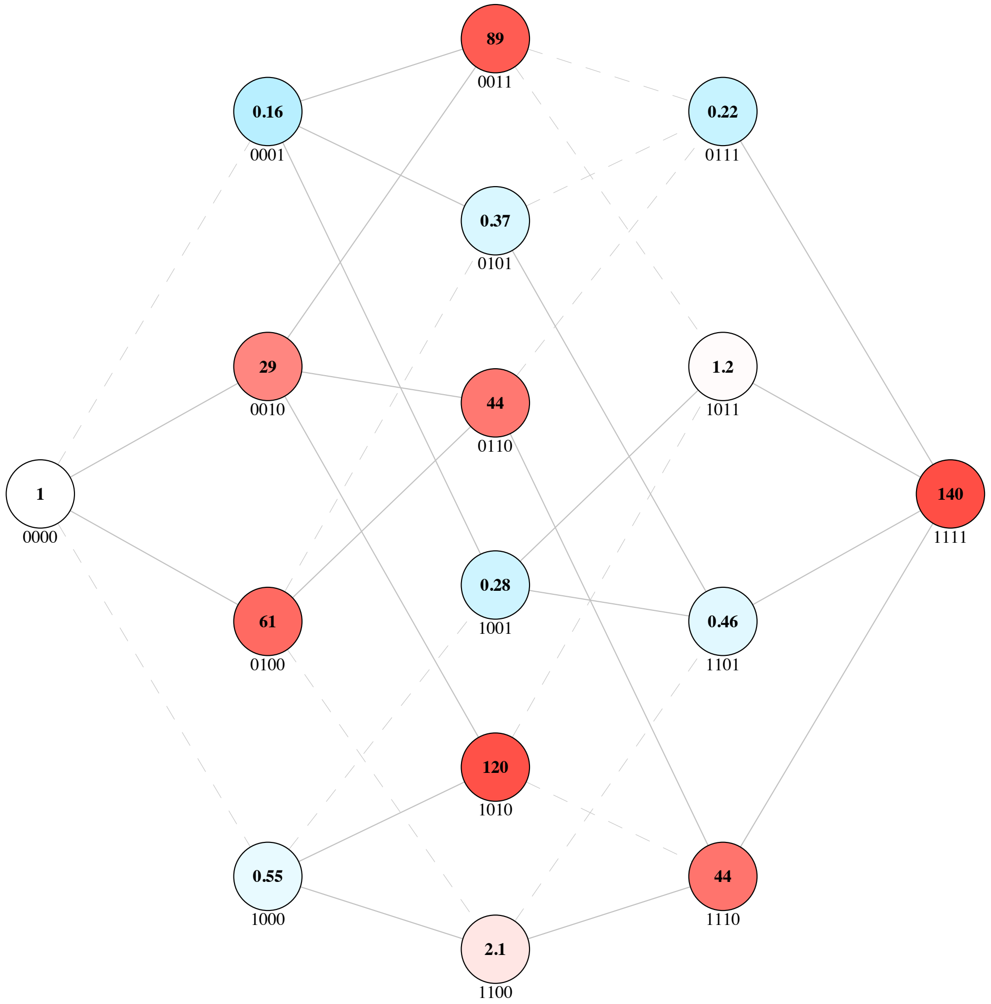


**Landscape 35 | Fitness Landscape of beta lactamase TEM in CRO antibiotic from Mira *et al.***

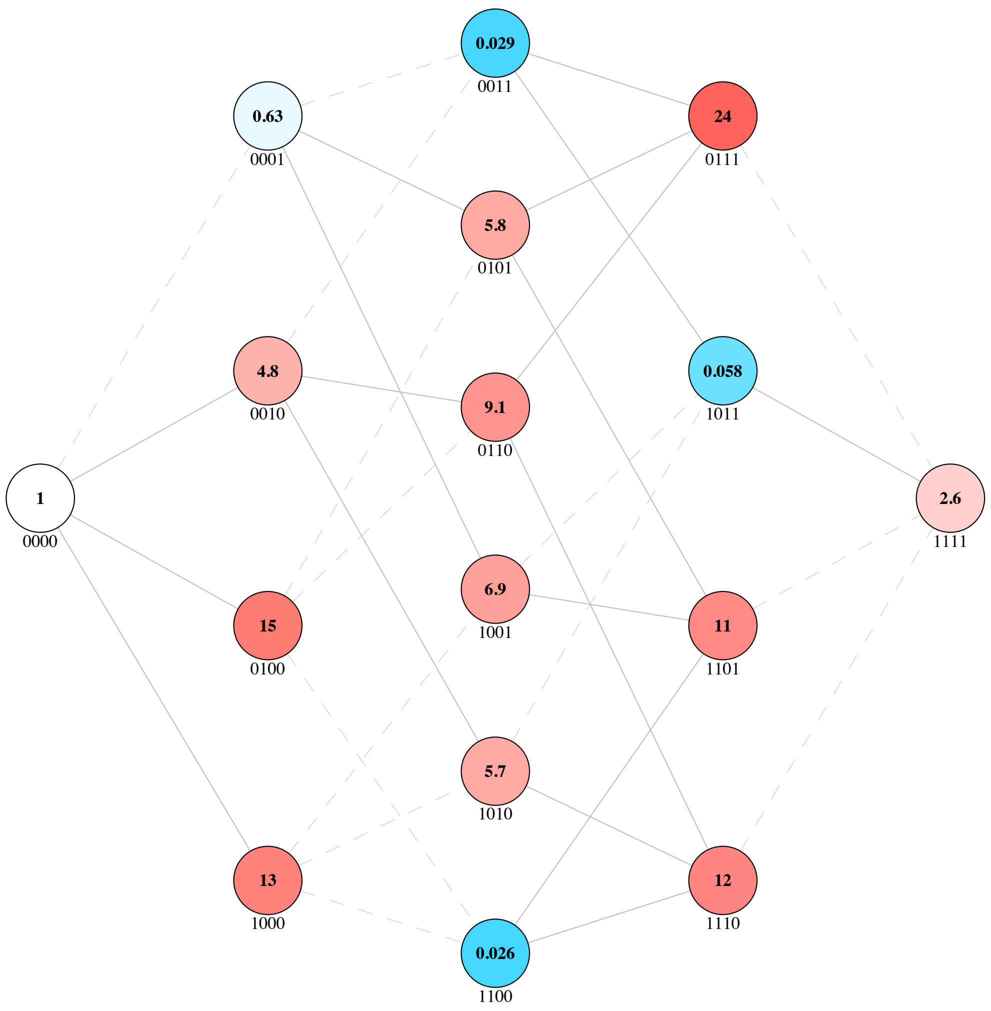


**Landscape 36 | Fitness Landscape of beta lactamase TEM in CTT antibiotic from Mira *et al.***

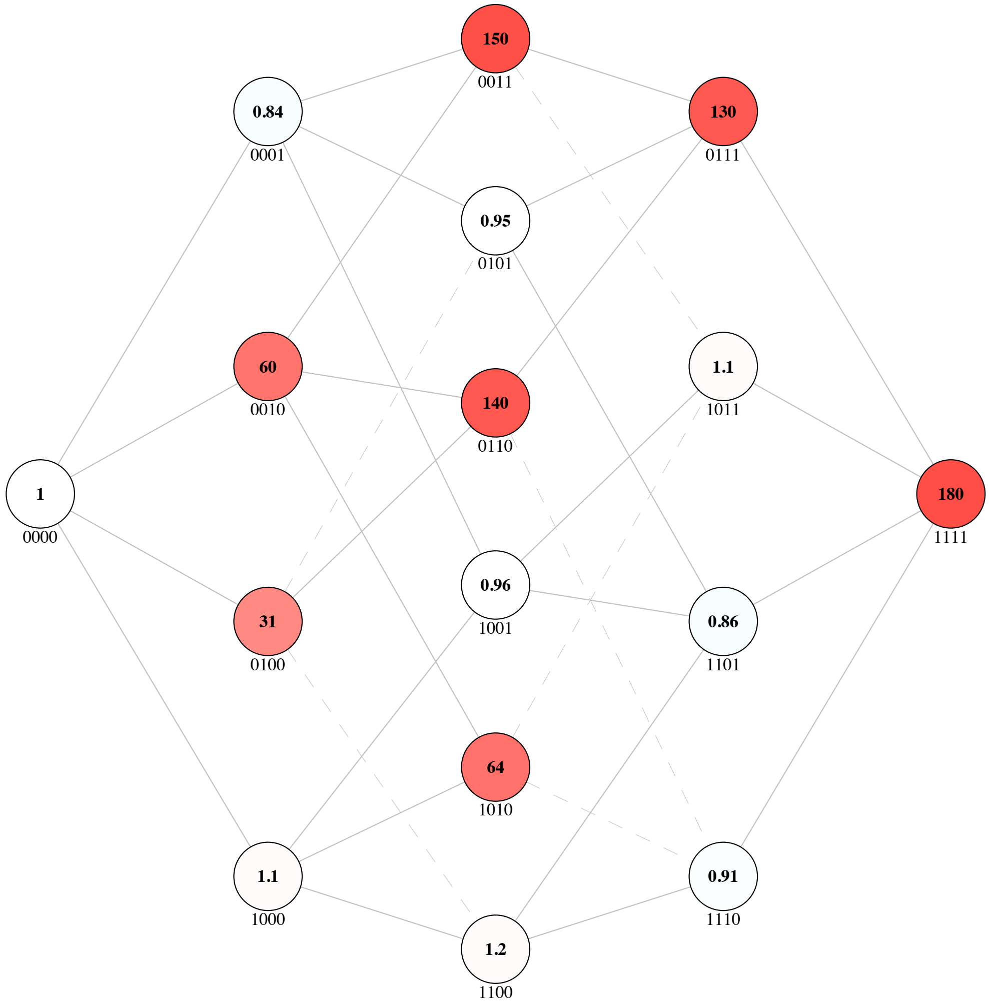


**Landscape 37 | Fitness Landscape of beta lactamase TEM in CTX antibiotic from Mira *et al.***

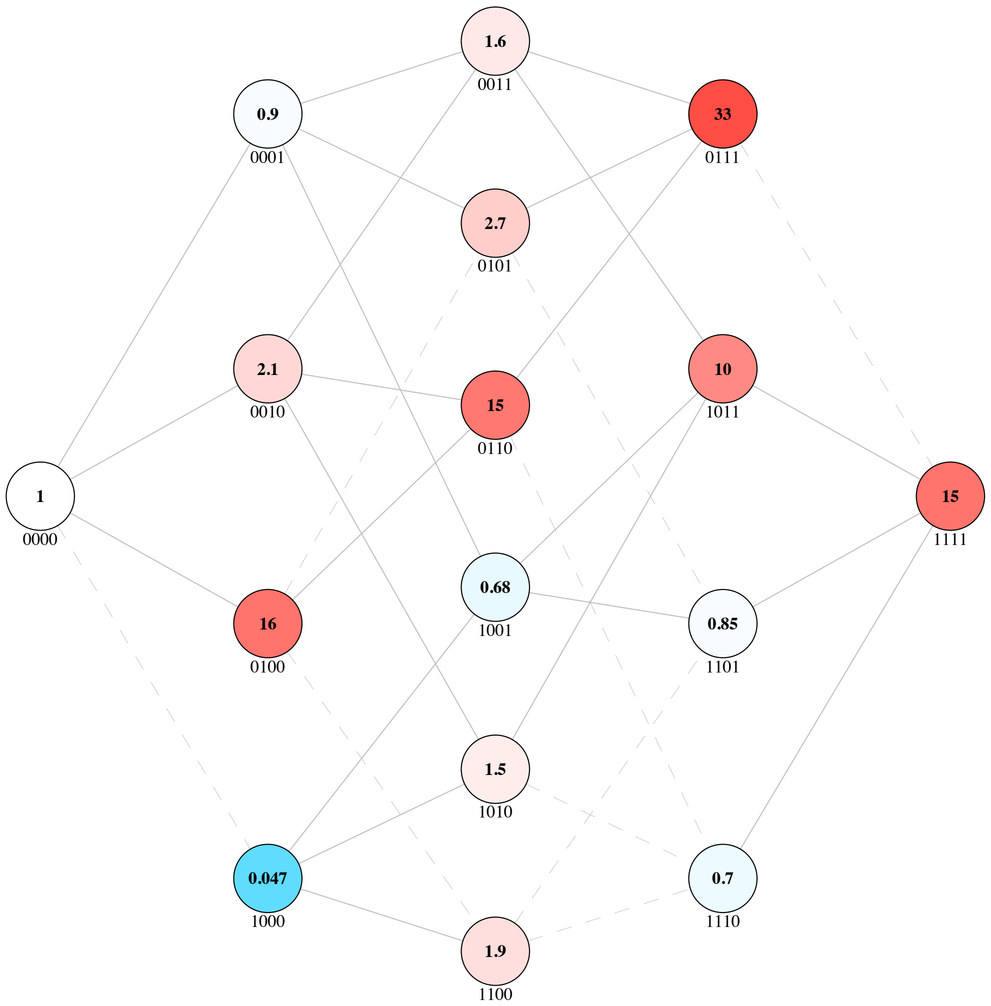


**Landscape 38 | Fitness Landscape of beta lactamase TEM in CXM antibiotic from Mira et al.**

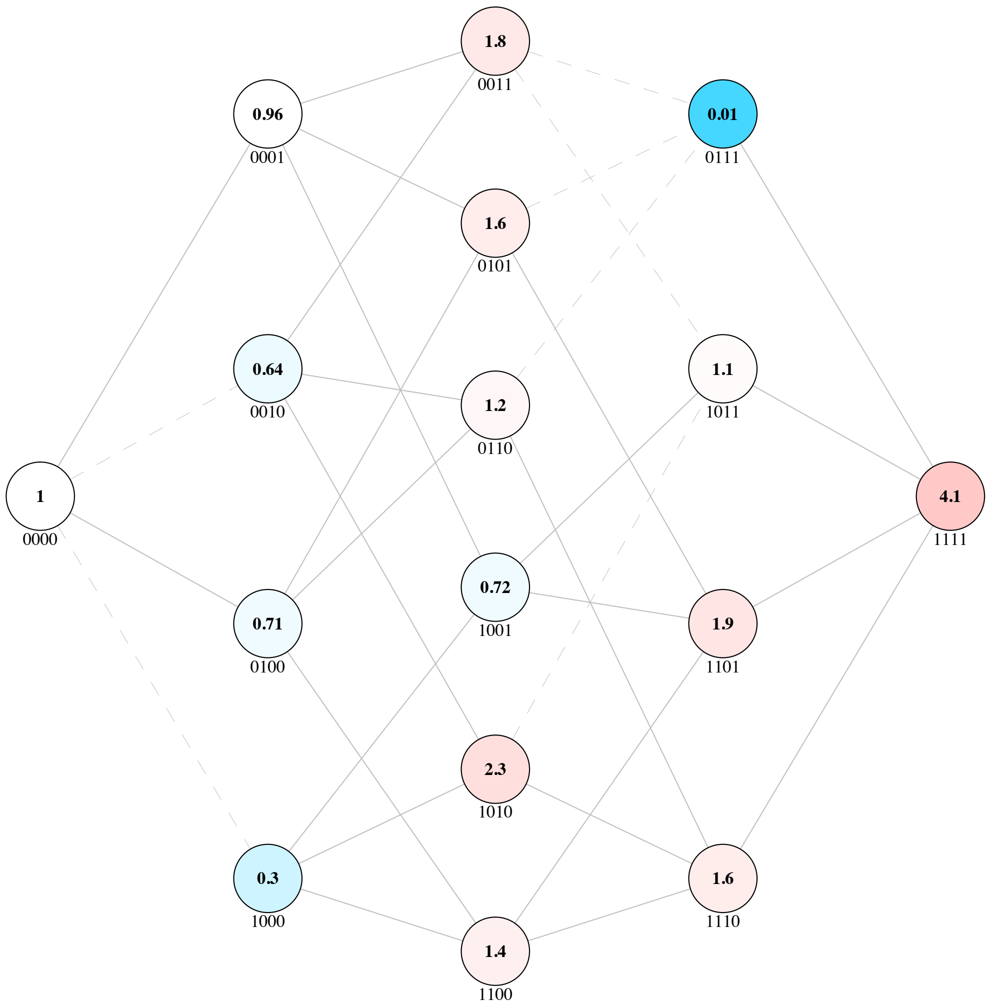


**Landscape 39 | Fitness Landscape of beta lactamase TEM in FEP antibiotic from Mira et al.**

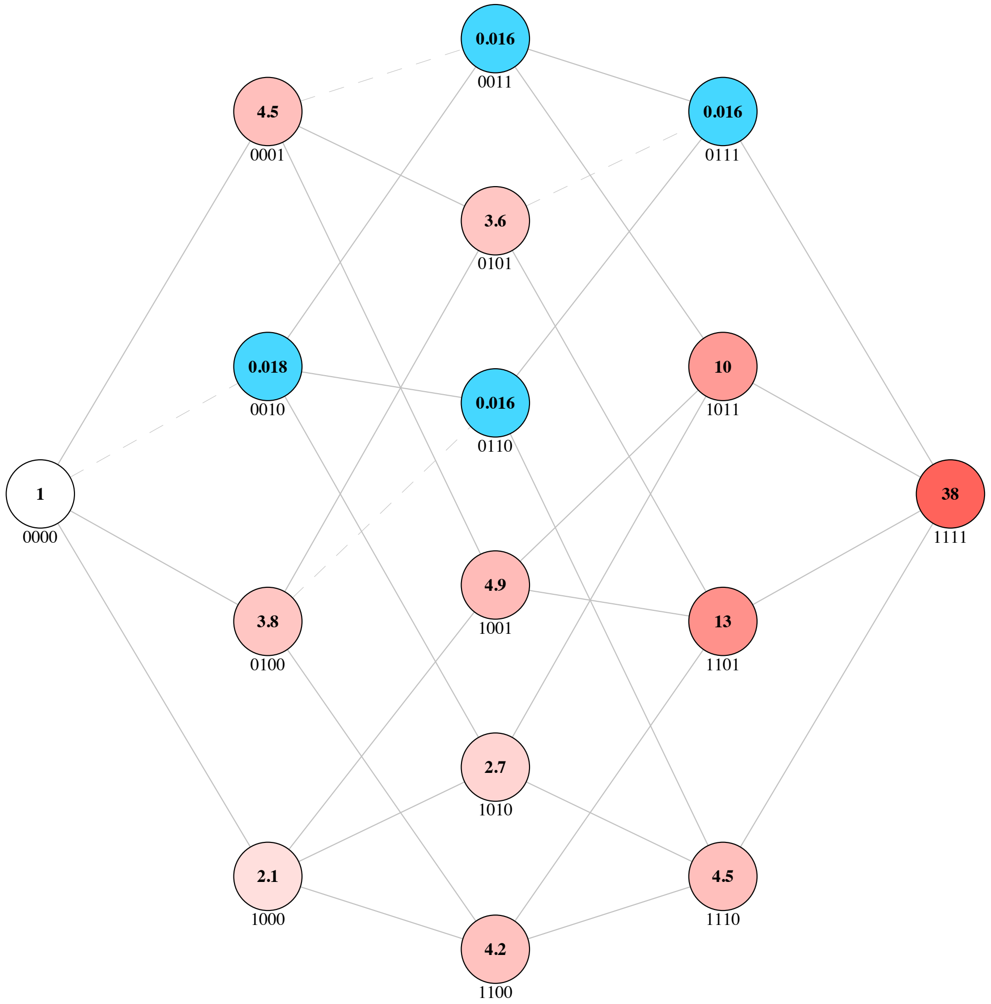


**Landscape 40 | Fitness Landscape of beta lactamase TEM in SAM antibiotic from Mira et al.**

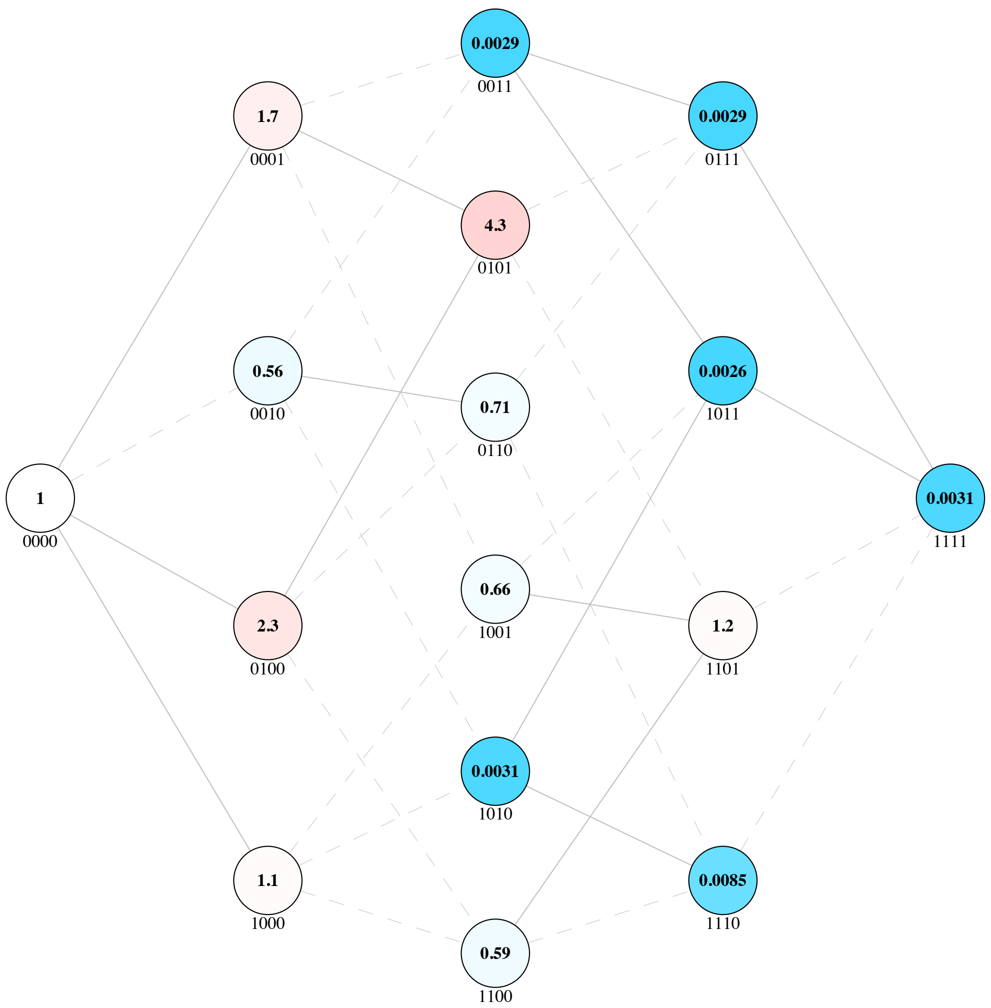


**Landscape 41 | Fitness Landscape of beta lactamase TEM in TZP antibiotic from Mira et al.**

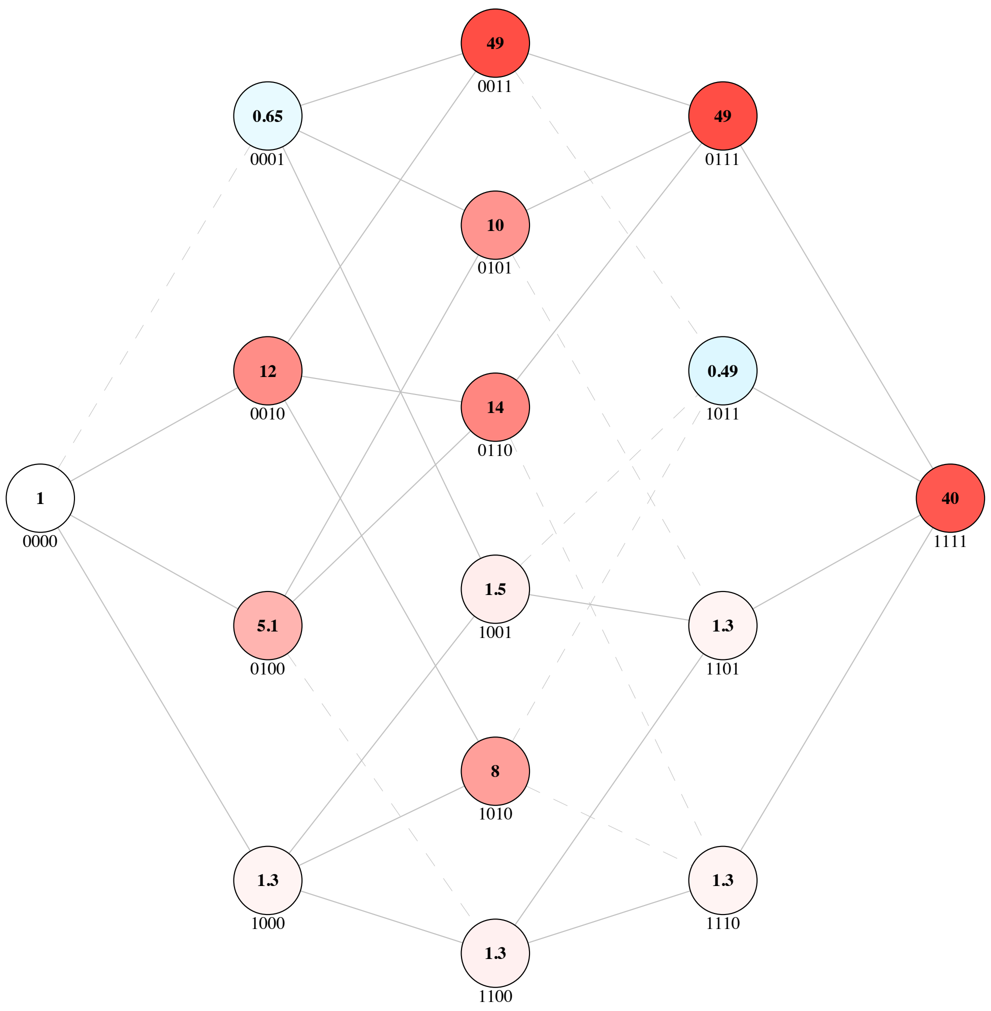


**Landscape 42 | Fitness Landscape of beta lactamase TEM in ZOX antibiotic from Mira et al.**

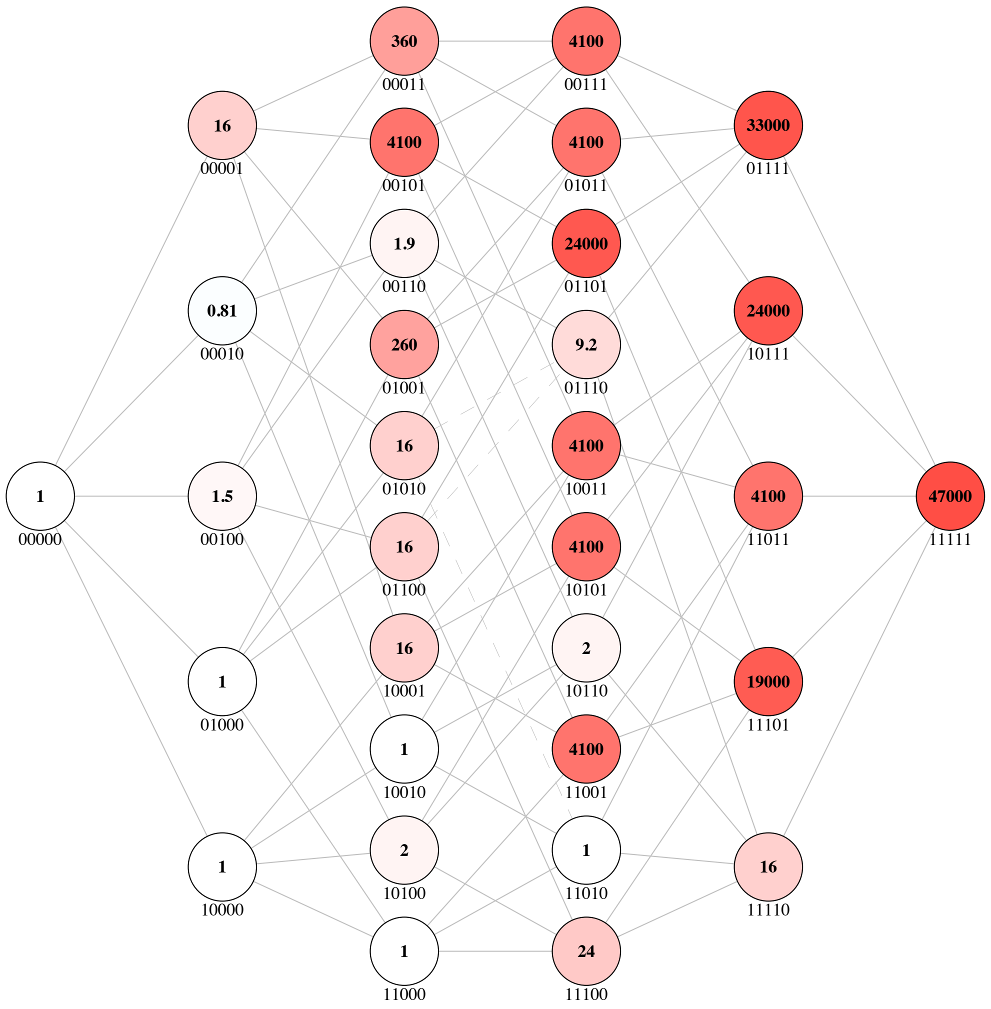


**Landscape 43 | Fitness Landscape of beta lactamase TEM from Weinreich *et al.***

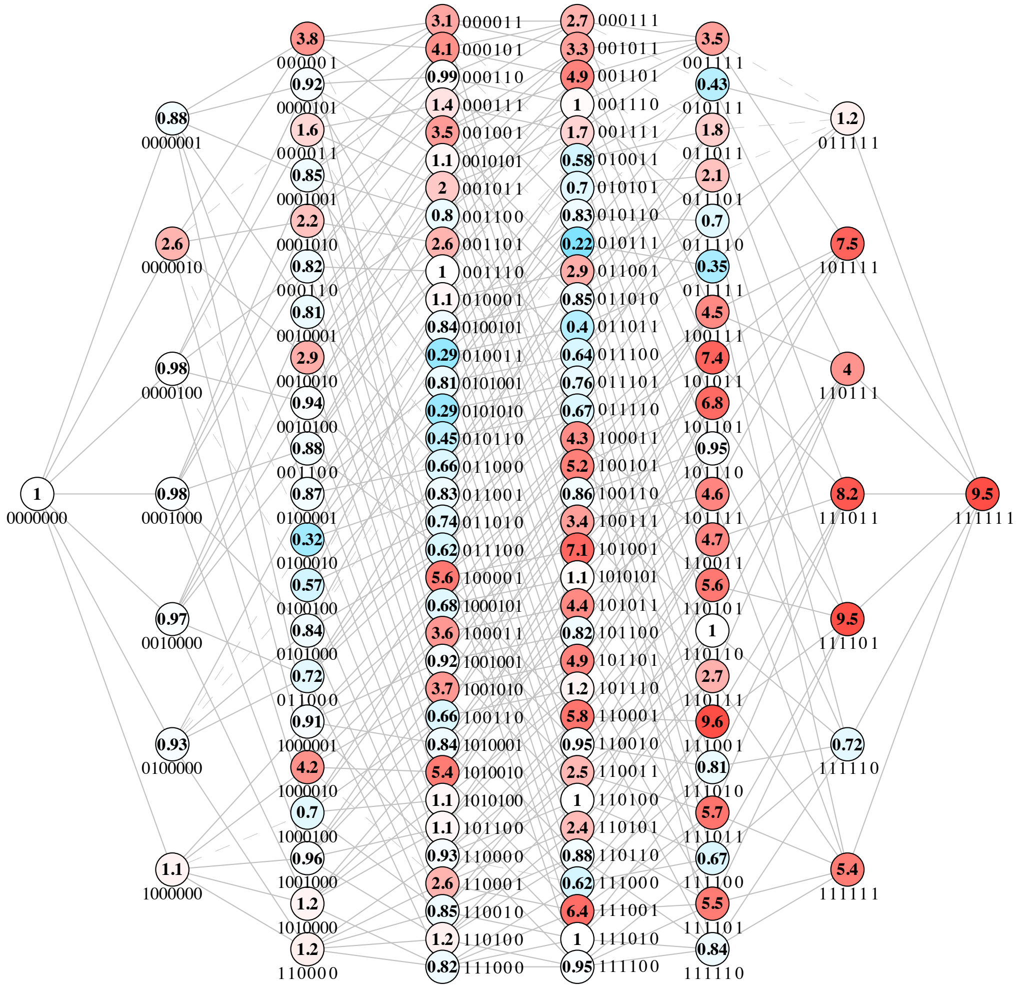

**Landscape 44 | Fitness Landscape of nitroreductase NfsA in the 20_39 trajectory from Hall *et al.***

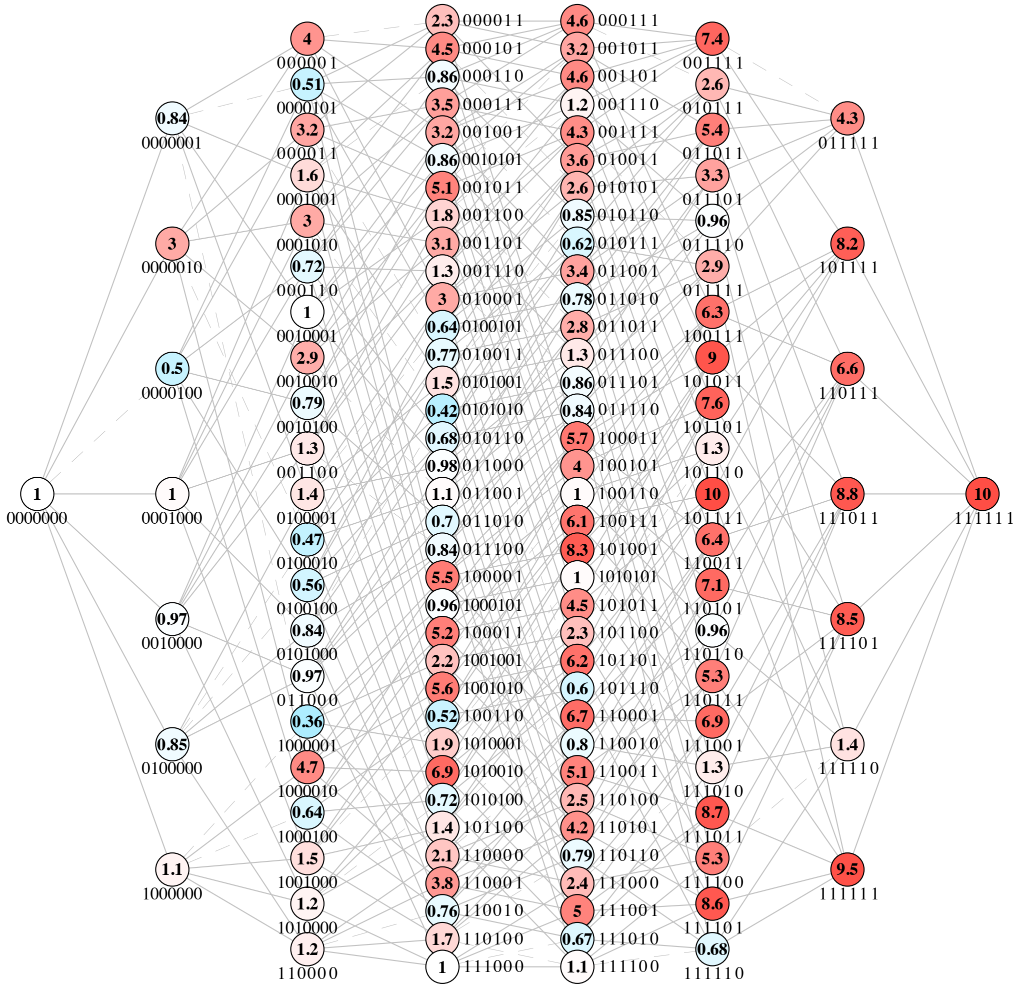


**Landscape 45 | Fitness Landscape of nitroreductase NfsA in the 36_37 trajectory from Hall *et al.***
