## Supplementary File 2 for "Higher-order epistasis creates idiosyncrasy, confounding predictions in protein evolution"

Representations of 45 functional contribution ($\Delta F$) and 45 epistatic (ε) box plots for each fitness landscape used in the study. Red bars represent mean $\Delta F$ or ε for a position or combination, respectively. Black line represents 0 and dashed lines represent positive and negative boundaries for our 1.5-fold significance threshold. $\Delta F$ and ε are both on a $\log_{10}$ scale except for all Mira *et al.* TEM boxplots due to their growth rate (additive *vs*. multiplicative) measurements.

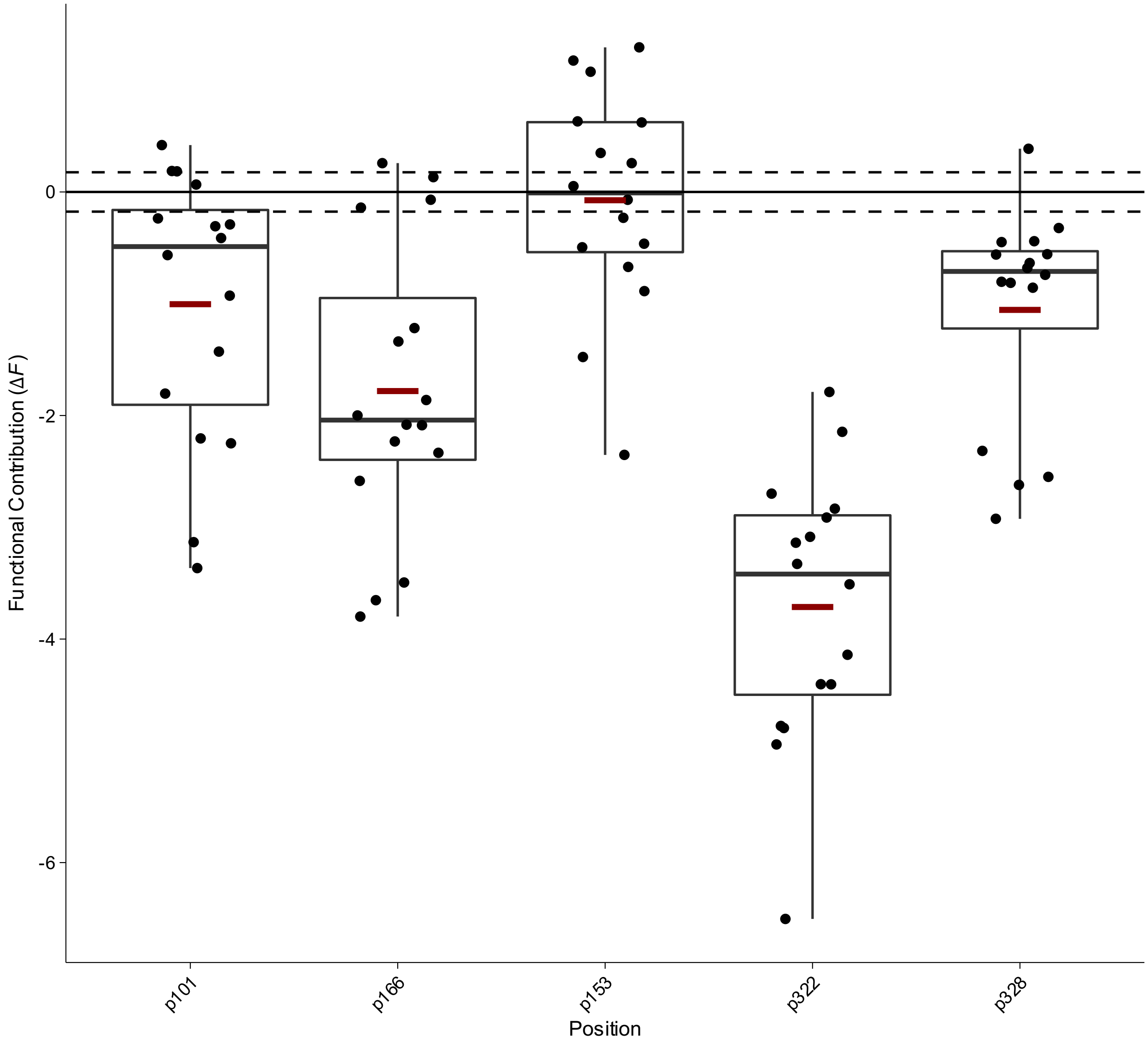


**Box Plot 1 | Functional contribution of positional mutations in Alkaline Phosphatase (AP) from Sunden *et al.***

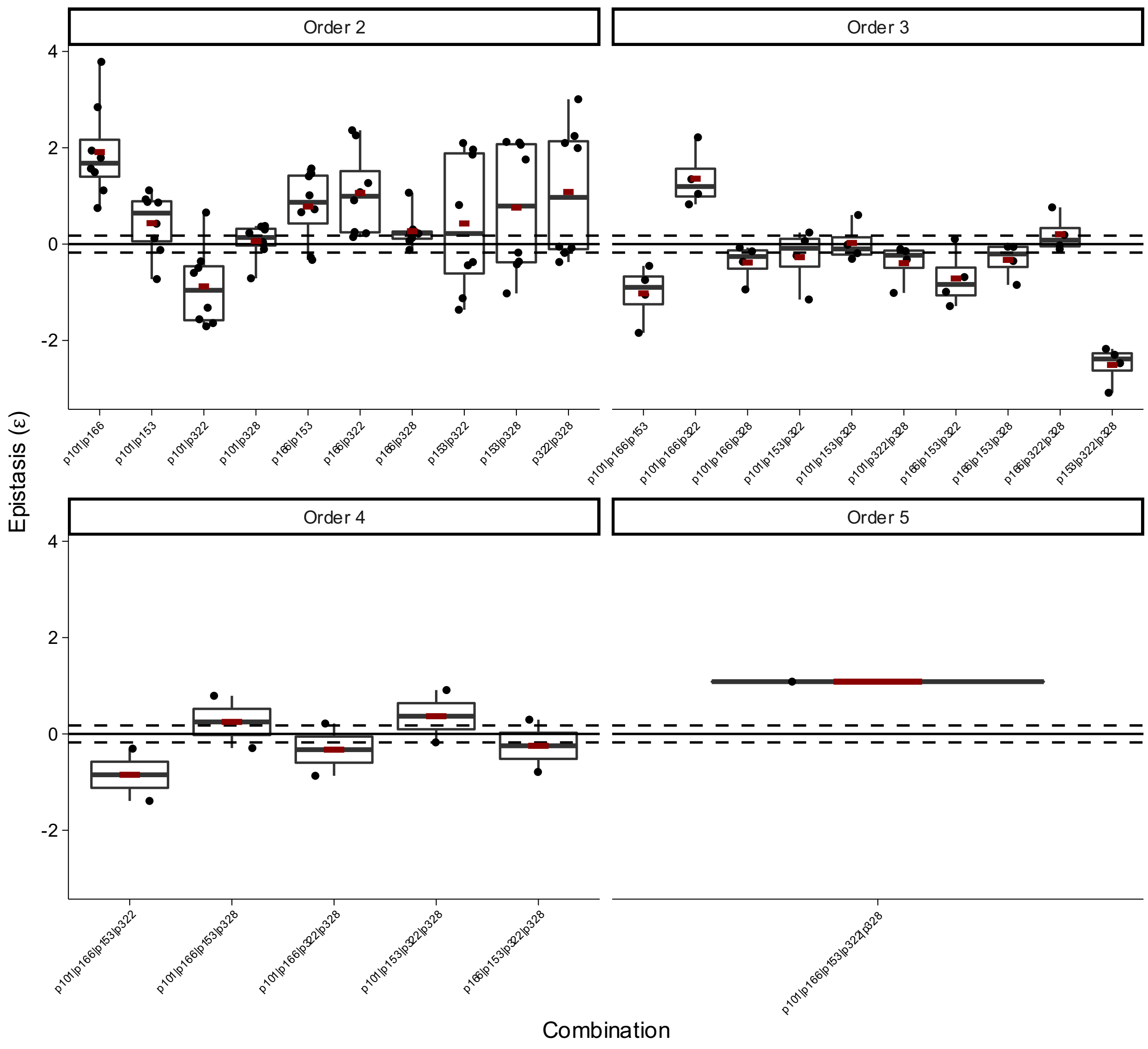


**Box Plot 2 | Epistasis of combinatorial mutations in Alkaline Phosphatase (AP) from Sunden *et al.***

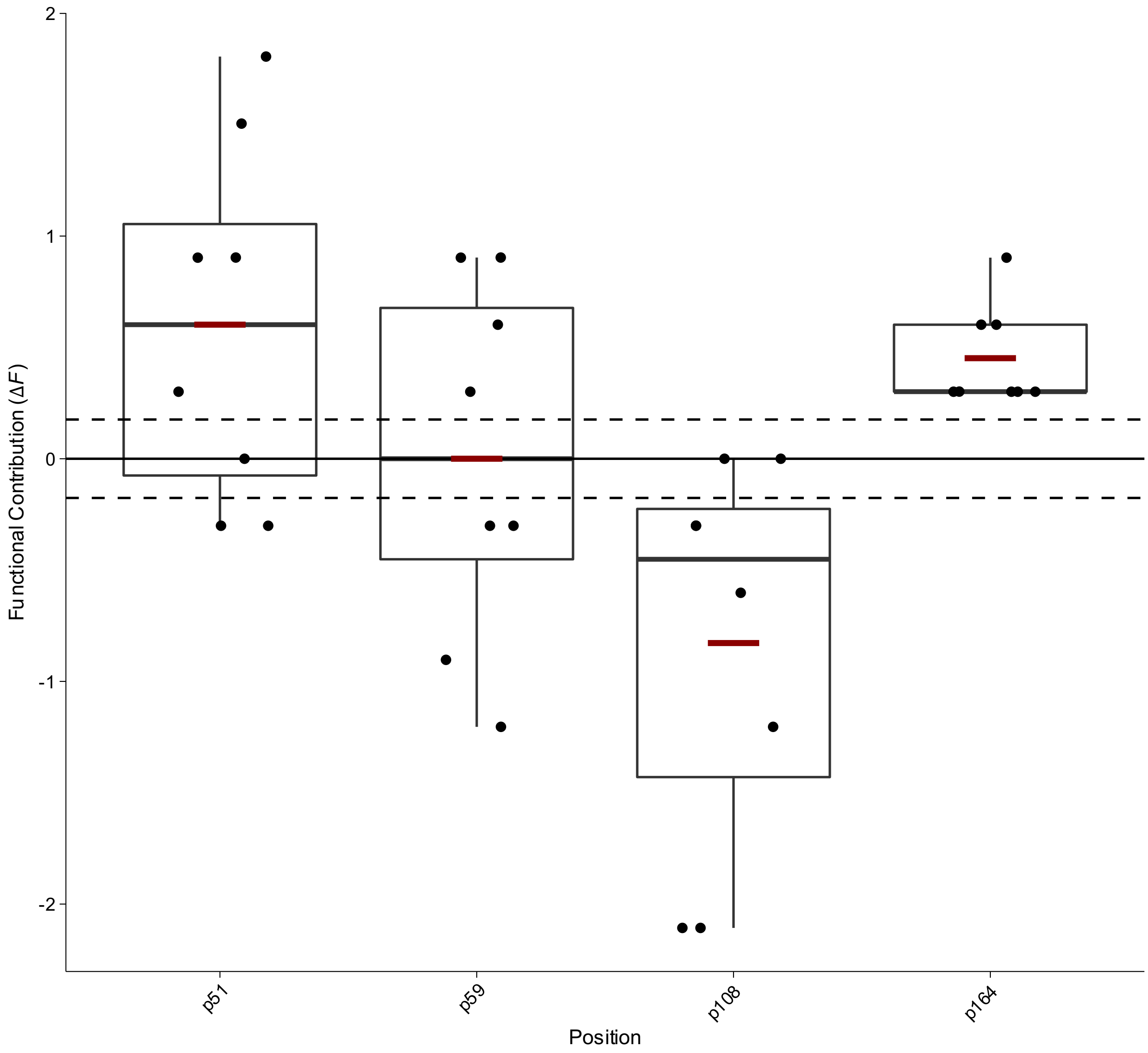


**Box Plot 3 | Functional contribution of positional mutations in Dihydrofolate Reductase (DHFR) for inhibitor c57 from Lozovsky *et al.***

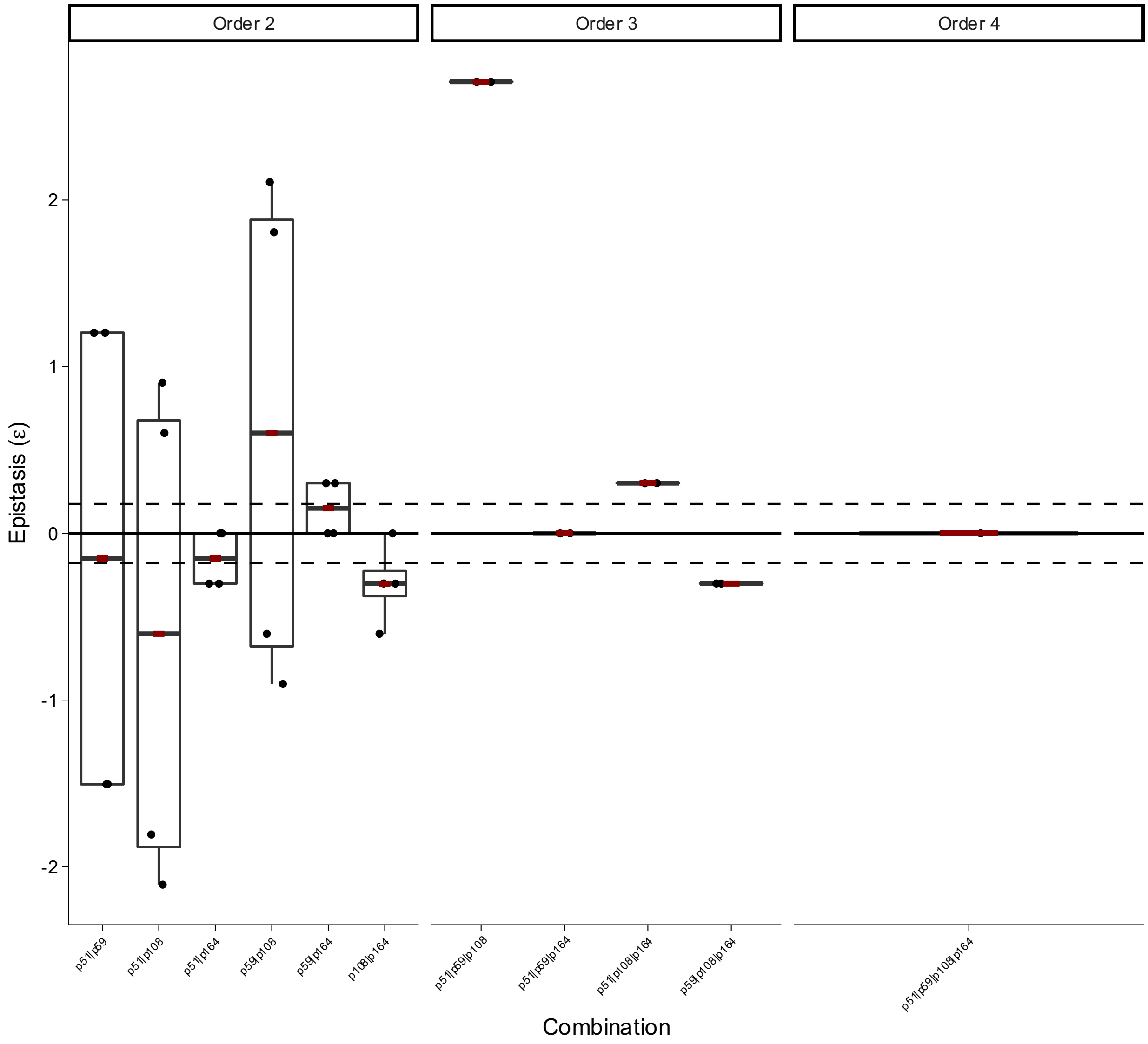


**Box Plot 4 | Epistasis of combinatorial mutations in Dihydrofolate Reductase (DHFR) for inhibitor c57 from Lozovsky *et al.***

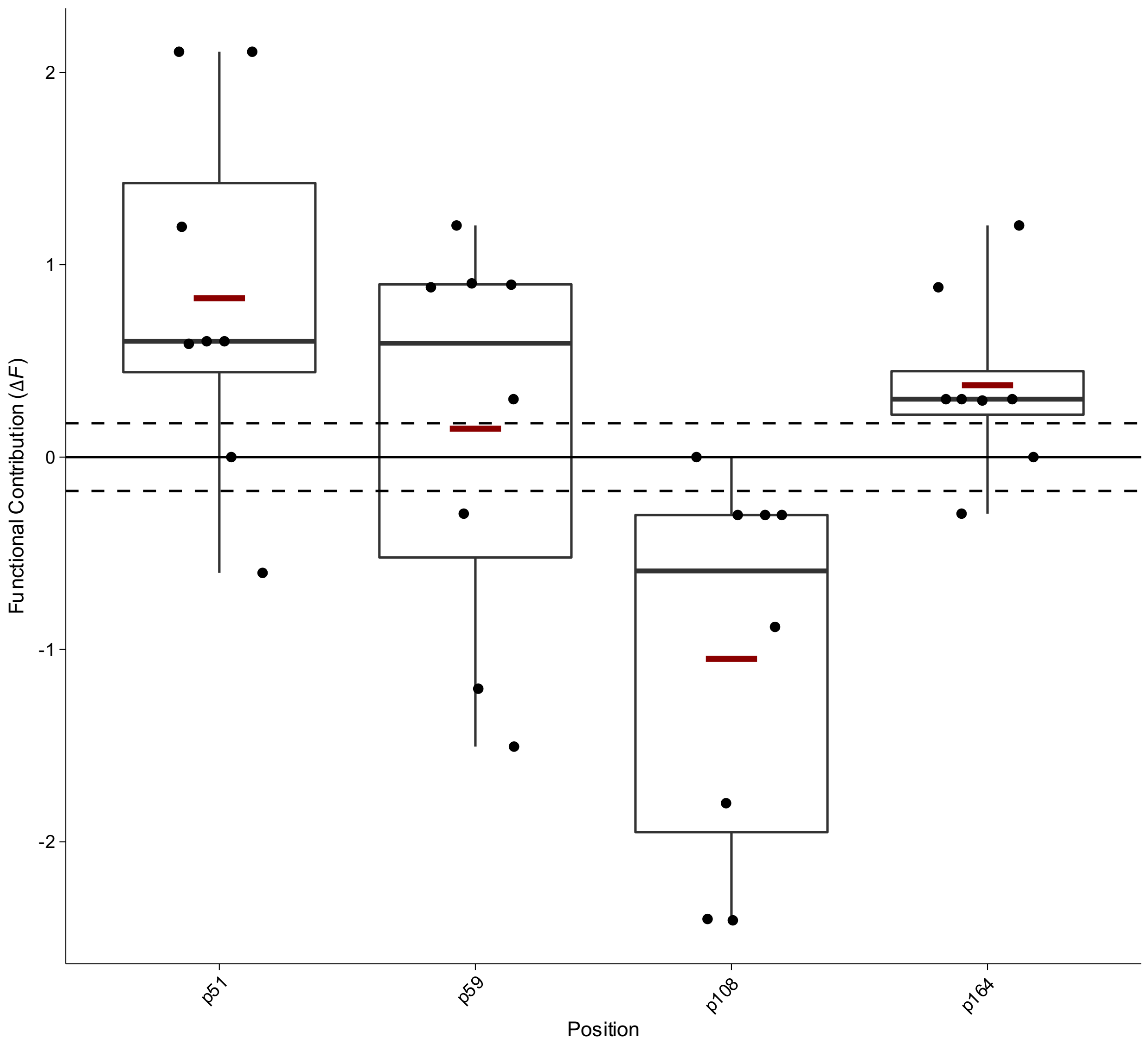


**Box Plot 5 | Functional contribution of positional mutations in Dihydrofolate Reductase (DHFR) for inhibitor c58 from Lozovsky *et al.***

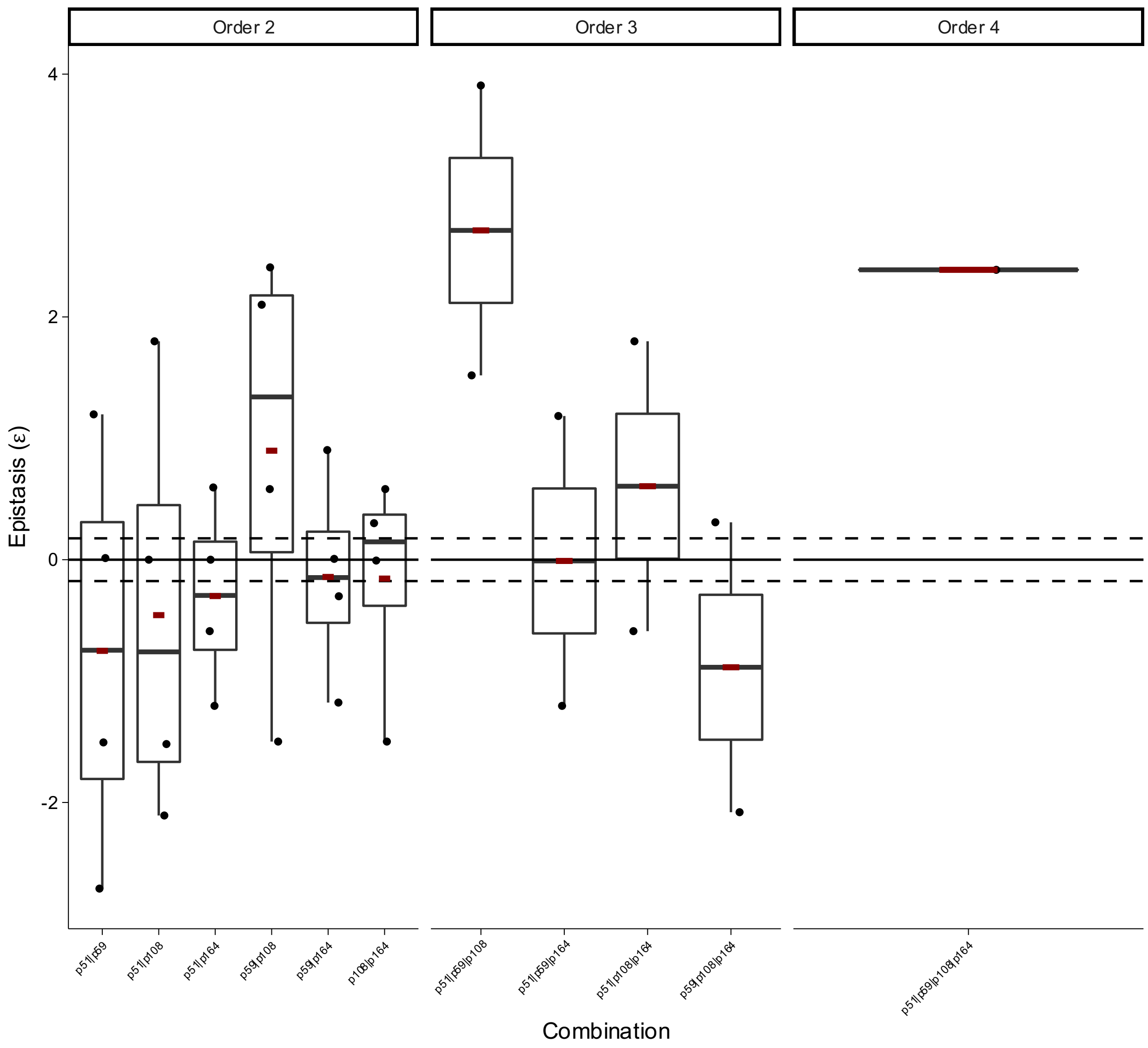


**Box Plot 6 | Epistasis of combinatorial mutations in Dihydrofolate Reductase (DHFR) for inhibitor c58 from Lozovsky *et al.***

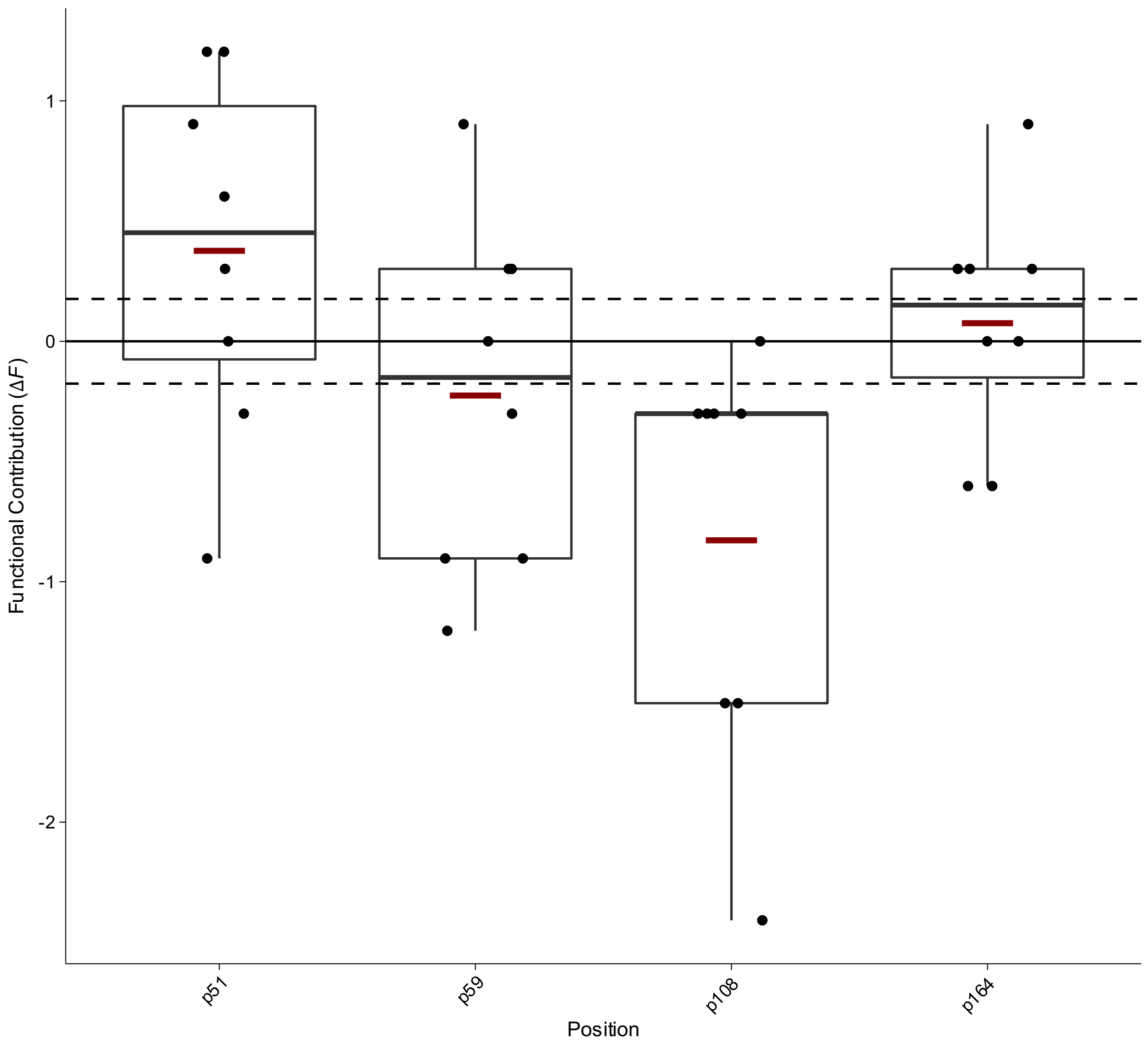


**Box Plot 7 | Functional contribution of positional mutations in Dihydrofolate Reductase (DHFR) for inhibitor c59 from Lozovsky *et al.***

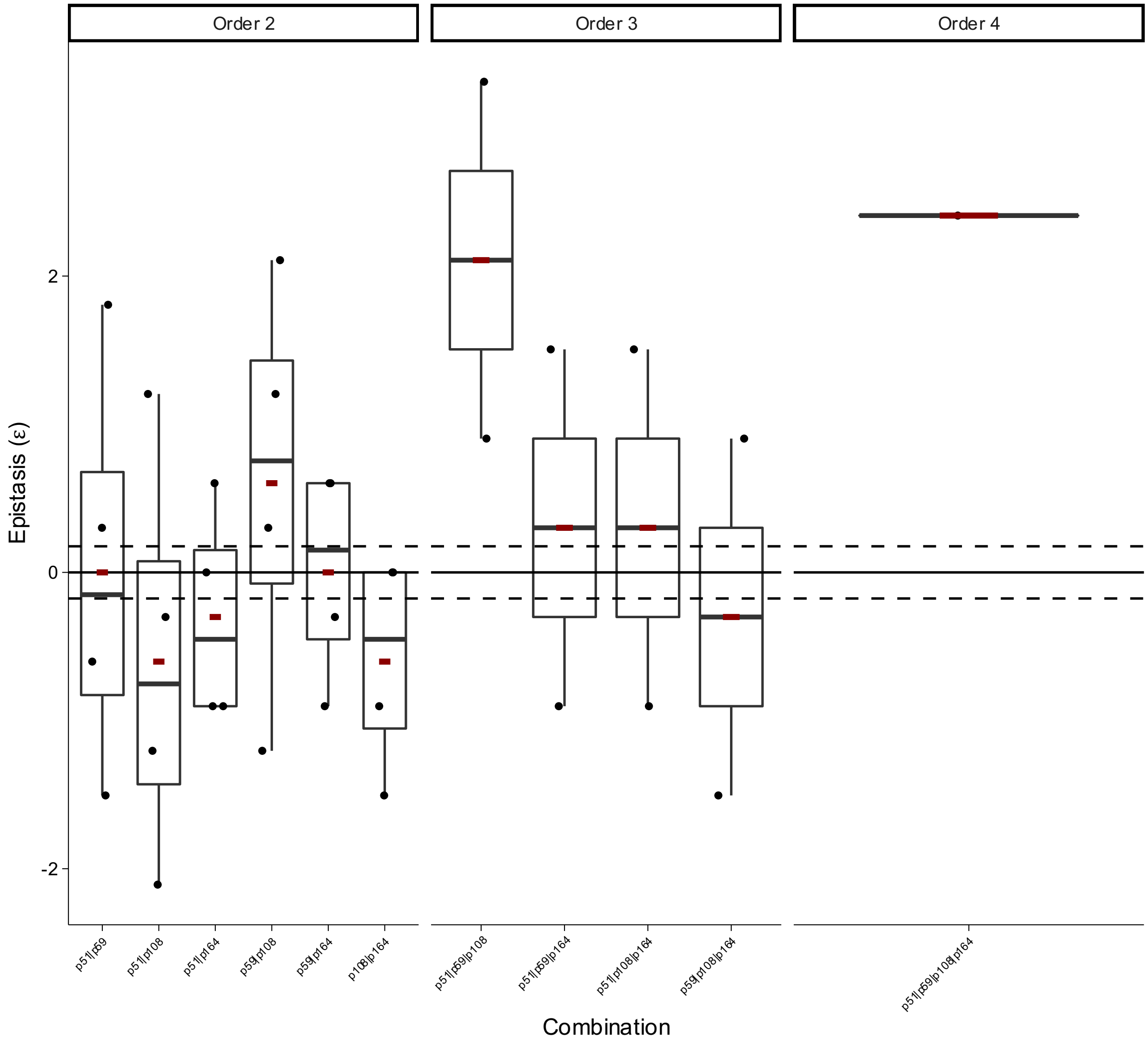


**Box Plot 8 | Epistasis of combinatorial mutations in Dihydrofolate Reductase (DHFR) for inhibitor c59 from Lozovsky *et al.***

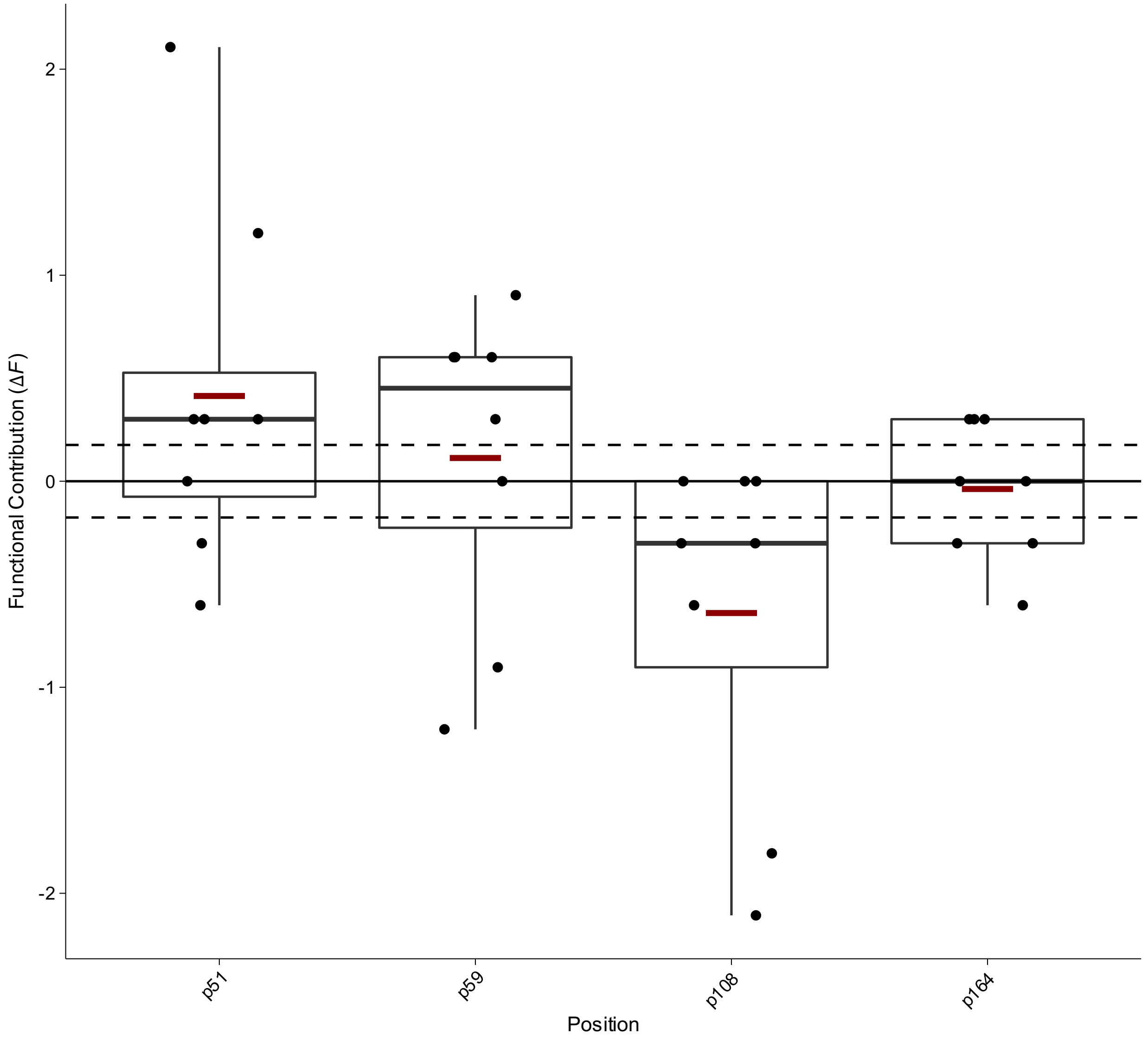


**Box Plot 9 | Functional contribution of positional mutations in Dihydrofolate Reductase (DHFR) for inhibitor c60 from Lozovsky *et al.***

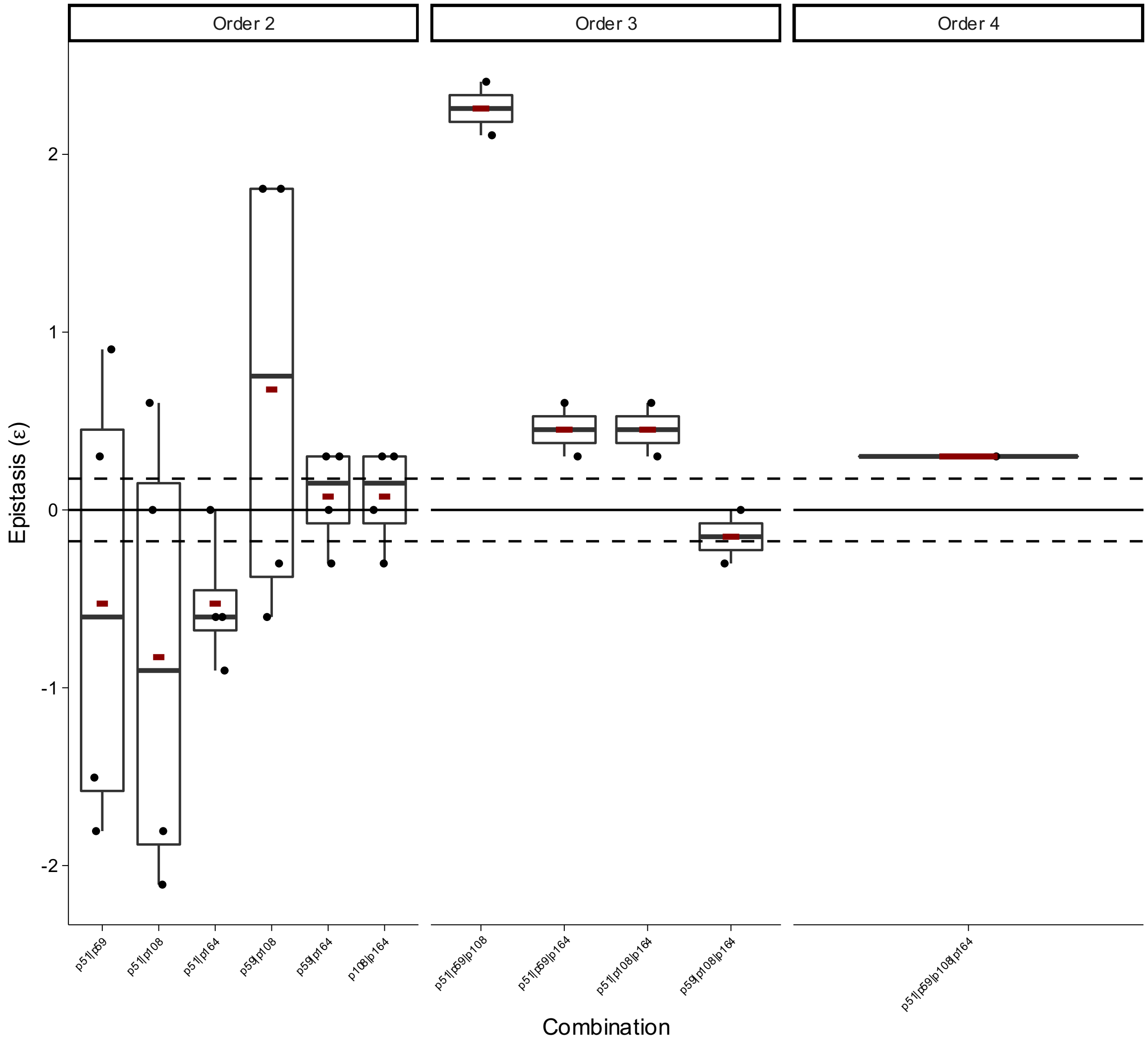


**Box Plot 10** | **Epistasis of combinatorial mutations in Dihydrofolate Reductase (DHFR) for inhibitor c60 from Lozovsky *et al.***

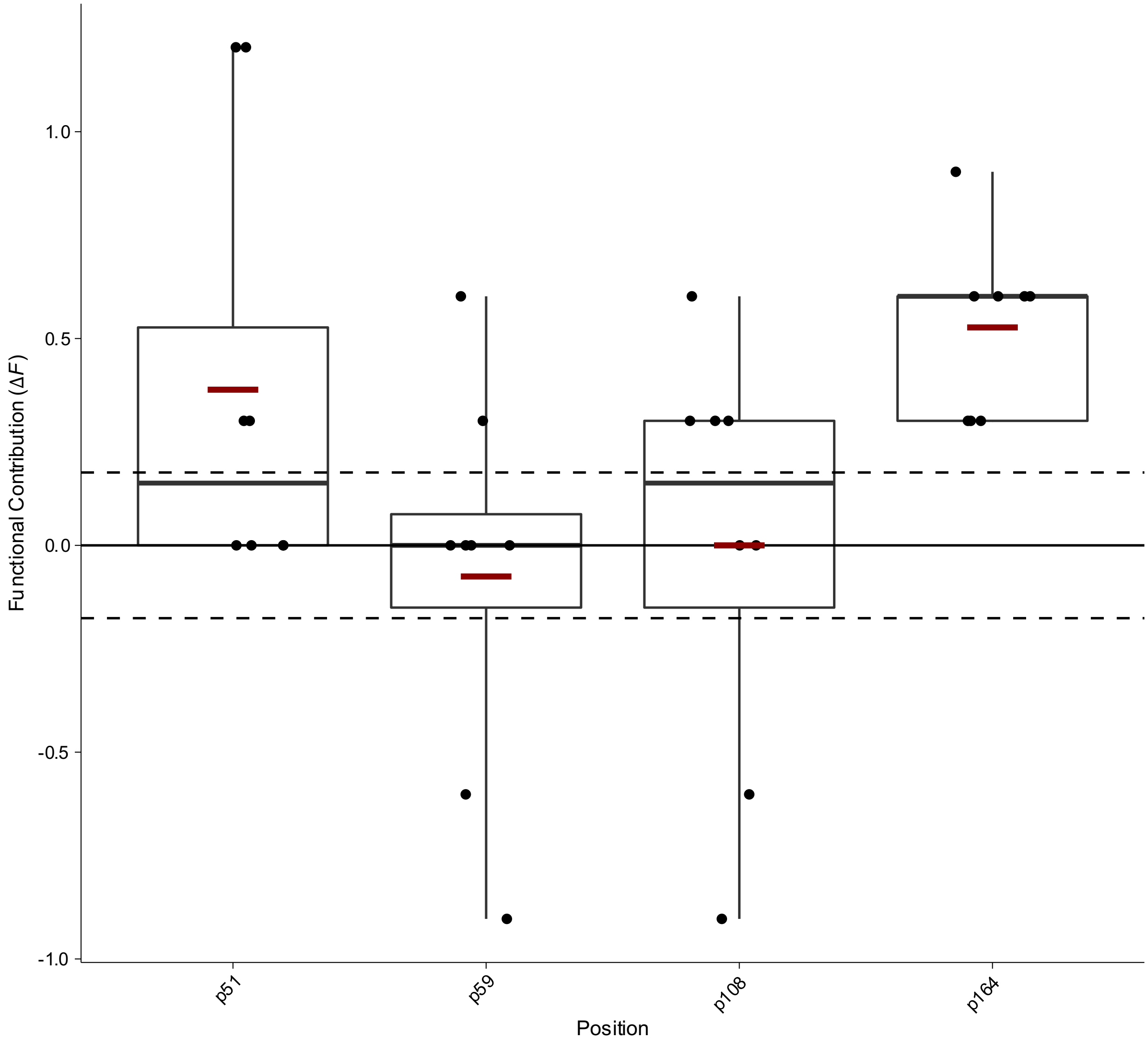


**Box Plot 11 | Functional contribution of positional mutations in Dihydrofolate Reductase (DHFR) for inhibitor c61 from Lozovsky *et al.***

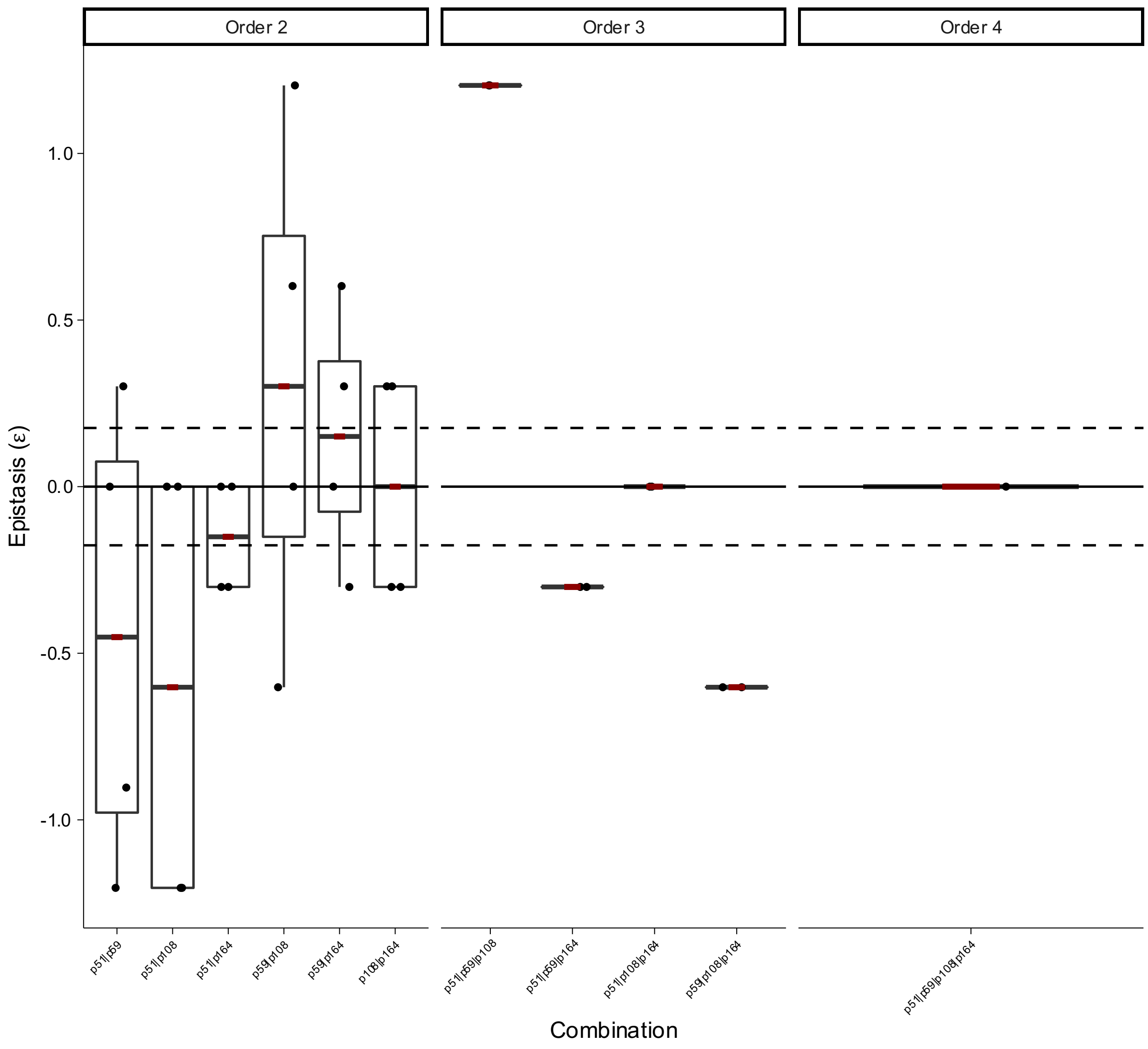


**Box Plot 12 | Epistasis of combinatorial mutations in Dihydrofolate Reductase (DHFR) for inhibitor c61 from Lozovsky *et al.***

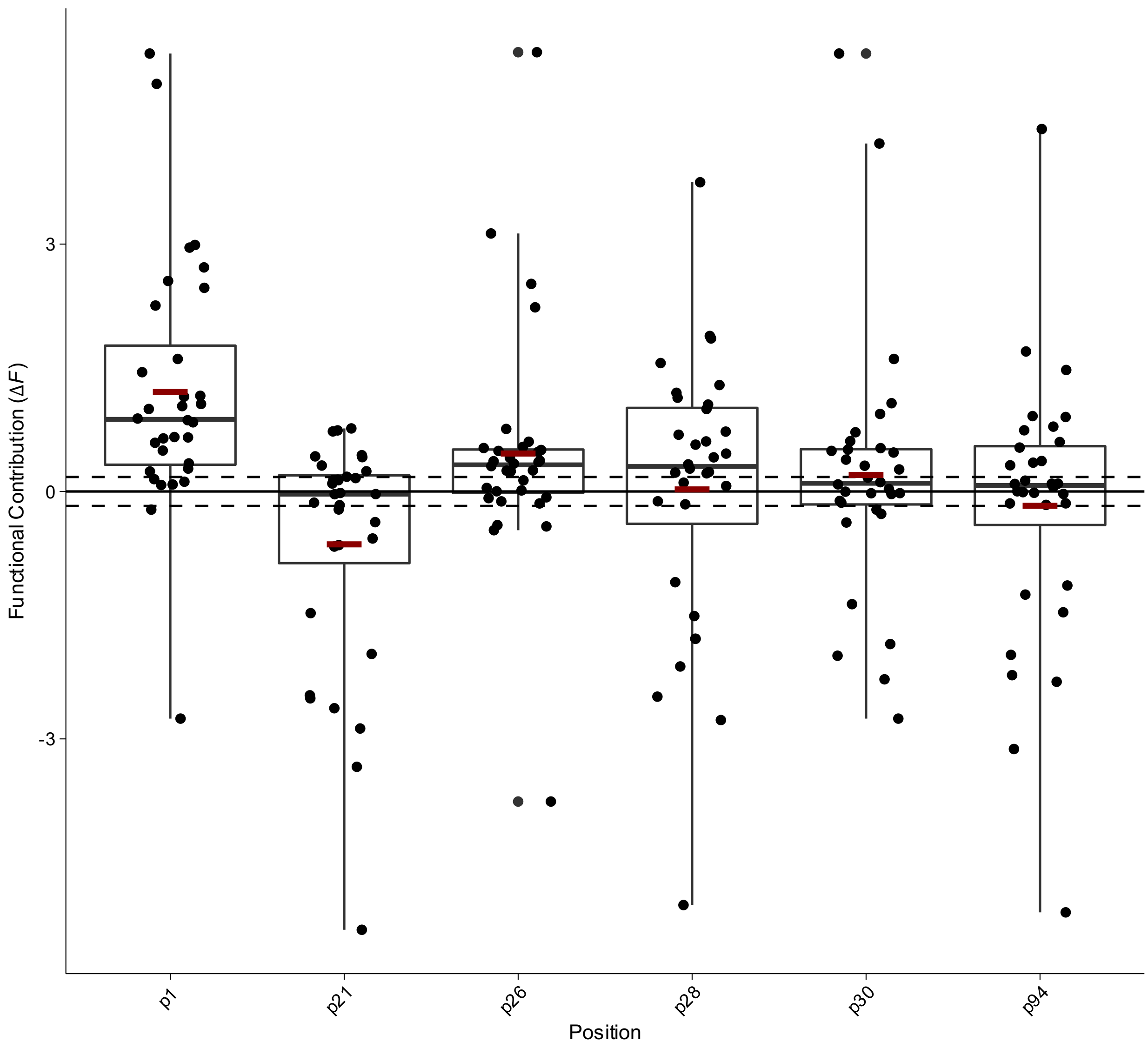


**Box Plot 13** | **Functional contribution of positional mutations in Dihydrofolate Reductase (DHFR) from Palmer *et al.***

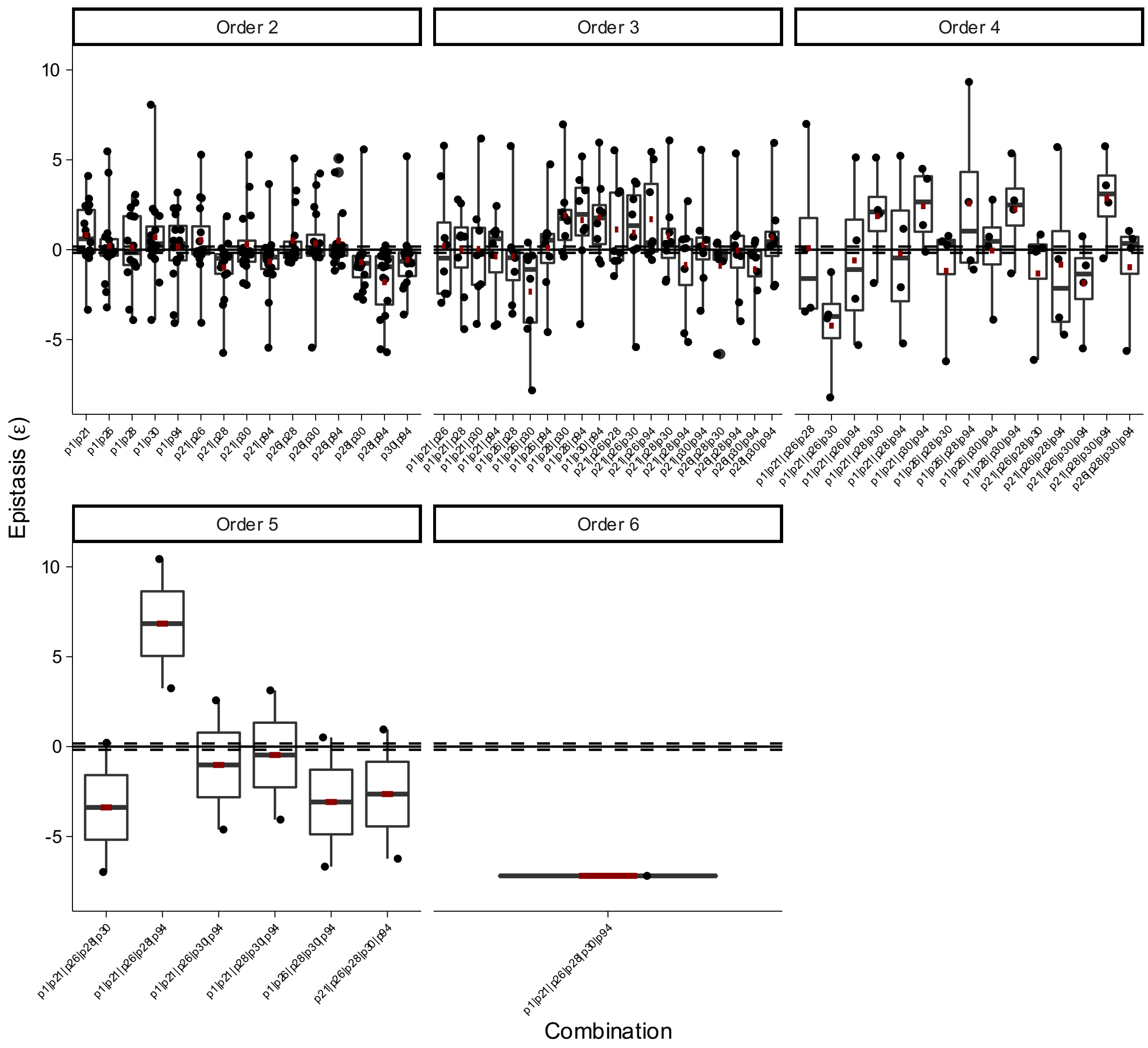


**Box Plot 14 | Epistasis of combinatorial mutations in Dihydrofolate Reductase (DHFR) from Palmer *et al.***

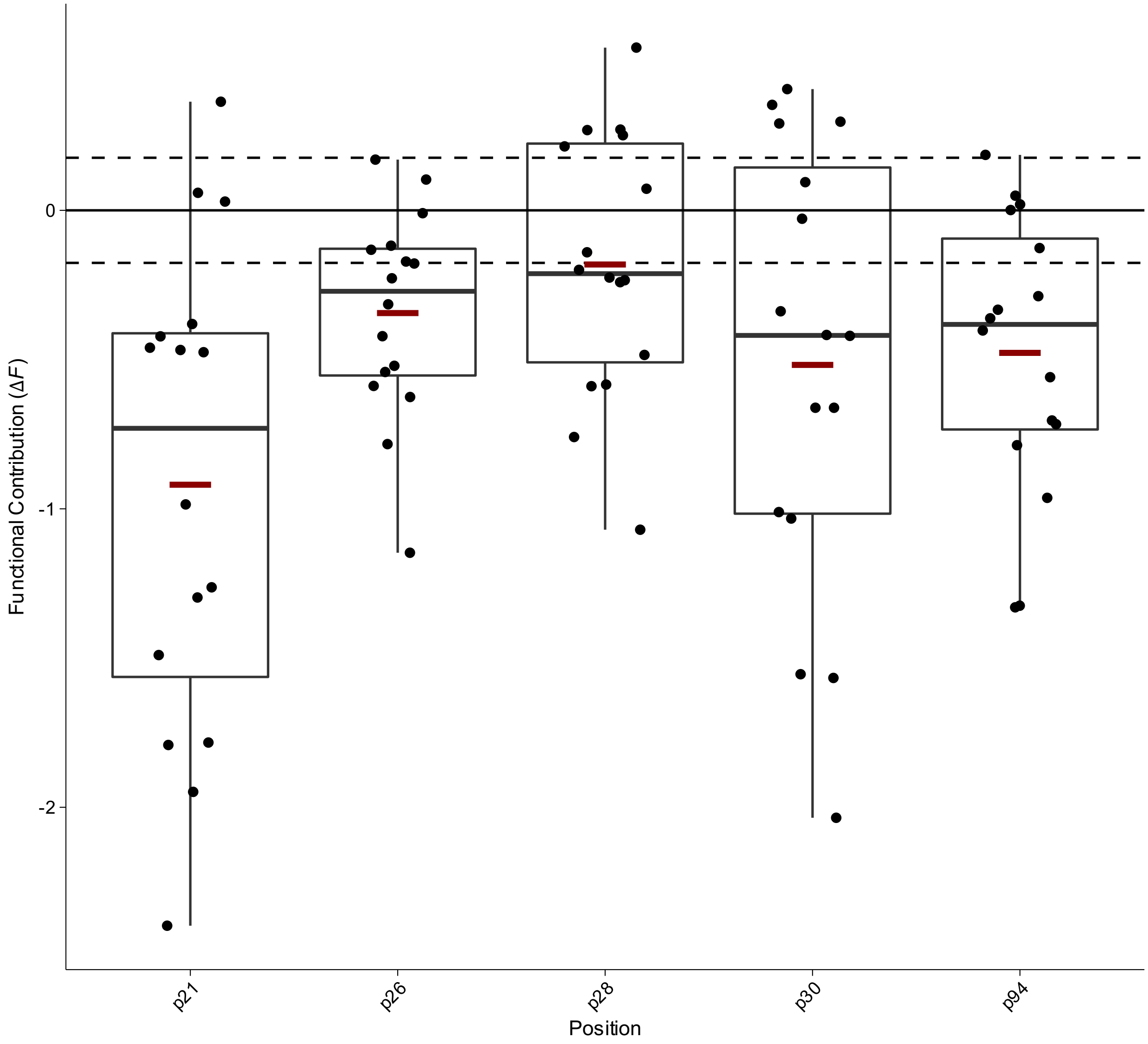


**Box Plot 15 | Functional contribution of positional mutations of Dihydrofolate Reductase (DHFR) of $k_{cat}$ in the glycine trajectory from Tamer *et al.***

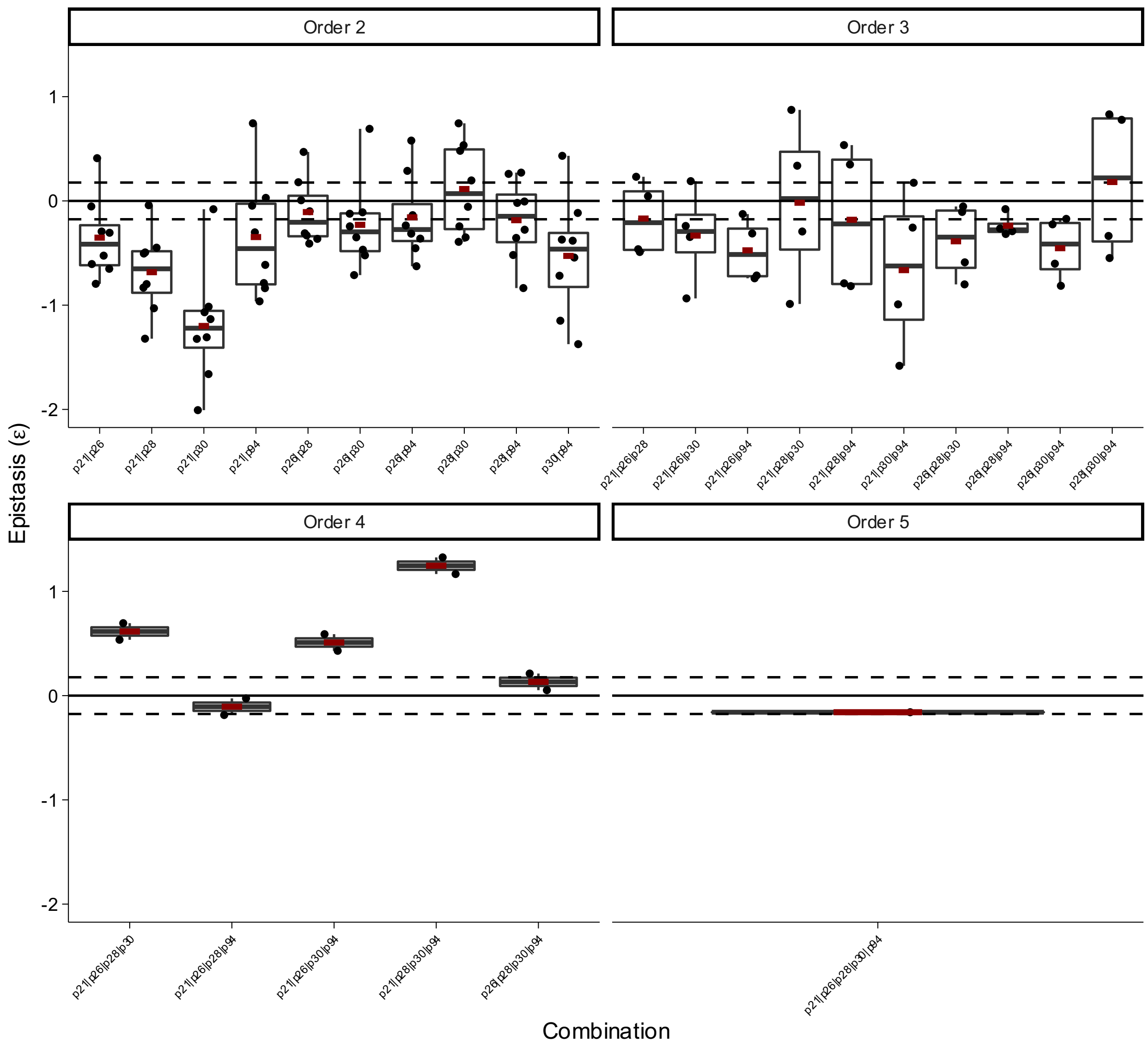


**Box Plot 16 | Epistasis of combinatorial mutations of Dihydrofolate Reductase (DHFR) of $k_{cat}$ in the glycine trajectory from Tamer *et al.***

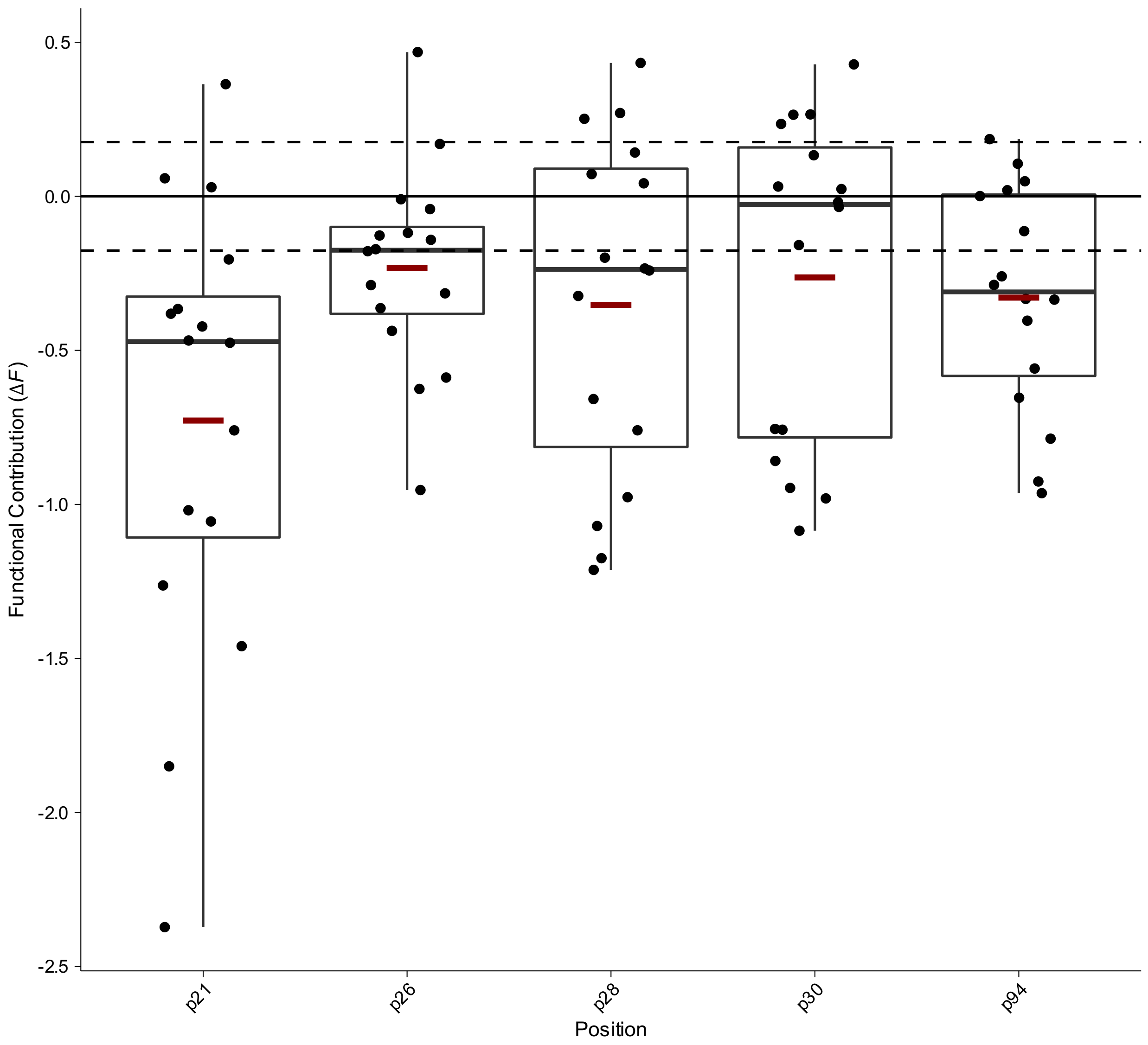


**Box Plot 17 | Functional contribution of positional mutations of Dihydrofolate Reductase (DHFR) of $k_{cat}$ in the arginine trajectory from Tamer *et al.***

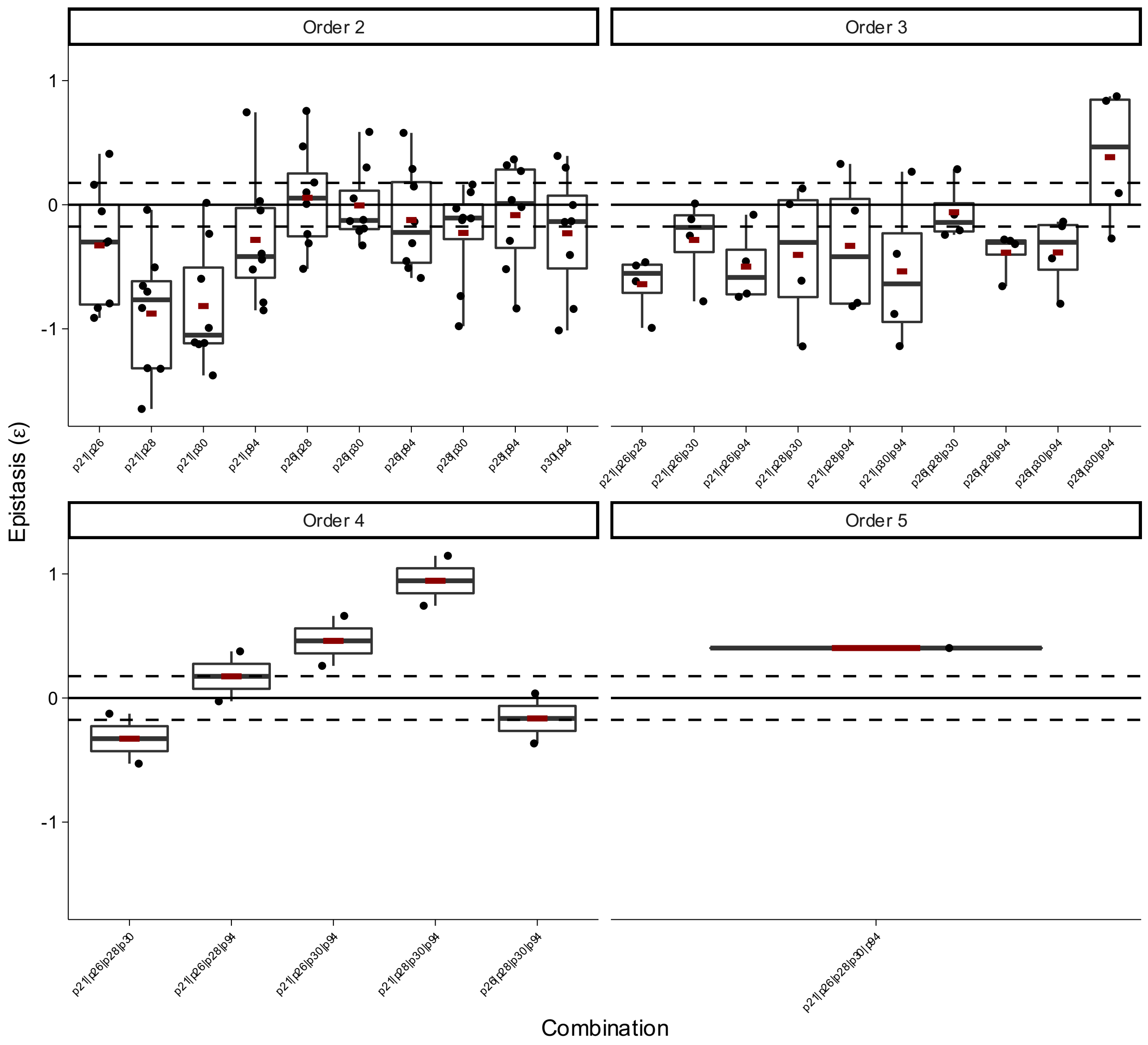


**Box Plot 18 | Epistasis of combinatorial mutations of Dihydrofolate Reductase (DHFR) of $k_{cat}$ in the arginine trajectory from Tamer *et al.***

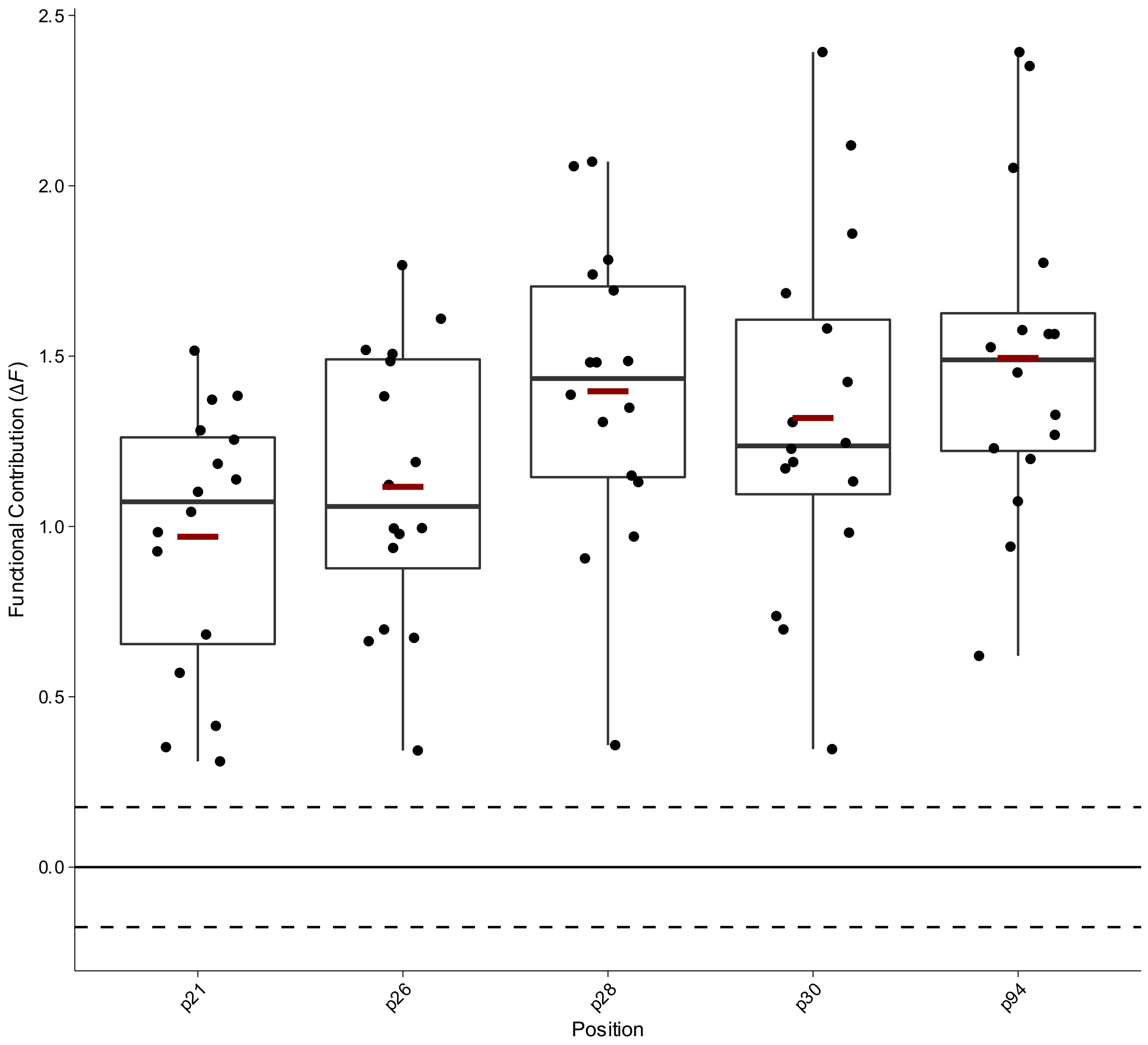


**Box Plot 19 | Functional contribution of positional mutations of Dihydrofolate Reductase (DHFR) of $K_I$ in the glycine trajectory from Tamer *et al.***

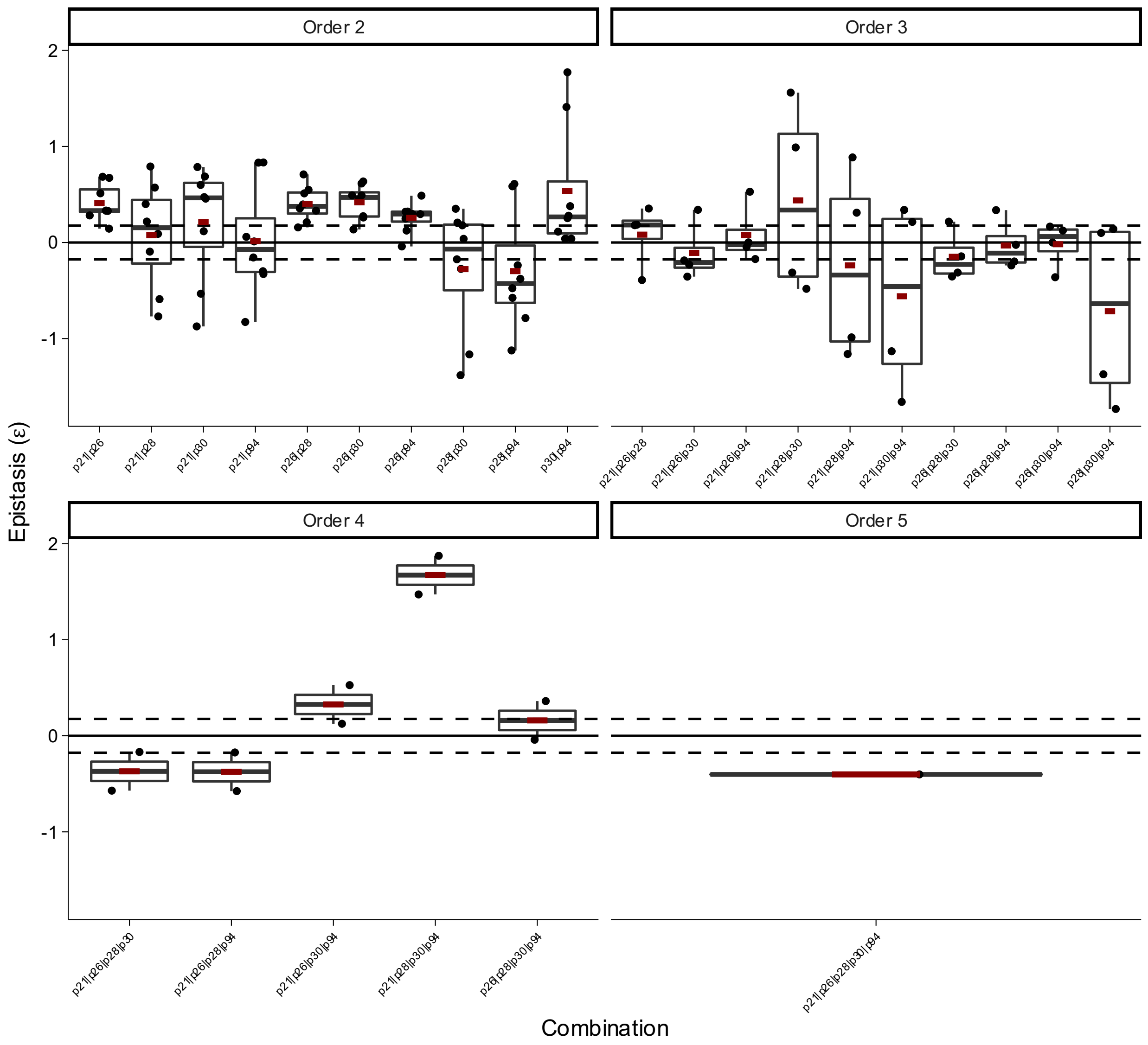


**Box Plot 20 | Epistasis of combinatorial mutations of Dihydrofolate Reductase (DHFR) of $K_I$ in the glycine trajectory from Tamer *et al.***

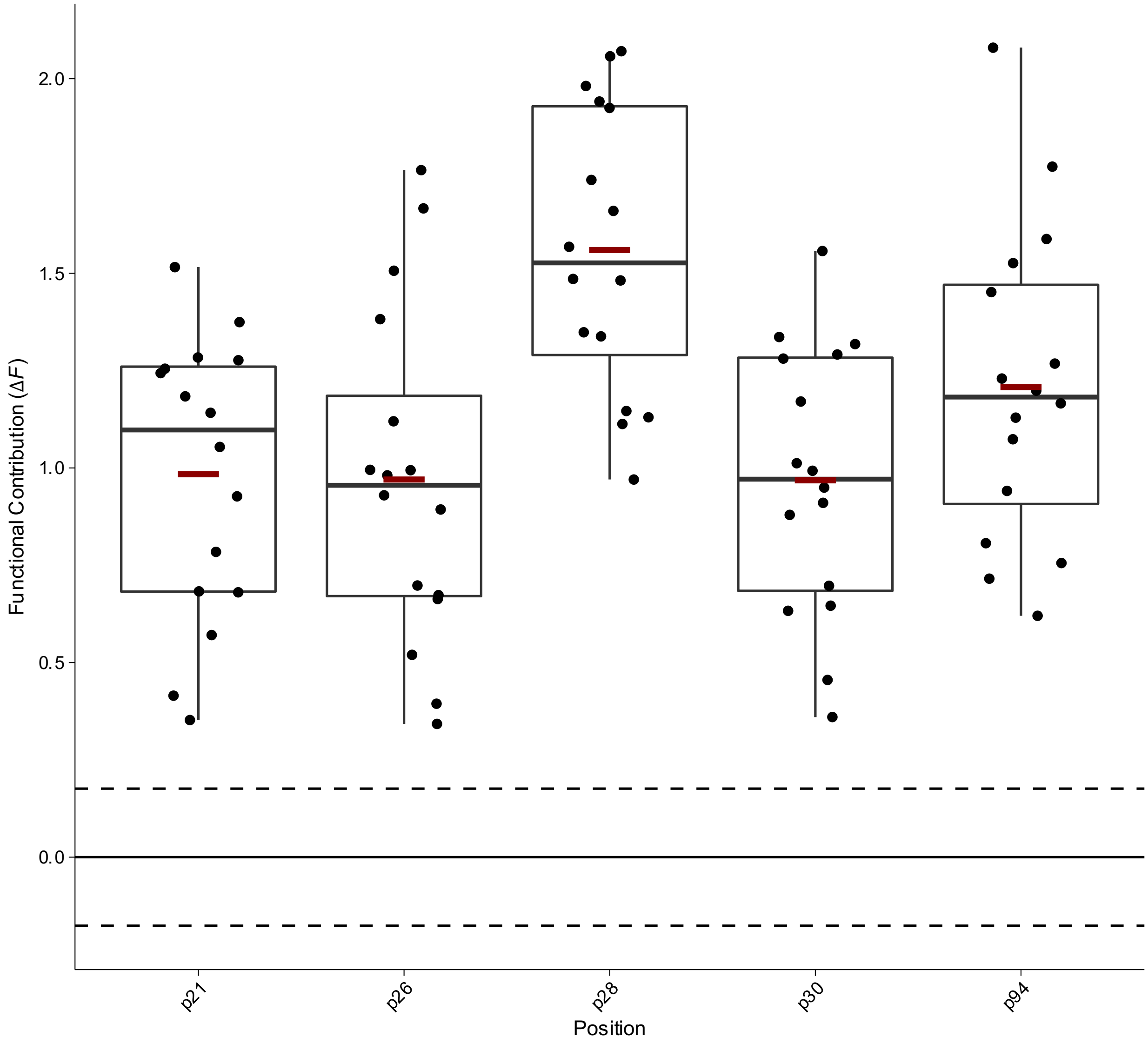


**Box Plot 21 | Functional contribution of positional mutations of Dihydrofolate Reductase (DHFR) of $K_I$ in the arginine trajectory from Tamer *et al.***

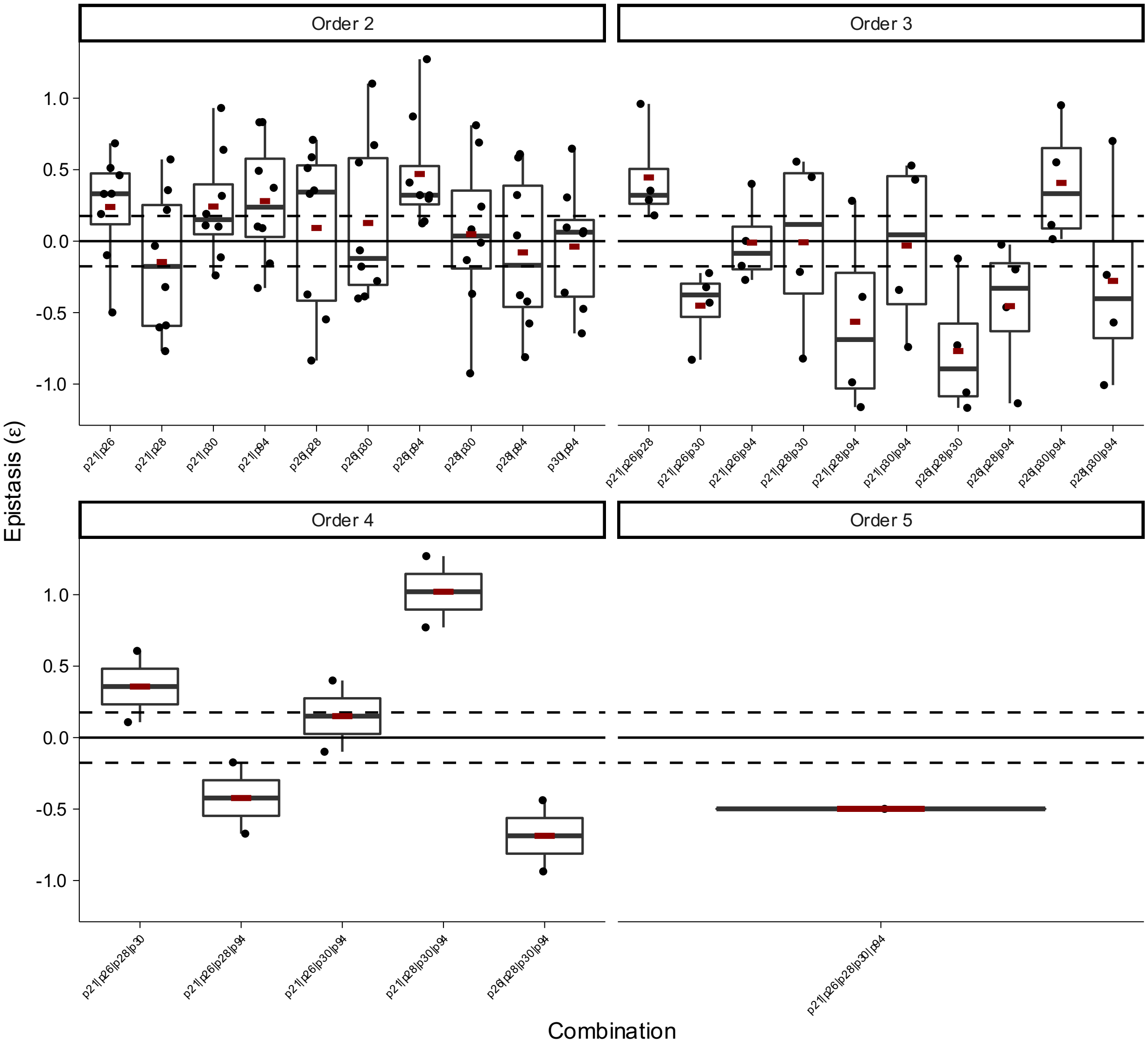


**Box Plot 22 | Epistasis of combinatorial mutations of Dihydrofolate Reductase (DHFR) of $K_I$ in the arginine trajectory from Tamer *et al.***

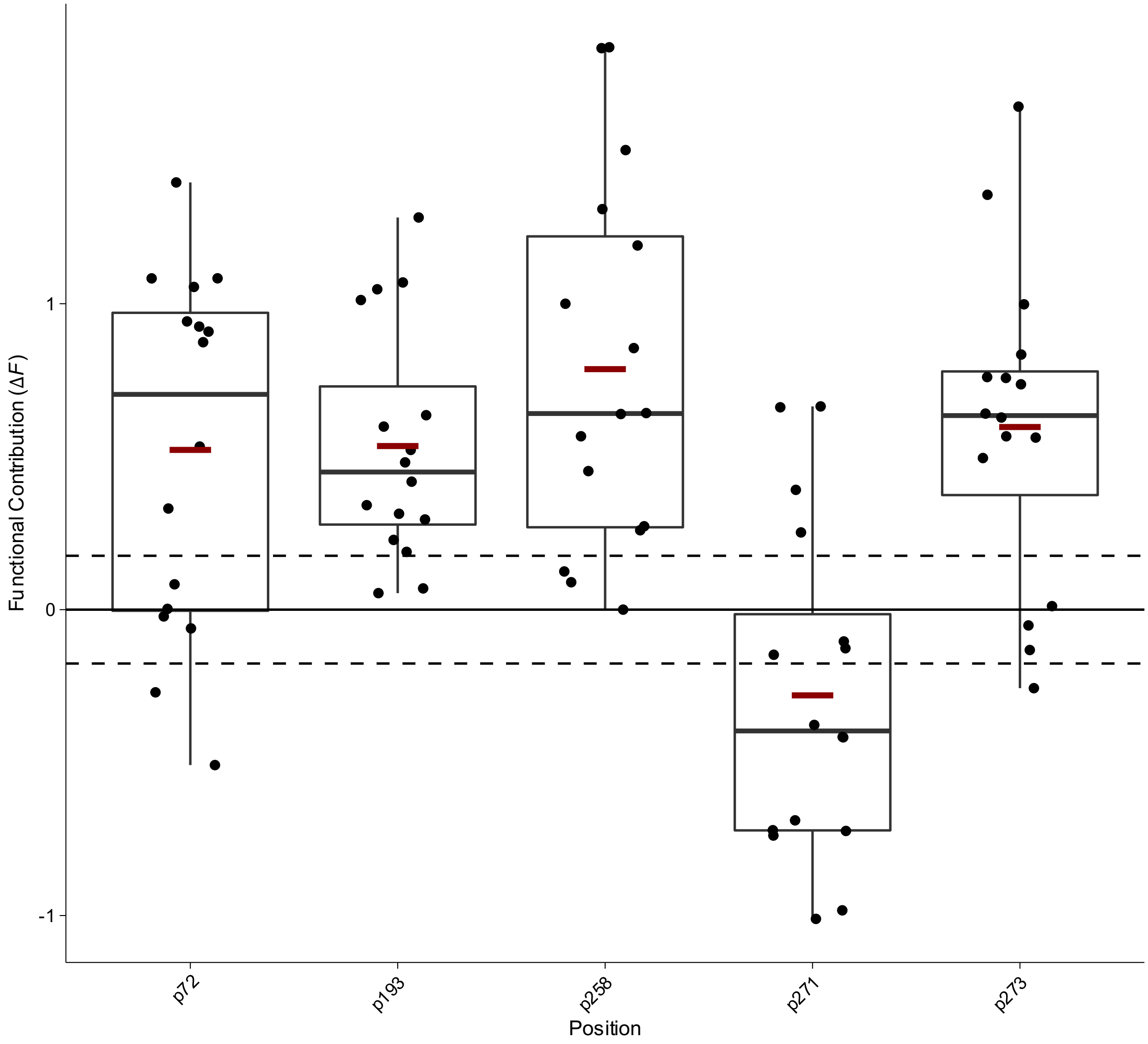


**Box Plot 23 | Functional contribution of positional mutations in Methyl Parathion Hydrolase (MPH) for methyl parathion in calcium metal conditions from Anderson *et al.***

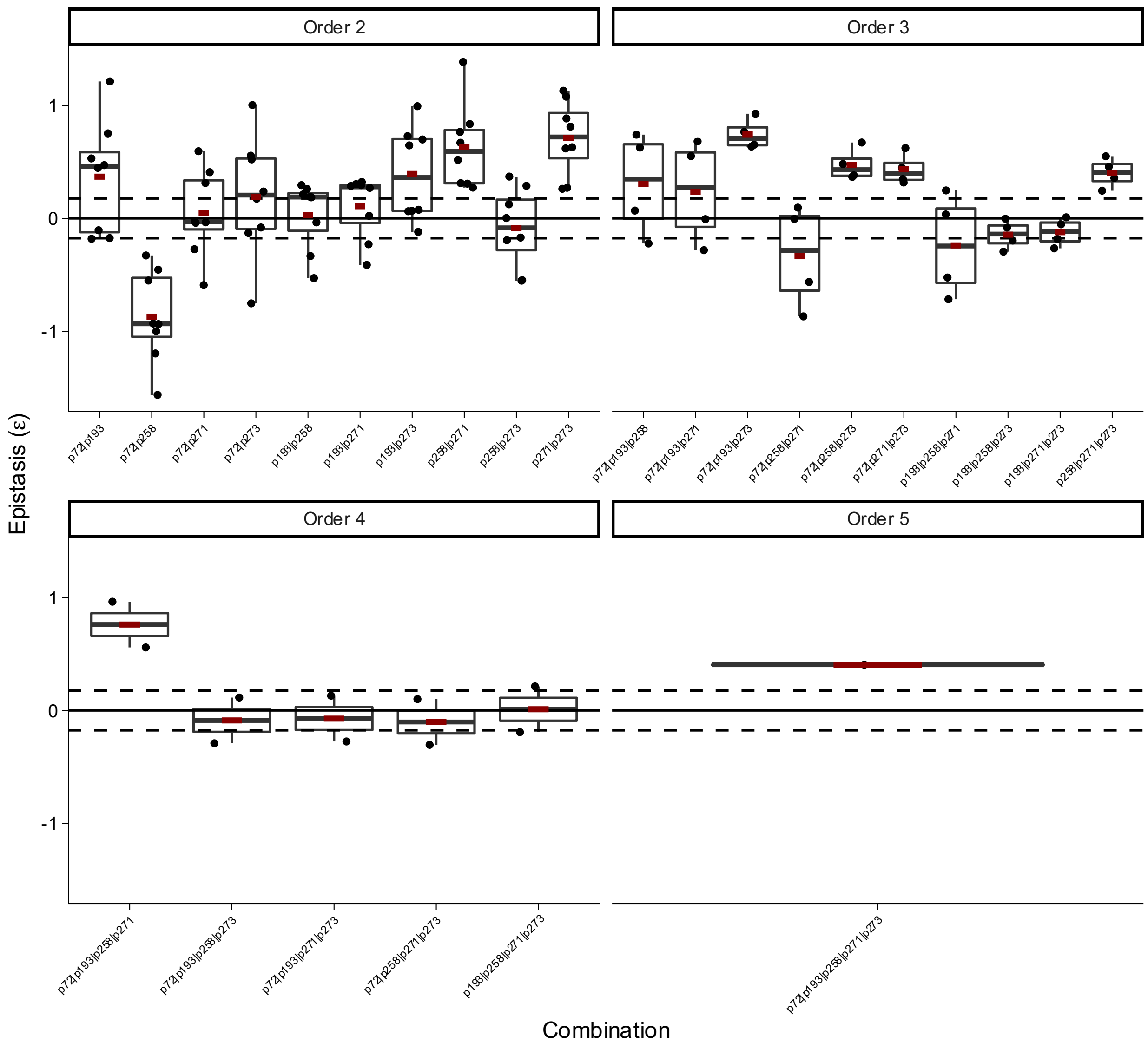


**Box Plot 24 | Epistasis of combinatorial mutations in Methyl Parathion Hydrolase (MPH) for methyl parathion in calcium metal conditions from Anderson *et al.***

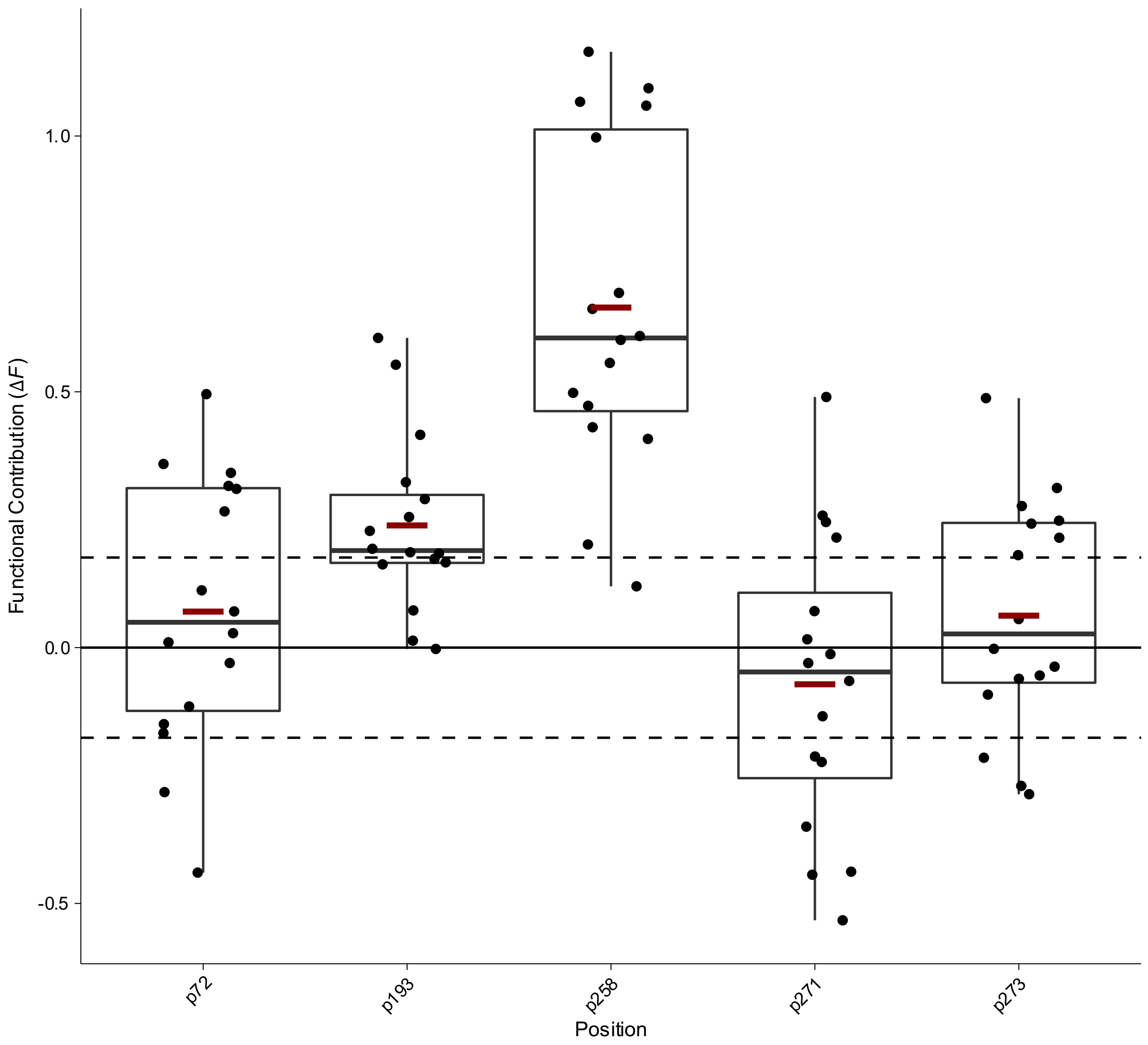


**Box Plot 25 | Functional contribution of positional mutations in Methyl Parathion Hydrolase (MPH) for methyl parathion in cadmium metal conditions from Anderson *et al.***

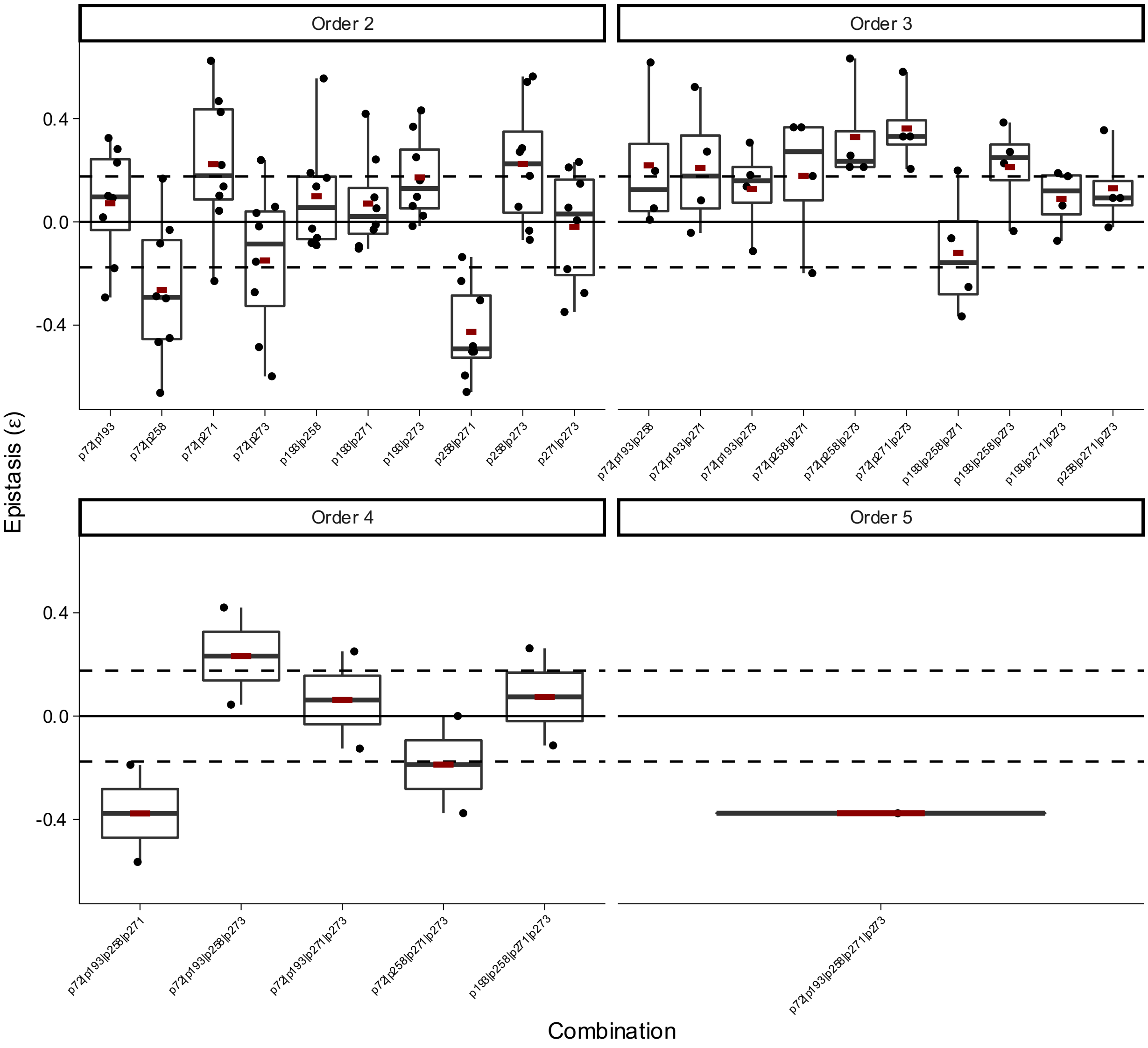


**Box Plot 26 | Epistasis of combinatorial mutations in Methyl Parathion Hydrolase (MPH) for methyl parathion in cadmium metal conditions from Anderson *et al.***

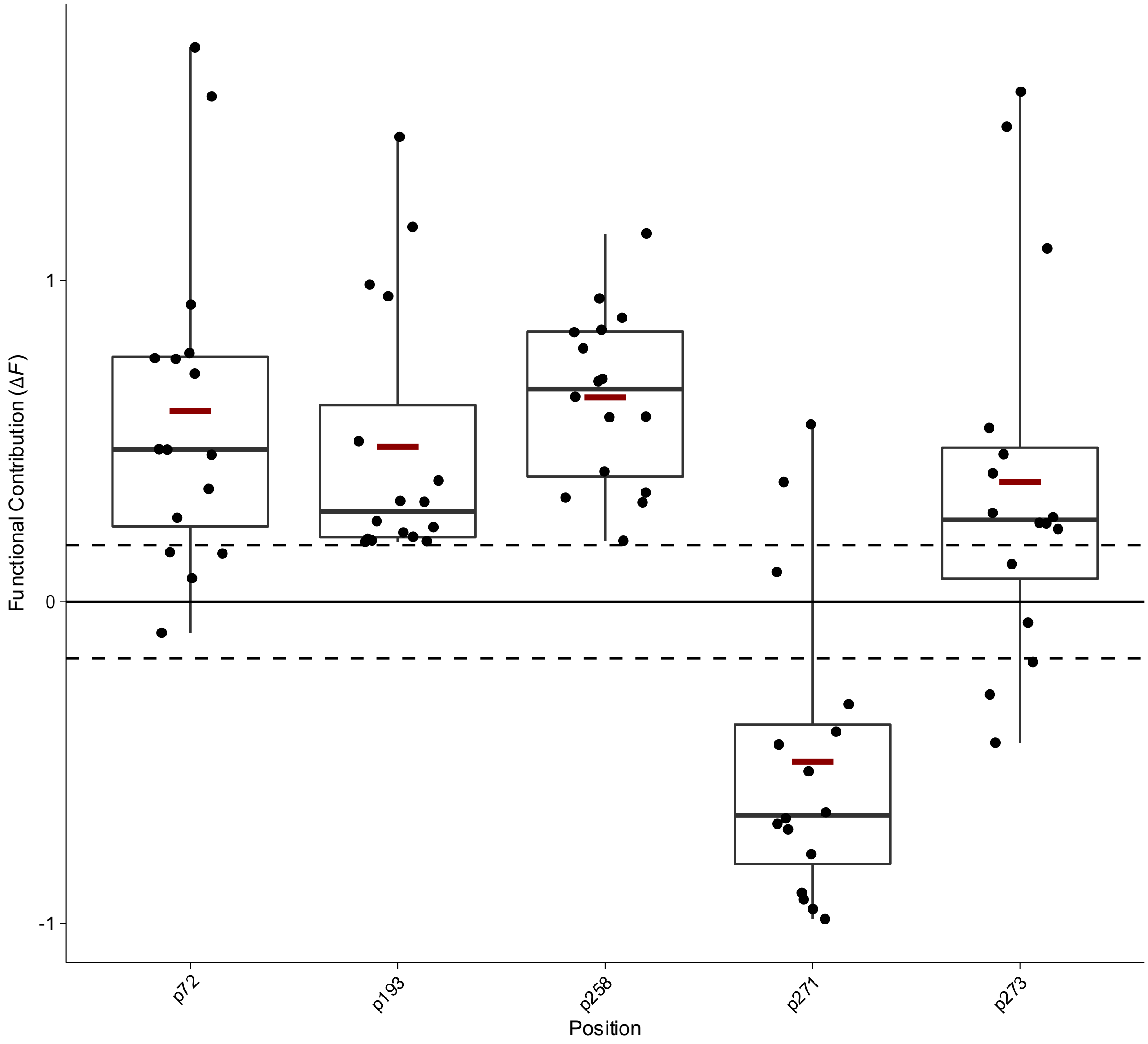


**Box Plot 27 | Functional contribution of positional mutations in Methyl Parathion Hydrolase (MPH) for methyl parathion in cobalt metal conditions from Anderson *et al.***

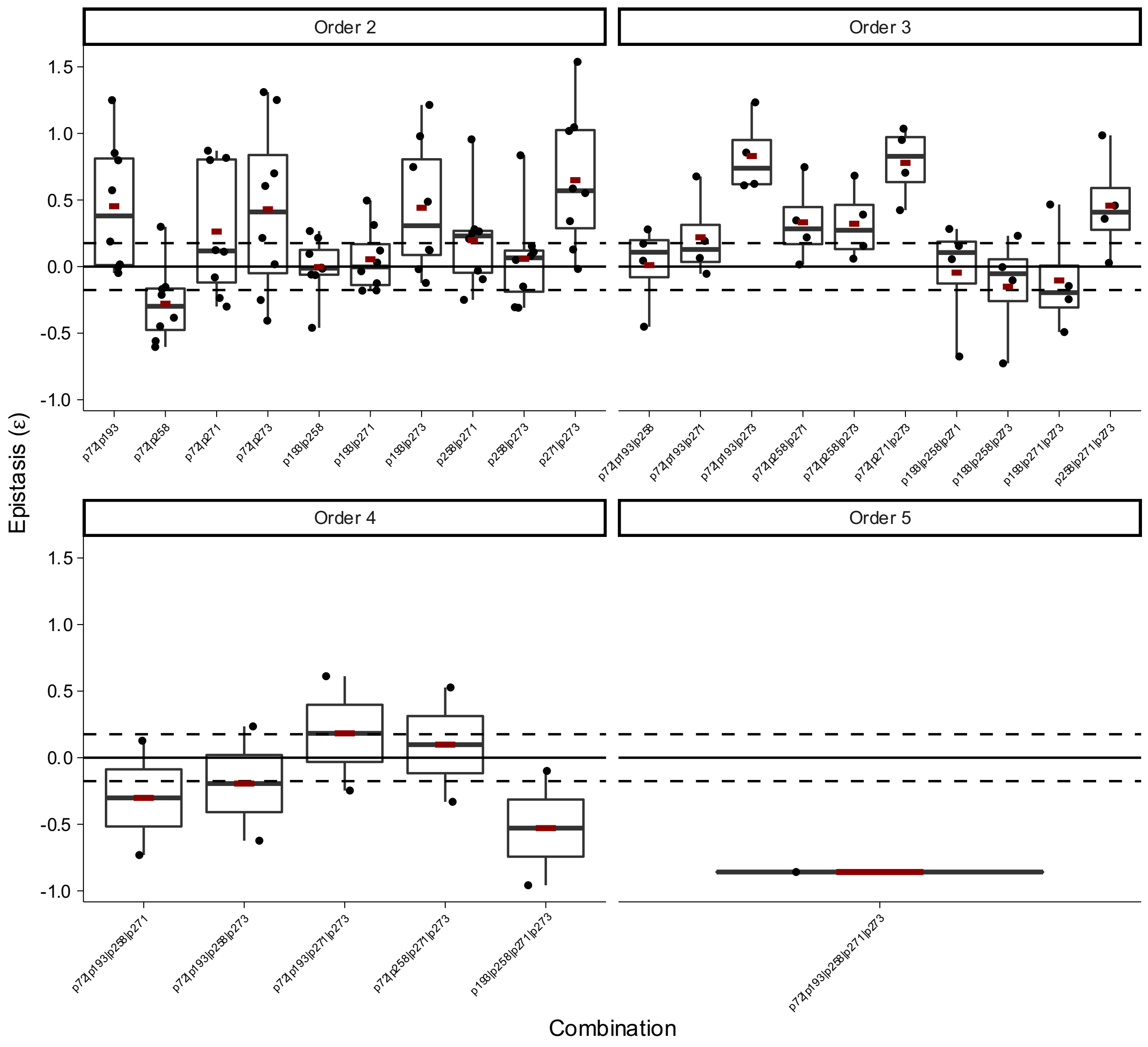


**Box Plot 28 | Epistasis of combinatorial mutations in Methyl Parathion Hydrolase (MPH) for methyl parathion in cobalt metal conditions from Anderson *et al.***

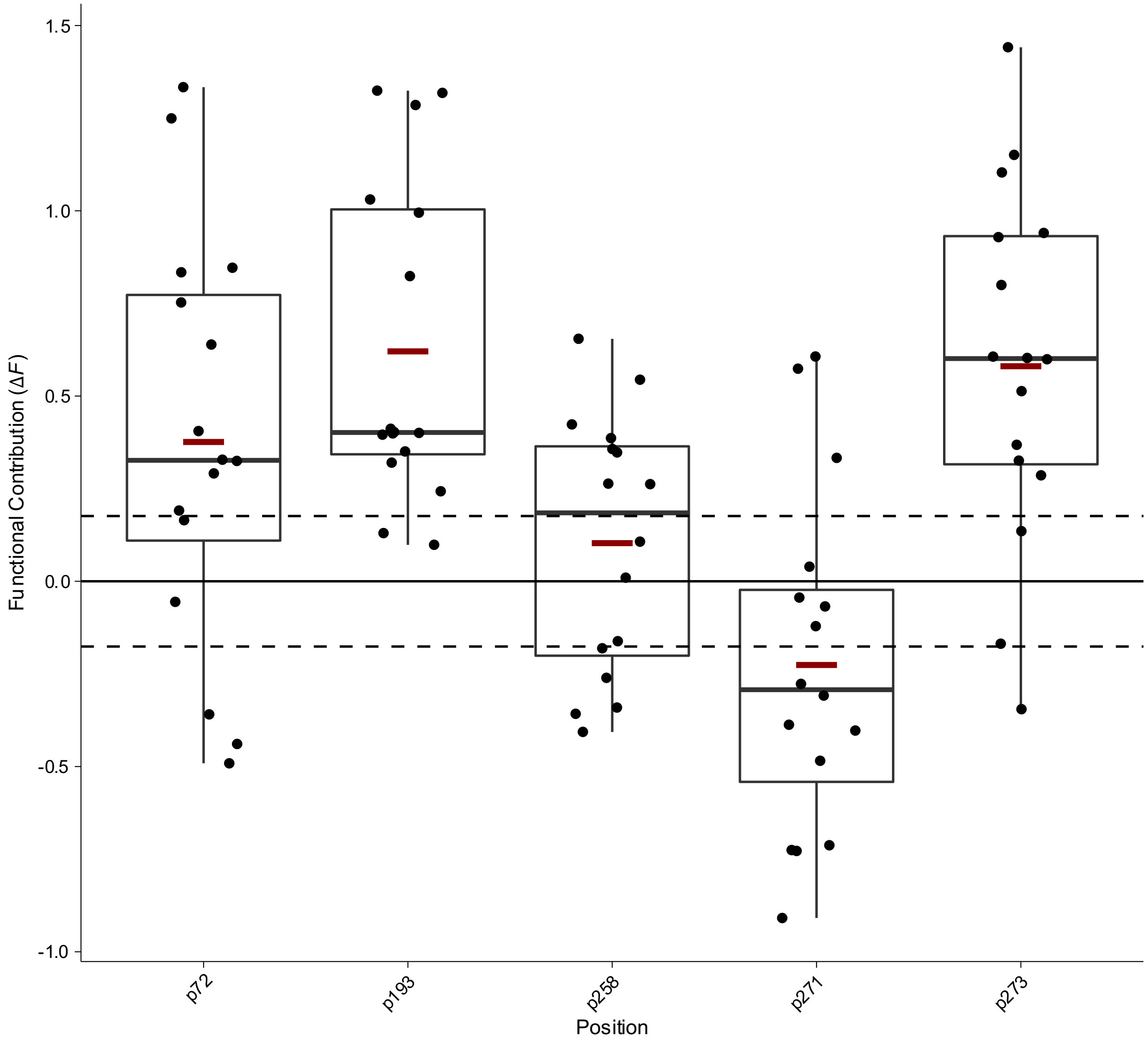


**Box Plot 29 | Functional contribution of positional mutations in Methyl Parathion Hydrolase (MPH) for methyl parathion in copper metal conditions from Anderson *et al.***

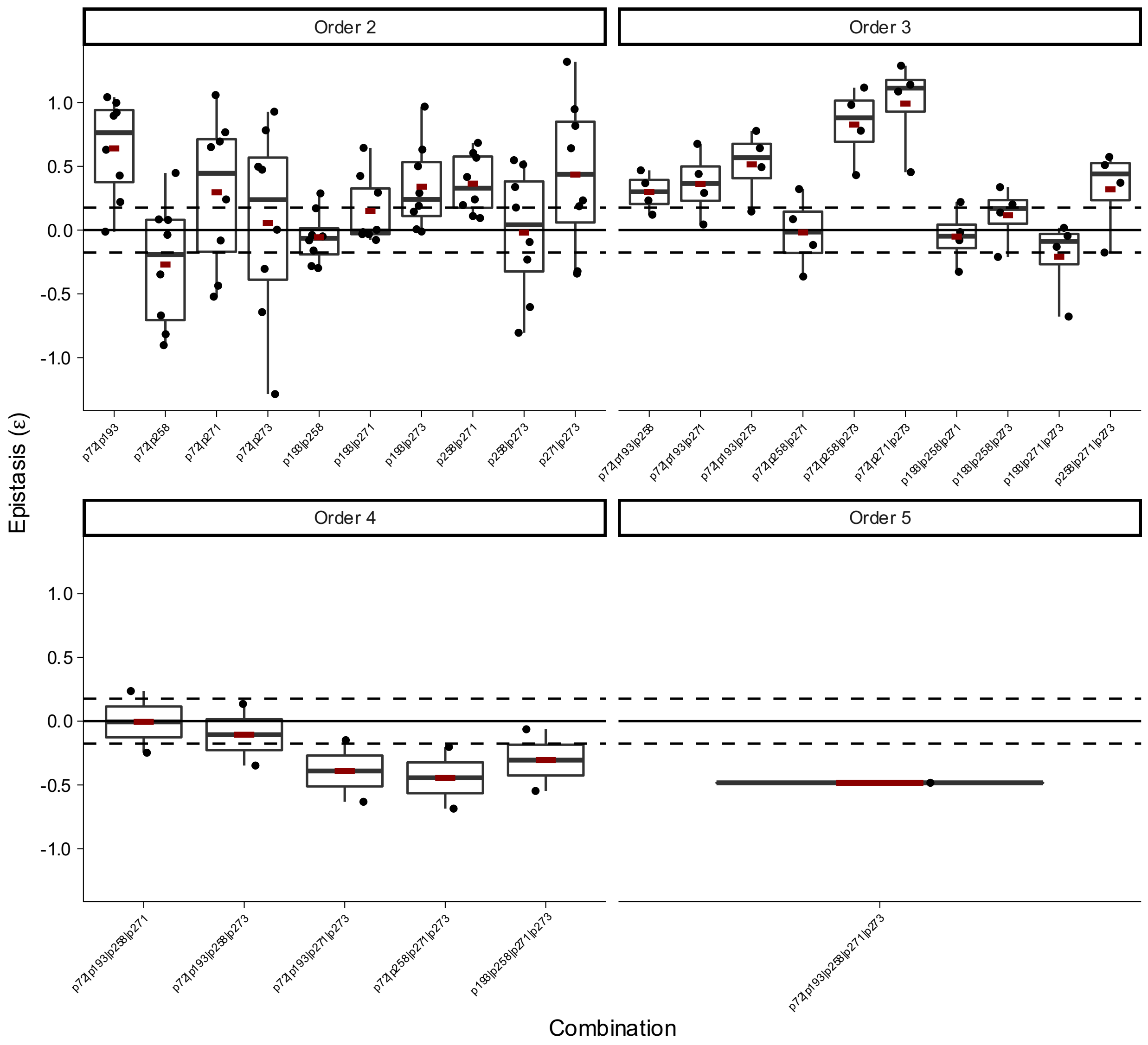


**Box Plot 30 | Epistasis of combinatorial mutations in Methyl Parathion Hydrolase (MPH) for methyl parathion in copper metal conditions from Anderson *et al.***

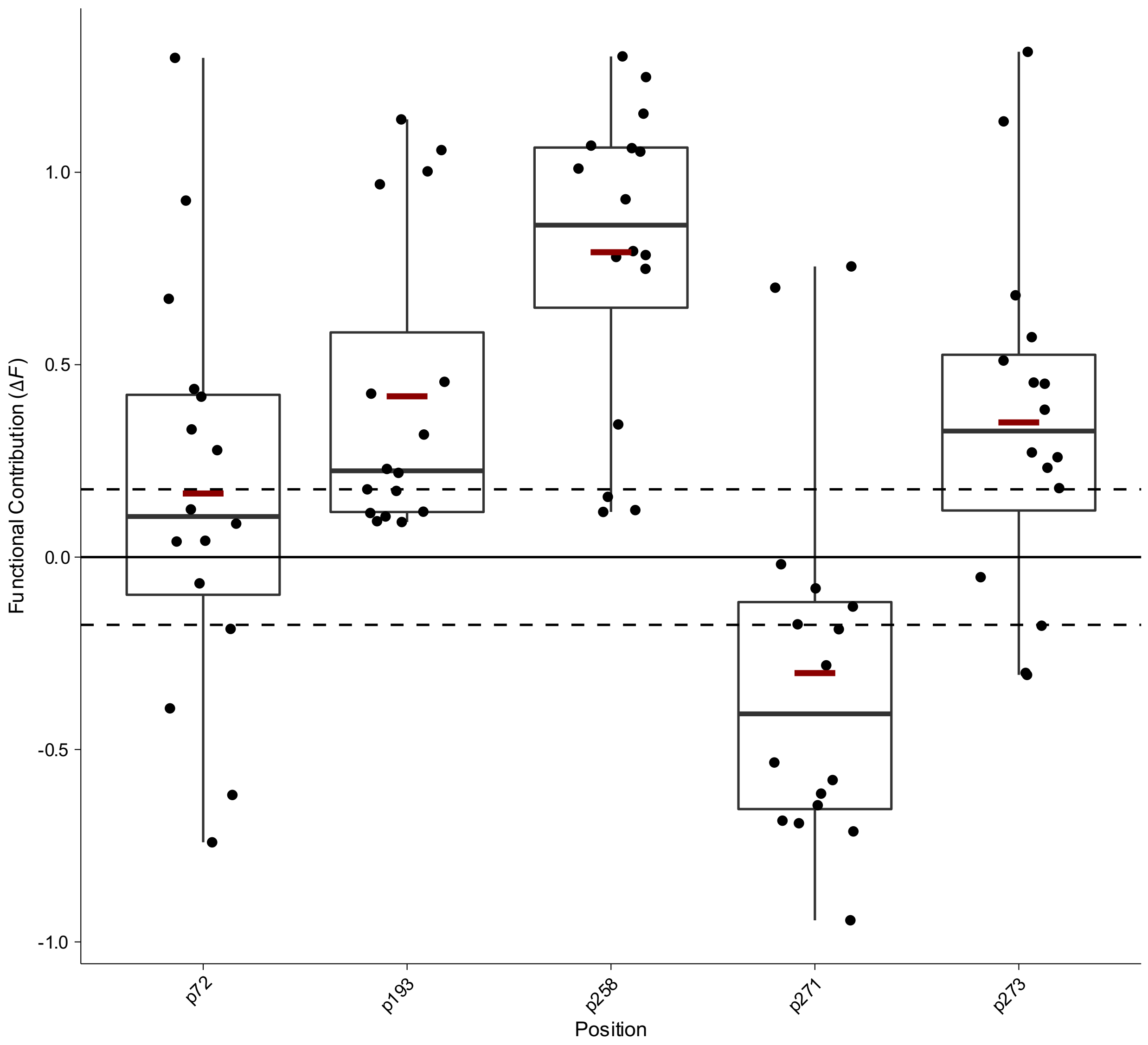


**Box Plot 31 | Functional contribution of positional mutations in Methyl Parathion Hydrolase (MPH) for methyl parathion in magnesium metal conditions from Anderson *et al.***

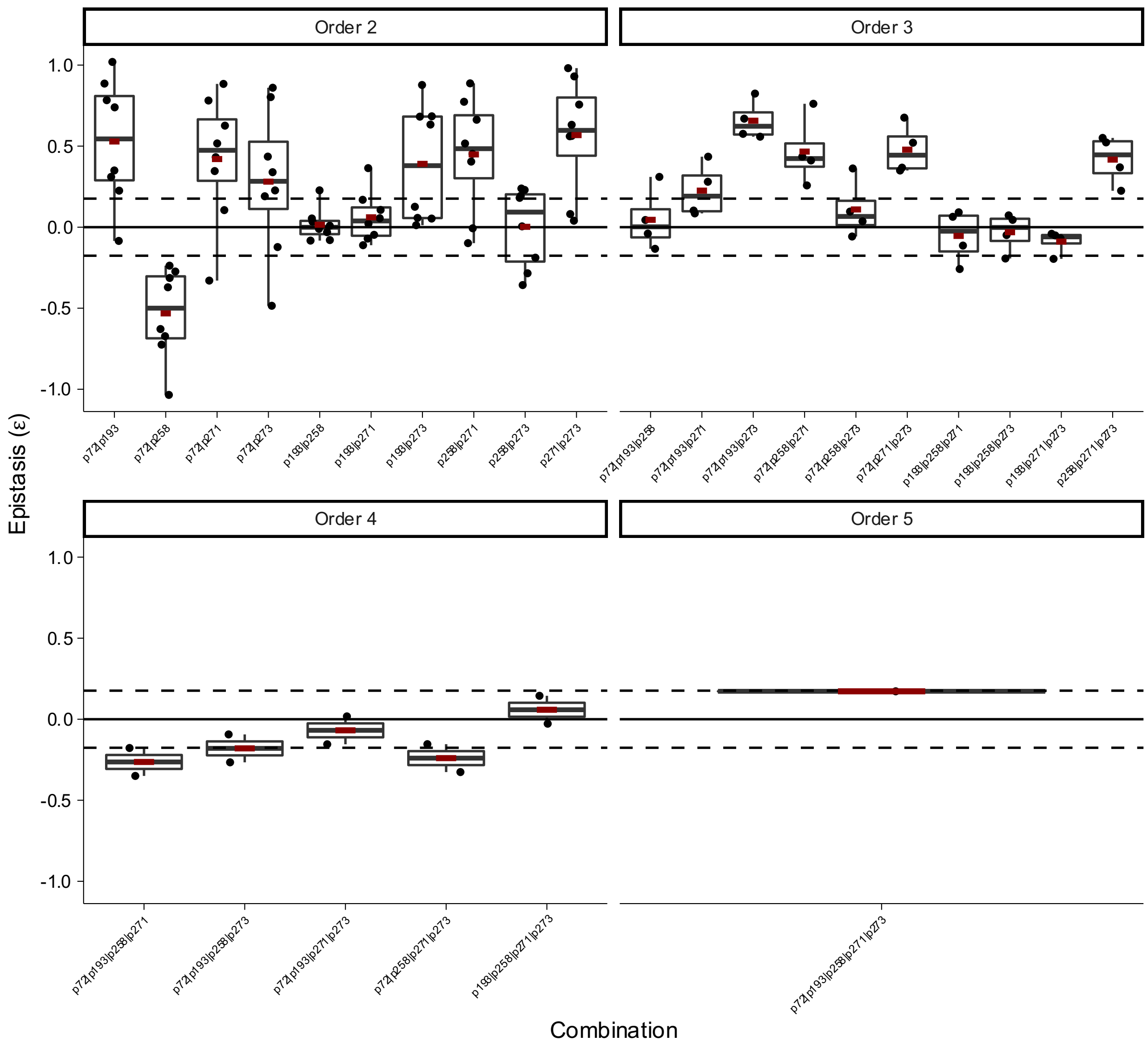


**Box Plot 32 | Epistasis of combinatorial mutations in Methyl Parathion Hydrolase (MPH) for methyl parathion in magnesium metal conditions from Anderson *et al.***

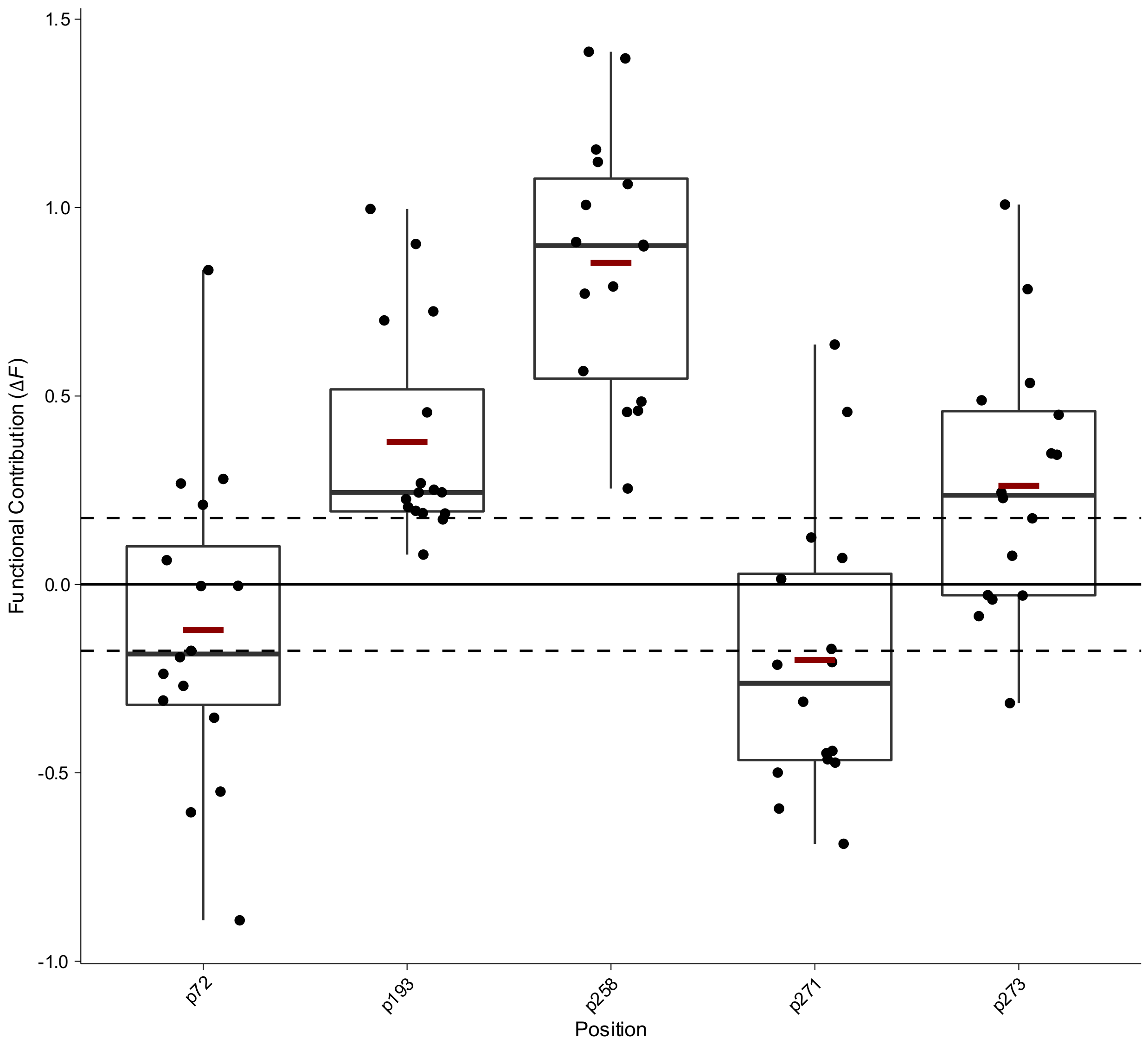


**Box Plot 33 | Functional contribution of positional mutations in Methyl Parathion Hydrolase (MPH) for methyl parathion in manganese metal conditions from Anderson *et al.***

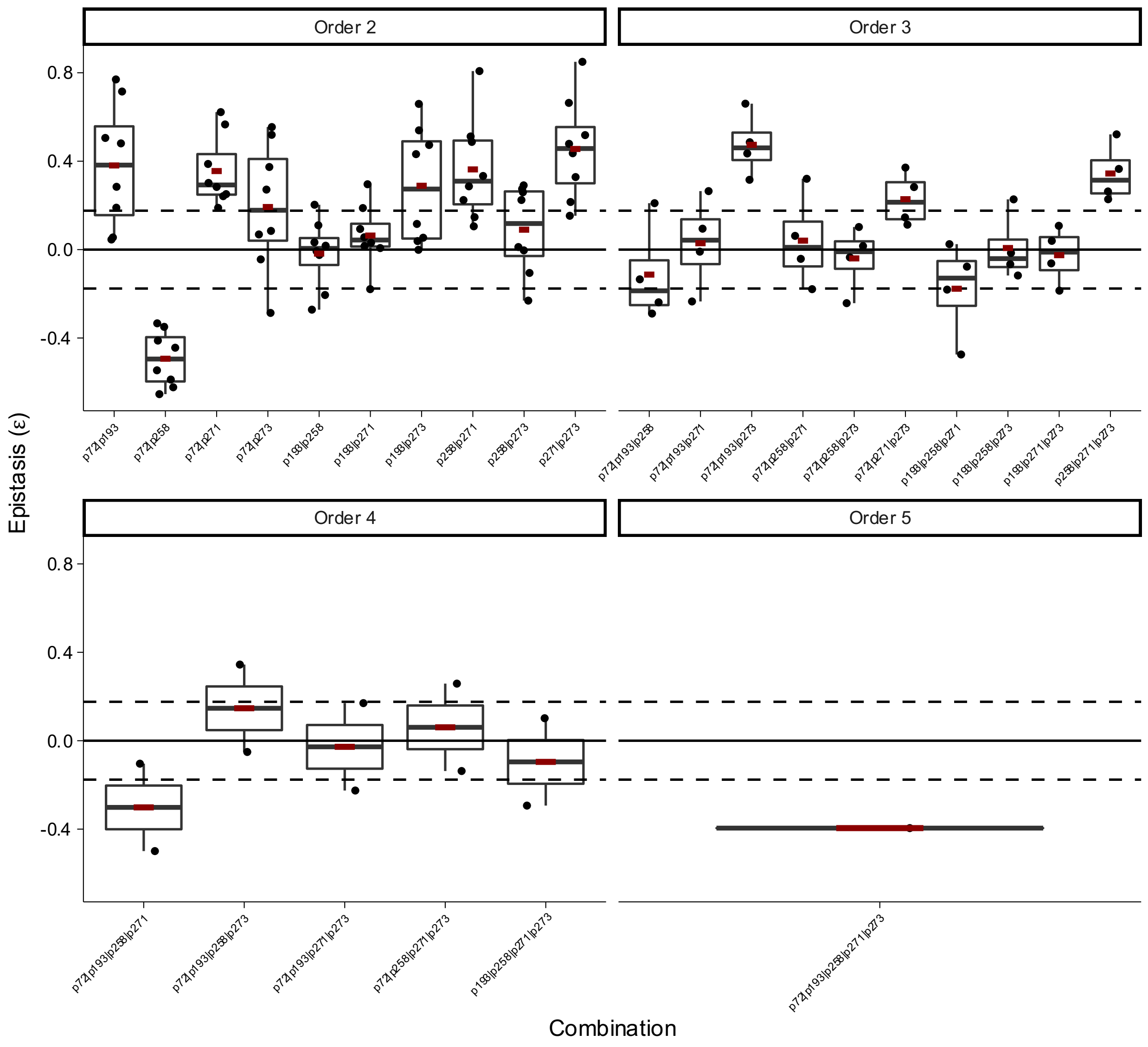


**Box Plot 34 | Epistasis of combinatorial mutations in Methyl Parathion Hydrolase (MPH) for methyl parathion in manganese metal conditions from Anderson *et al.***

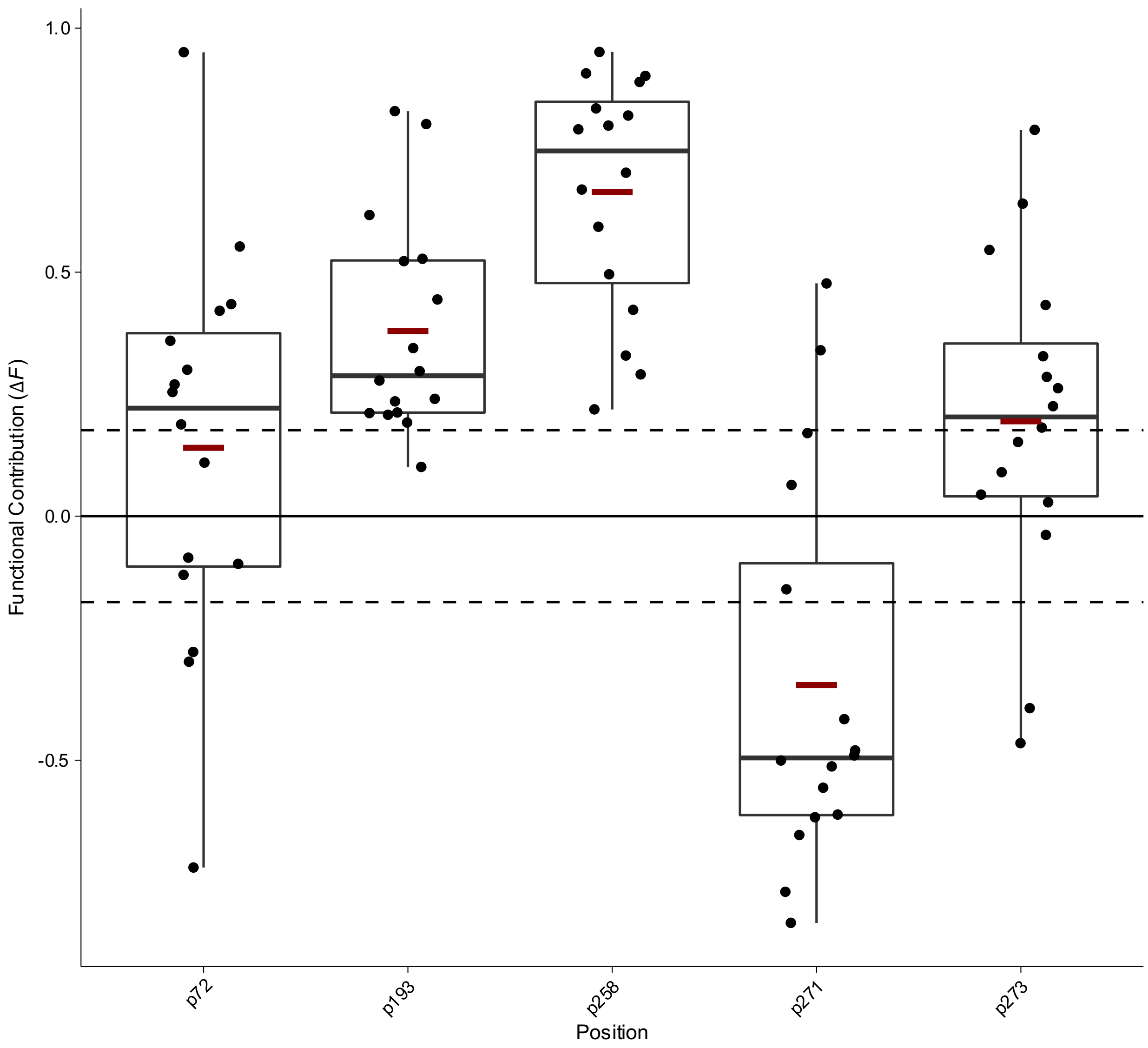


**Box Plot 35 | Functional contribution of positional mutations in Methyl Parathion Hydrolase (MPH) for methyl parathion in nickel metal conditions from Anderson *et al.***

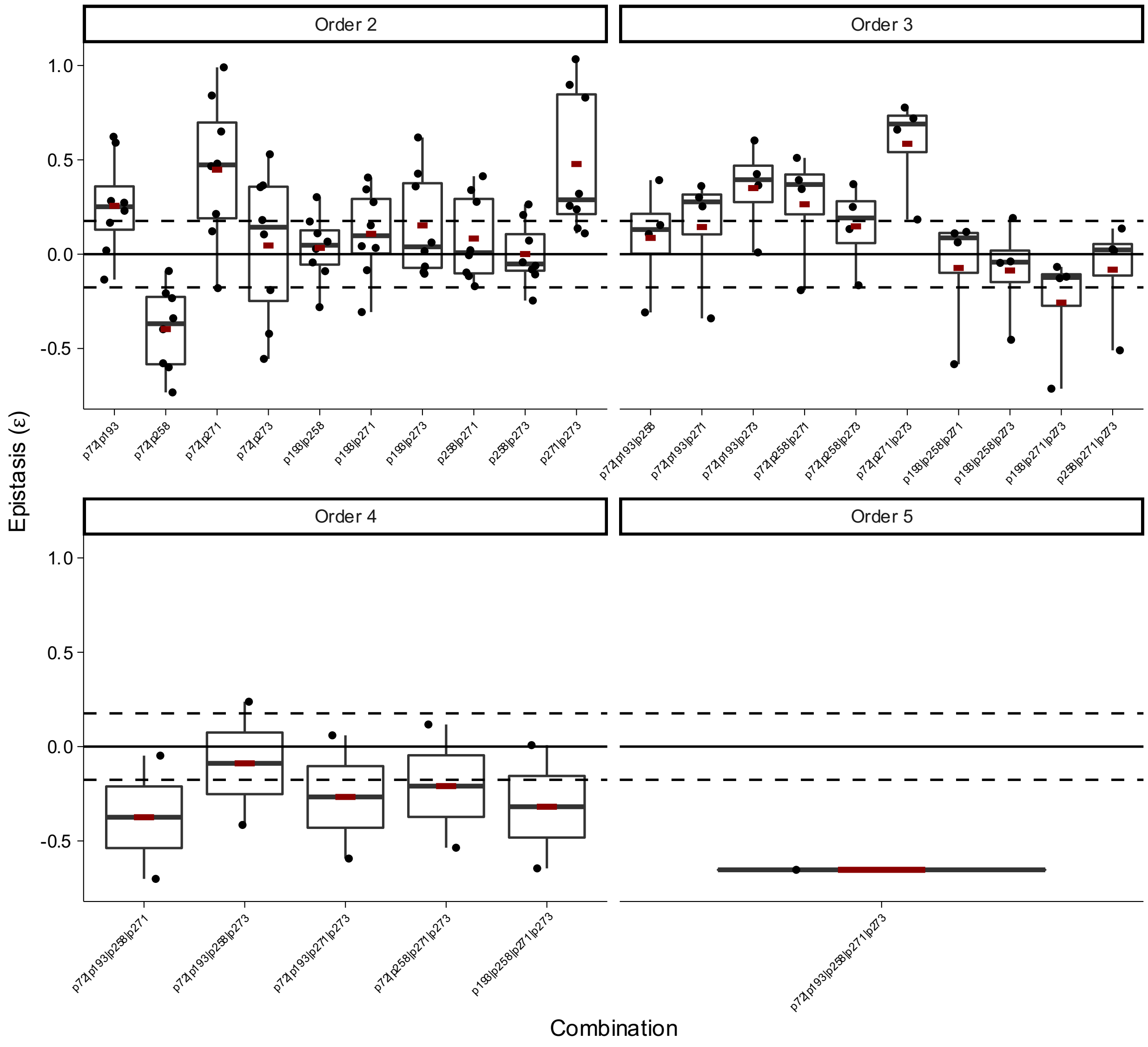


**Box Plot 36 | Epistasis of combinatorial mutations in Methyl Parathion Hydrolase (MPH) for methyl parathion in nickel metal conditions from Anderson *et al.***

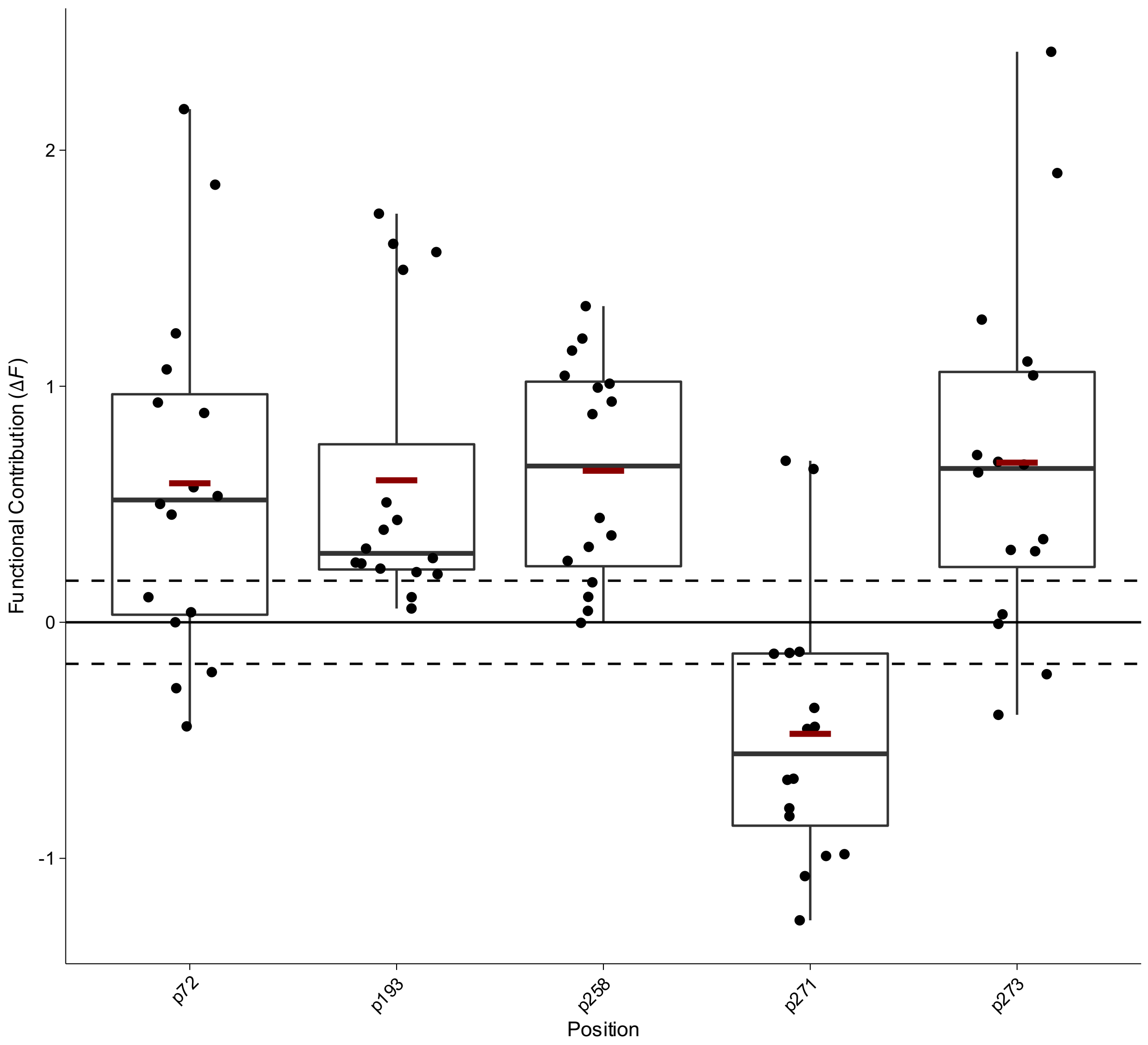


**Box Plot 37 | Functional contribution of positional mutations in Methyl Parathion Hydrolase (MPH) for methyl parathion in zinc metal conditions from Anderson *et al.***

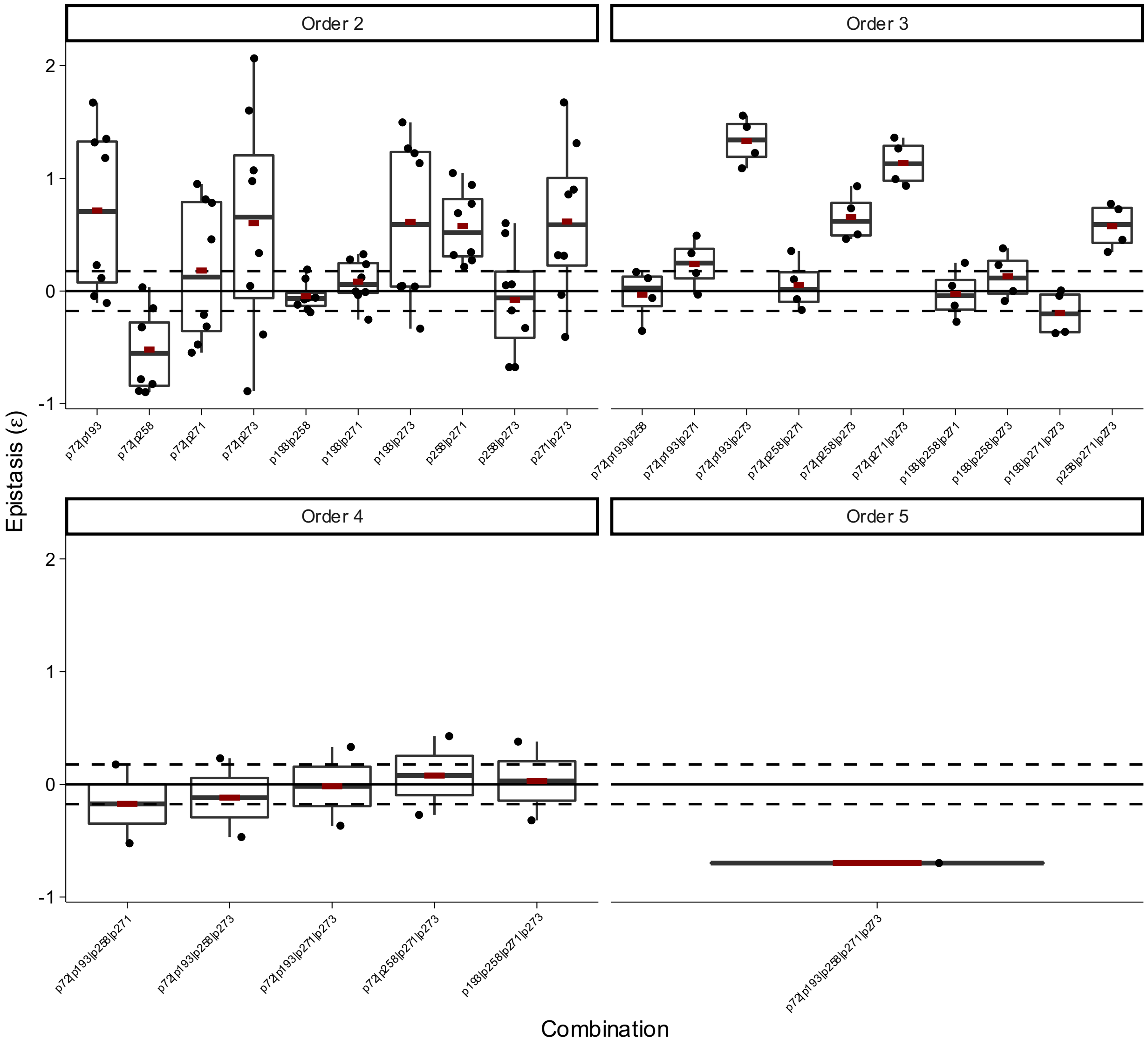


**Box Plot 38 | Epistasis of combinatorial mutations in Methyl Parathion Hydrolase (MPH) for methyl parathion in zinc metal conditions from Anderson *et al.***

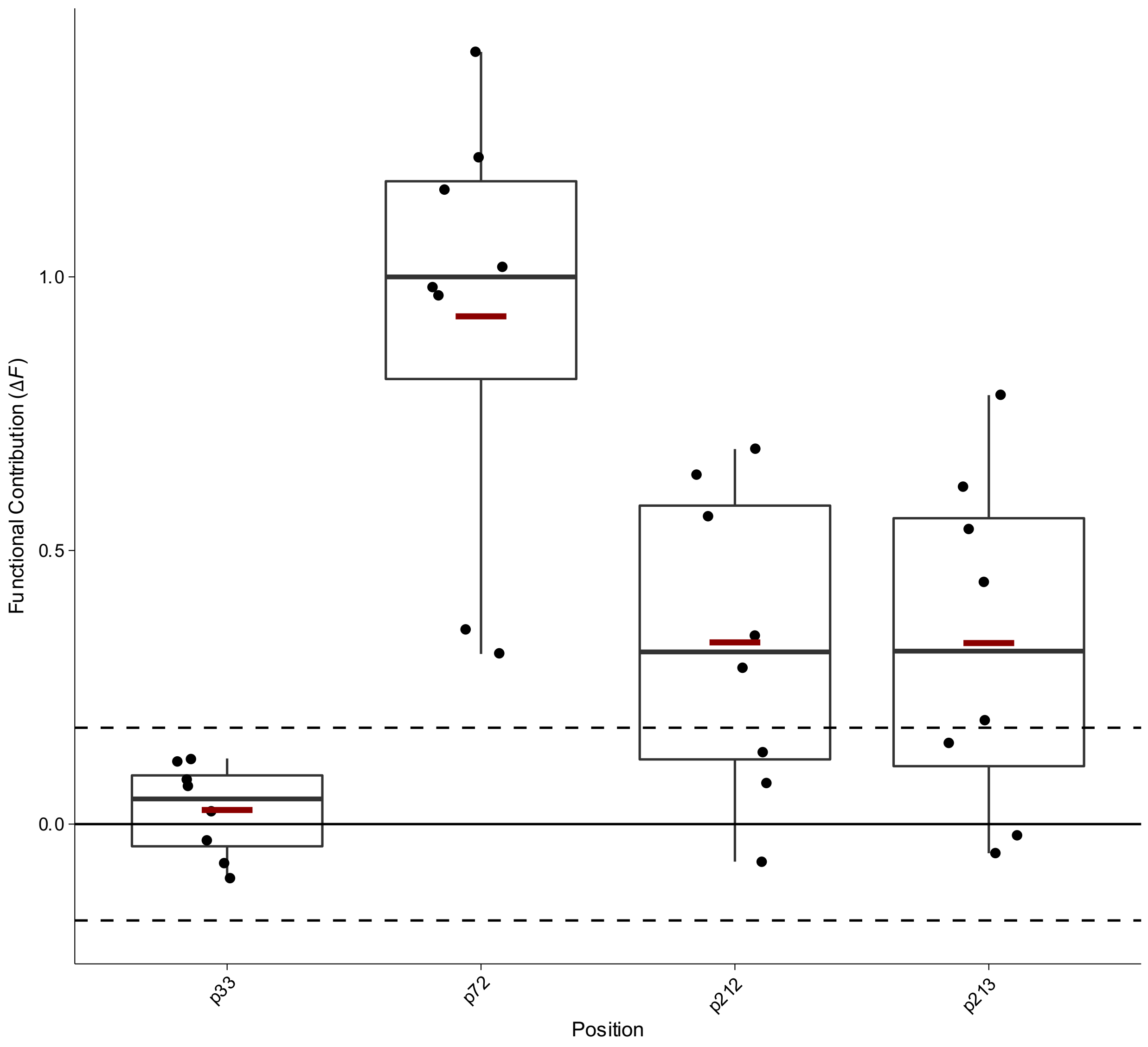


**Box Plot 39 | Functional contribution of positional mutations in beta lactamase OXA-48 for ceftazidime hydrolysis in trajectory 1 from Fröhlich *et al.***

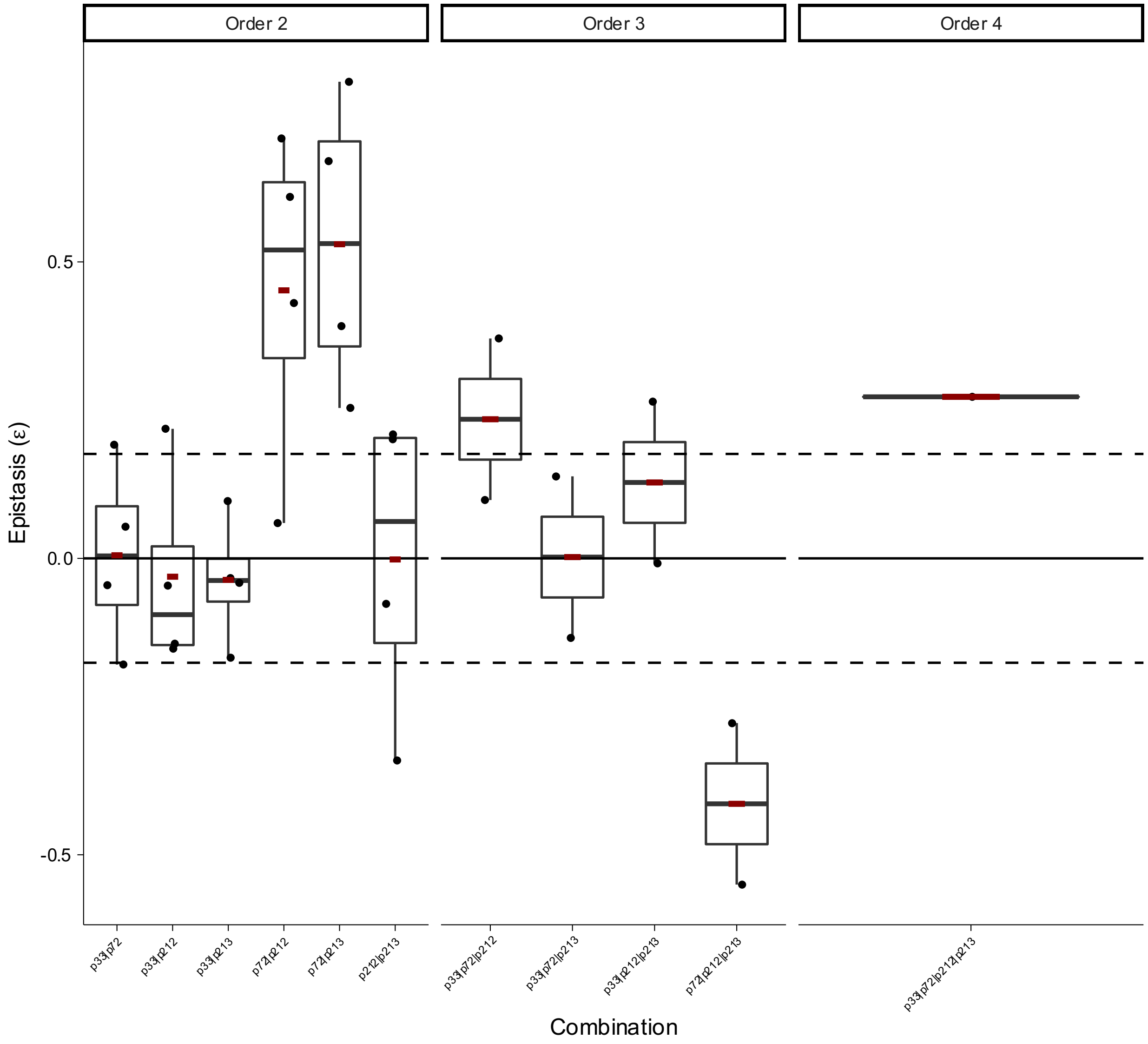


**Box Plot 40 | Epistasis of combinatorial mutations in beta lactamase OXA-48 for ceftazidime hydrolysis in trajectory 1 from Fröhlich *et al.***

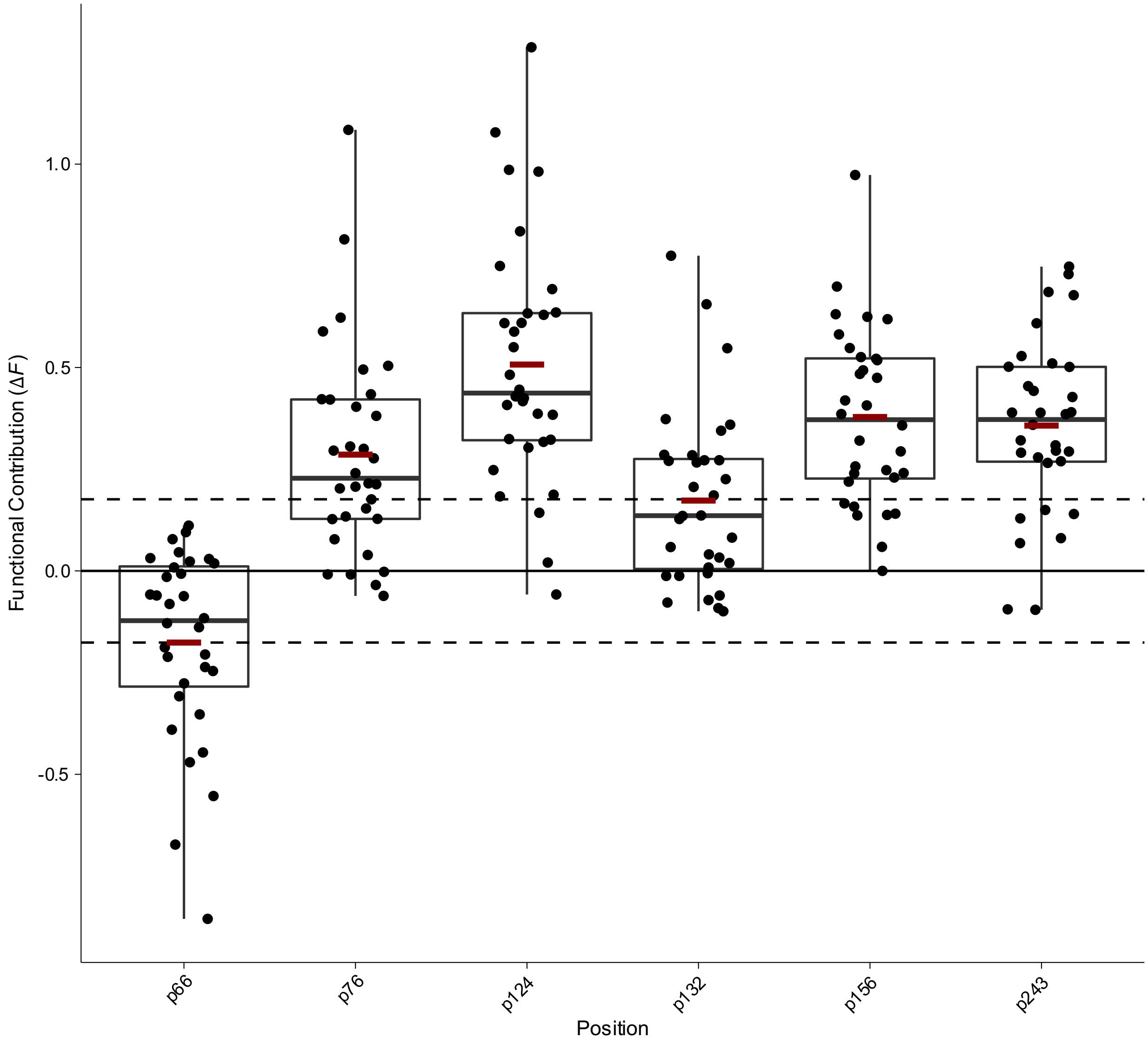


**Box Plot 41 | Functional contribution of positional mutations in beta lactamase OXA-48 for ceftazidime hydrolysis in trajectory 2 from Fröhlich *et al.***

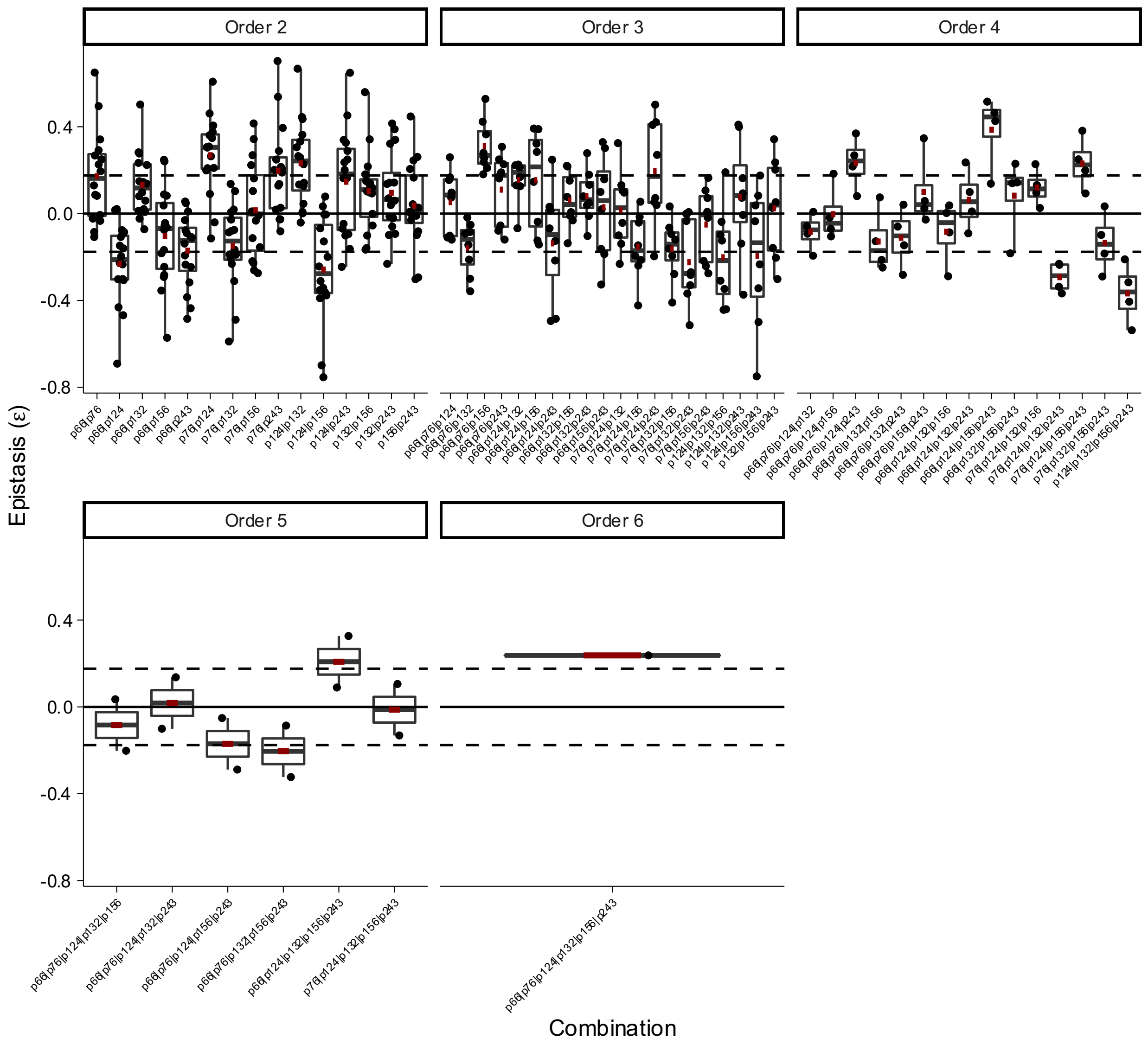


**Box Plot 42 | Epistasis of combinatorial mutations in beta lactamase OXA-48 for ceftazidime hydrolysis in trajectory 2 from Fröhlich *et al.***

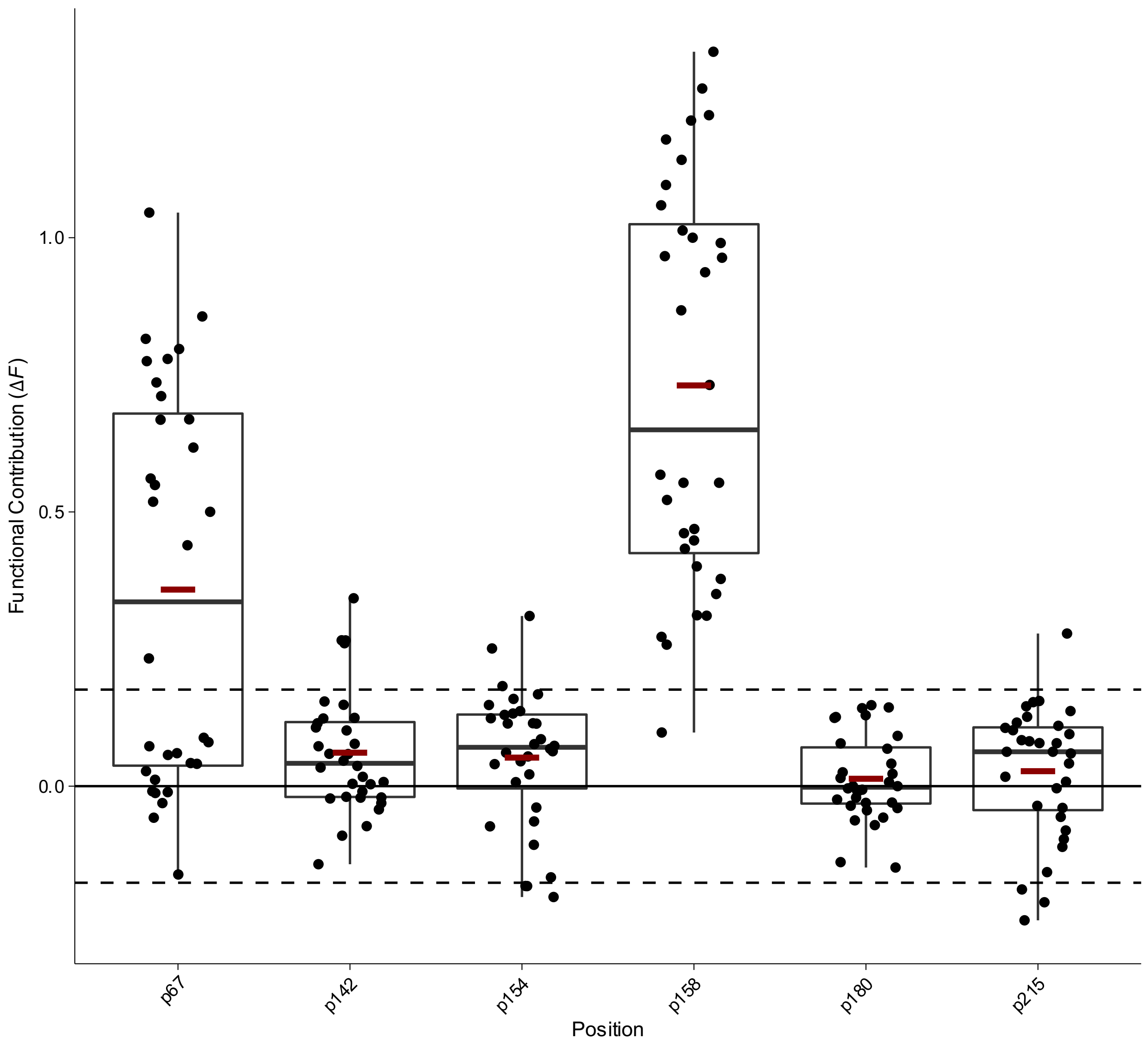


**Box Plot 43 | Functional contribution of positional mutations in beta lactamase OXA-48 for ceftazidime hydrolysis in trajectory 3 from Fröhlich *et al.***

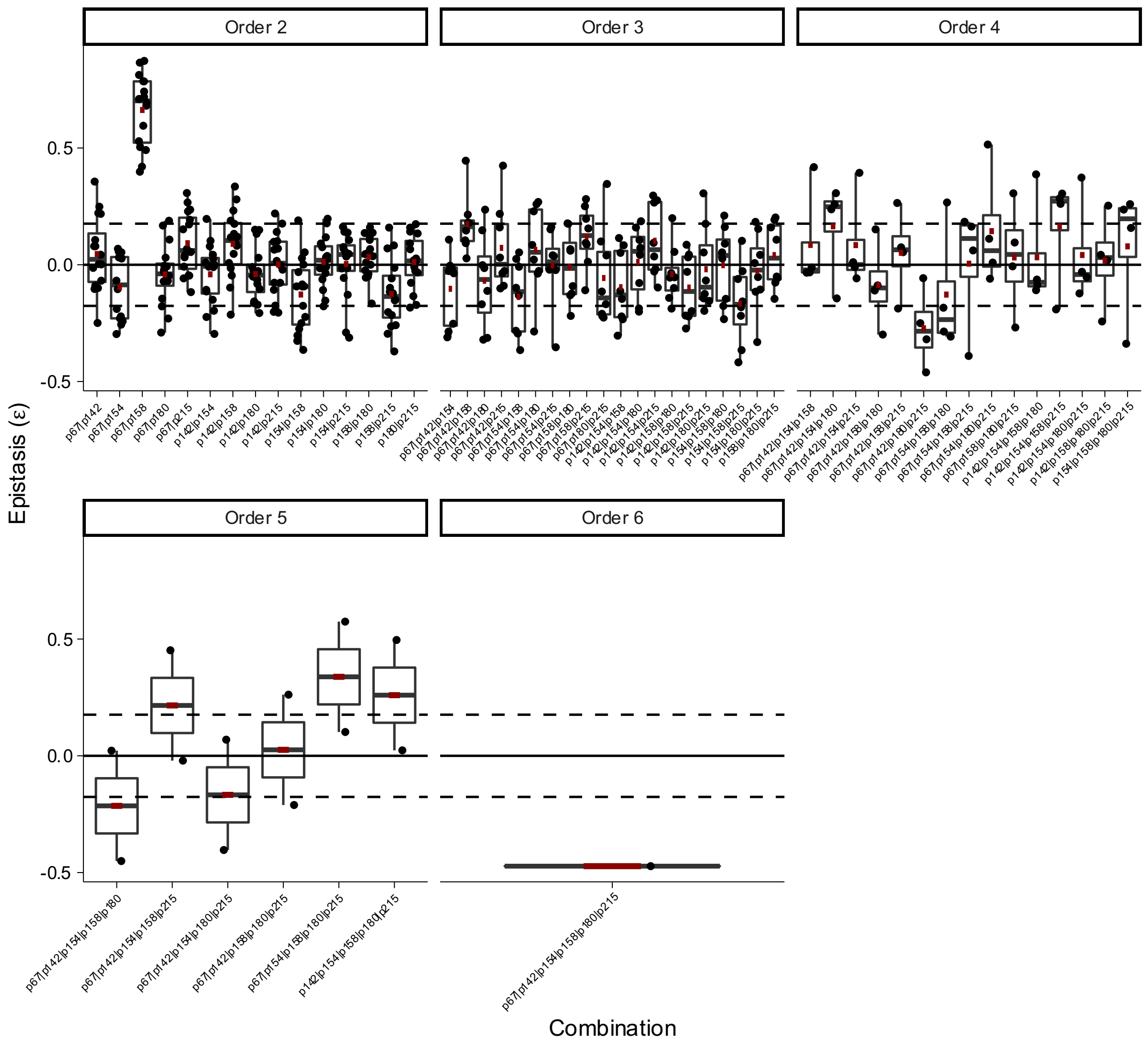


**Box Plot 44 | Epistasis of combinatorial mutations in beta lactamase OXA-48 for ceftazidime hydrolysis in trajectory 3 from Fröhlich *et al.***

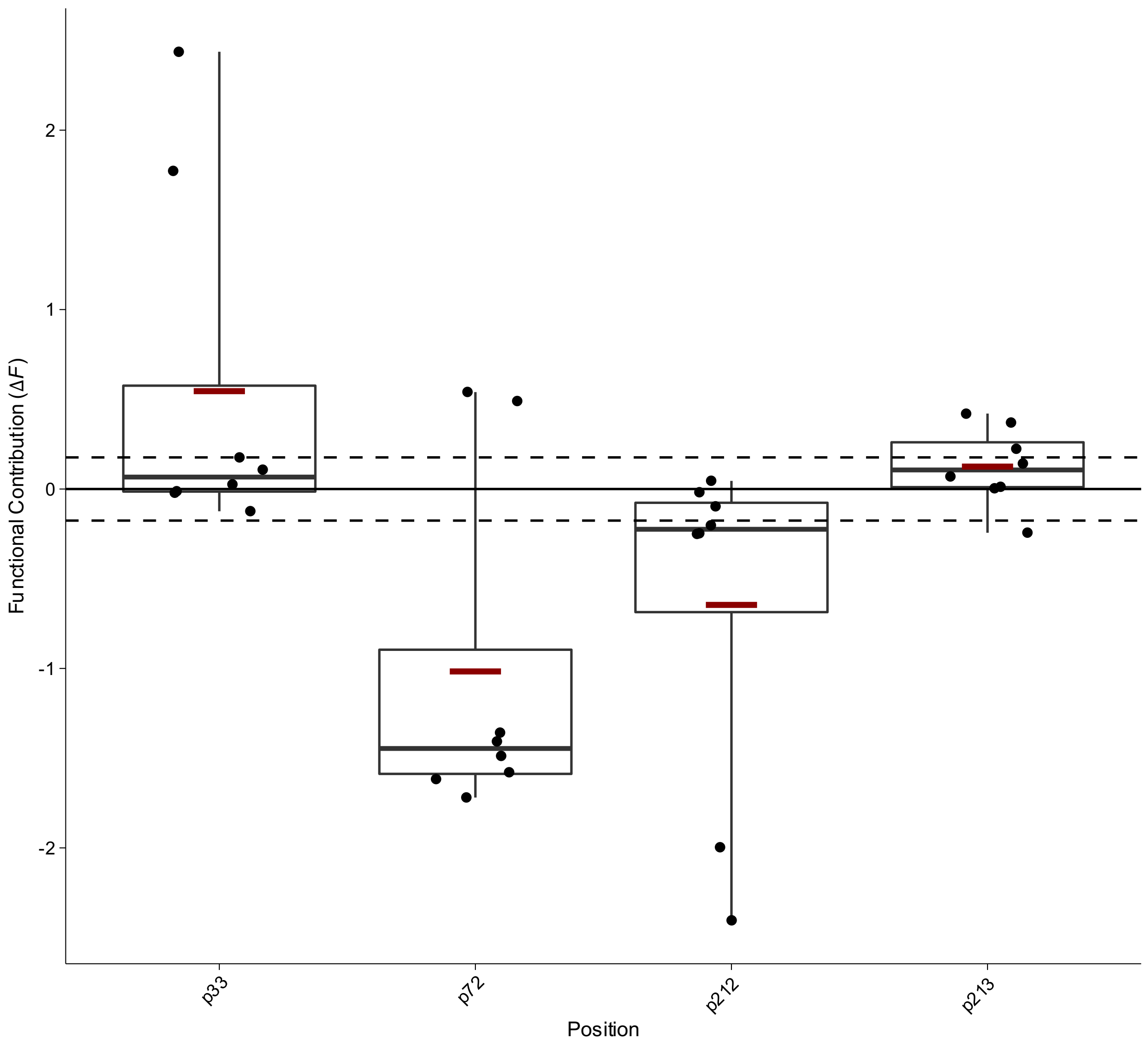


**Box Plot 45 | Functional contribution of positional mutations in beta lactamase OXA-48 for piperacillin hydrolysis in trajectory 1 from Fröhlich *et al.***

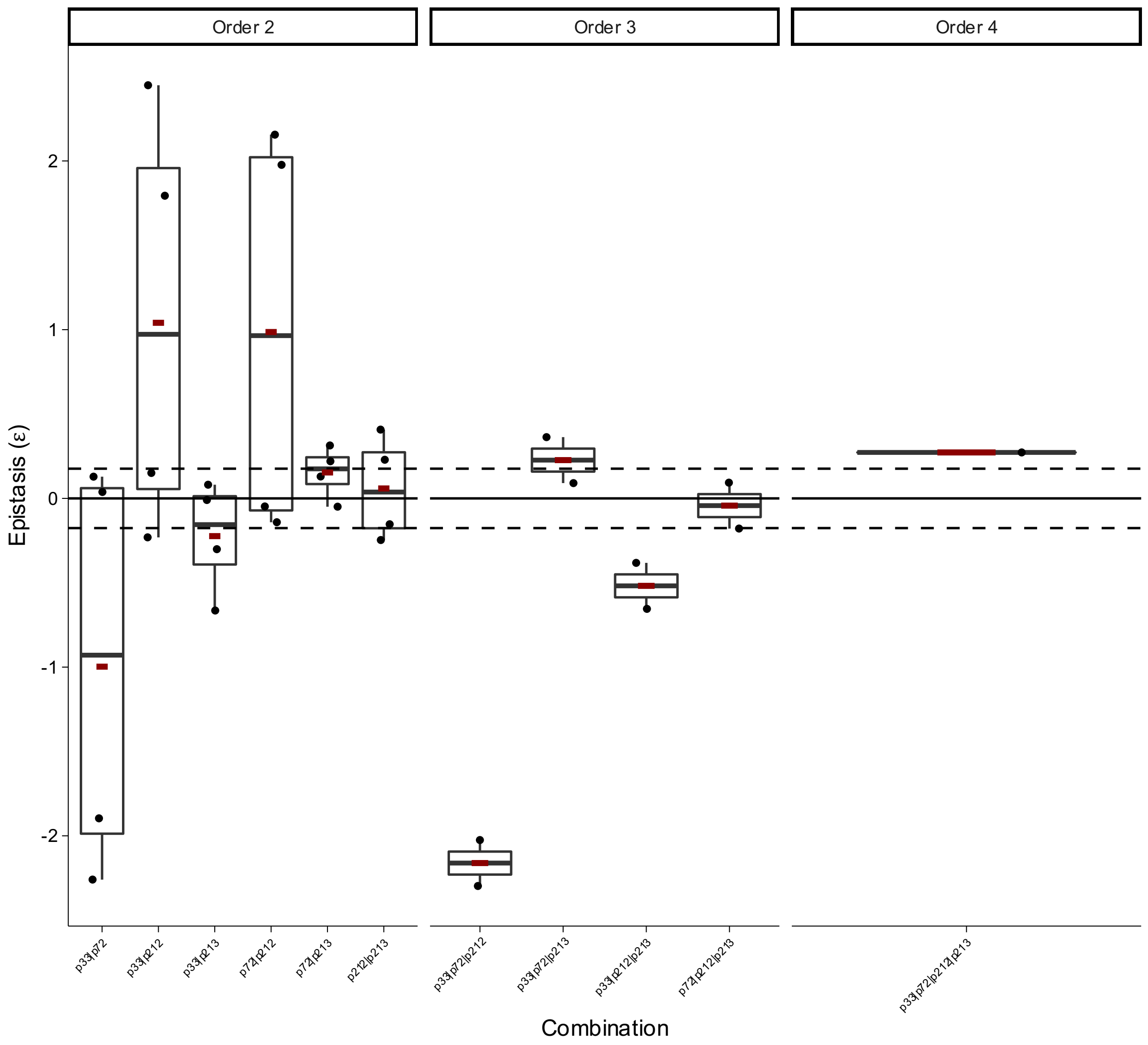


**Box Plot 46 | Epistasis of combinatorial mutations in beta lactamase OXA-48 for piperacillin hydrolysis in trajectory 1 from Fröhlich *et al.***

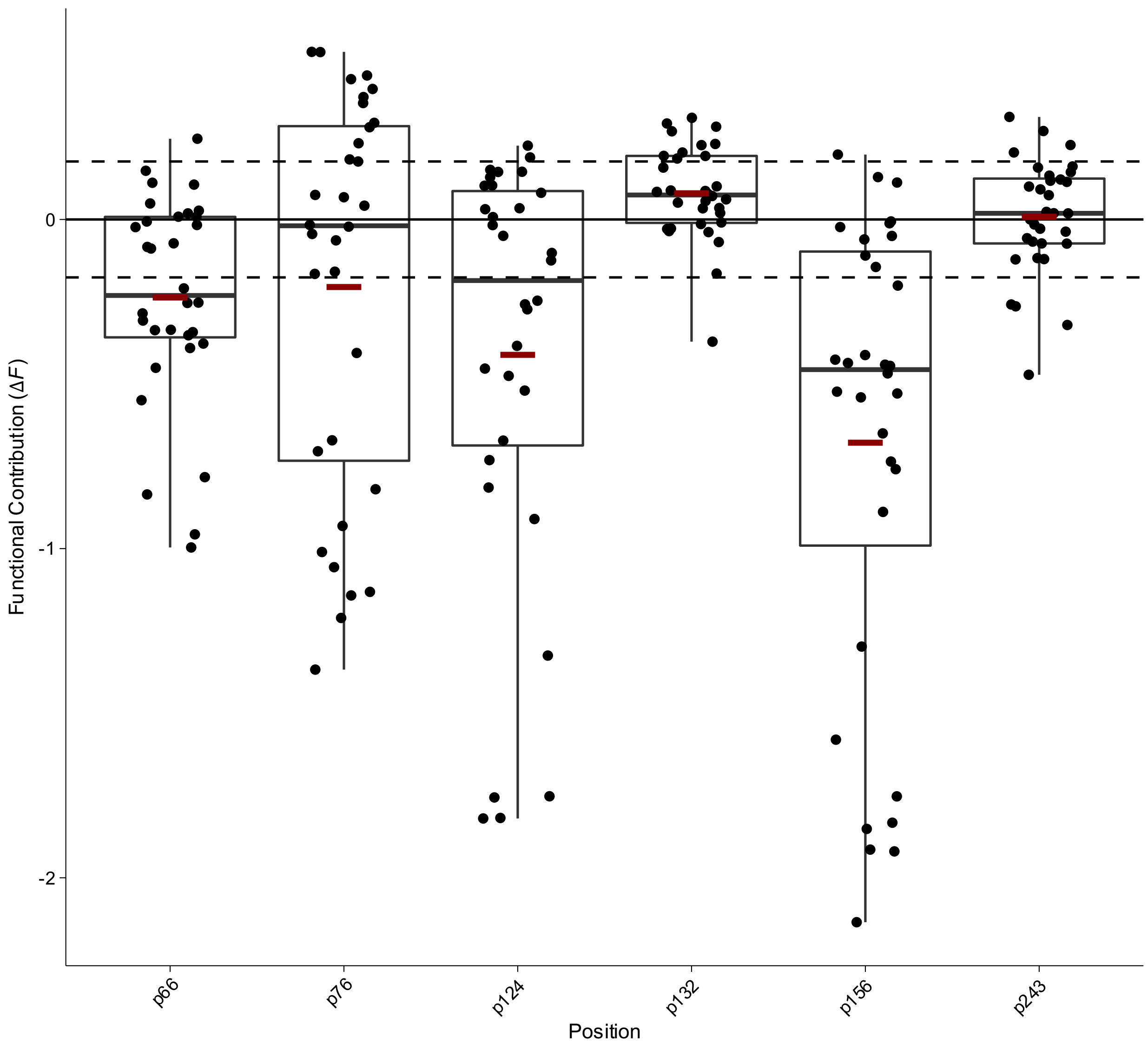


**Box Plot 47 | Functional contribution of positional mutations in beta lactamase OXA-48 for piperacillin hydrolysis in trajectory 2 from Fröhlich *et al.***

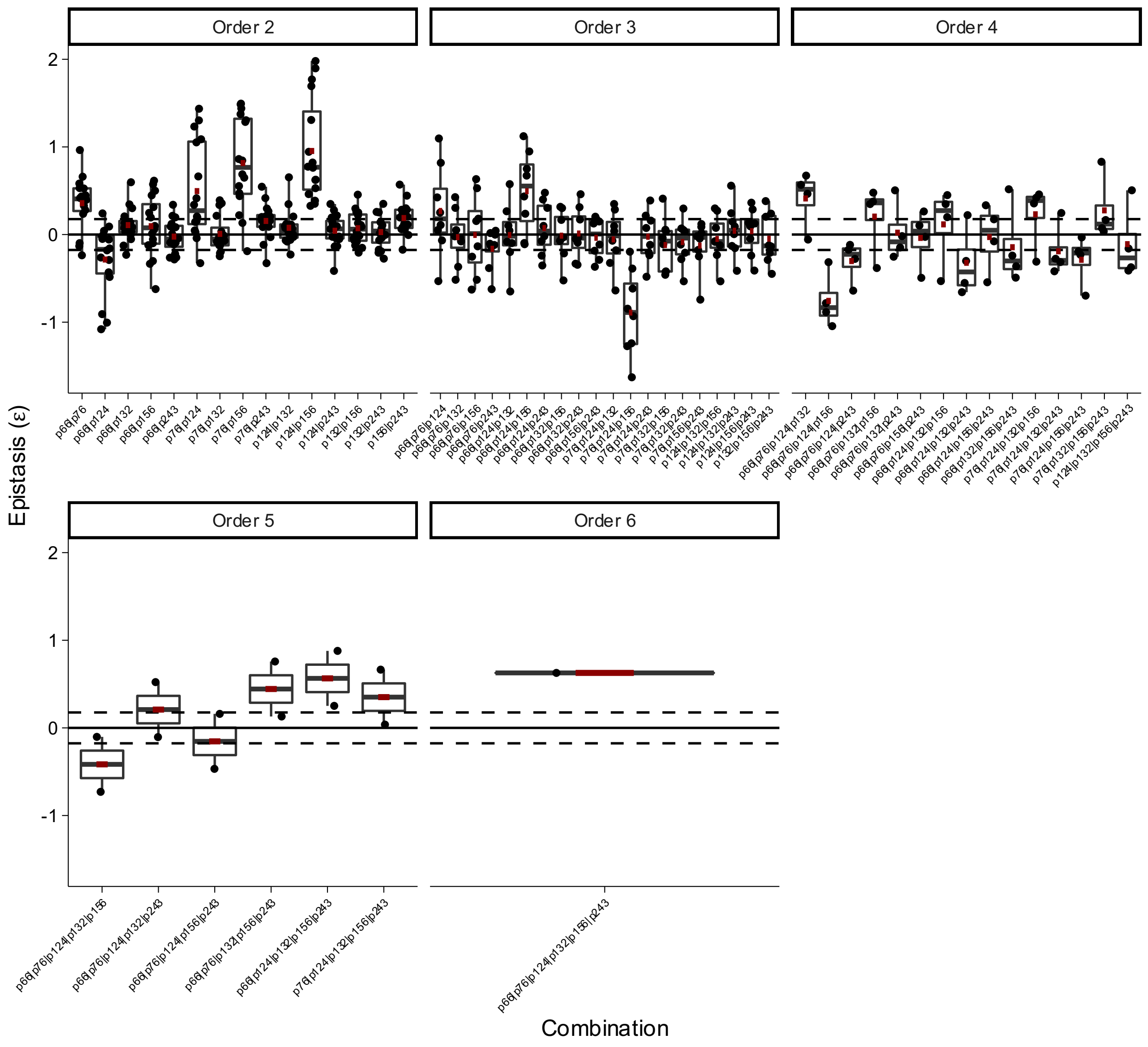


**Box Plot 48 | Epistasis of combinatorial mutations in beta lactamase OXA-48 for piperacillin hydrolysis in trajectory 2 from Fröhlich *et al.***

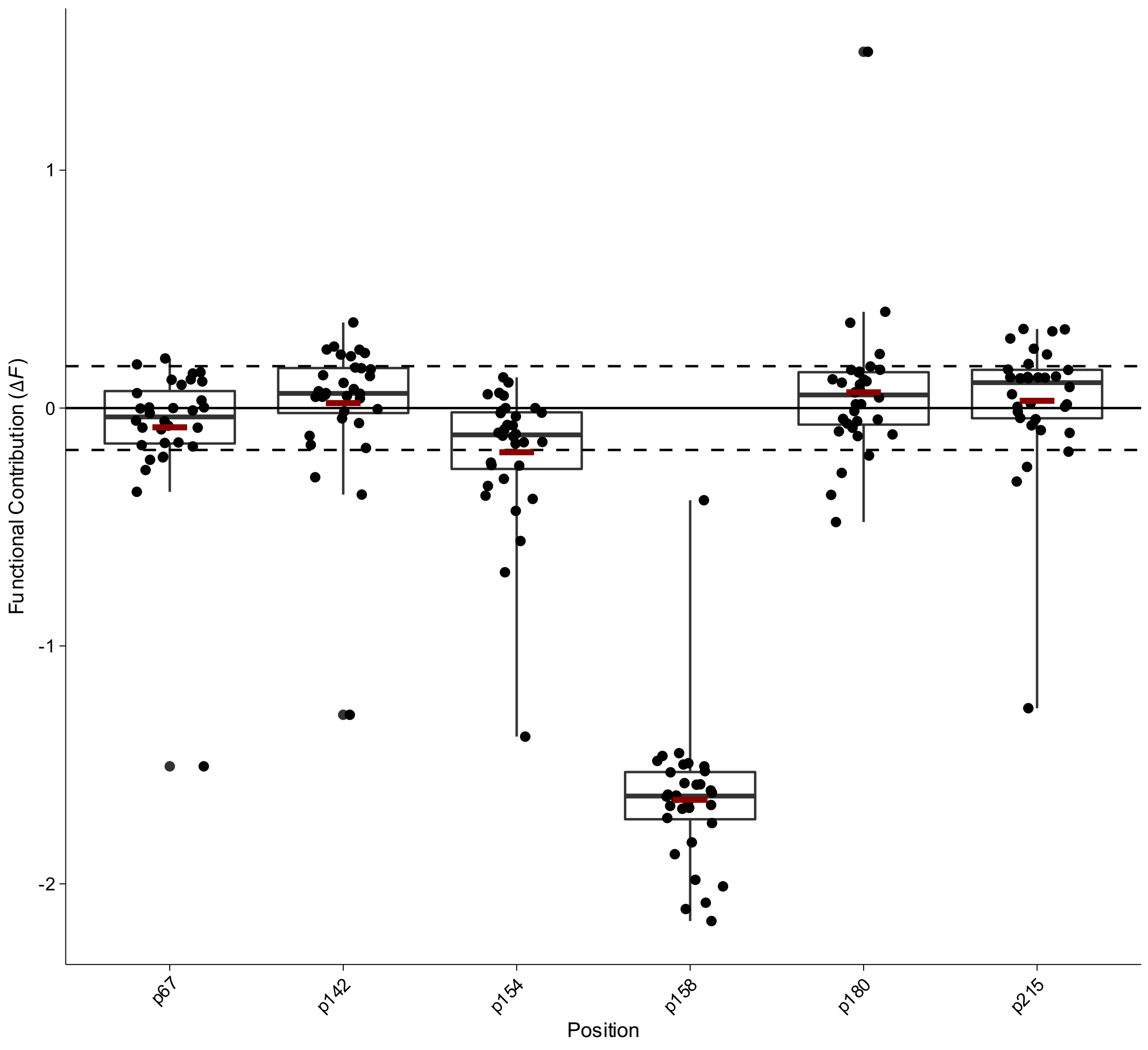


**Box Plot 49 | Functional contribution of positional mutations in beta lactamase OXA-48 for piperacillin hydrolysis in trajectory 3 from Fröhlich *et al.***

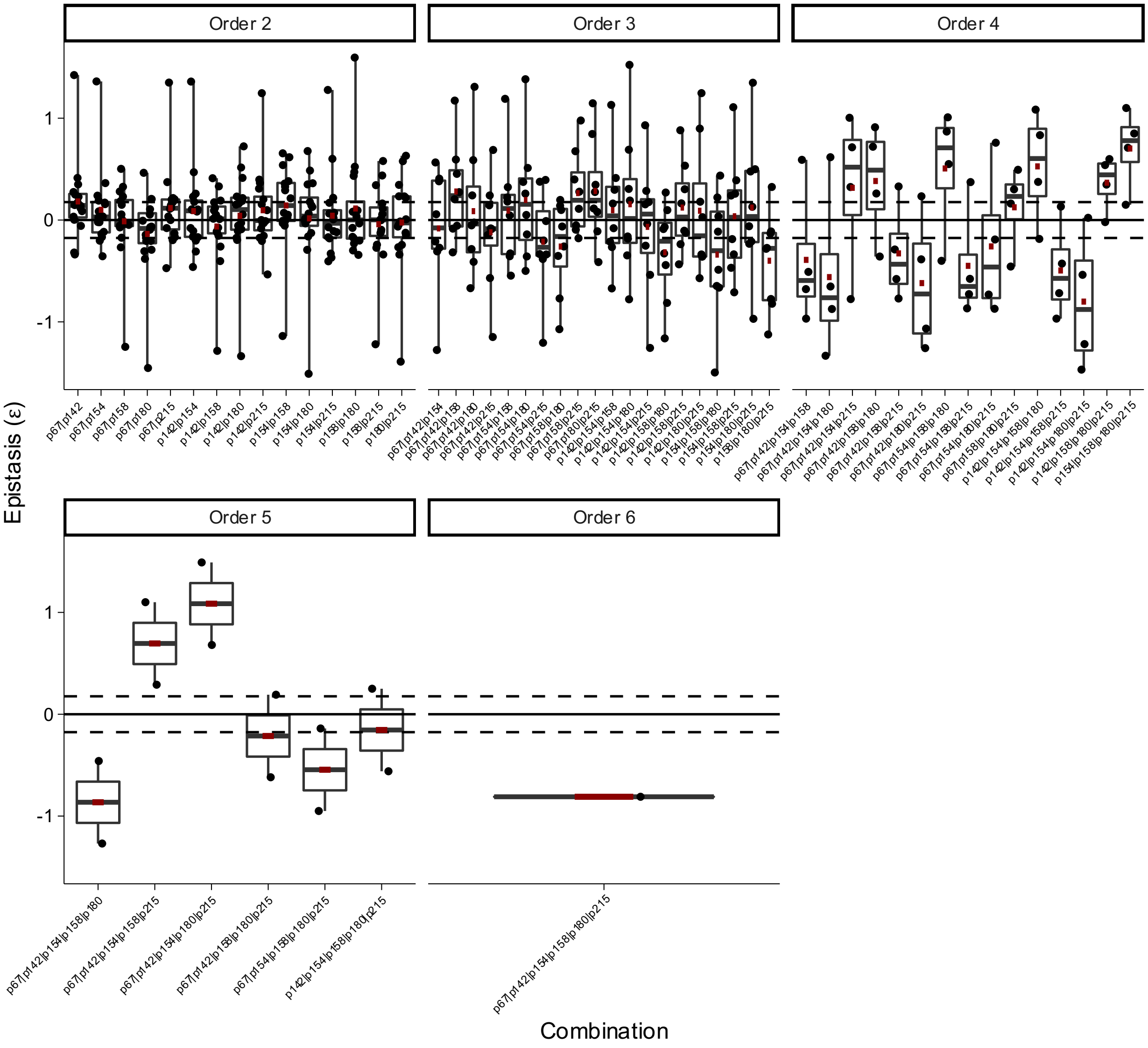


**Box Plot 50 | Epistasis of combinatorial mutations in beta lactamase OXA-48 for piperacillin hydrolysis in trajectory 3 from Fröhlich *et al.***

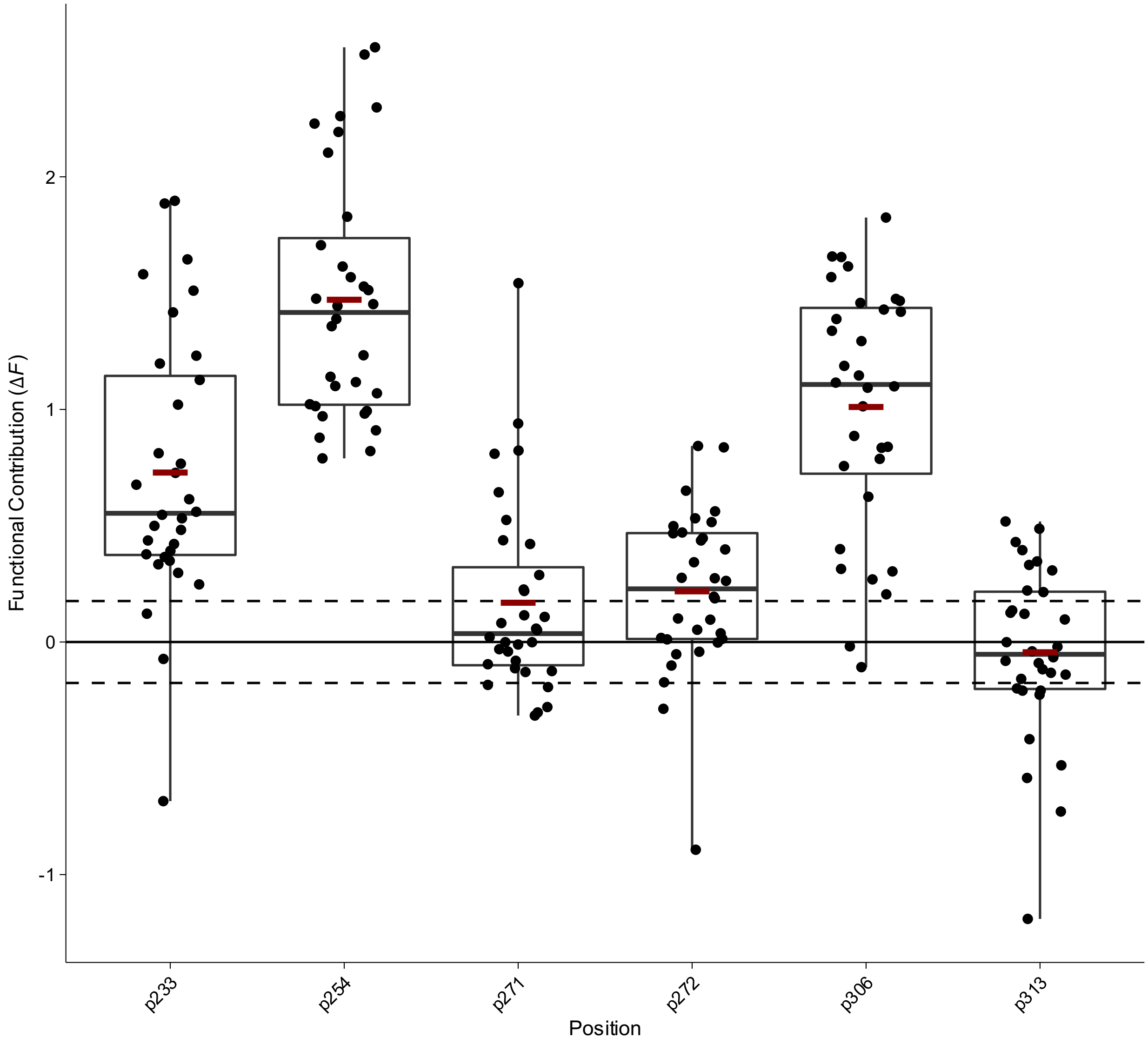


**Box Plot 53 | Functional contribution of positional mutations in phosphotriesterase (PTE) for 2-napthyl hexanoate (2NH) hydrolysis from this publication**

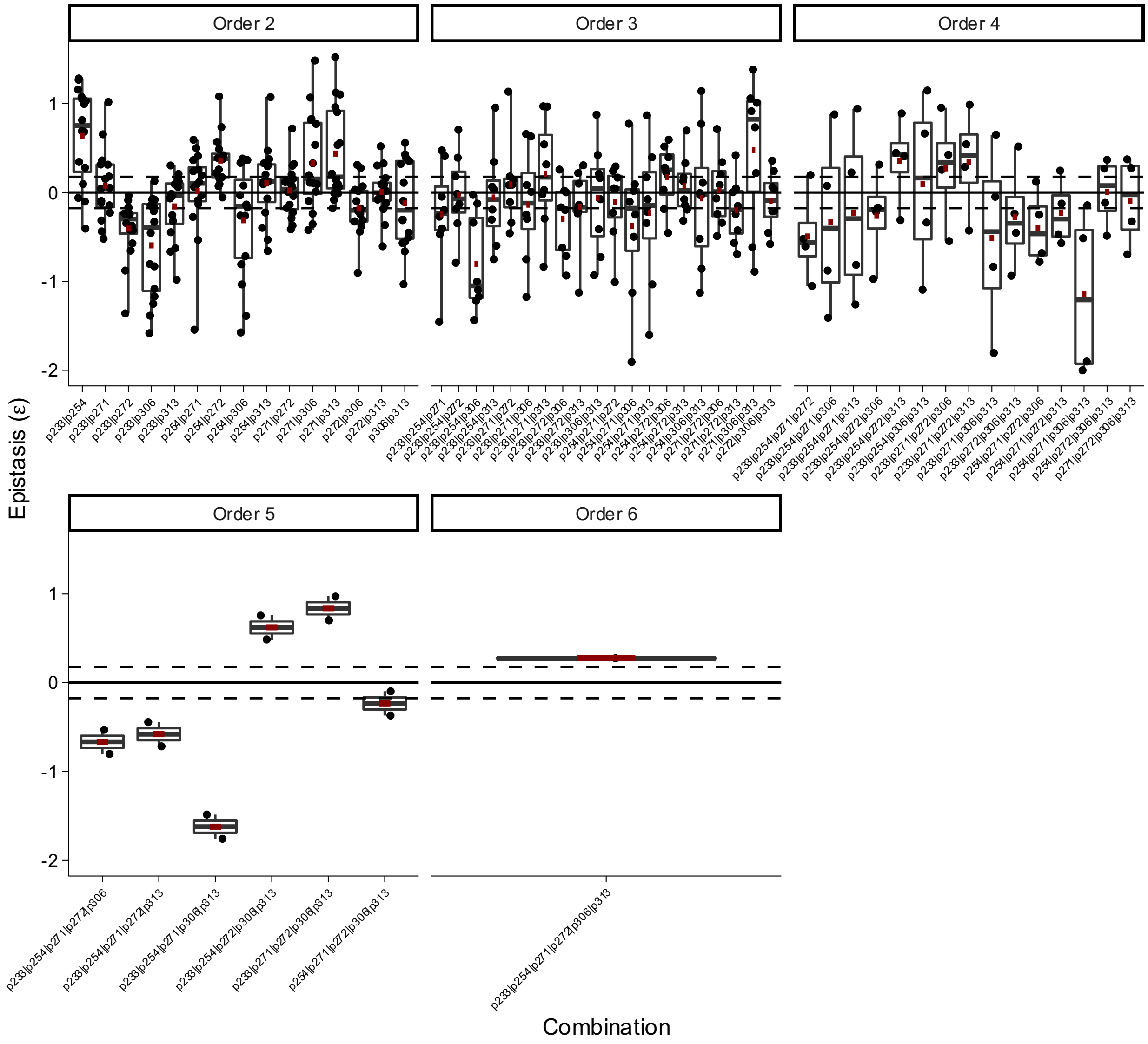


**Box Plot 52 | Epistasis of combinatorial mutations in phosphotriesterase (PTE) for 2-napthyl hexanoate (2NH) hydrolysis from this publication**

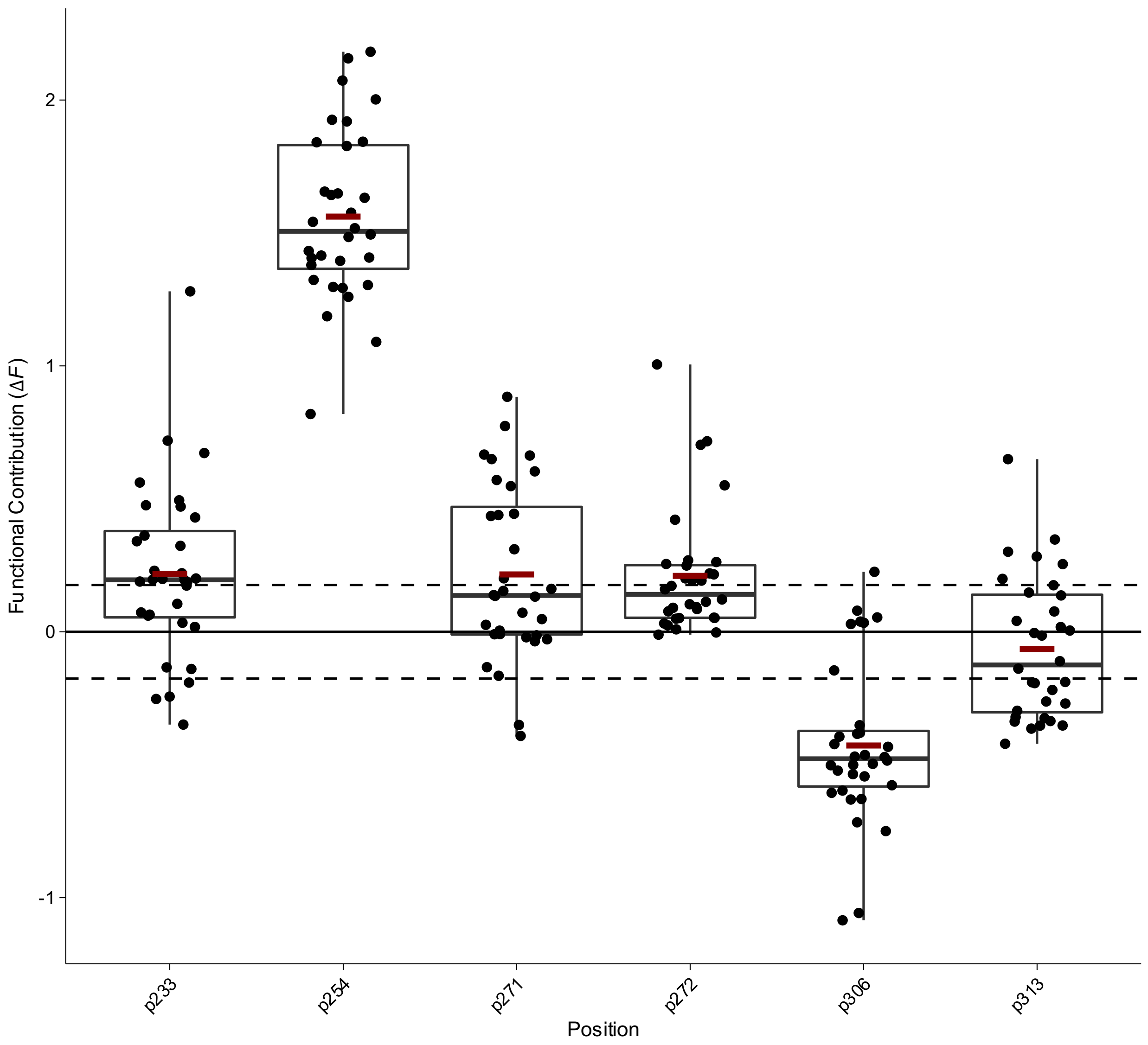


**Box Plot 53 | Functional contribution of positional mutations in phosphotriesterase (PTE) for butyrate hydrolysis from Miton *et al.***

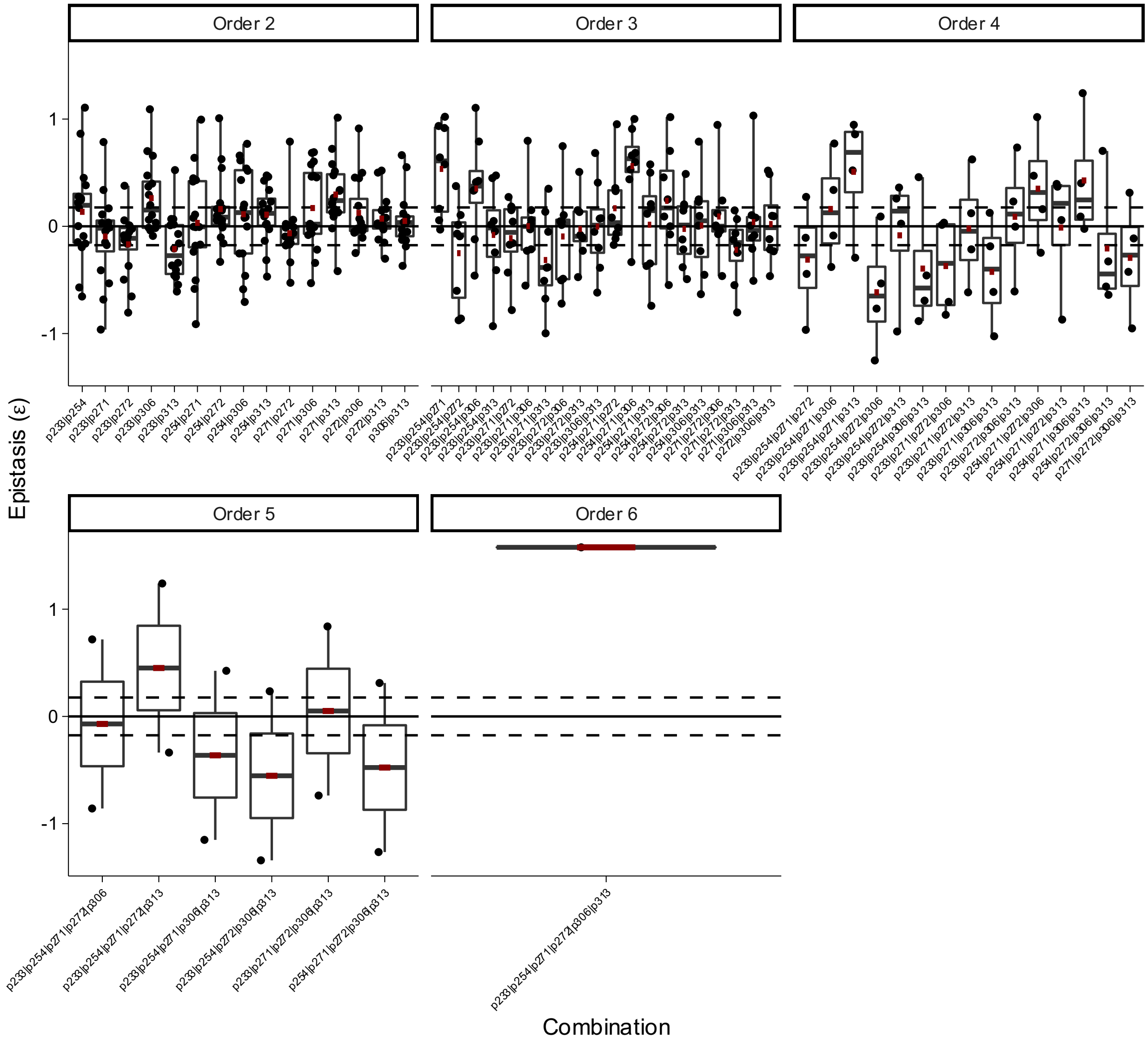


**Box Plot 54 | Epistasis of combinatorial mutations in phosphotriesterase (PTE) for butyrate hydrolysis from Miton *et al.***

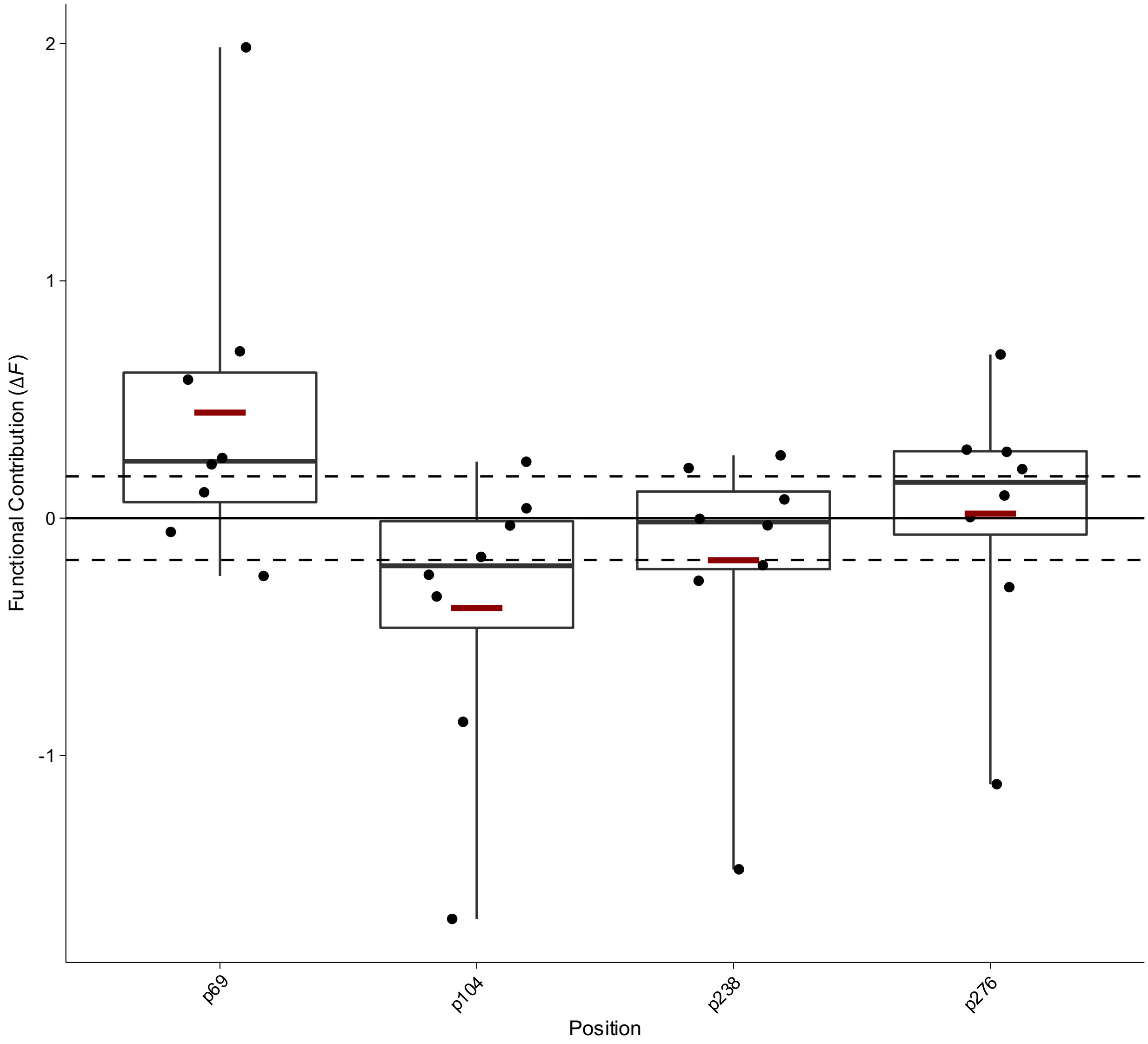


**Box Plot 55 | Functional contribution of positional mutations in of beta lactamase TEM in AM antibiotic from Mira *et al.***

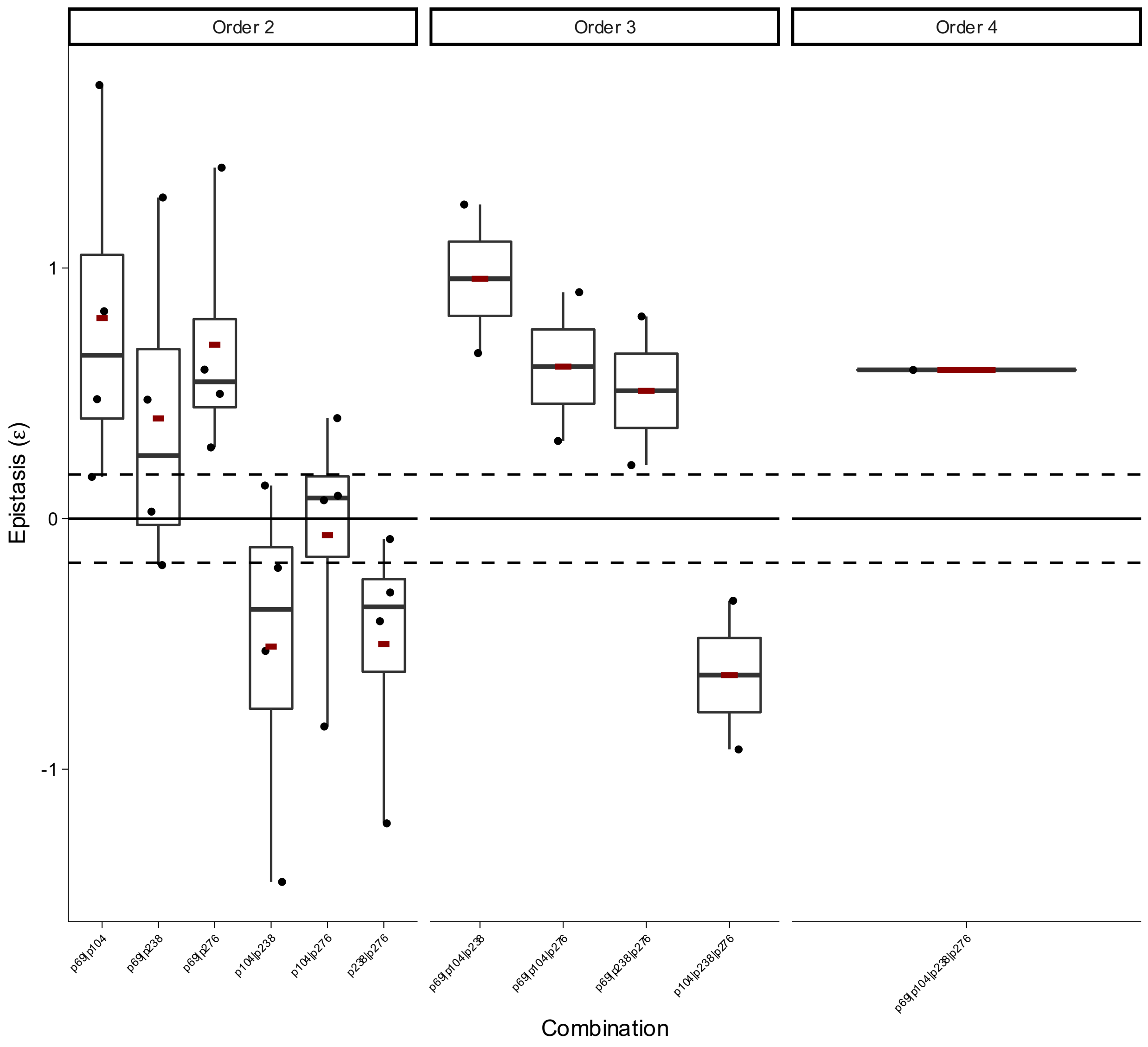


**Box Plot 56 | Epistasis of combinatorial mutations in beta lactamase TEM in AM antibiotic Mira *et al.***

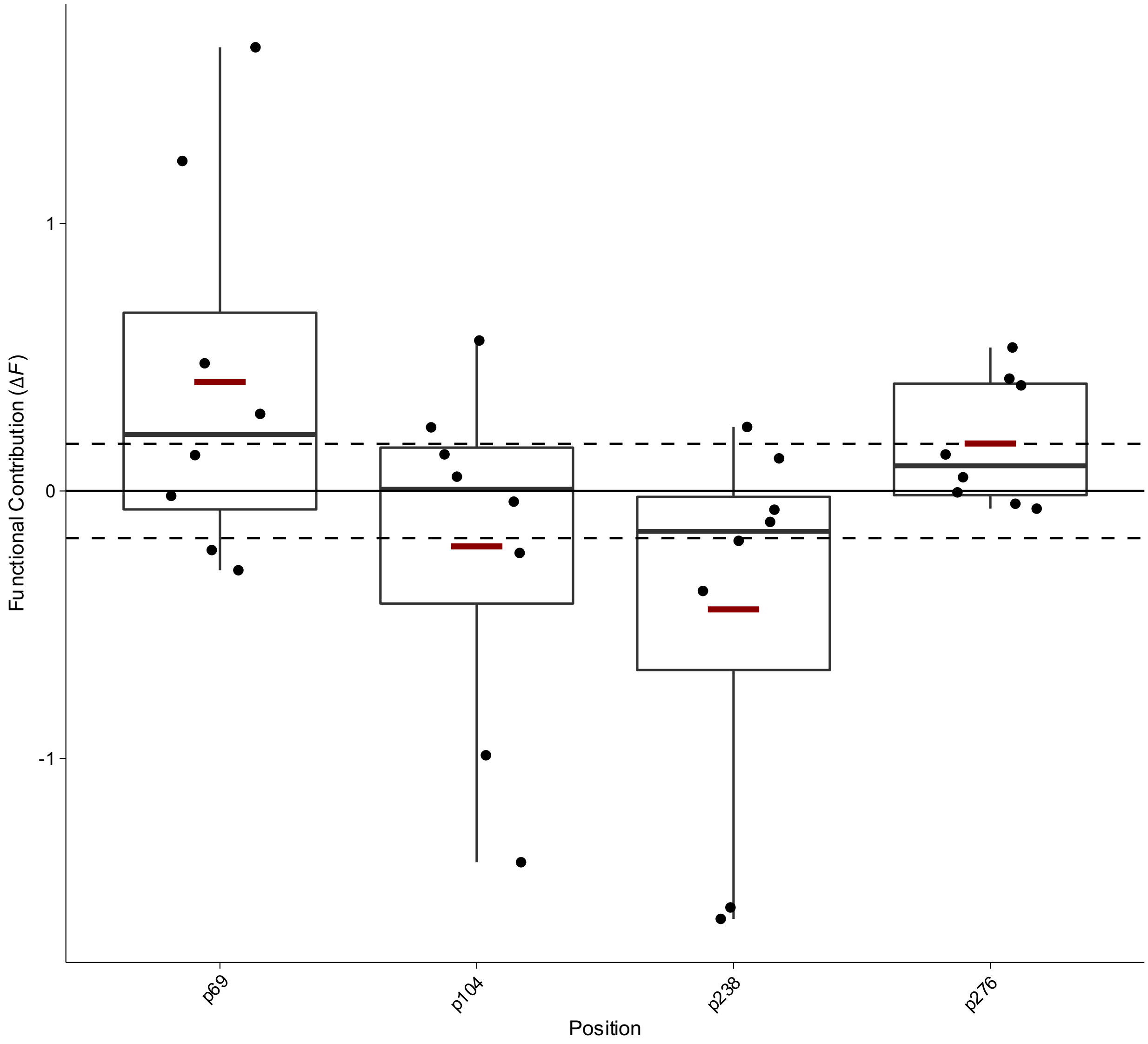


**Box Plot 57 | Functional contribution of positional mutations in of beta lactamase TEM in AMC antibiotic Mira *et al.***

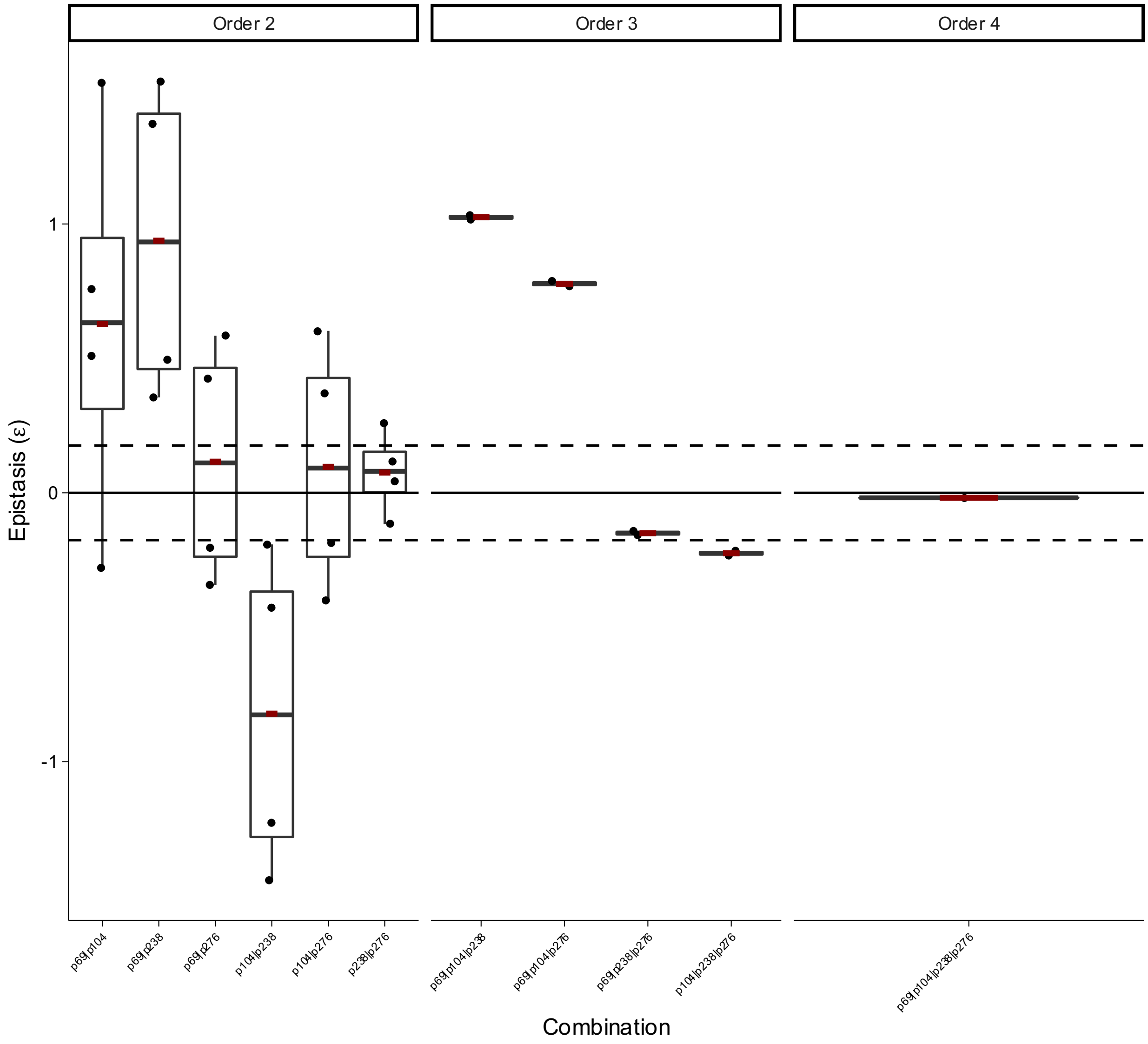


**Box Plot 58 | Epistasis of combinatorial mutations in beta lactamase TEM in AMC antibiotic Mira *et al.***

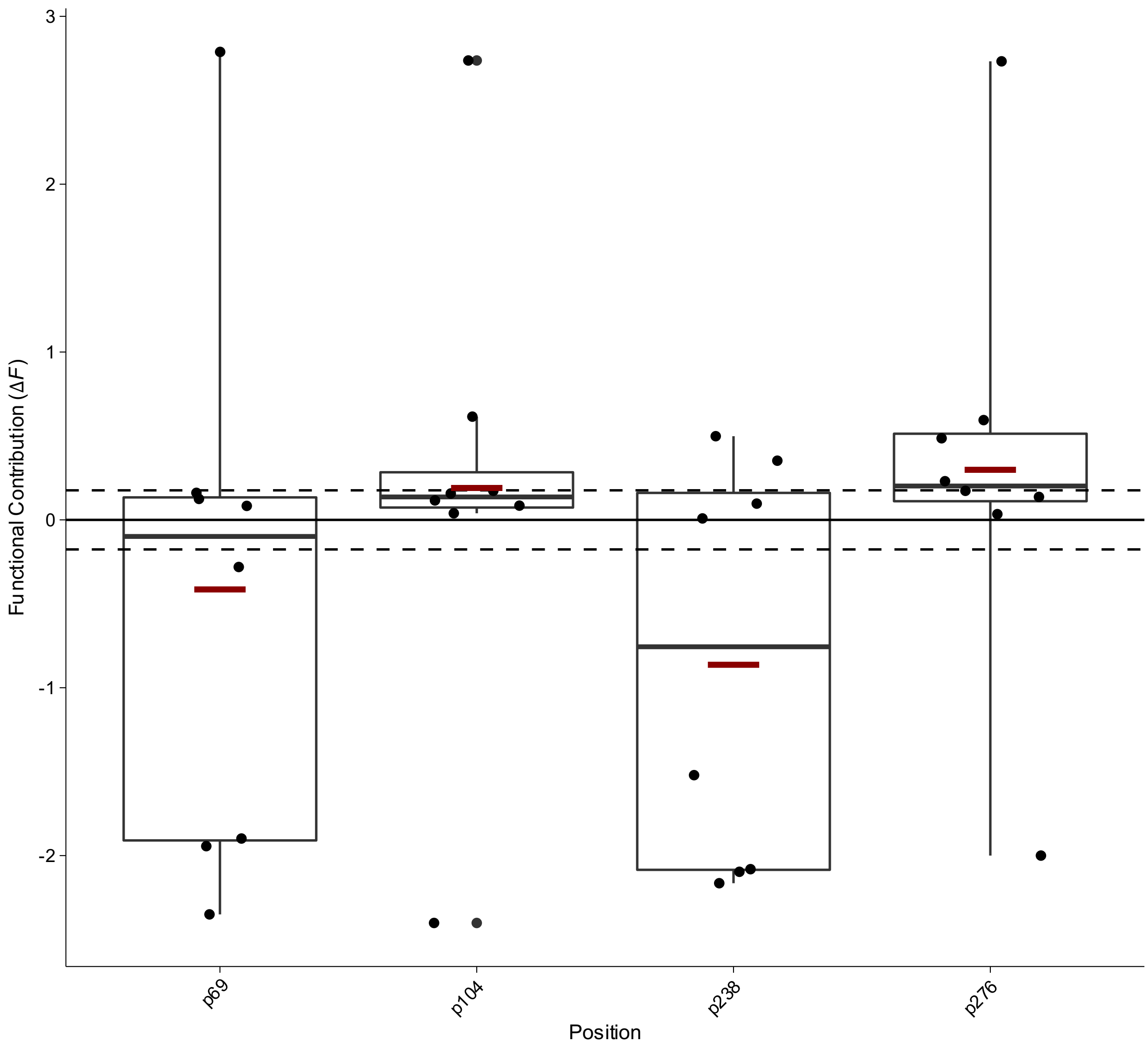


**Box Plot 59 | Functional contribution of positional mutations in of beta lactamase TEM in AMP antibiotic Mira *et al.***

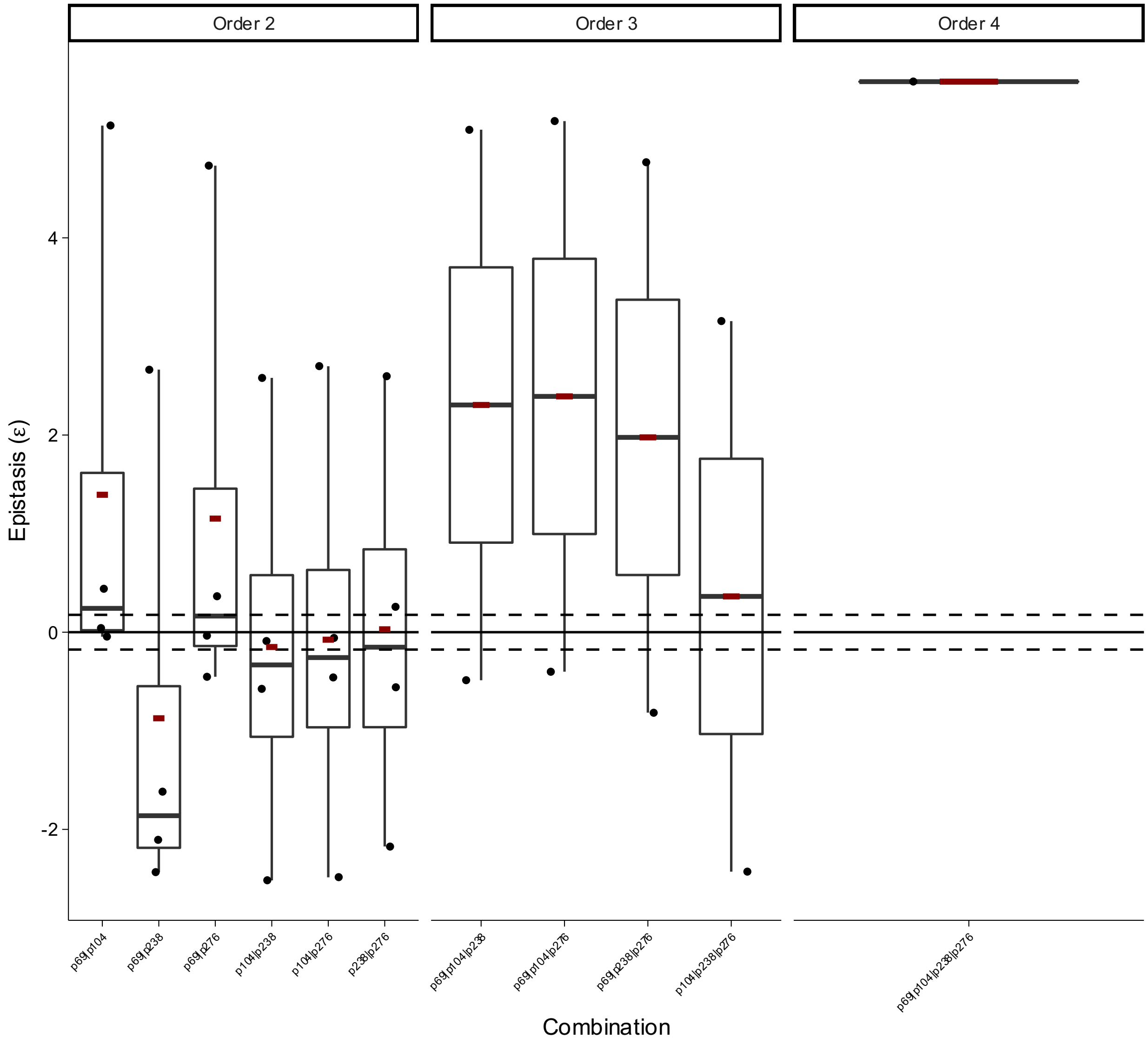


**Box Plot 60 | Epistasis of combinatorial mutations in beta lactamase TEM in AMP antibiotic Mira *et al.***

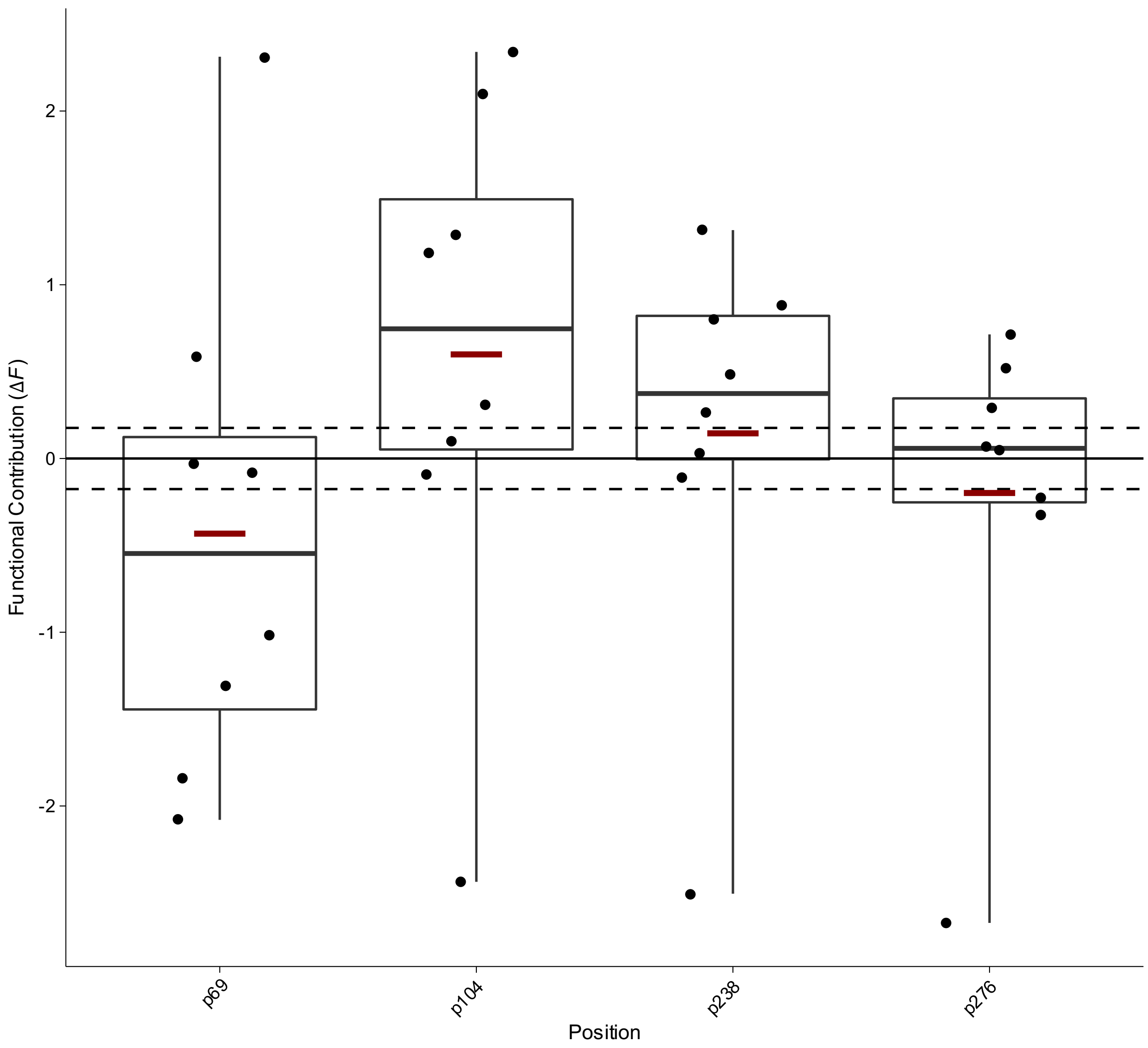


**Box Plot 61 | Functional contribution of positional mutations in of beta lactamase TEM in CAZ antibiotic Mira *et al.***

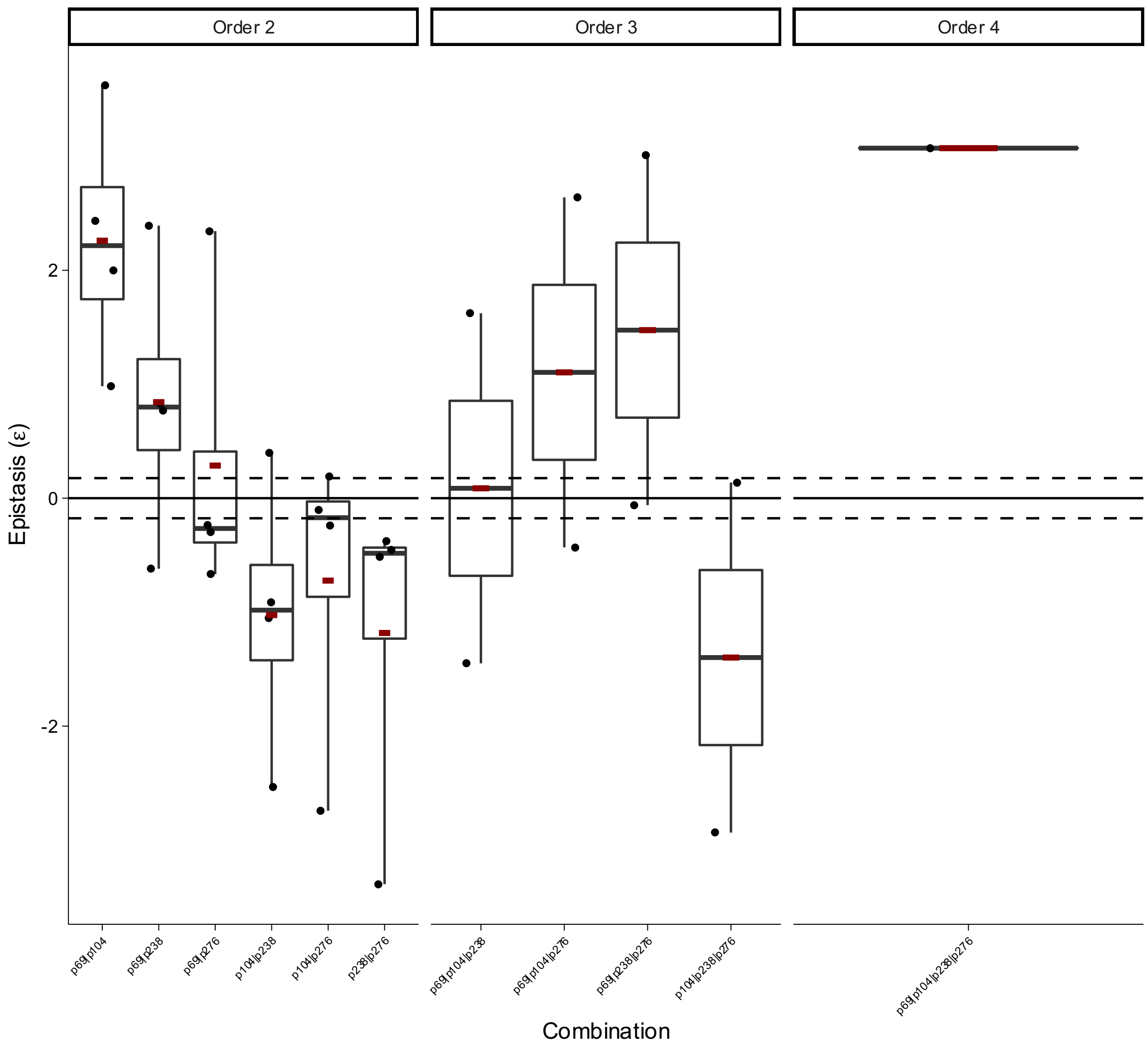


**Box Plot 62 | Epistasis of combinatorial mutations in beta lactamase TEM in CAZ antibiotic Mira *et al.***

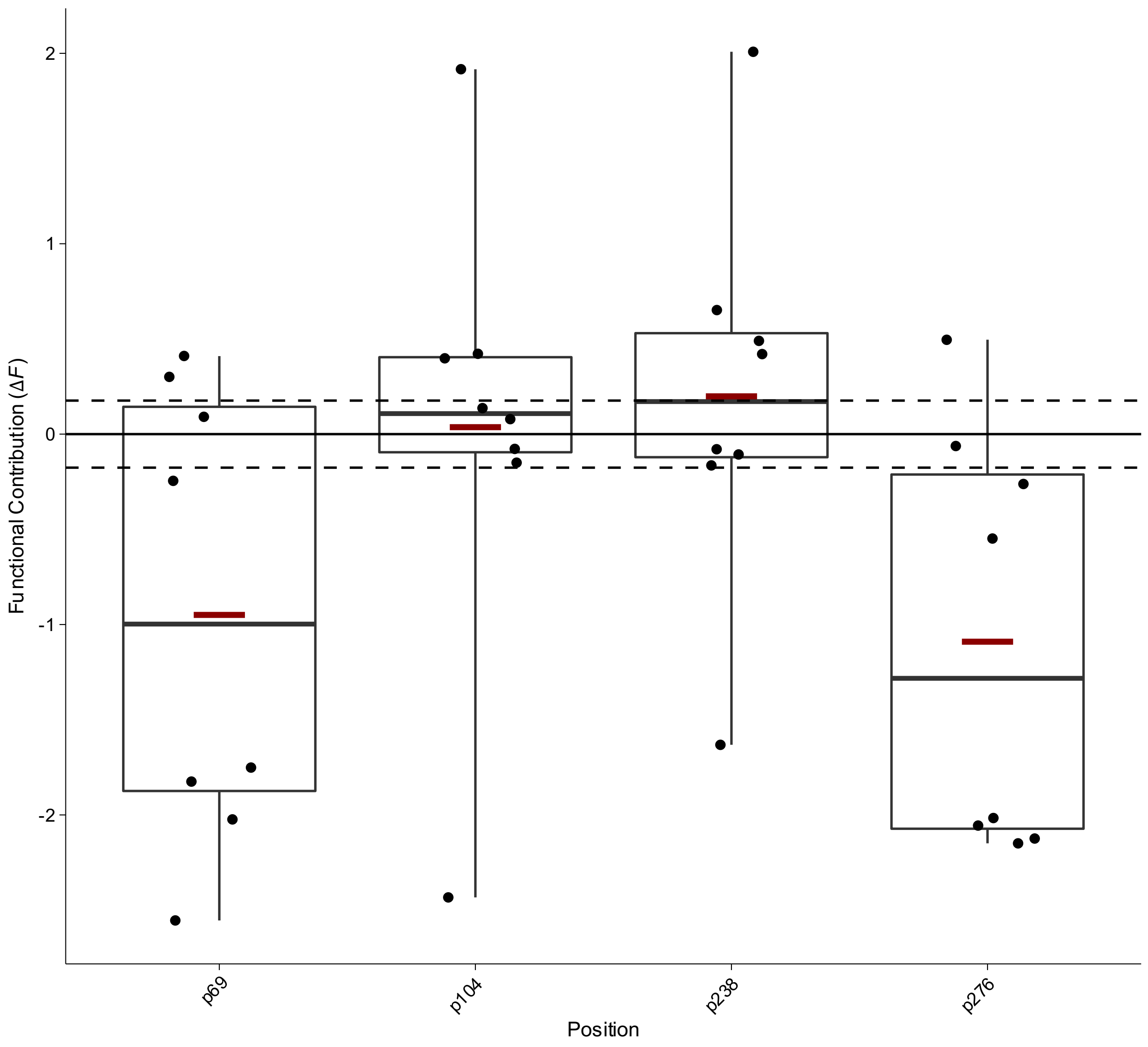


**Box Plot 63 | Functional contribution of positional mutations in of beta lactamase TEM in CEC antibiotic Mira *et al.***

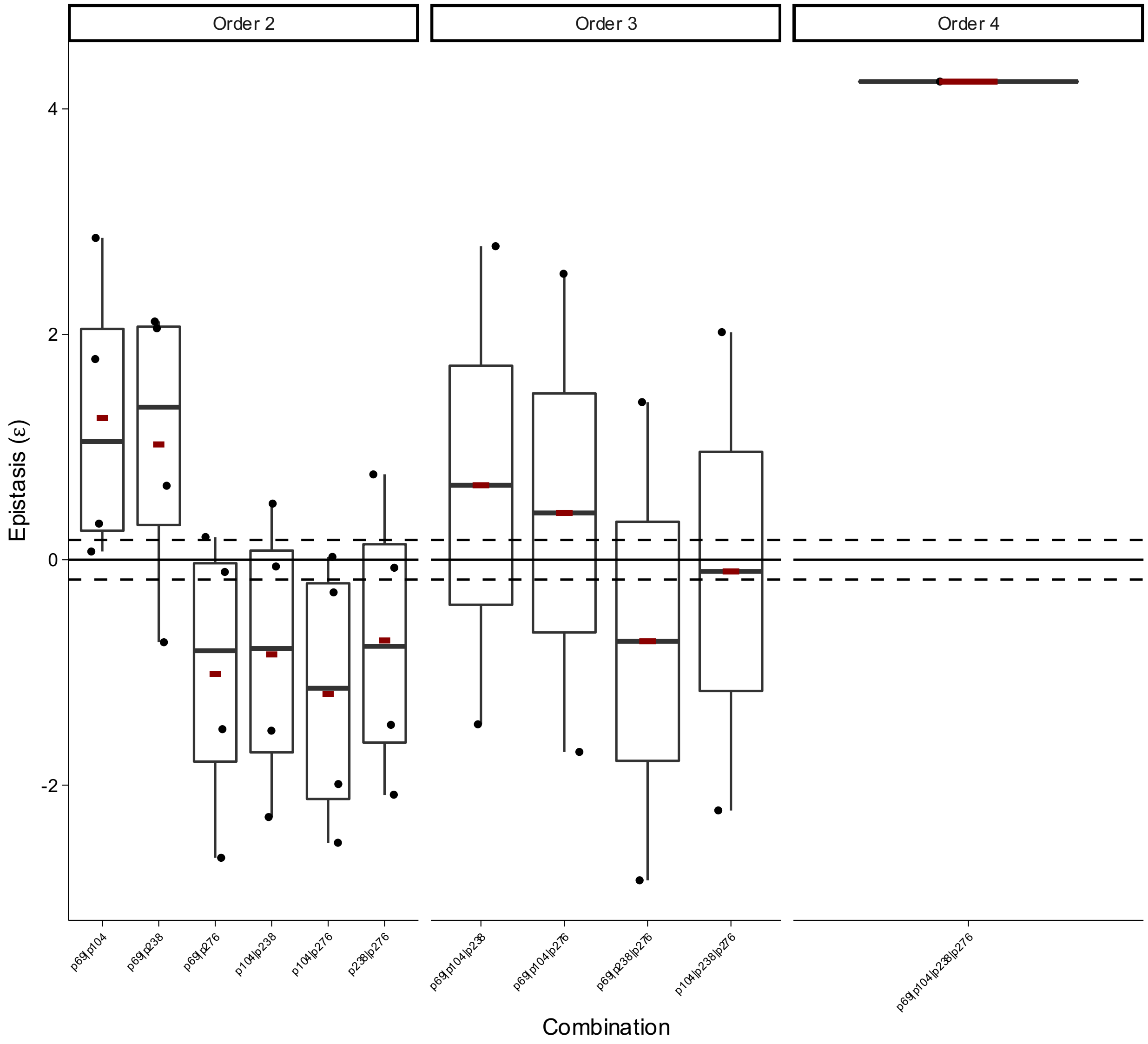


**Box Plot 64 | Epistasis of combinatorial mutations in beta lactamase TEM in CEC antibiotic Mira *et al.***

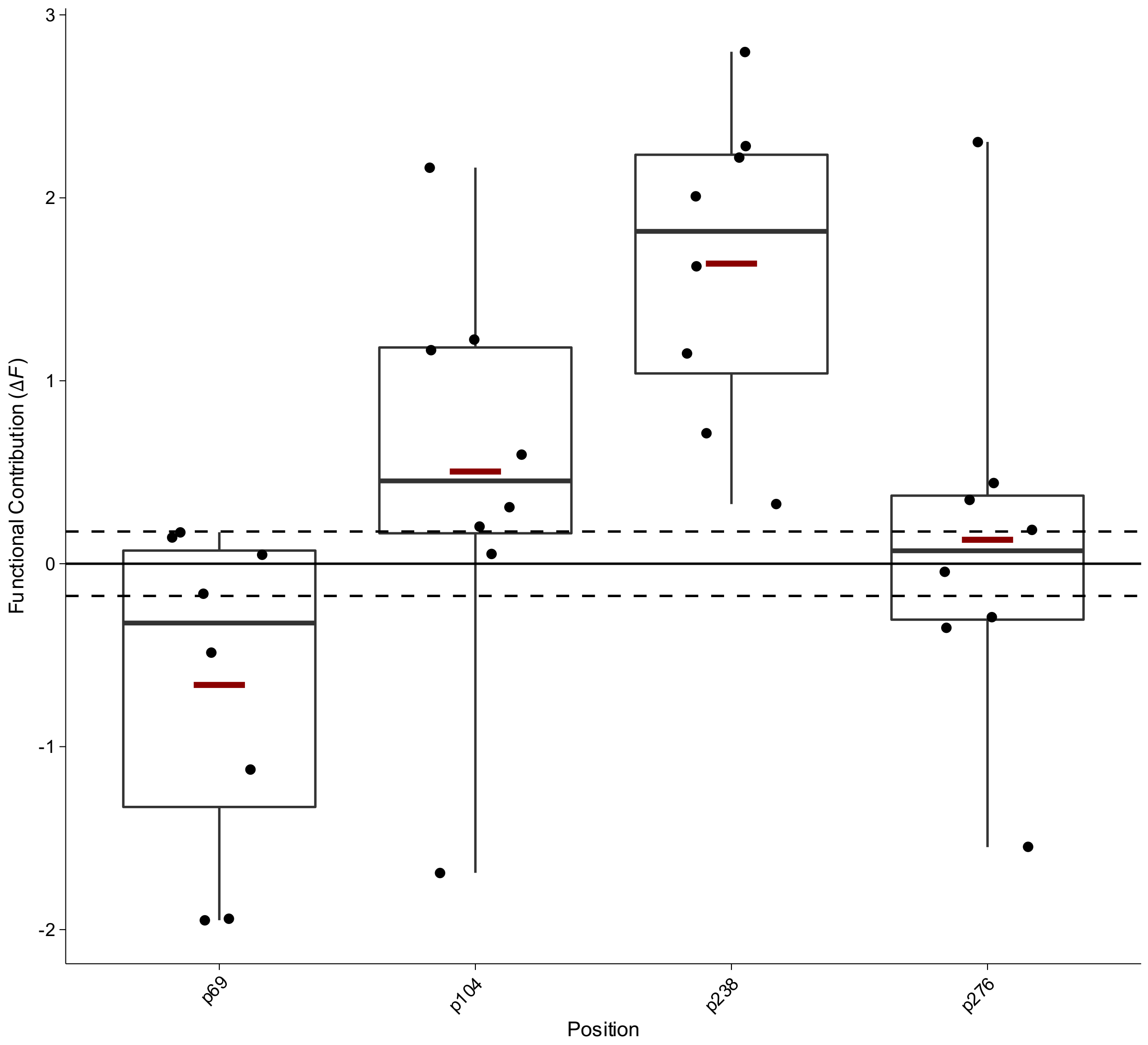


**Box Plot 65 | Functional contribution of positional mutations in of beta lactamase TEM in CPD antibiotic Mira *et al.***

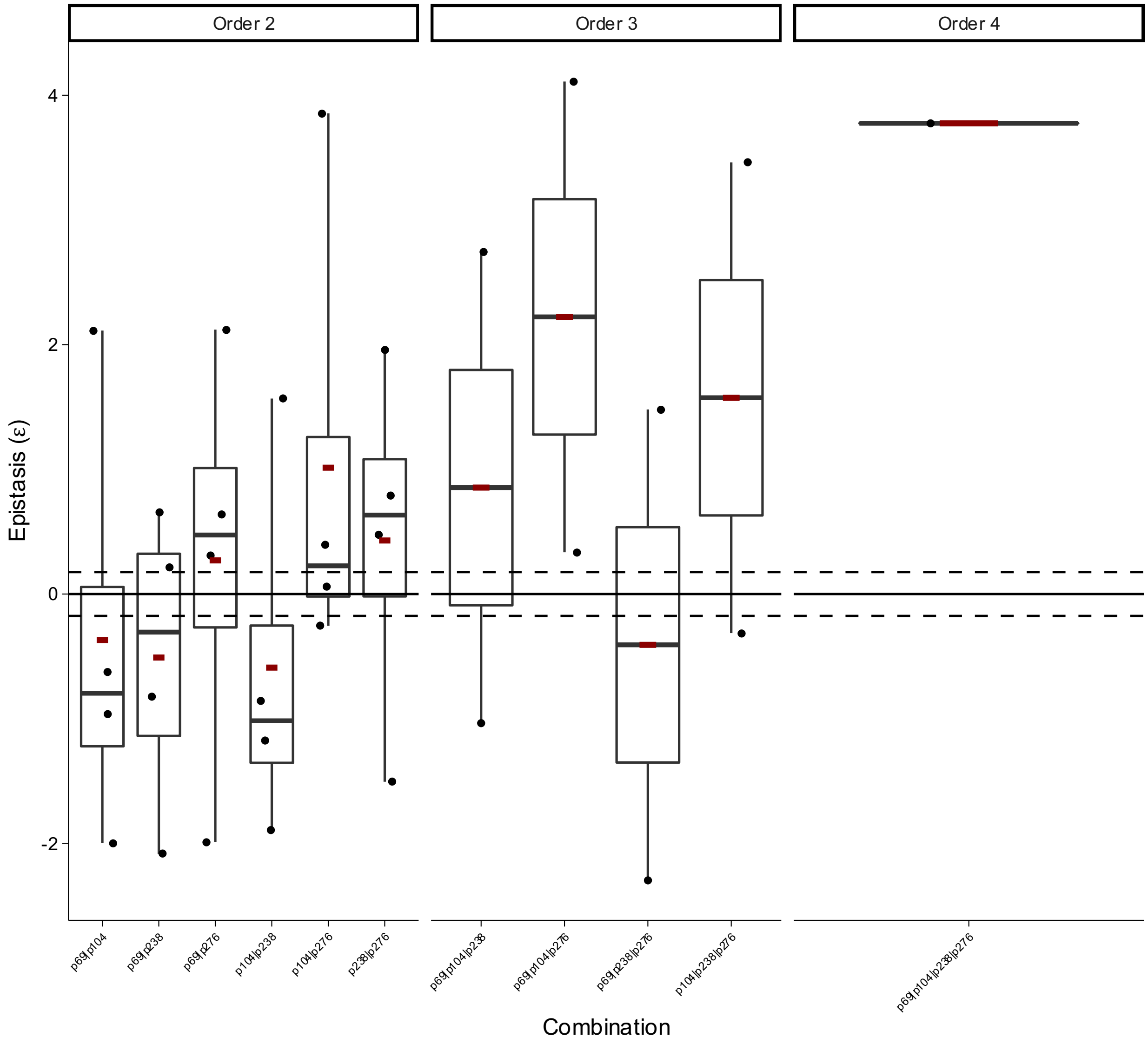


**Box Plot 66 | Epistasis of combinatorial mutations in beta lactamase TEM in CPD antibiotic Mira *et al.***

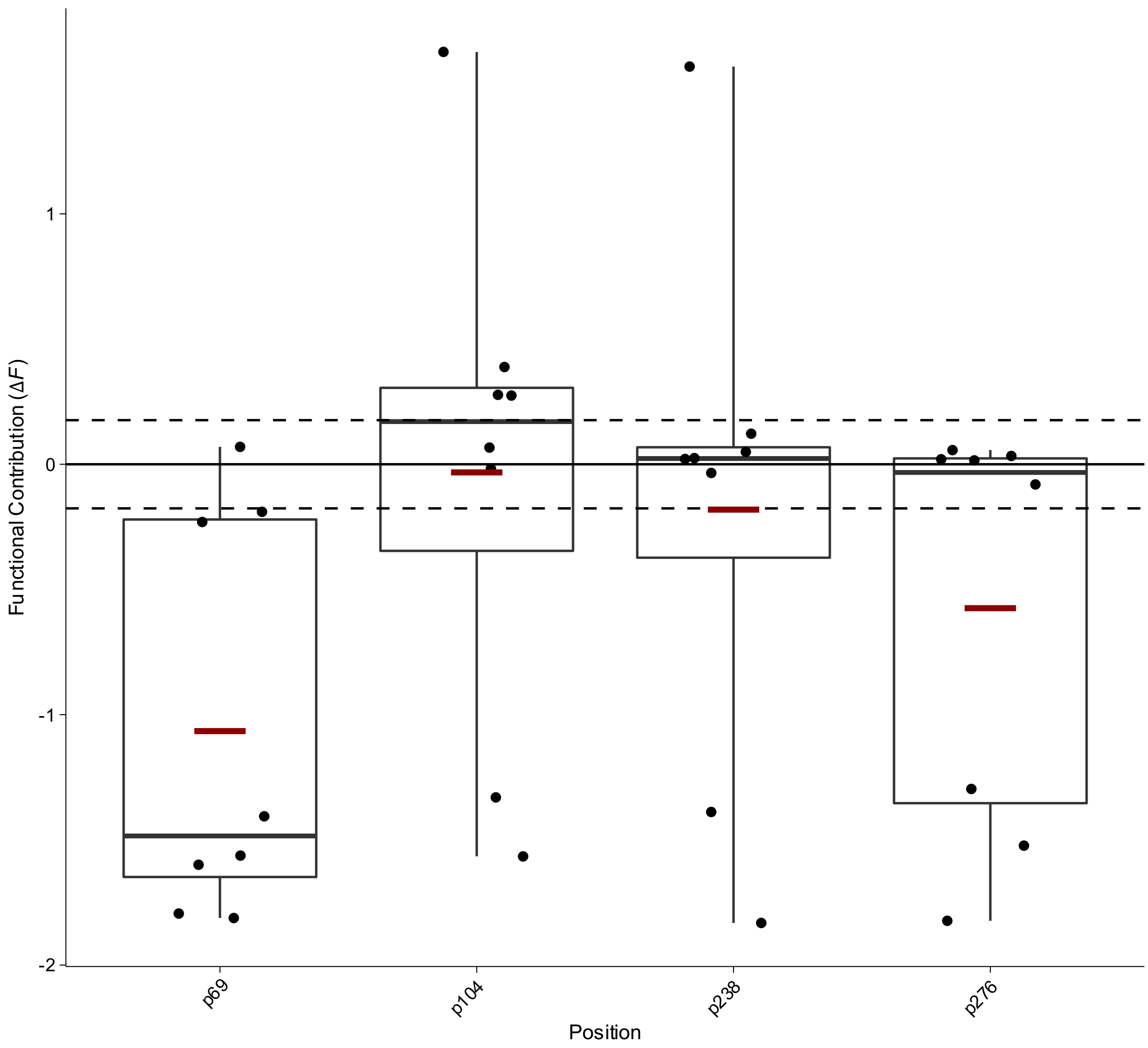


**Box Plot 67 | Functional contribution of positional mutations in of beta lactamase TEM in CPR antibiotic Mira *et al.***

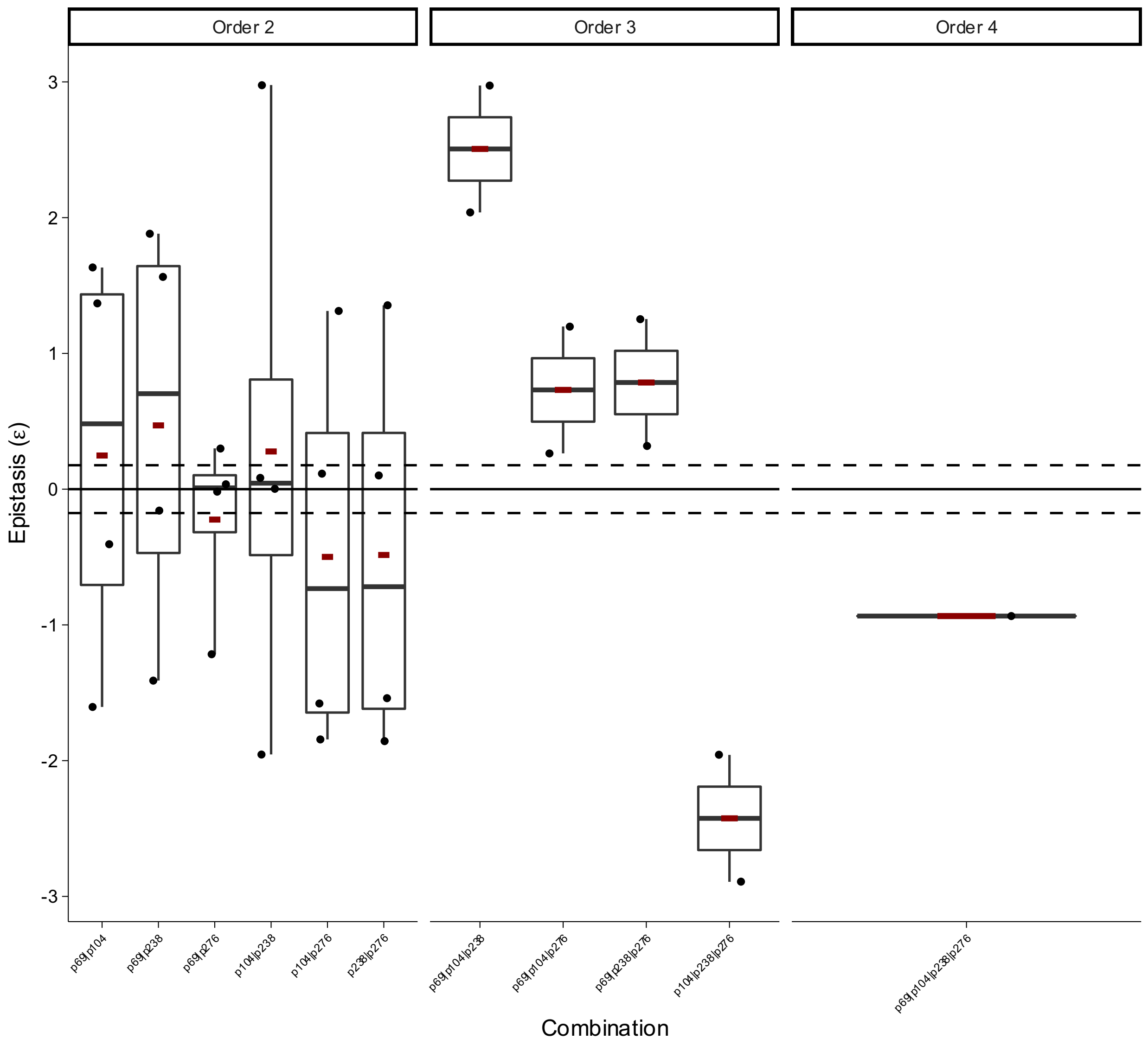


**Box Plot 68 | Epistasis of combinatorial mutations in beta lactamase TEM in CPR antibiotic Mira *et al.***

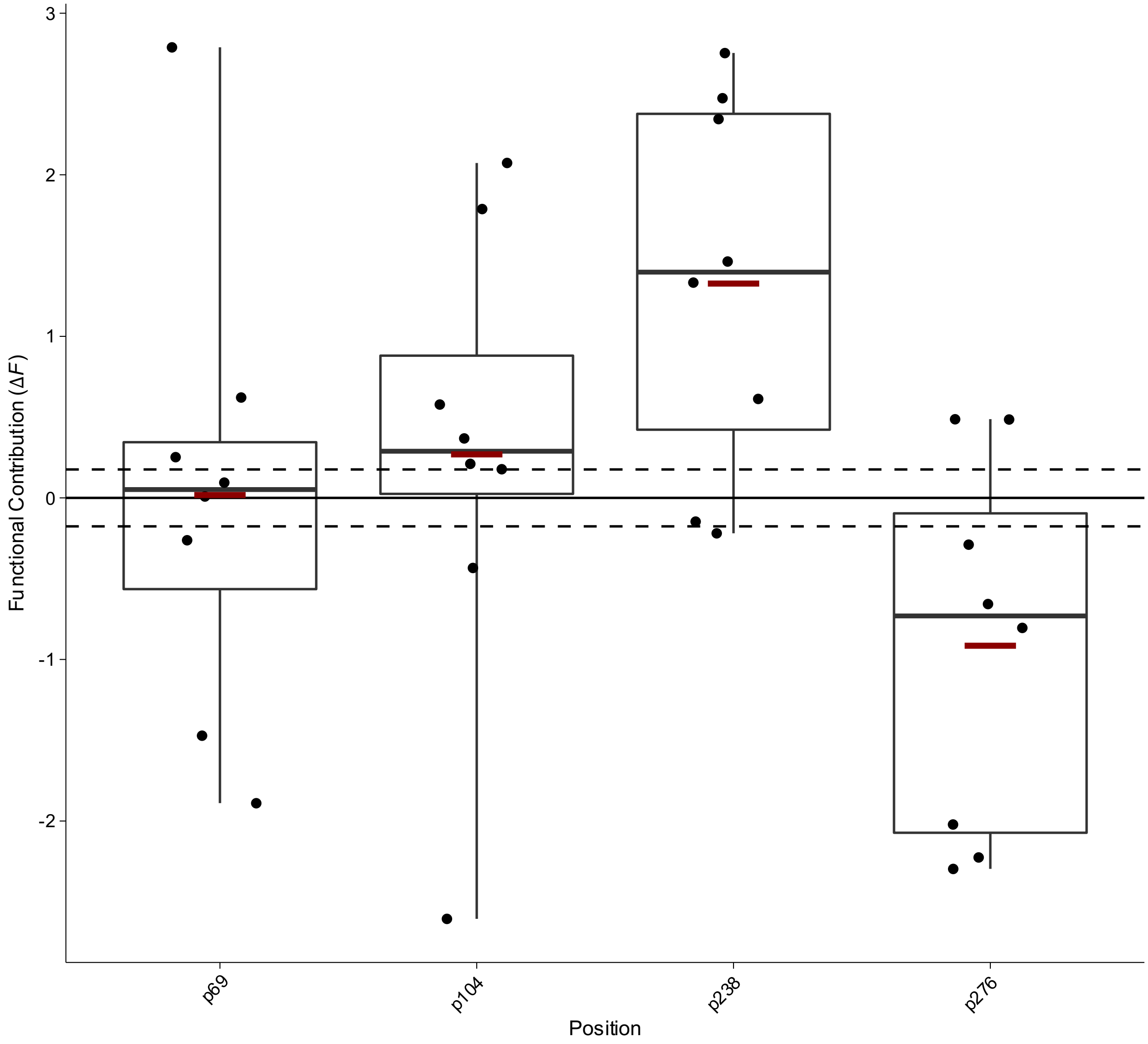


**Box Plot 69 | Functional contribution of positional mutations in of beta lactamase TEM in CRO antibiotic Mira *et al.***

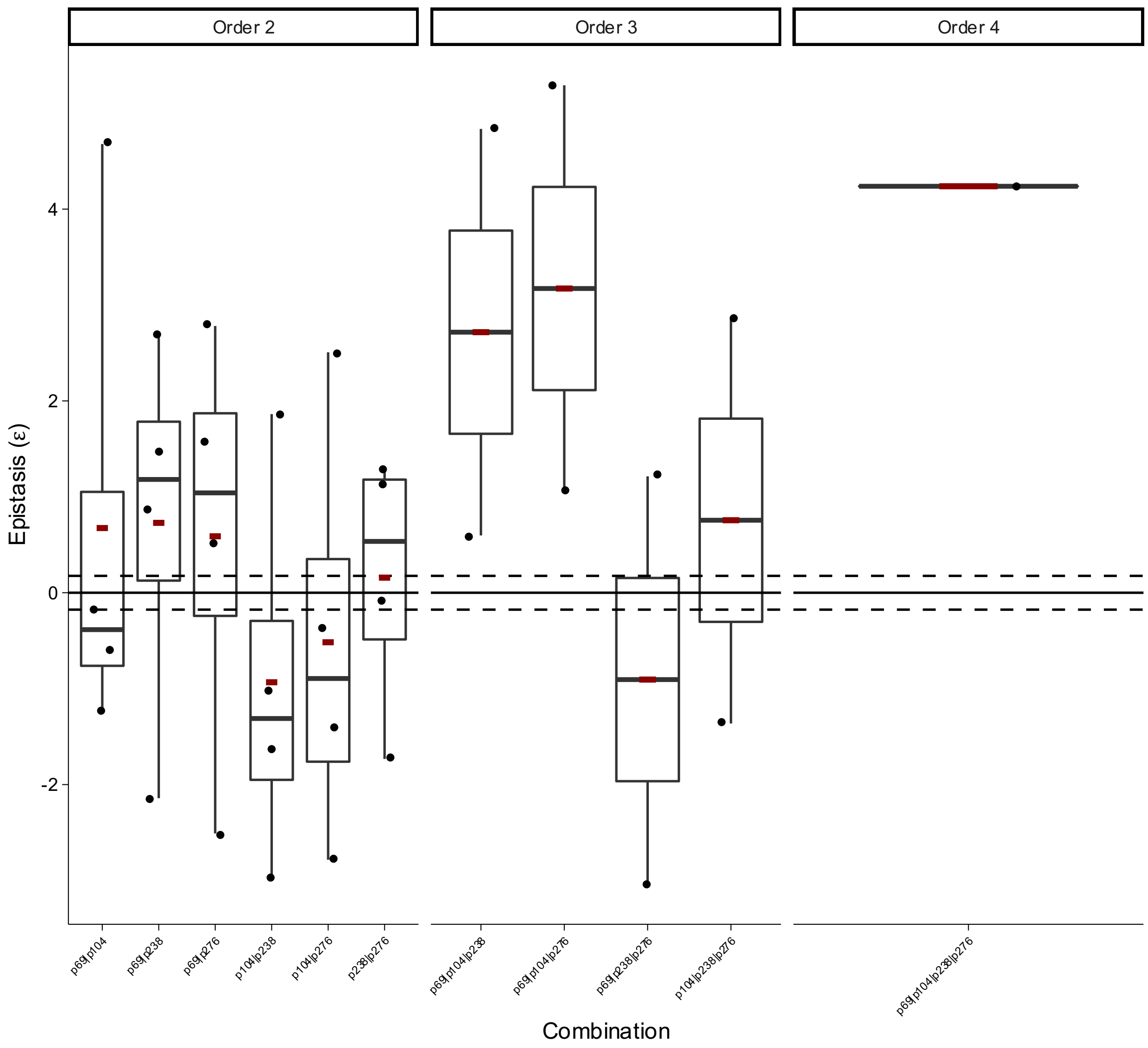


**Box Plot 70 | Epistasis of combinatorial mutations in beta lactamase TEM in CRO antibiotic Mira *et al.***

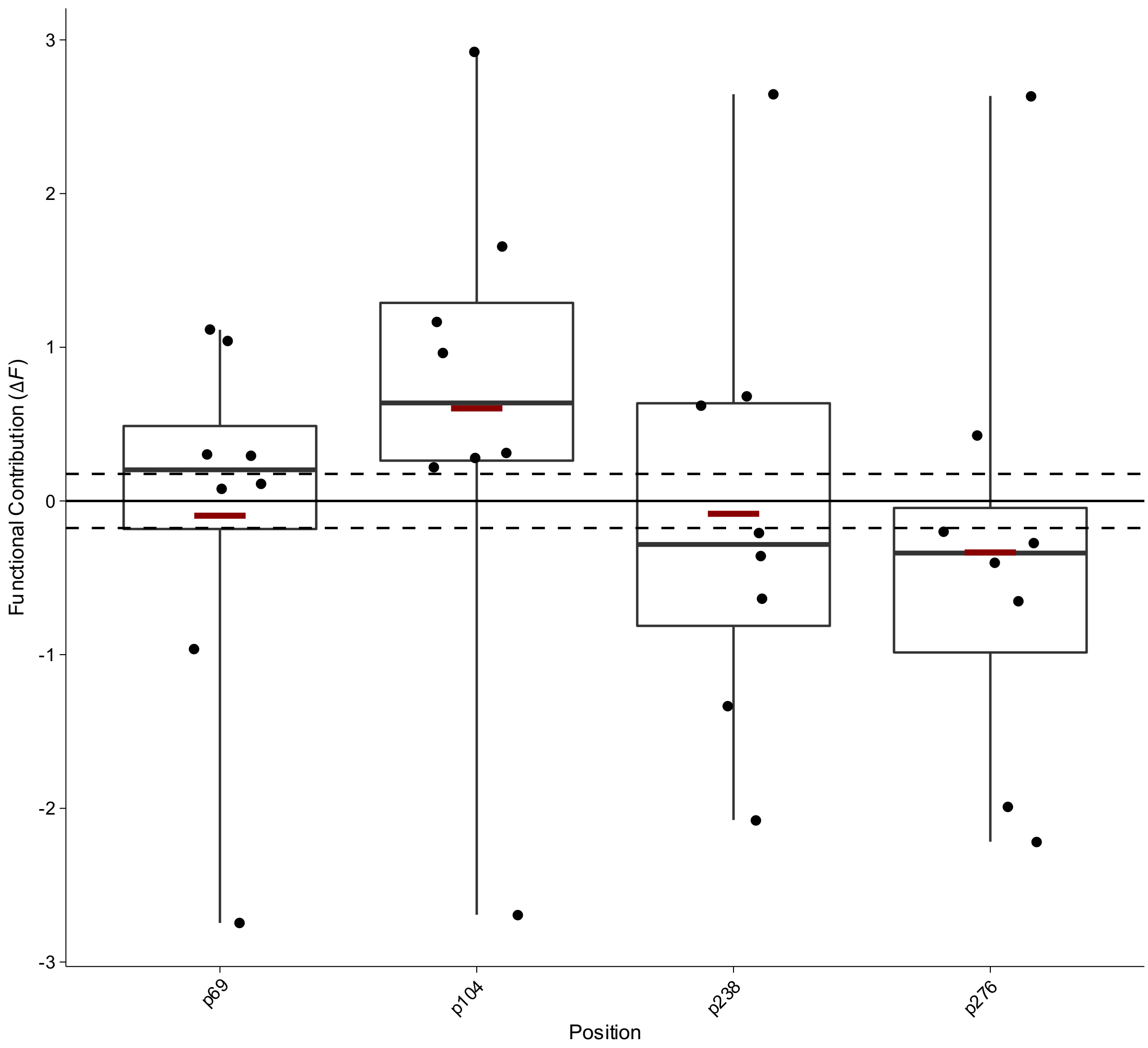


**Box Plot 71 | Functional contribution of positional mutations in of beta lactamase TEM in CTT antibiotic Mira *et al.***

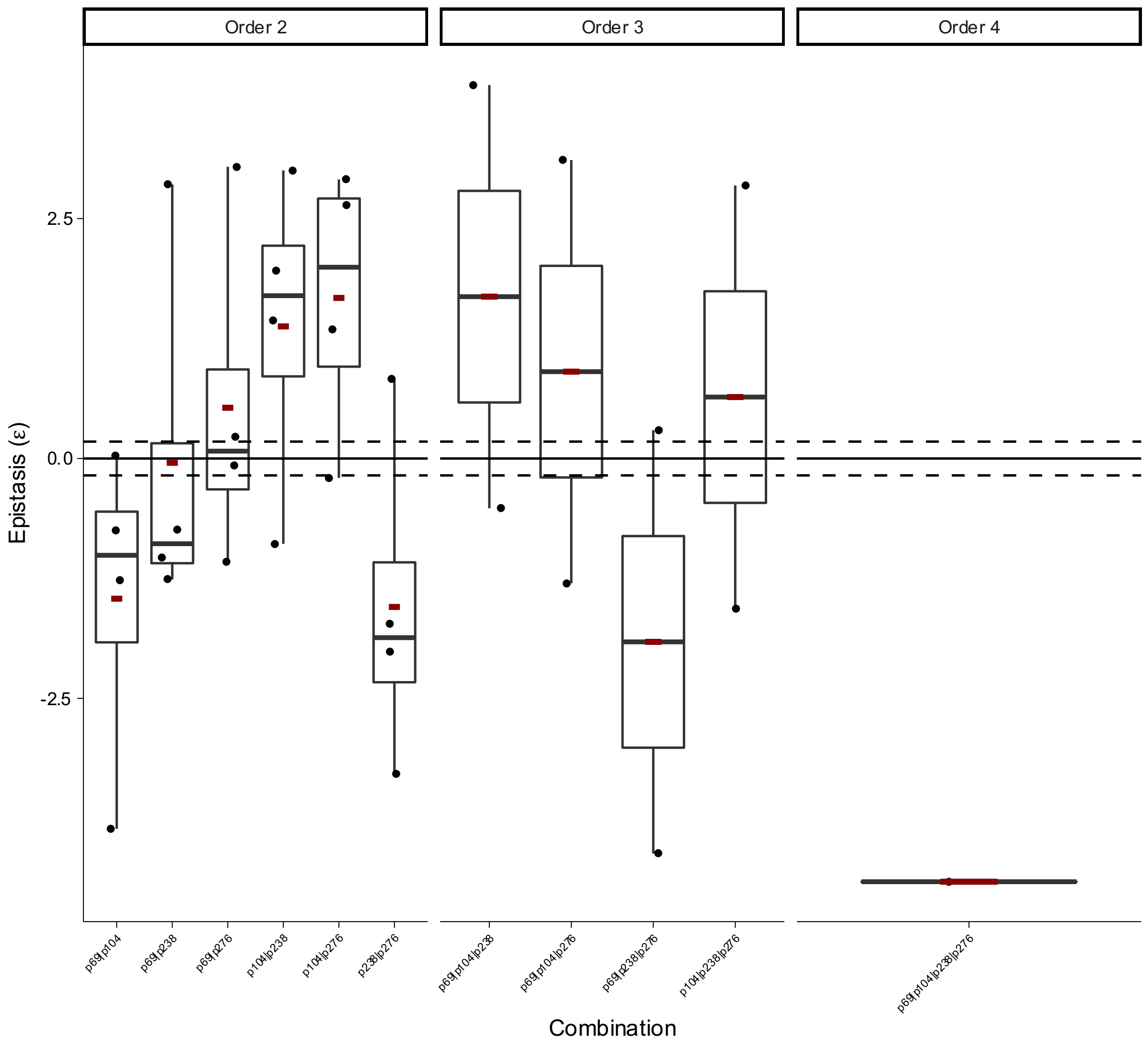


**Box Plot 72 | Epistasis of combinatorial mutations in beta lactamase TEM in CTT antibiotic Mira *et al.***

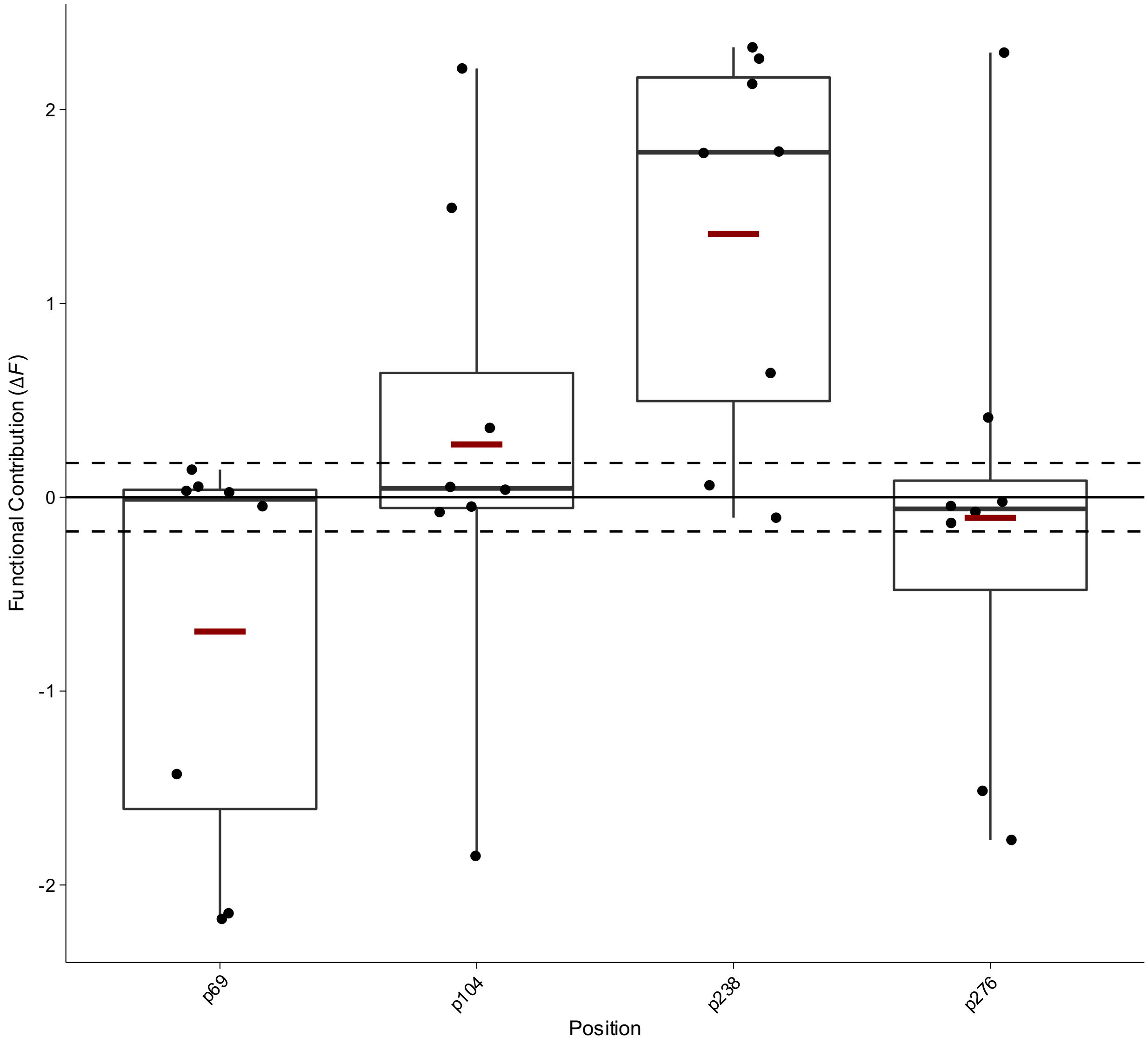


**Box Plot 73 | Functional contribution of positional mutations in of beta lactamase TEM in CTX antibiotic Mira *et al.***

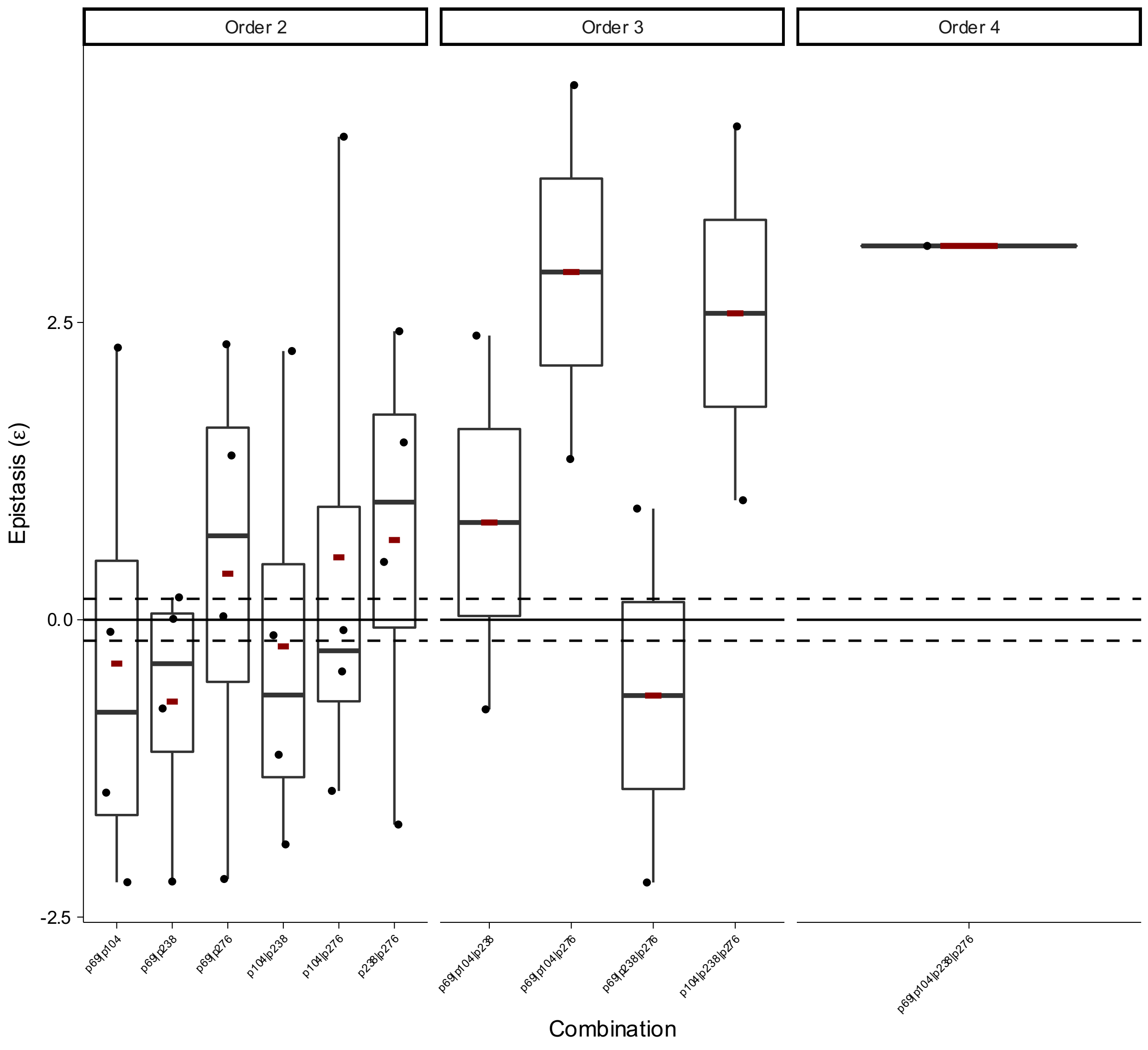


**Box Plot 74 | Epistasis of combinatorial mutations in beta lactamase TEM in CTX antibiotic Mira *et al.***

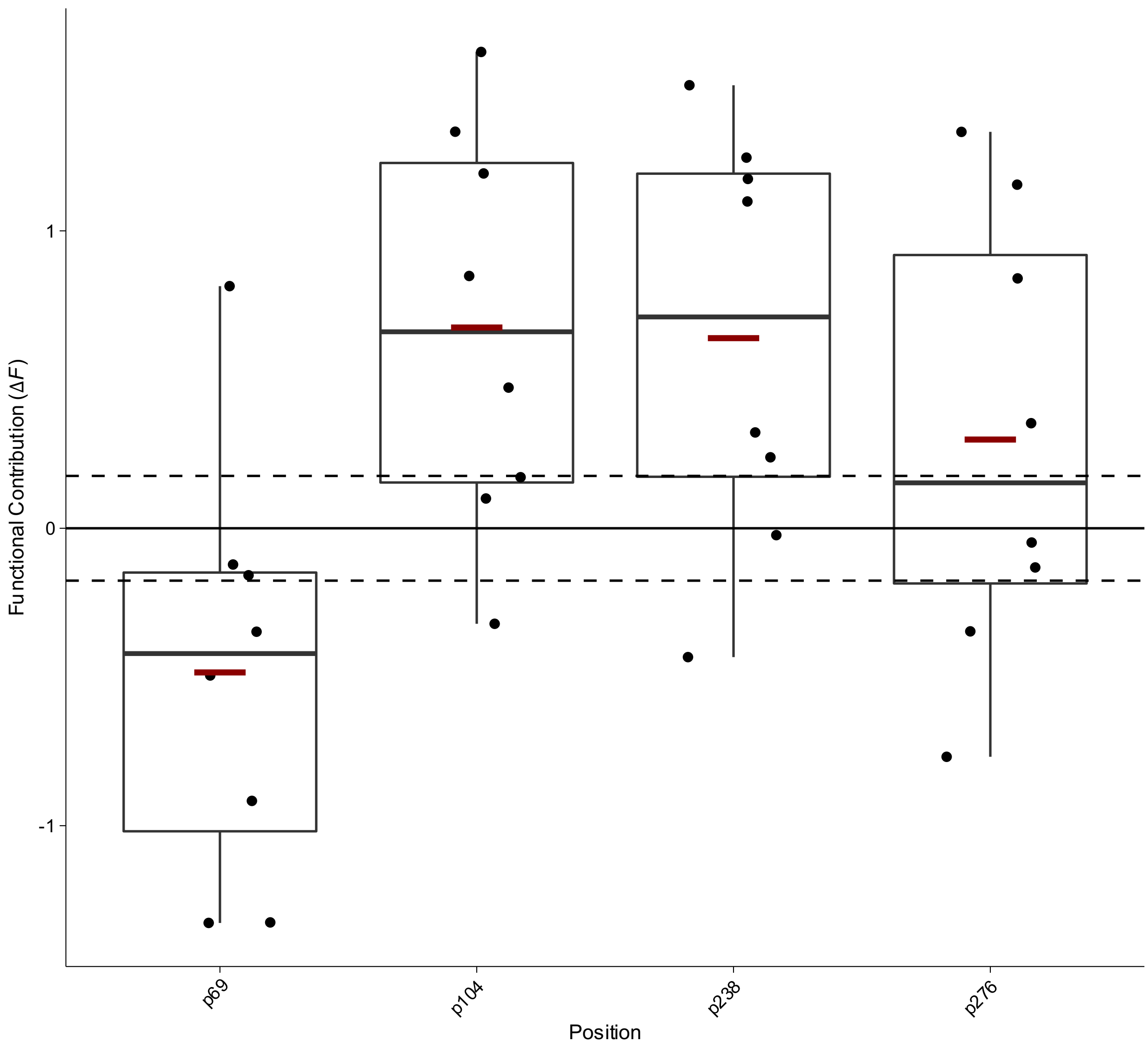


**Box Plot 75 | Functional contribution of positional mutations in of beta lactamase TEM in CXM antibiotic Mira *et al.***

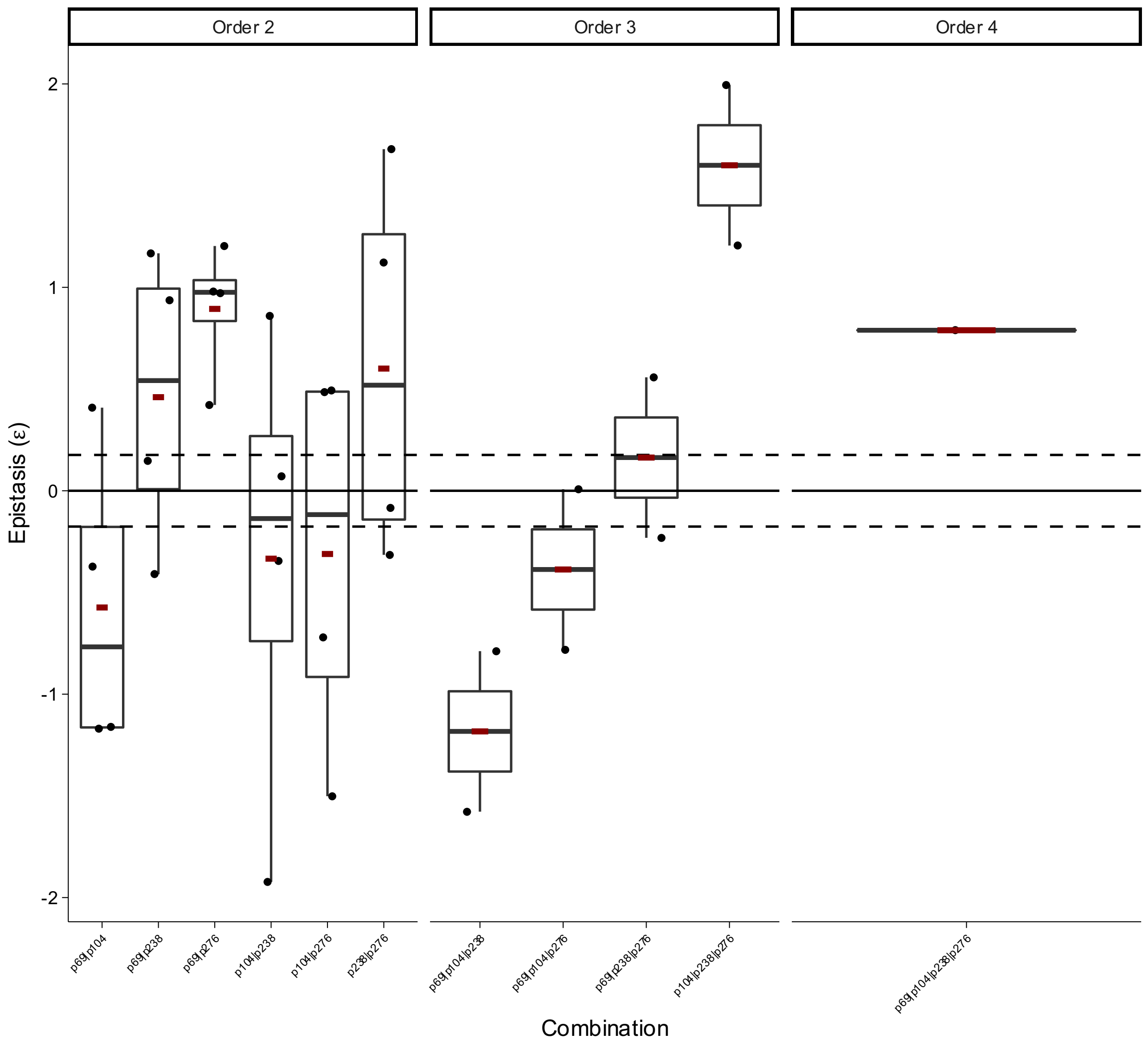


**Box Plot 76 | Epistasis of combinatorial mutations in beta lactamase TEM in CXM antibiotic Mira *et al.***

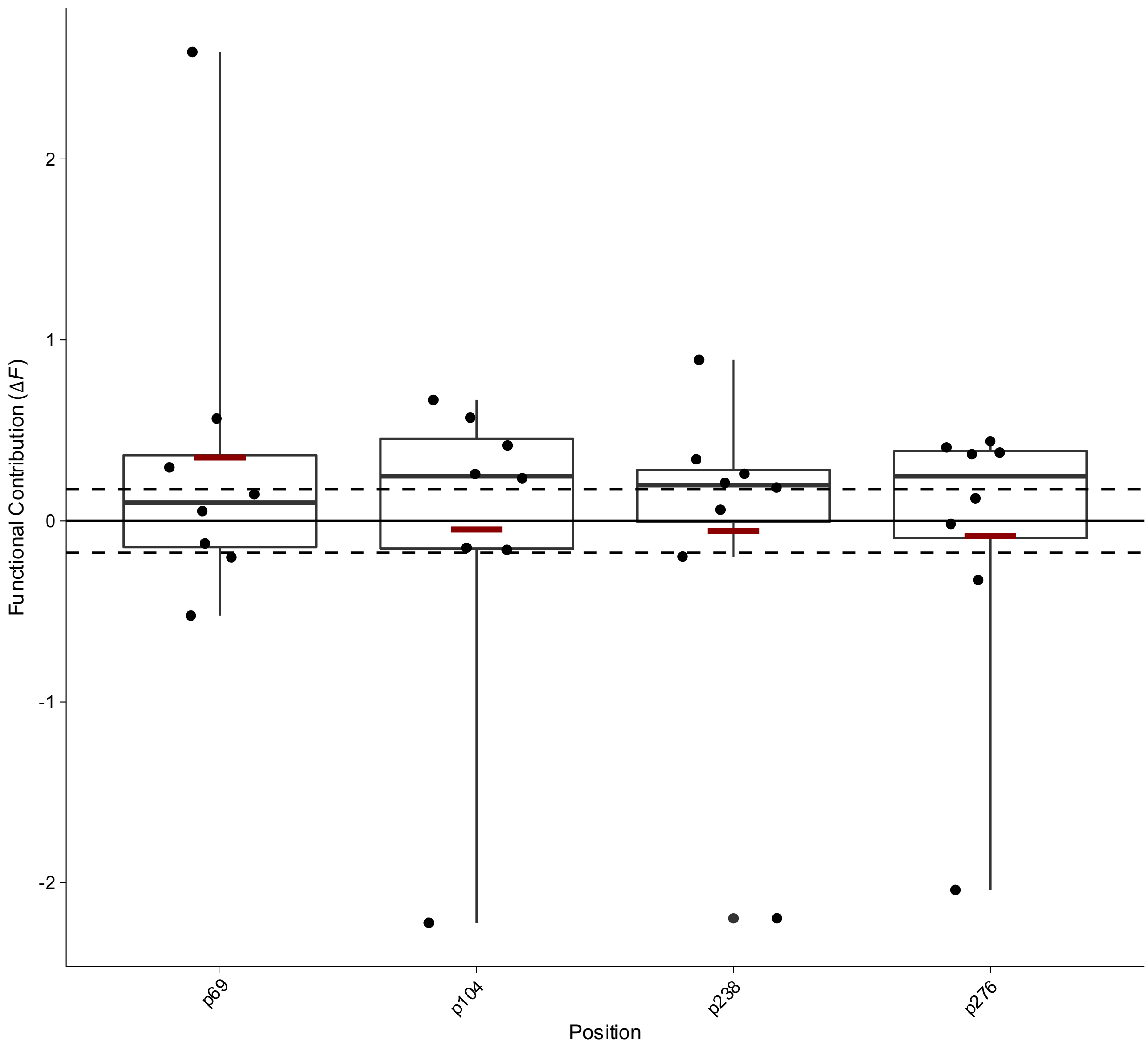


**Box Plot 77 | Functional contribution of positional mutations in of beta lactamase TEM in FEP antibiotic Mira *et al.***

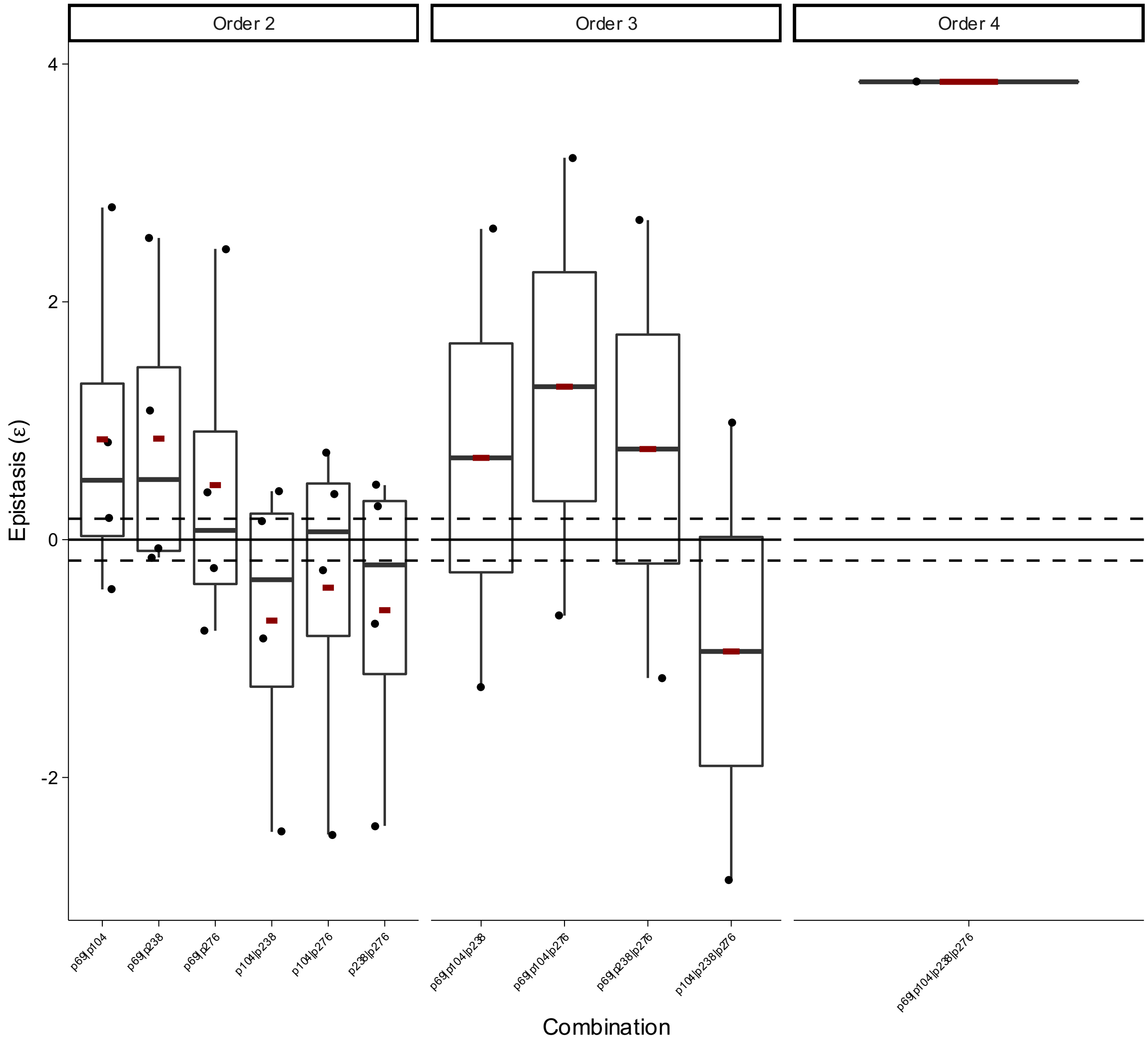


**Box Plot 78 | Epistasis of combinatorial mutations in beta lactamase TEM in FEP antibiotic Mira *et al.***

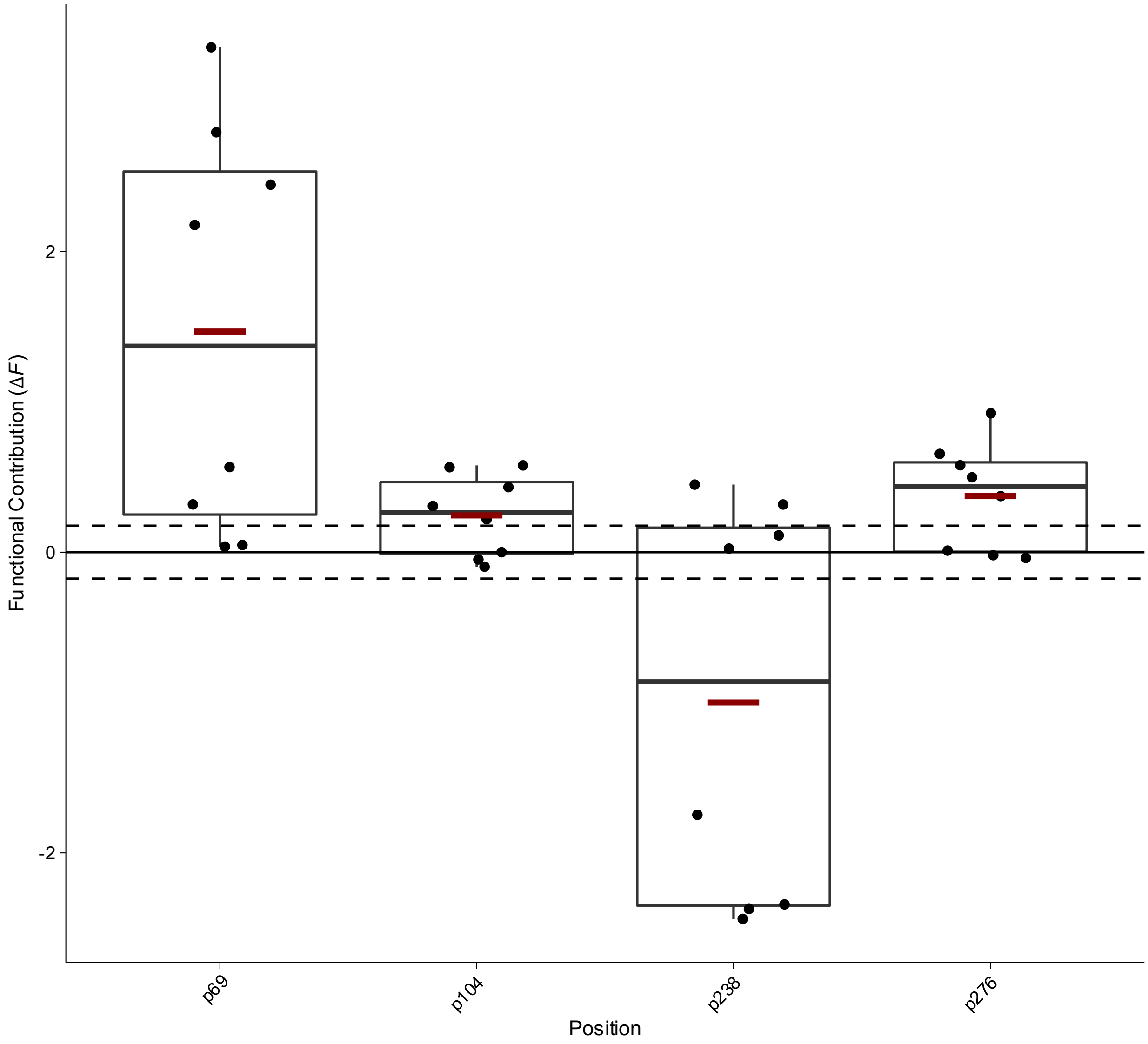


**Box Plot 79 | Functional contribution of positional mutations in of beta lactamase TEM in SAM antibiotic Mira *et al.***

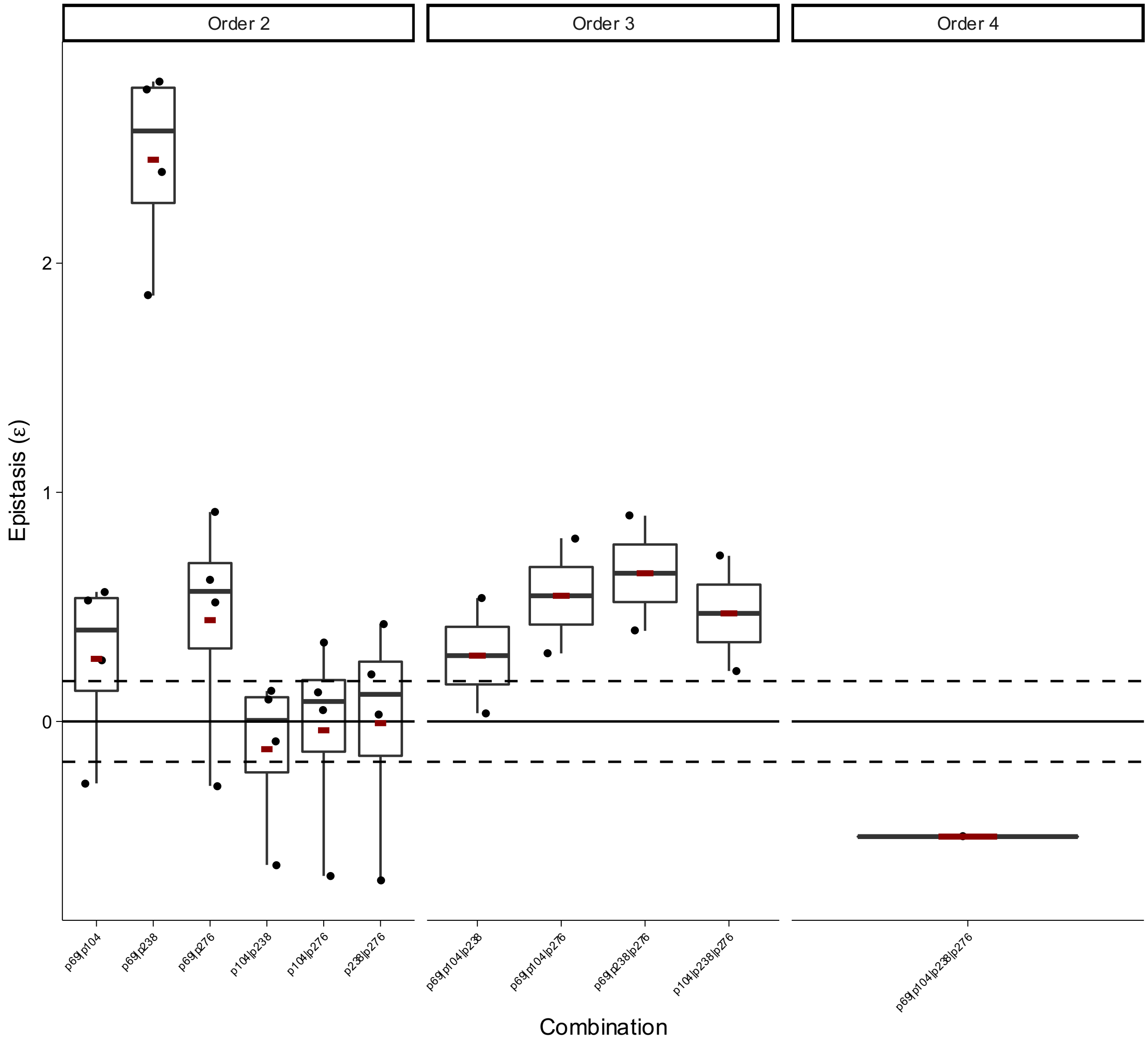


**Box Plot 80 | Epistasis of combinatorial mutations in beta lactamase TEM in SAM antibiotic Mira *et al.***

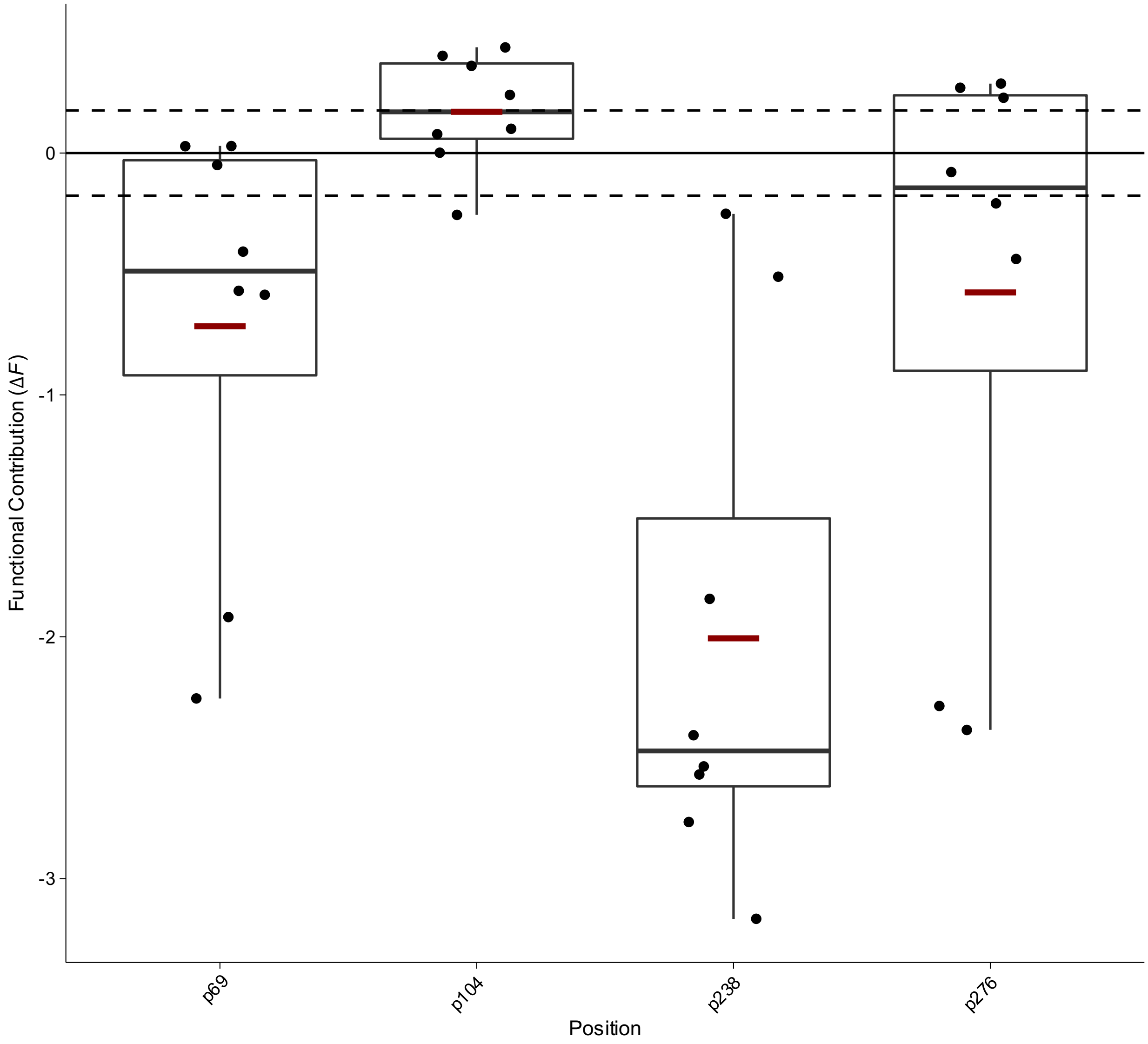


**Box Plot 81 | Functional contribution of positional mutations in of beta lactamase TEM in TZT antibiotic Mira *et al.***

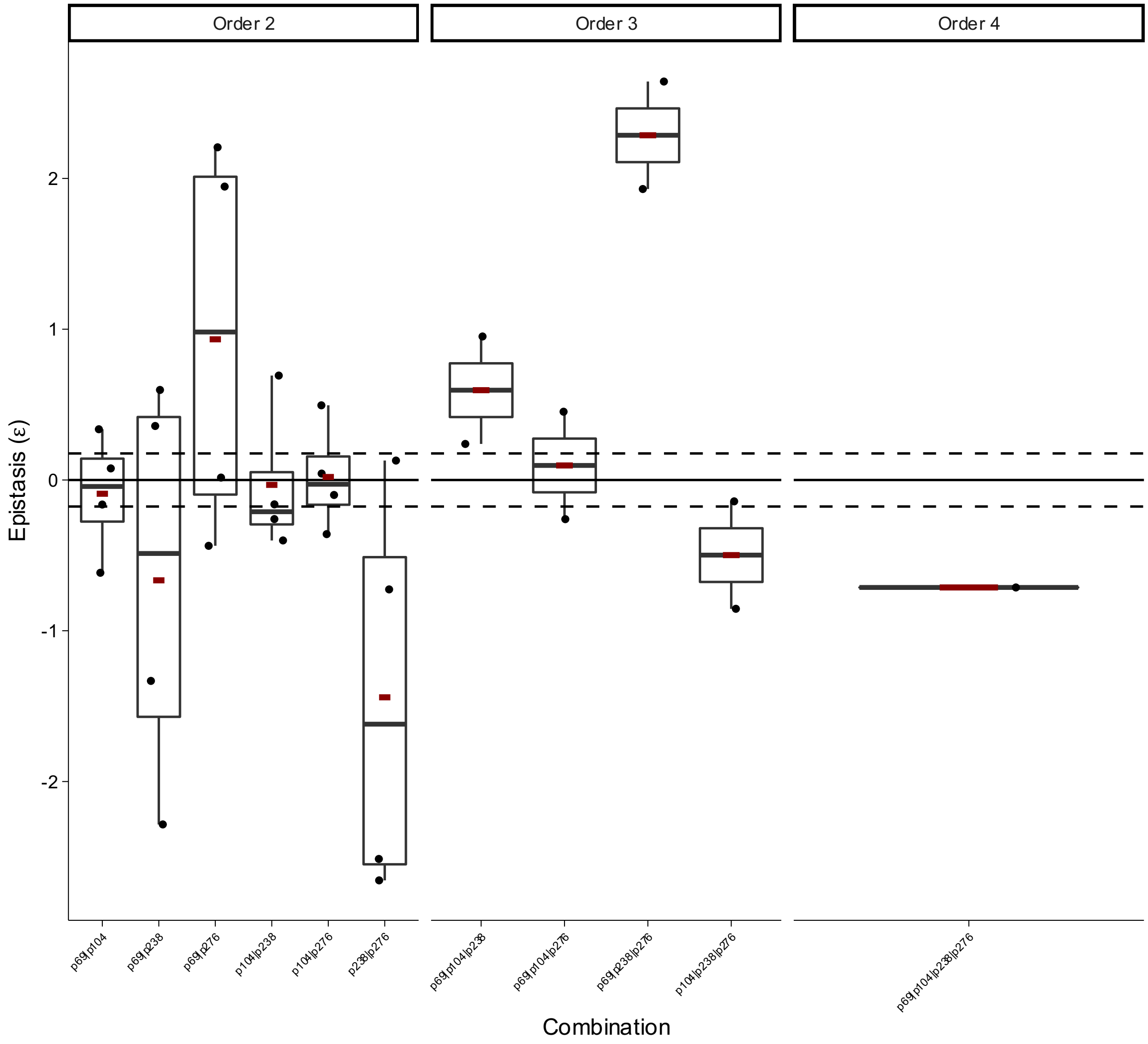


**Box Plot 82 | Epistasis of combinatorial mutations in beta lactamase TEM in TZP antibiotic Mira *et al.***

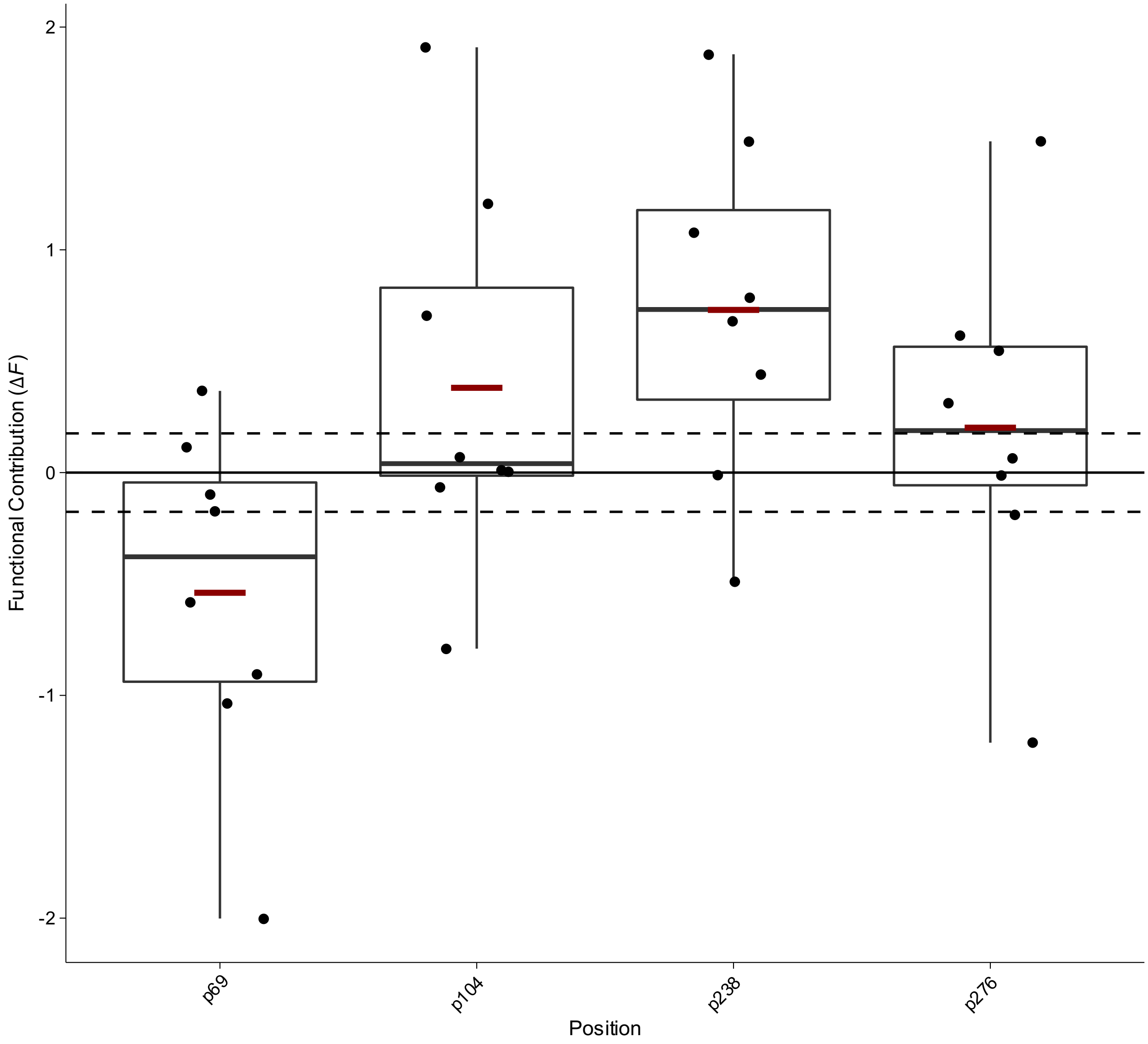


**Box Plot 83 | Functional contribution of positional mutations in beta lactamase TEM in ZOX antibiotic Mira *et al.***

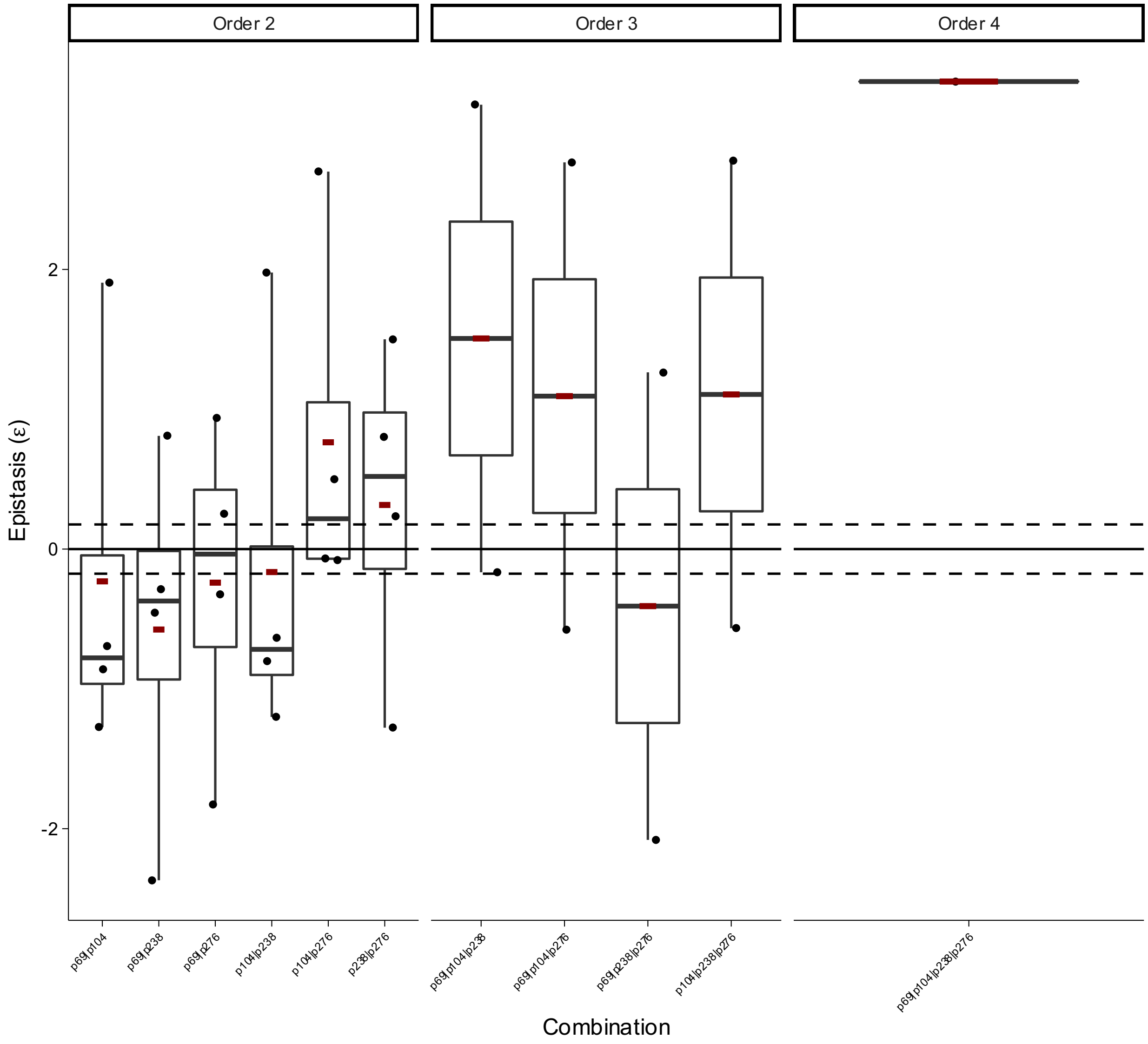


**Box Plot 84 | Epistasis of combinatorial mutations in beta lactamase TEM in ZOX antibiotic Mira *et al.***

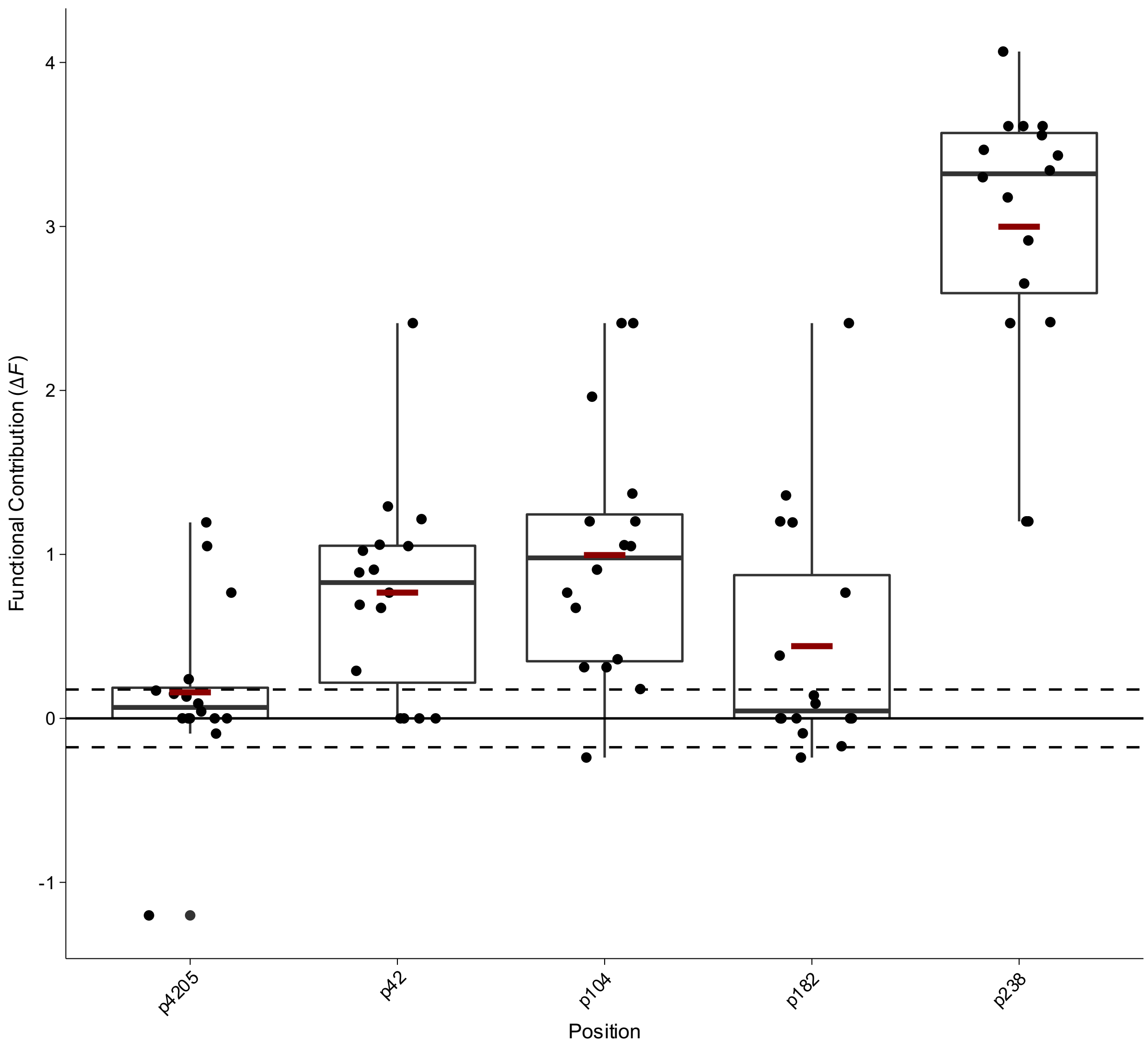


**Box Plot 85 | Functional contribution of positional mutations in beta lactamase TEM from Weinreich *et al.***

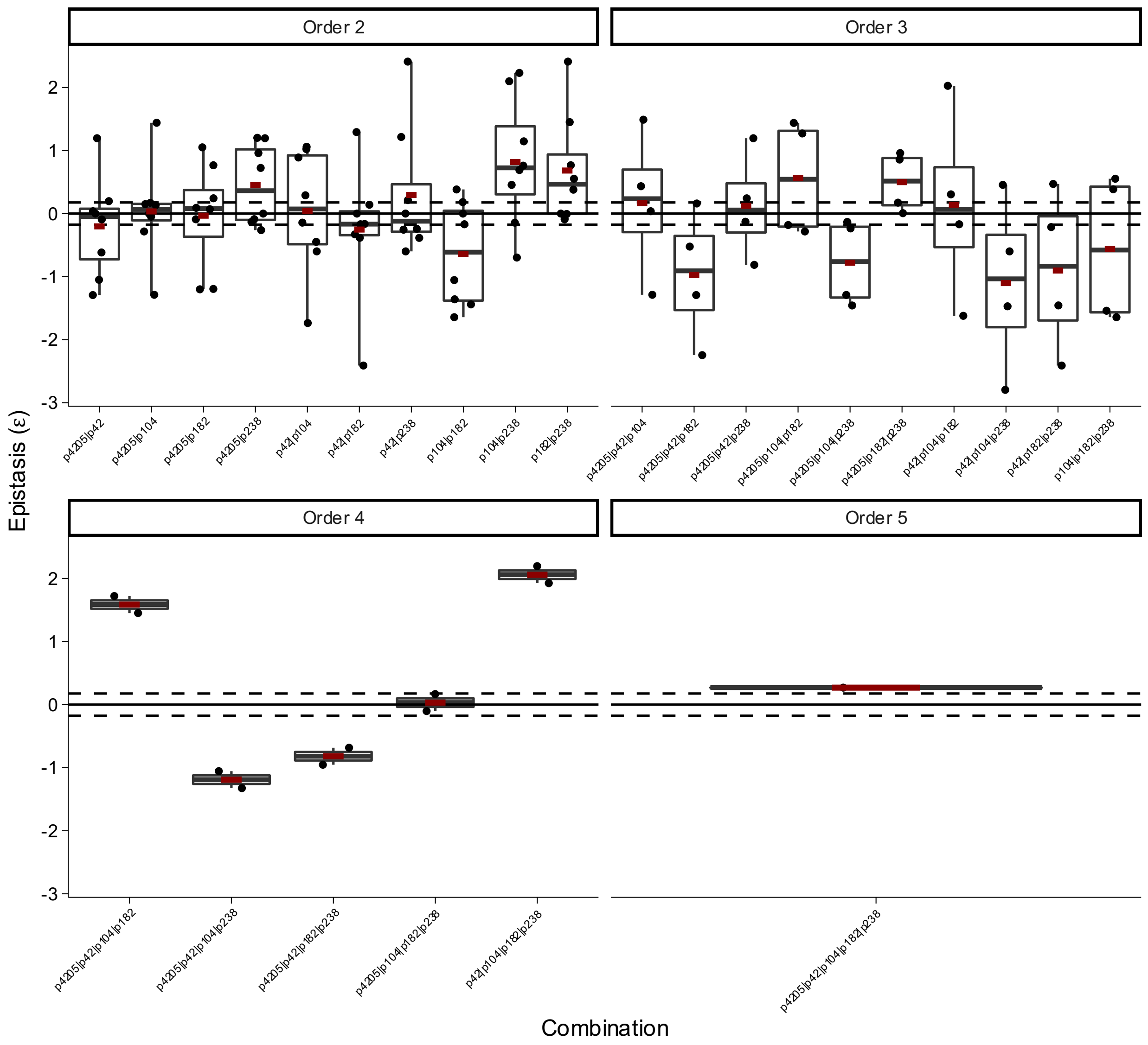


**Box Plot 86 | Epistasis of combinatorial mutations in beta lactamase TEM from Weinreich *et al.***

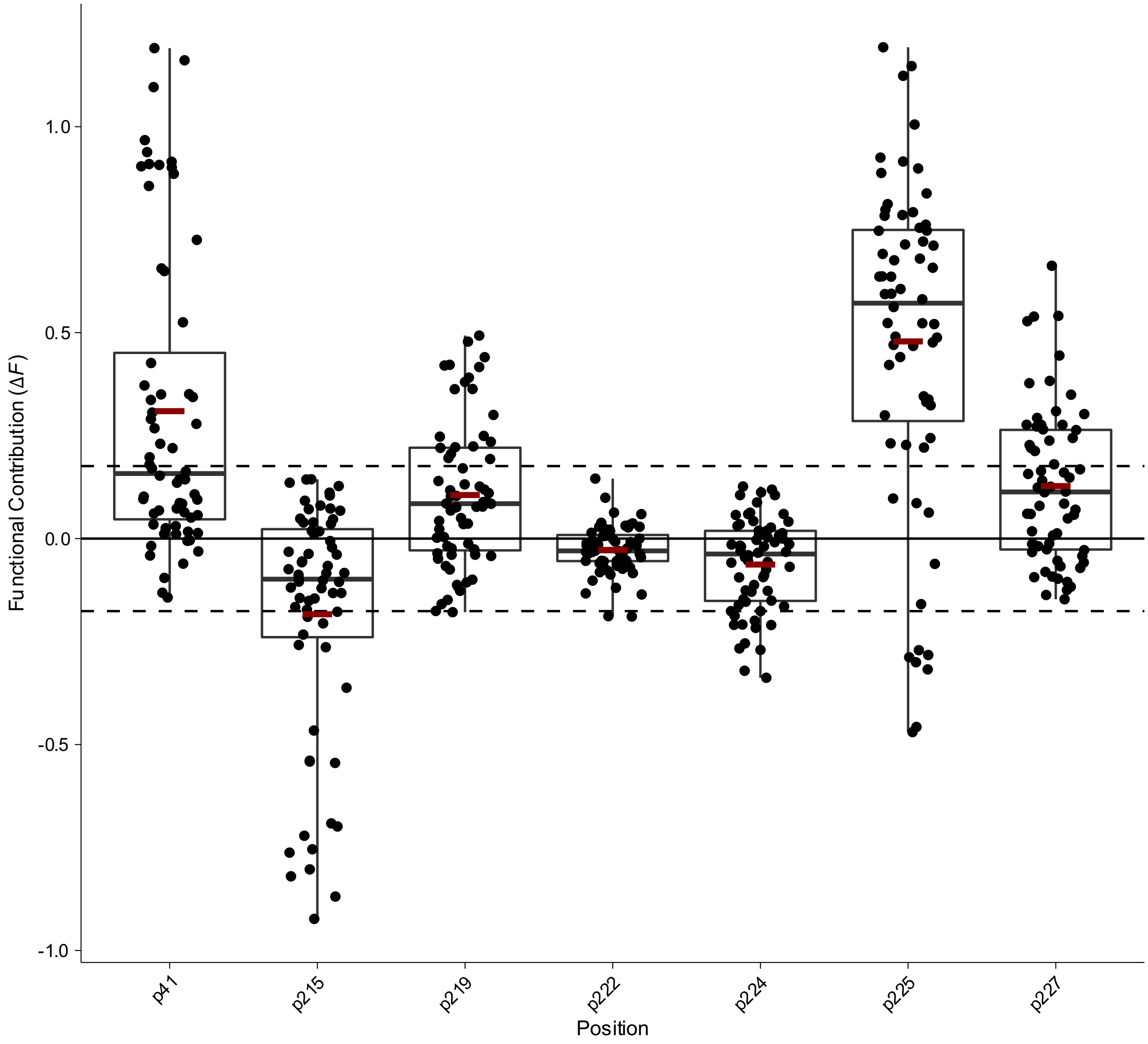


**Box Plot 87 | Functional contribution of positional mutations in nitroreductase NfsA in the 20_39 trajectory from Hall *et al.***

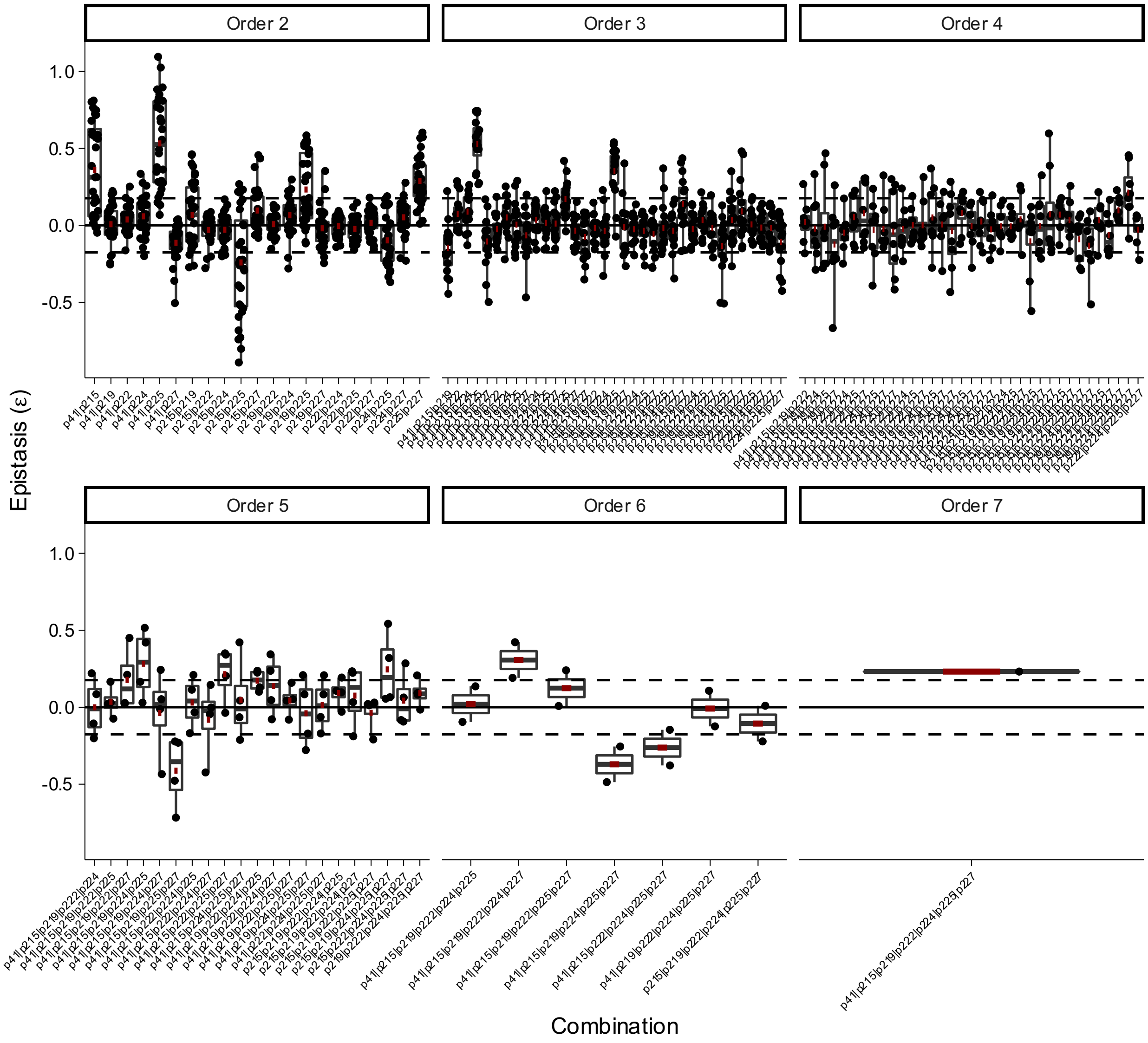


**Box Plot 88 | Epistasis of combinatorial mutations in nitroreductase NfsA in the 20_39 trajectory from Hall *et al.***

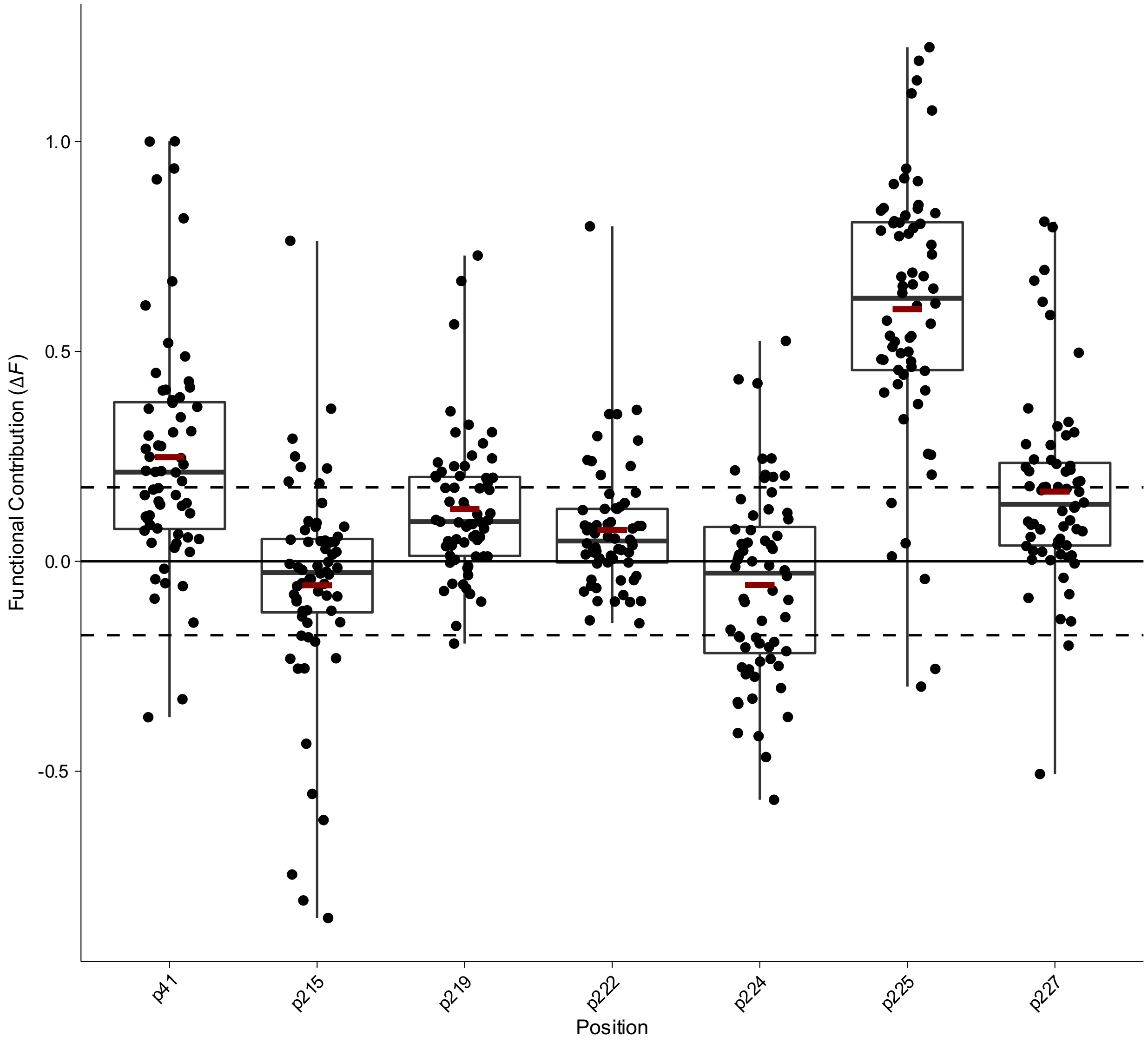


**Box Plot 89 | Functional contribution of positional mutations in nitroreductase NfsA in the 36_37 trajectory from Hall *et al.***

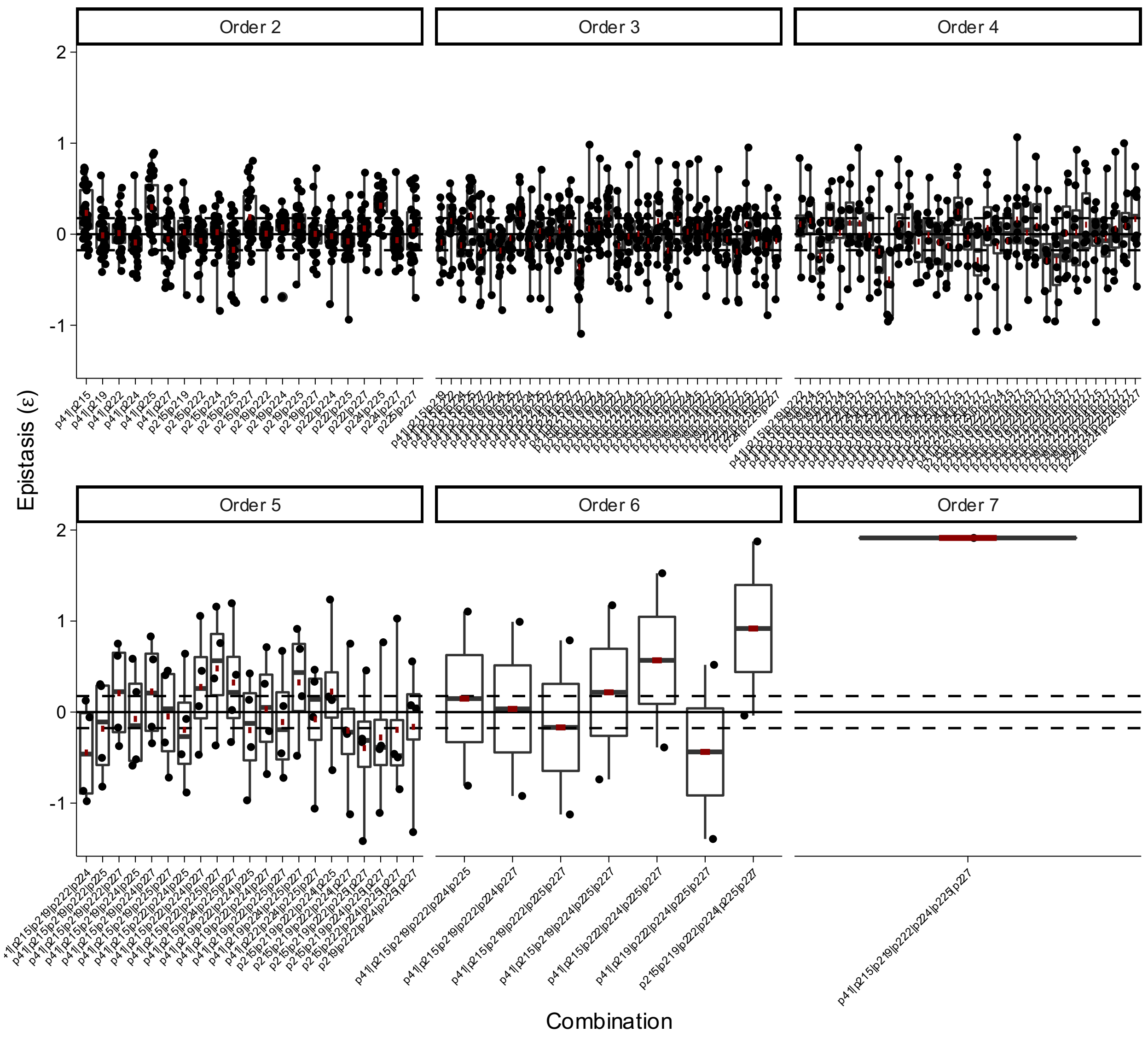


**Box Plot 90 | Epistasis of combinatorial mutations in nitroreductase NfsA in the 36_37 trajectory from Hall *et al.***
