## Supplementary File 3 for "Higher-order epistasis creates idiosyncrasy, confounding predictions in protein evolution"

### Supplementary information for the non-linear transformation

#### Part 1: Simulation

We simulated a genotype-phenotype map with 4 mutated positions using the following script.

```
set.seed(123) # Constant Seed

# Create a library of 4 mutants
sim_df = as.data.frame(gtools::permutations(2, 4, c(-1, 1), repeats = TRUE))

colnames(sim_df) = c("p1", "p2", "p3", "p4")

geno_tab = gtools::permutations(2, 4, c(-1, 1), repeats = TRUE)
geno_tab[which(geno_tab == -1)] = 2

geno_tab = matrix(c("Y", "X")[geno_tab], nrow = 16, ncol = 4)

geno_tab = apply(geno_tab, 1, function(x) paste(x, collapse = ""))

### MAKE SURE GENOTYPES MATCH ORDER OF EFFECT COLUMNS

# Defined epistatic coefficents

pos1 = 2
pos2 = 4
pos3 = 0.5
pos4 = 1
pos1_pos2 = 1
pos1_pos3 = 1
pos1_pos4 = 5
pos2_pos3 = 1
pos2_pos4 = 0.5
pos3_pos4 = 1
pos1_pos2_pos3 = 1
pos1_pos2_pos4 = 3
pos1_pos3_pos4 = 1
pos2_pos3_pos4 = 1
pos1_pos2_pos3_pos4 = 1

# Define positional combinations

sim_df$p1_p2 = sim_df$p1 == 1 & sim_df$p2 == 1
sim_df$p1_p3 = sim_df$p1 == 1 & sim_df$p3 == 1
sim_df$p1_p4 = sim_df$p1 == 1 & sim_df$p4 == 1
sim_df$p2_p3 = sim_df$p2 == 1 & sim_df$p3 == 1
```

```
sim_df$p2_p4 = sim_df$p2 == 1 & sim_df$p4 == 1
sim_df$p3_p4 = sim_df$p3 == 1 & sim_df$p4 == 1

sim_df$p1_p2_p3 = sim_df$p1 == 1 & sim_df$p2 == 1 & sim_df$p3 == 1
sim_df$p1_p2_p4 = sim_df$p1 == 1 & sim_df$p2 == 1 & sim_df$p4 == 1
sim_df$p1_p3_p4 = sim_df$p1 == 1 & sim_df$p3 == 1 & sim_df$p4 == 1
sim_df$p2_p3_p4 = sim_df$p2 == 1 & sim_df$p3 == 1 & sim_df$p4 == 1

sim_df$p1_p2_p3_p4 = sim_df$p1 == 1 & sim_df$p2 == 1 & sim_df$p3 == 1 &
sim_df$p4 == 1

# Simulate effects based on a starting genotype (start)

effect = c()
for(i in 1:dim(sim_df)[1]){
  start = 10
  if(sim_df[i,1] == 1) {
    start = start*pos1
  }
  if(sim_df[i,2] == 1) {
    start = start*pos2
  }
  if(sim_df[i,3] == 1) {
    start = start*pos3
  }
  if(sim_df[i,4] == 1) {
    start = start*pos4
  }
  if(sim_df[i,5] == 1) {
    start = start*pos1_pos2
  }
  if(sim_df[i,6] == 1) {
    start = start*pos1_pos3
  }
  if(sim_df[i,7] == 1) {
    start = start*pos1_pos4
  }
  if(sim_df[i,8] == 1) {
    start = start*pos2_pos3
  }
  if(sim_df[i,9] == 1) {
    start = start*pos2_pos4
  }
  if(sim_df[i,10] == 1) {
    start = start*pos3_pos4
  }
  if(sim_df[i,11] == 1) {
    start = start*pos1_pos2_pos3
  }
```

```
  if(sim_df[i,12] == 1) {
    start = start*pos1_pos2_pos4
  }
  if(sim_df[i,13] == 1) {
    start = start*pos1_pos3_pos4
  }
  if(sim_df[i,14] == 1) {
    start = start*pos2_pos3_pos4
  }
  if(sim_df[i,15] == 1) {
    start = start*pos1_pos2_pos3_pos4
  }
  effect = c(effect, start)
}

# Log transform the effects

sim_df_effect = log10(effect)

## Formatting and output to mimic datasets used in epistatic analysis code

out_df = data.frame("first" = c("WT", "XXXX", "Positions", 1, "Genotype",
rep(geno_tab, each = 1)), "second" = c(rep(NA, 3), 2, "Effect",
sim_df_effect),
                    "third" = c(rep(NA, 3), 3, rep(NA, length(sim_df_effect)
+ 1)), "fourth" = c(rep(NA, 3), 4, rep(NA, length(sim_df_effect) + 1)))


## Exporting

write.table(out_df, "sim.csv", row.names = F, col.names = F, na = "", sep =
",")
```

**Part 2: Data Processing of Simulation**

We tested the non-linear transformation technique[1] from Sailer & Harms on non-idiosyncratic, linear (additive), epistatic data generated by the code above.

Coefficients were as follows:

- p1: 2-fold
- p2: 4-fold
- p3: 0.5-fold
- p4: 1-fold
- p1p4: 5-fold
- p2p4: 0.5-fold
- p1p2p4: 3-fold

First, we imported the csv file generated by the simulation script

```
d = read_csv("sim.csv",
             show_col_types = F)

d = d[5:dim(d)[1], 1:2]
colnames(d) = c("Genotype", "Phenotype")
d$Phenotype = as.numeric(d$Phenotype)

head(d)

## # A tibble: 6 × 2
##   Genotype Phenotype
##   <chr>        <dbl>
## 1 XXXX         1
## 2 XXXY         1
## 3 XXYX         0.699
## 4 XXYY         0.699
## 5 XYXX         1.60
## 6 XYXY         1.30
```

Then we processed the genotypes to 0 and 1 coefficients instead of X's and Y's

```
# Split Genotype into 0s and 1s to take the mean

geno = do.call("rbind", str_split(d$Genotype, ""))
geno[geno == "X"] = 0
geno[geno == "Y"] = 1

geno = as_tibble(geno)
colnames(geno) = c("p1", "p2", "p3", "p4")
geno$p1 = as.numeric(geno$p1)
geno$p2 = as.numeric(geno$p2)
geno$p3 = as.numeric(geno$p3)
geno$p4 = as.numeric(geno$p4)

geno$Phenotype = d$Phenotype

head(geno)

## # A tibble: 6 × 5
##      p1    p2    p3    p4 Phenotype
##   <dbl> <dbl> <dbl> <dbl>     <dbl>
## 1     0     0     0     0     1
## 2     0     0     0     1     1
## 3     0     0     1     0     0.699
## 4     0     0     1     1     0.699
## 5     0     1     0     0     1.60
## 6     0     1     0     1     1.30
```

Got the average functional contribution of each position (p1-4) and used an additive model to predict function.

```r
## Average of each position
dif = function(x){return(x[2]-x[1])}
params = c()

# Loop ever position and get the parameter for its average effect

for(pos in 1:4) {
  params = c(params, dif(geno %>%
  group_by_at(pos) %>%
  summarise(mean_pheno = mean(Phenotype)) %>%
  pull(mean_pheno)))
}

# Predict

pred_pheno = c()
for(row in 1:dim(geno)[1]) {
  pred_pheno = c(pred_pheno, geno$Phenotype[1] + sum(geno[row,1:4] * params))
}

# Output is a vector of pred_pheno

geno$Predicted = pred_pheno

# Plot of Padd vs Pobs

plot(Phenotype ~ Predicted, geno, pch = 16)
abline(0, 1)
```

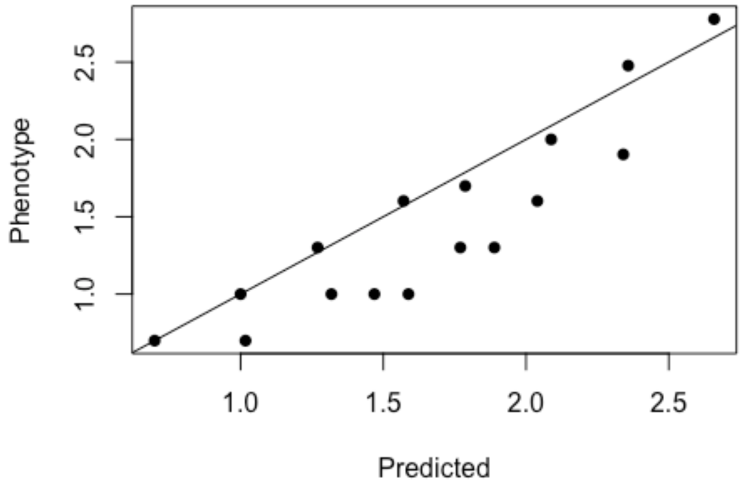


We see apparent epistasis skewing the average functional effect prediction on the landscape. Keep in mind that Predicted here is the same as predictions stemming from the 1st order coefficients of a linear model.

**Part 3: Spline Transform**

What is the current $R^2$ based on the plot we see above?

Currently it is 67.6%

We can try to remove the non-linearity with a spline transform by looping from 1 to 10 knots and determining best fit by AIC

```
library(splines)

aics = c()
for(knot in 1:10) {
  sm = lm(Predicted ~ bs(Phenotype, knots = knot), data = geno)
  aics = c(aics, AIC(sm))
}
```

Best performing model is one with 3 knots

```
sm = lm(Predicted ~ bs(Phenotype, knots = which.min(aics)), data = geno)

plot(Predicted ~ Phenotype, geno, xlab = "Phenotype", ylab = "Predicted", pch
= 16)
nlt <- seq(min(geno$Predicted), max(geno$Predicted), length.out = 200)
lines(nlt, predict(sm, data.frame(Phenotype = nlt)))
```

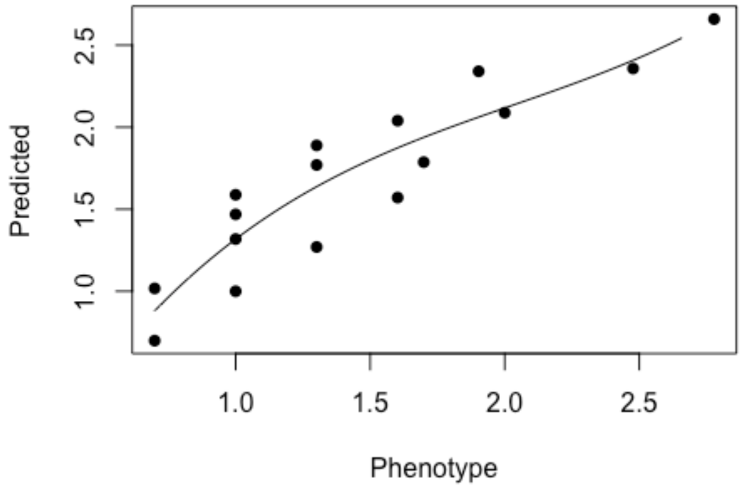


So how does this transformation look like?

```
plot(predict(sm) ~ geno$Predicted, xlab = "Predicted", ylab = "Linearized
Phenotype", pch = 16)
abline(0, 1)
```

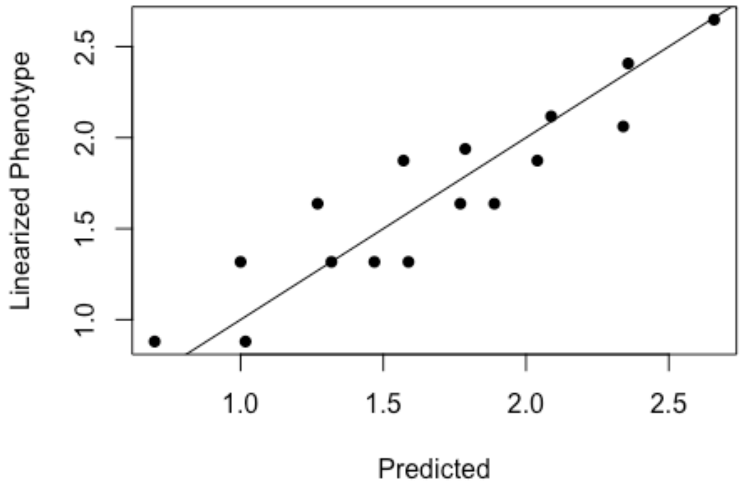


Has the fit improved according to the $R^2$?

Now it is 81.7%

And how much have we transformed the phenotype? Let’s look at the from original to non-linear transformed

```
plot(predict(sm) - geno$Phenotype, xlab = "Index", ylab =
expression(Delta*"Phenotype"), pch = 16)
abline(0, 0)
abline(log10(1.5), 0, lty = 2)
abline(log10(1/1.5), 0, lty = 2)
```

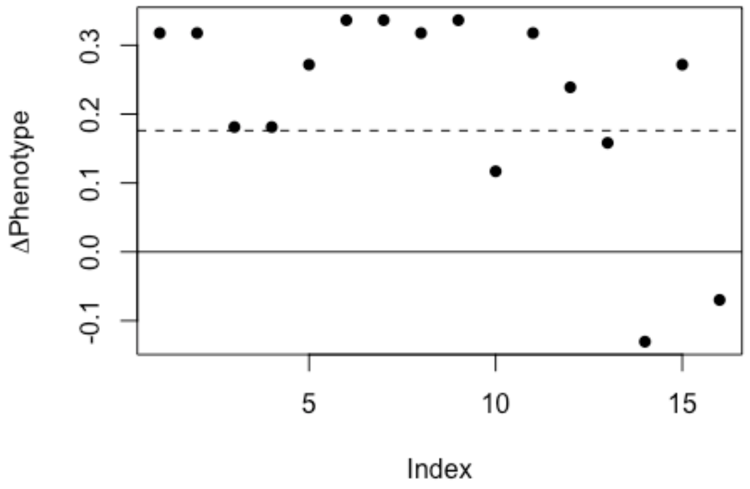


This is a large transformation beyond the 1.5-fold error (shown as dashed line) that is discussed in the main text.

We can also visualize this by plotting the previous pre-linearized phenotype in red.

```
plot(geno$Phenotype ~ geno$Predicted, xlab = "Predicted", ylab = "Linearized
Phenotype", pch = 16, col = "red")
points(predict(sm) ~ geno$Predicted, pch = 16)
abline(0, 1)
```

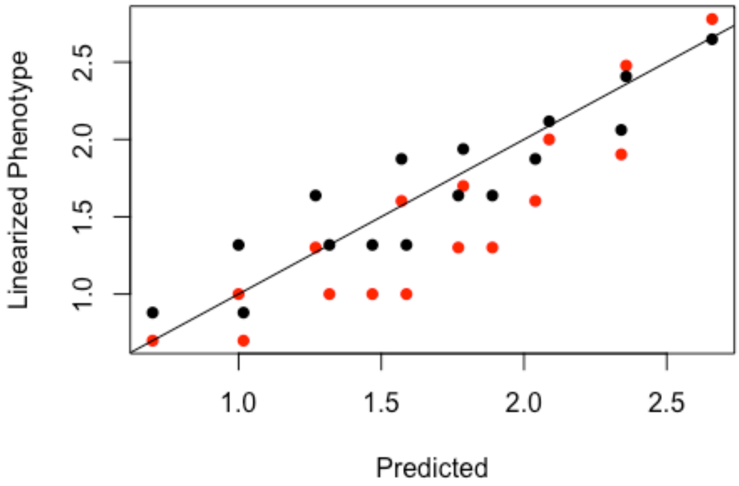


Since we introduced no inherent non-linearity into the genotype-phenotype map, the non-linear transform is flattening the “true” epistasis we simulated.

Although appreciate the value in applying the non-linear transform to various data in order to reduce the effect of various non-linear properties on the system as much as possible, we chose not to use it for our data in order not to skew or minimize the inferred epistasis.
